# Cation-Cation Photosensitization for Protein Ligation and Intracellular Catalysis

**DOI:** 10.1101/2025.11.03.686045

**Authors:** Vishal Agarwal, Hieu Pham, Samuel G. Bartko, Michael T. Taylor

## Abstract

We report here a photosensitized strategy for protein labelling in which N-substituted pyridinium salts are activated using a 2,4-diaryl-N-methyl quinolinium scaffold. Structure-reactivity relationships were performed to optimize the sensitizer structure, and ultimately generated a system that gives protein labelling in minutes at micromolar reagent concentrations. Mechanistic studies suggest a photo-induced electron transfer-based sensitization mechanism. The mildness of this system enabled us to assay sensitization both on individual biomolecules and in complex proteomes and demonstrated excellent compatibility with lysate- and the live-cell-based systems. Imaging of photolabelled HeLa cells were performed and revealed that catalysis occurs in multiple cellular compartments. Chemical proteomics performed at the lysate level resulted in the enrichment of 319 proteins with a 93% selectivity to Tryptophan residues. Live cell labelling resulted in 101 enriched proteins, primarily from the nucleus.

---

Photocatalytic technologies are established tools for photodynamic therapy^1^ and are rapidly emerging tools for biomolecule conjugate synthesis^2^, bioorthogonal chemistry^3^, and biochemical discovery^4^. As tools for protein ligation, Kodadek showed that photocatalytic cross-linking of proteins was possible using palladium and Ruthenium photocatalysts^4a,b^. Later, Nakamura showed that Ru(bpy)_3_-based catalysts could be targeted to individual recombinant proteins and label these selectively^4c^. More recently Macmillan and Rovis have shown that Ir- and Os-photoredox catalysts can enable cell-surface proximity labelling^4d,e^. Since these reports, multiple reports have shown that multiple catalysts structures, including classical organic dyes and enzymes, can be targeted for labelling in extra- and intra-cellular environments^4f-l^. Whilst elegant, these approaches are reliant on comparatively few reactive intermediates: nitrenes, carbenes, aryloxy radicals, and ^1^O_2_. New catalytic approaches that provide complementary labelling outcomes could enhance the scope of applications.

We have developed an approach towards protein ligation that harnesses photo-induced electron transfer (PET) by pairing N-substituted pyridinium salts with Tryptophan (Trp)-containing proteins to achieve Trp-selective ligation^5^. This process is biocompatible and enabled intracellular Trp-selective proteome profiling for the first time. During the course of our studies, we noticed a phenomenon in which previously reported pyridinium salt **1**, when selectively excited using 427 nm LED in the presence of equimolar **2**, which is not activated by this lightsource, could sensitize labelling with **2** (**Figure 1A**). **1** and **2** have similar calculated LUMO energies^6^, leading us to postulate that electron transfer (ET) between **1•** (generated by reductive quenching of [**1**]^*^) and **2** could occur rapidly and reversibly, leading to fragmentation of the resulting **2•** and subsequent labelling (**Figure 1B**). Given that pyridinium compounds structurally related to **1** and **2** rapidly fragment^5,7^, electron transfer between these species must happen with exceptional rapidity and thus prompted us to search for a catalytic variant of **2** that could harness these rapid kinetics. In 2015, Studer elegantly showed that simple N-carbamoyl pyridinium salts could be activated photocatalytically using Ru(bpy)_3_ for heterocycle functionalization under small molecule synthesis conditions^8^; but these conditions were not amenable to protein ligation (see SI Section 4). Screening several sensitizer classes led us to identify 1,3-diarylquinolinium salts as potential catalysts (see SI Section 4). These moieties appeared ideal for photo-catalysis as (1) related analogs have demonstrated biocompatibility^9^ and (2) they provide a simple organic framework for tuning in specific photophysical properties.

**Figure 1.**
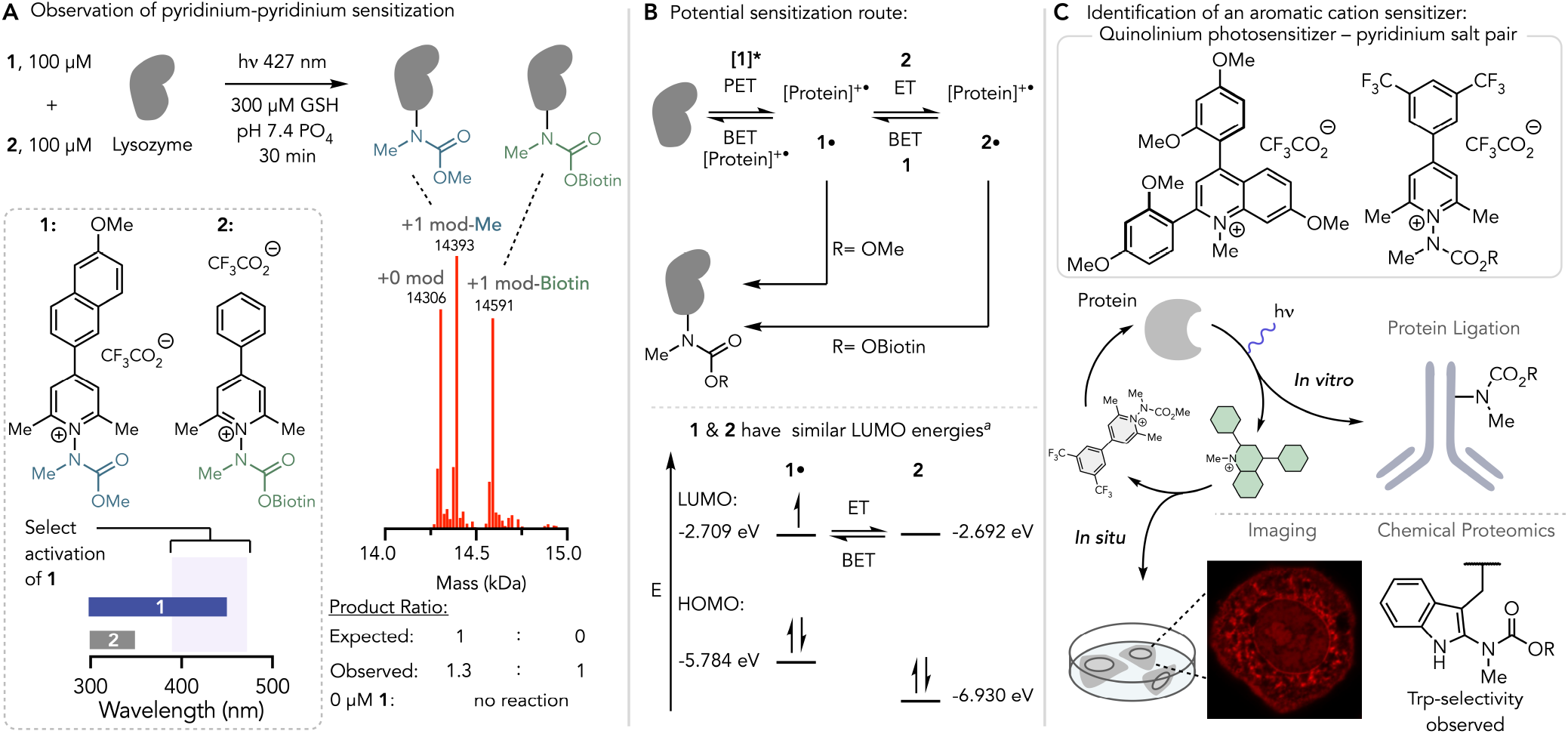
(**A**) Observation of Pyridinium-pyridinium photosensitization. (**B**) Potential mechanism of sensitization. BET=back electron transfer. (**C**) Identification and optimization of a diarylquinolinium aromatic cation sensitizer. ^*a*^Calculated at the UMN12-SX 6-31g(d) SMD=H_2_O level of theory.

We therefore sought to identify an optimal pair of quinolinium sensitizer and stoichiometric pyridinium salt as defined by a system that uses minimal amounts of sensitizer and stoichiometric pyridinium as well as minimal irradiation times. We assembled a small bank of 2,4biaryl-*N*-methyl-quinolinium salts possessing various aromatic groups and functional decorations designed to perturb the electronics of the catalyst. This was facilitated by extending regioselective cross-coupling approaches used by Chenoweth and Petersson^9^ (**Figure 2A**, inset). Our screening assay consisted of irradiation of model protein lysozyme (10 µM), glutathione (GSH) (300 µM), a specific photocatalyst (**3**-**17**, Figure 2A) at a range of concentrations (5-200 µM), and one of pyridinium salts **18**-**20** (100 µM) in 20 mM pH 7.5 Na_2_HPO_4_/NaH_2_PO_4_ for 5 minutes. Reaction mixtures were analyzed by measuring the ratio of labelled:unlabelled protein by intact protein mass spectrometry. Studies were performed this way to both quickly identify photosensitizers that label proteins under kinetically demanding regimes and to provide mechanistic insight.

**Figure 2.**
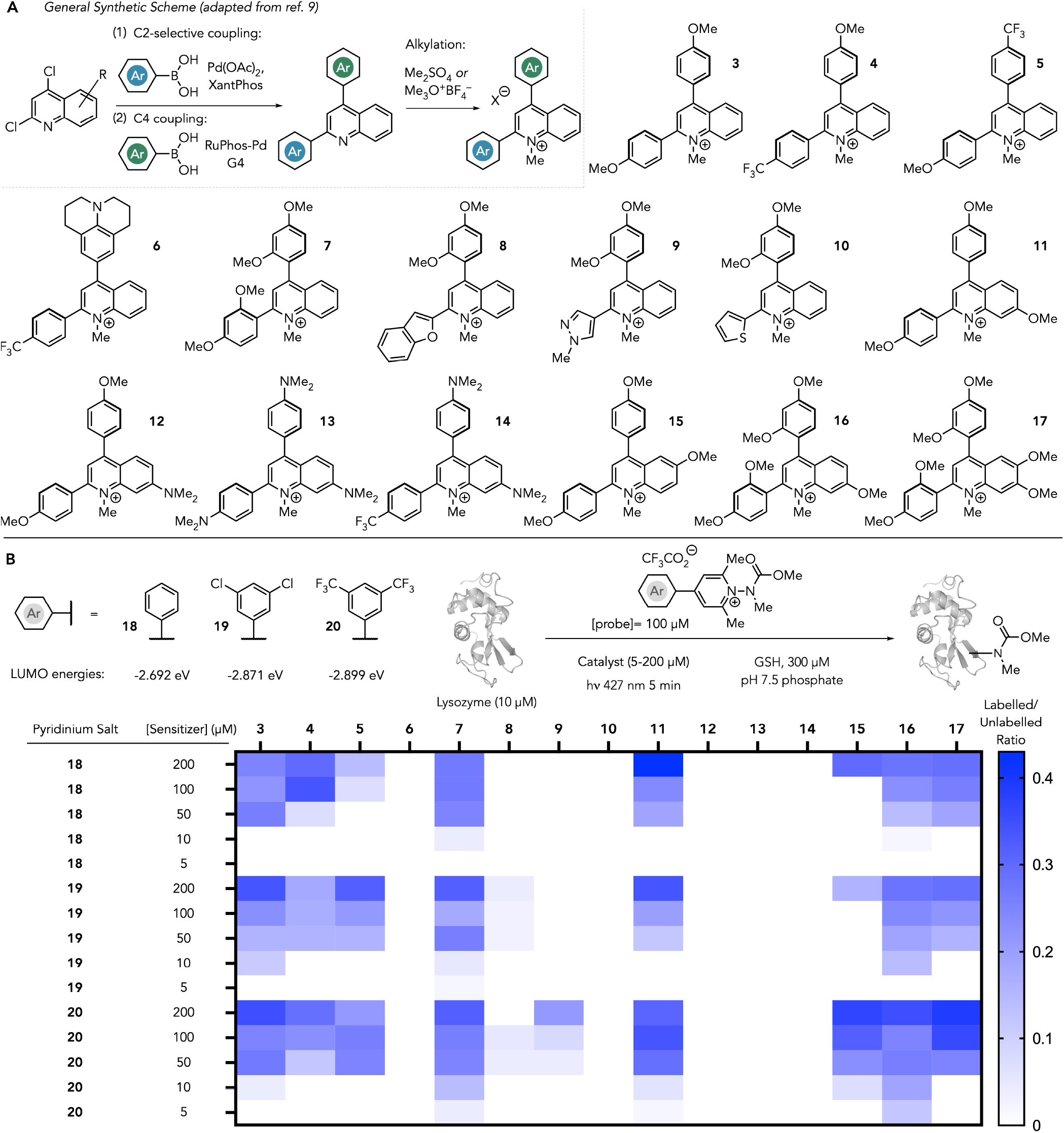
(**A**) 2,4-diaryl-*N*-methyl-quinolinium catalysts synthesized and screened this study. All quinolinium salts possess CF_3_CO^−^ counterions (**B**) Screening of photocatalysts for lysozyme labelling with pyridinium salts **18-20**. Calculated LUMO energies of **18**-**20** provide insight into electronic structure. ^*a*^Calculated at the UMN12-SX 6-31g(d) SMD=H_2_O level of theory.

These efforts revealed clear structure reactivity relationships for sensitized labelling (**Figure 2B**). Structures possessing strong, dialkylamino-electron-donating groups failed to promote labelling (**6, 12**-**14**) under any conditions. By contrast, structures containing moderately donating MeO– groups (**3, 7, 11, 15**-**17**) gave overall the highest levels of labelling. Mixtures of donating and withdrawing groups on the 2– and 4– aromatic substituents gave comparatively less labelling (**4, 5**) and sensitizers possessing heteroaromatic groups (**8**-**10**) gave little to no labelling. Whilst all three stoichiometric pyridinium salts (**18**-**20**) showed labelling activities at higher photosensitizer concentrations (50-200 µM), discernable differences became clearer at lower concentrations (5, 10 µM). Specifically, **18** gave two hits with measurable labelling at 10 µM sensitizer and none at 5 µM sensitizer, **19** gave three hits at 10 µM, and one at 5 µM, whilst **20** gave four hits at 10 µM, and three hits at 5 µM sensitizer. Out of these structures, sensitizers **7, 11**, and **16** gave measurable labelling at 5 and 10 µM concentrations, and the highest levels using stoichiometric salt **20**. Of these, sensitizer **16** gave the most labelling and thus we selected to primarily use the pairing of sensitizer **16** and pyridinium salt **20** for further studies.

Next, we sought to evaluate the mechanistic basis of photosensitization with the **16**/**20** pair. **16** possesses significant absorption out to ∼450 nm (λ_max_=379 nm), a very large stokes shift (164 nm) but a very low fluorescence quantum yield (<0.01) as well as a moderate photooxidation potential ([**16**+]^*^/**16**• of +1.4 eV) (**Figure 3A**). We performed perturbation studies in which we monitor the percentage of unconverted **20** as a function of condition after 20 minutes of irradiation time using LC/MS and an internal standard (**Figure 3B**). Under standard conditions (10 µM lysozyme, 100 µM **20**, 10 µM **16**, 300 µM GSH, 20 mM Na_2_HPO_4_/NaH_2_PO_4_), **20** is almost entirely converted (10% **20** remaining, entry 1), whilst removing the protein from the solution had little effect upon **20** consumption (9% remaining, entry 2). Removal of GSH resulted in lower conversion (92% and 82 % **20** remaining, entries 3 & 4, respectively), and removal of sensitizer, protein, GSH, and buffer results in lower conversions of **20** (entry 5). Incubation of the reaction mixture in the absence of light gave no conversion of **20** and confirms the need for photoactivation of the sensitizer to drive reactivity (entry 6). Substitution of **20** for trimethylpyridinium salt **21**, which has a calculated LUMO energy 581 mV higher than **20** (entry 7) resulted in loss of pyridinium conversion (>95% **21** remaining); suggesting a need to match orbital energies for sensitization. Taken together, we posit that the primary means of photosensitization by **16** is through an electron transfer mechanism in which [**16**]^*^ is reductively quenched by an exogenous electron donor to give **16**^•^ which subsequently undergoes SET with pyridinium salt **20**. [**20**]^•^ then undergoes rapid and irreversible fragmentation to yield a short-lived carbamoyl radical^10^ that reacts with aromatic nucleophiles to yield carbamylated proteins (**Figure 3C**).

**Figure 3.**
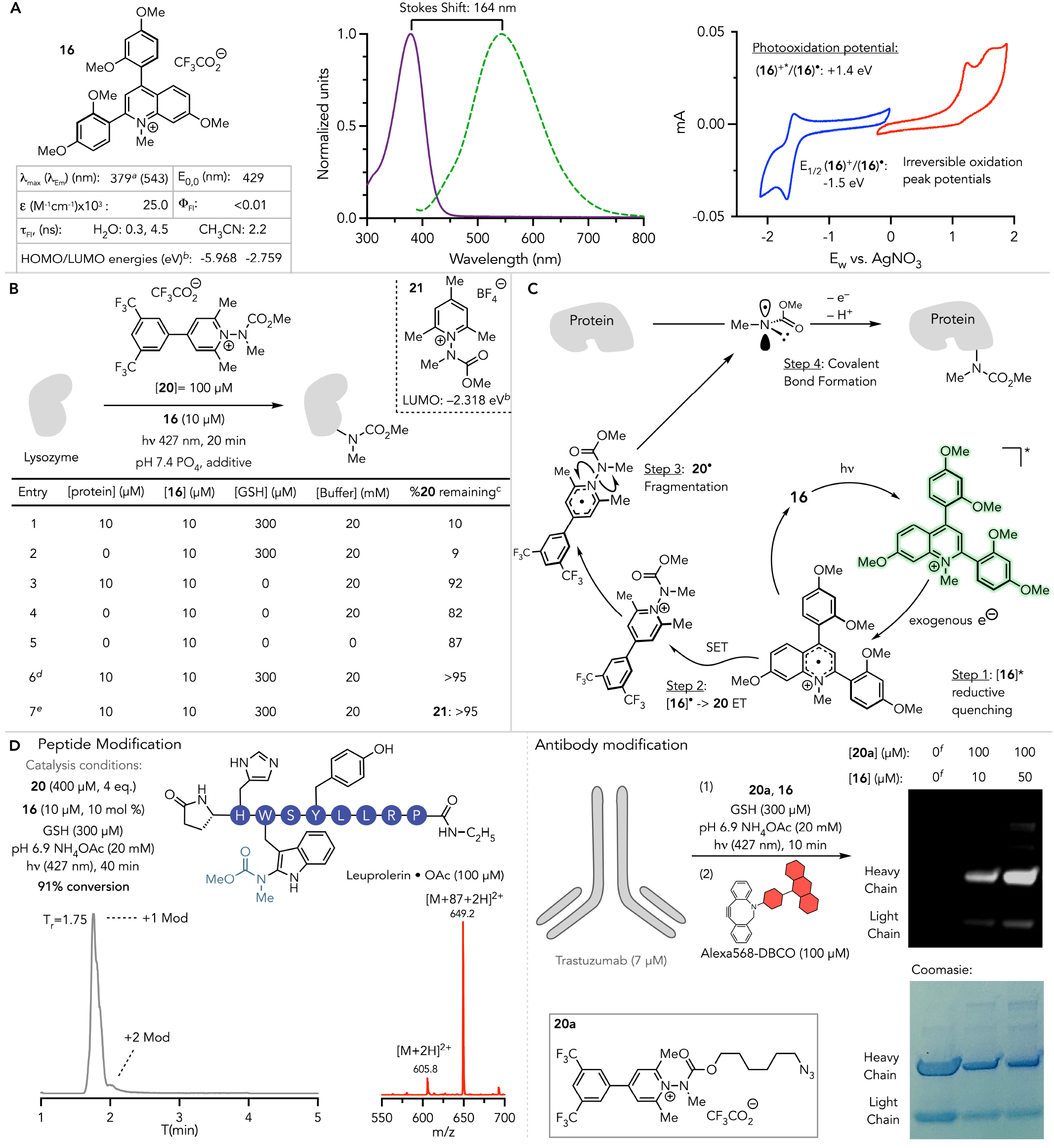
(**A**) Select properties of **16** in 20 mM pH 7.5 Na_2_HPO_4_/NaH_2_PO_4_. Cyclic voltammetry was measured in CH_3_CN. (**B**) Perturbation studies to establish mechanism. (**C**) Proposed sensitization mechanism with **16.** (**D**) Photosensitized labelling of a small peptide (Leuprolerin•OAc) and a large protein (Trastuzumab). ^*a*^Lowest energy transition. ^*b*^Calculated at the UMN12-SX 6-31g(d) SMD=H_2_O level of theory. ^*c*^Measured by LC/MS against an internal standard: entries 1-6: **20a**, entry 7: **20**. ^*d*^No irradiation. ^*e*^100 µM **21** used instead of 100 µM **20**. ^*f*^Antibody not exposed to reaction conditions.

Next, we explored reactivity against therapeutic biomolecules. To show catalytic sensitization, we performed labelling against GnRH agonist peptide Leuprolide•OAc (1 Trp, 1 Tyr, 1 His). Irradiation of a 100 µM solution of peptide using 10 µM **16** (10 mol% sensitizer) and 400 µM **20** (4 eq.) in NH_4_OAc buffer and irradiation with 427 nm light for 40 minutes resulted in 91% conversion to labelled peptide: with MS2 of the resultant conjugate showing modification at Trp. This result highlights that **16** can act catalytically and highlights the potential of applying this chemistry towards peptide derivatization. We then explored labelling on a larger biologic by irradiating a solution of Trastuzumab (7 µM), **16** (10-50 µM), and azide-derived **20a** (100 µM) in 20 mM NH_4_OAc buffer for 10 min with 427 nm light. Installation of Alexa568^11^ and measurement of in-gel fluorescence revealed modifications at both catalyst concentrations to primarily the heavy-chains of trastuzumab (∼7:1 ratio). Taken in conjunction with our initial screening studies, the catalytic pair of **16** and **20** demonstrates competent labelling against both small and large biomolecules under mild conditions with favorable stoichiometries of both reagents.

Given the mildness of these conditions, we posited that **16** and **20** may enable photosensitized labelling in more complex biological environments. We therefore commenced lysate-level experiments in which HEK293T lysates (2 mg/mL) were first treated with sensitizer **16** in varying concentrations and biotinylated **20b** (100 µM) followed by irradiation for 20 minutes and labelling outcomes assessed by western-analysis (**Figure 4A**). Under these conditions, the **16**/**20b** combination gives robust labelling at 5-50 µM concentrations of **16** with a clear dose-response in signal (lanes 1-3). Control studies in which either only sensitizer **16** (lane 4), pyridinium **20b** (lane 5) are irradiated produced no measurable signal, whilst the presence of **16** and **20b**, but in the absence of light also produce no measurable signal (lane 7). These studies demonstrate that the observed lysate labelling is a direct consequence of photosensitization with **16**. We then scanned varying concentrations ratios of **16** and **20a** over which labelling can be observed using western analysis. Screening of three different ratios of **20a**:**16** of 10:1, 4:1, and 1:1 ratios at higher concentrations ([total small molecules]=100-110 µM) yielded similar levels of labelling (see SI Figure S23). Maintaining these same ratios but decreasing total small molecule content to 50-55 µM resulted in fainter signal, but still maintained robust labelling. Decreasing small molecule content further to 10-12.5 µM gives weaker labelling. Additional experiments also show that the rapid reactivity observed for *in vitro* protein ligation is maintained in complex proteomes, with measurable signal between 1-5 minutes (see SI Figures S20, S26). We also found that **2**^5b^ is also a suitable, but slightly less reactive, pyridinium precursor for proteome labelling (see SI Figures S19, S20).

**Figure 4.**
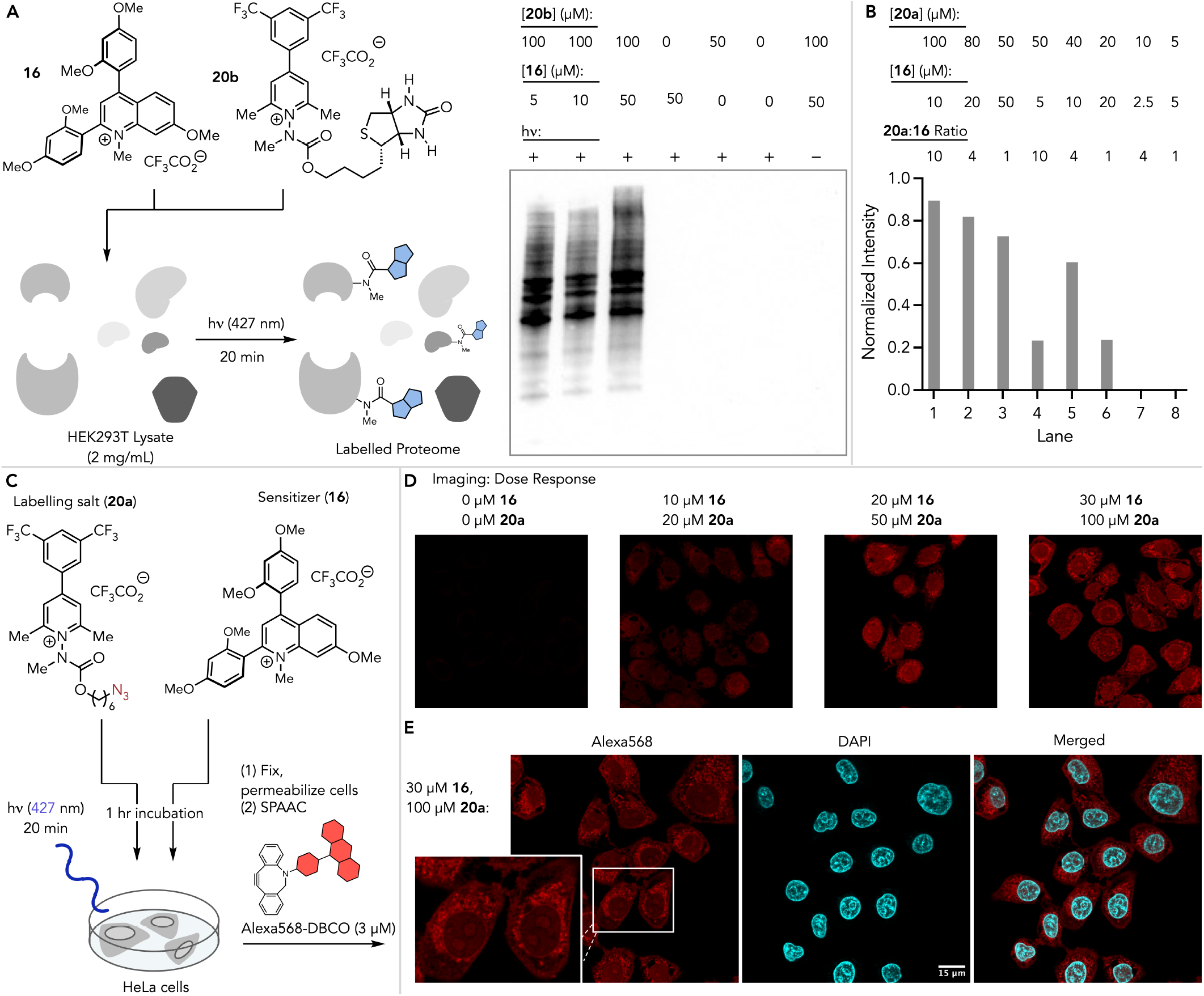
(**A**) Lysate-level reactivity of 16/20b. (**B**) Relative labelling intensities at varying concentrations and ratios of **16** and **20a**. (**C**) Intracellular labelling and fixed cell imaging workflow. (**D**) Dose response at varying concentrations of **16** and **20a**. (**E**) Dual color imaging with Alexa568 (labelled sites) and DAPI (nucleus).

We next assessed the capacity of the **16**/**20** system for intracellular sensitization using an imaging-based approach. Hela cell cultures were incubated with **16** and azide-derived **20a** for 60 minutes, followed by irradiation with 427 nm light for 20 minutes. Cells were then fixed and permeabilized for subsequent introduction of a fluorophore by click chemistry. Multiple iterations of CuAAC were attempted but resulted in significant non-specific labelling^12^ that obfuscated labelling assessments (see SI figure S16). Switching to a strain-promoted cycloaddition with dibenzocyclooctyne-Alexafluor-568 addressed this issue and allowed for facile imaging by confocal microscopy (**Figure 4C**). These studies revealed the capability of both **16** and **20a** to cross cellular membranes and sufficiently co-localize to enable photosensitization in live cells. Aromatic, hydrophobic cations are generally presumed to localize in mitochondria and have been used extensively to deliver payloads to this organelle^13^. Our imaging studies in Figure 4 which feature two hydrophobic, aromatic cations, indicate that these two specific reagents can localize in multiple subcellular locations, including nuclear compartments. Taken in combination with proteomic datasets generated by us^5b^, as well as other imaging-based studies using aromatic cations^14^, indicate that localization of this class of structure are nuanced and structure-dependent. In agreement with our gel-base studies, we observed dose response between fluorescence intensity and reagent concentration up to 50 µM **20a** and 30 µM **16**.

We concluded our studies through chemical proteomic profiling of the **16**/**20a** sensitization system, with the intent of gaining preliminary understanding of the proteomic-subsections and amino acid site selectivity enriched by this combination. HEK293T lysates were incubated with **16** (30 µM) and **20** (100 µM) for 30 min. at room temperature followed by irradiation with 427 nm light for 20 minutes and then sample preparation for peptide-level enrichment. Briefly, labelled proteins were biotinylated using CuAAC and an acid-cleavable biotin tag^15^, followed by digestion and peptide enrichment with neutravidin. Bound peptides were released with formic acid and, following cleanup, were analyzed by LC/MSMS using data-dependent acquisition (DDA) parameters (**Figure 5A**). Experiments were performed in triplicate, and data were analyzed using the Fragpipe software suite^16^ with closed searching, label-free quantitation (LFQ), and sample variance stabilization normalization. This analysis identified 319 enriched proteins (log_2_-fold change ≥ 1) and 615 sites harboring carbamoyl modifications. Of these modifications, 521 were harbored on Trp (**Figures 5B**,**C**, 93% selectivity) with remainder being harbored on low levels on histidine, phenylalanine, and tyrosine. Gene Ontology (GO) analysis of the molecular function of the 319 enriched proteins with Enrichr^17^ showed this system enriches mRNA binding proteins significantly (**Figure 5E**). Proteome-wide annotation of functional Trp sites is lacking, and we highlight the capability of the **16**/**20a** sensitization system to probe this underexplored area by highlighting labelled, but functionally unannotated, Trp residues on three functionally distinct proteins including Nup205 (nuclear protein trafficking)^18^, PPIF (mitochondrial dynamics and apoptosis)^19^, and EIF4A1 (RNA binding)^20^ (**Figure 5F**).

**Figure 5.**
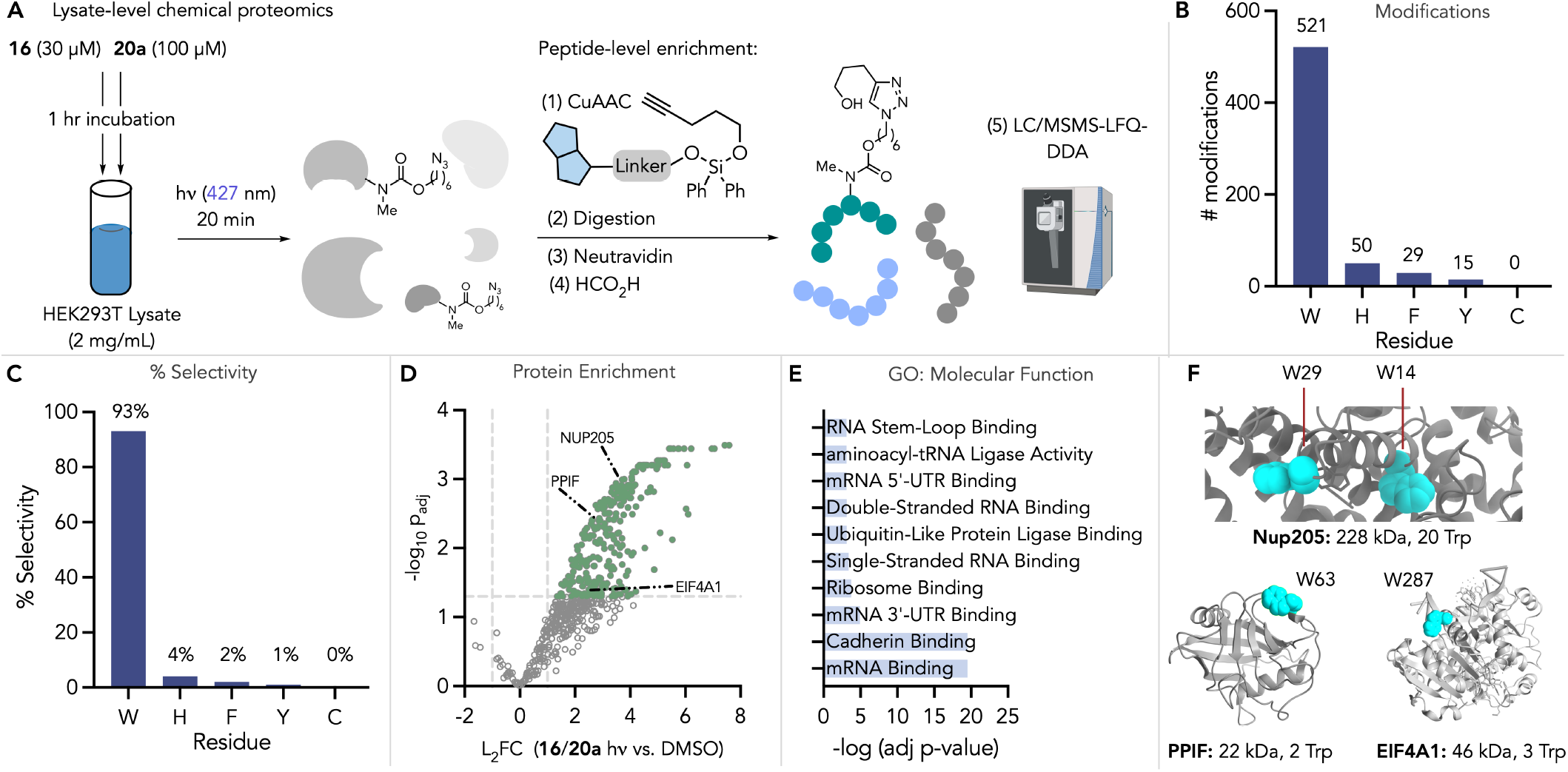
(**A**) Workflow for lysate-level proteome profiling with 16/20a. Experiments were performed in triplicate. (**B**) Summary of detected modifications. (**C**) Selectivity based on amino acid frequency. (**D**) Volcano plot showing differential expression. (**E**) GO molecular function analysis of enriched proteins. (**F**) Representative examples of modified Trp residues lacking functional annotation.

We then extended our proteome profiling work to live HEK293T cells. Live cultures were incubated with **16** and **20a** for 60 minutes at 37 °C, followed by irradiation for 20 minutes at 4 °C, and sample preparation/analysis as described for lysate level experiments (**Figure 6A**). Analysis of enriched proteins compared to controls of DMSO treatment and **16**/**20a** treatment, but no light irradiation revealed 101 proteins differentially expressed compared to both negative controls and validates the need for photoactivation in combination with gel-based studies (see SI Figures S27-S30). GO cellular component analysis shows enrichment of nuclear proteins (57% of all enriched proteins when analyzed in UniProt, Figure S36). This data, when combined with the high Trp selectivity observed from lysate studies and the very short lifetimes of any of the reactive intermediates likely generated during the catalytic cycle (**Figure 3C**), leads us to posit that preorganization of the three reactants is required for a labelling event (**Figure 6C**).

**Figure 6.**
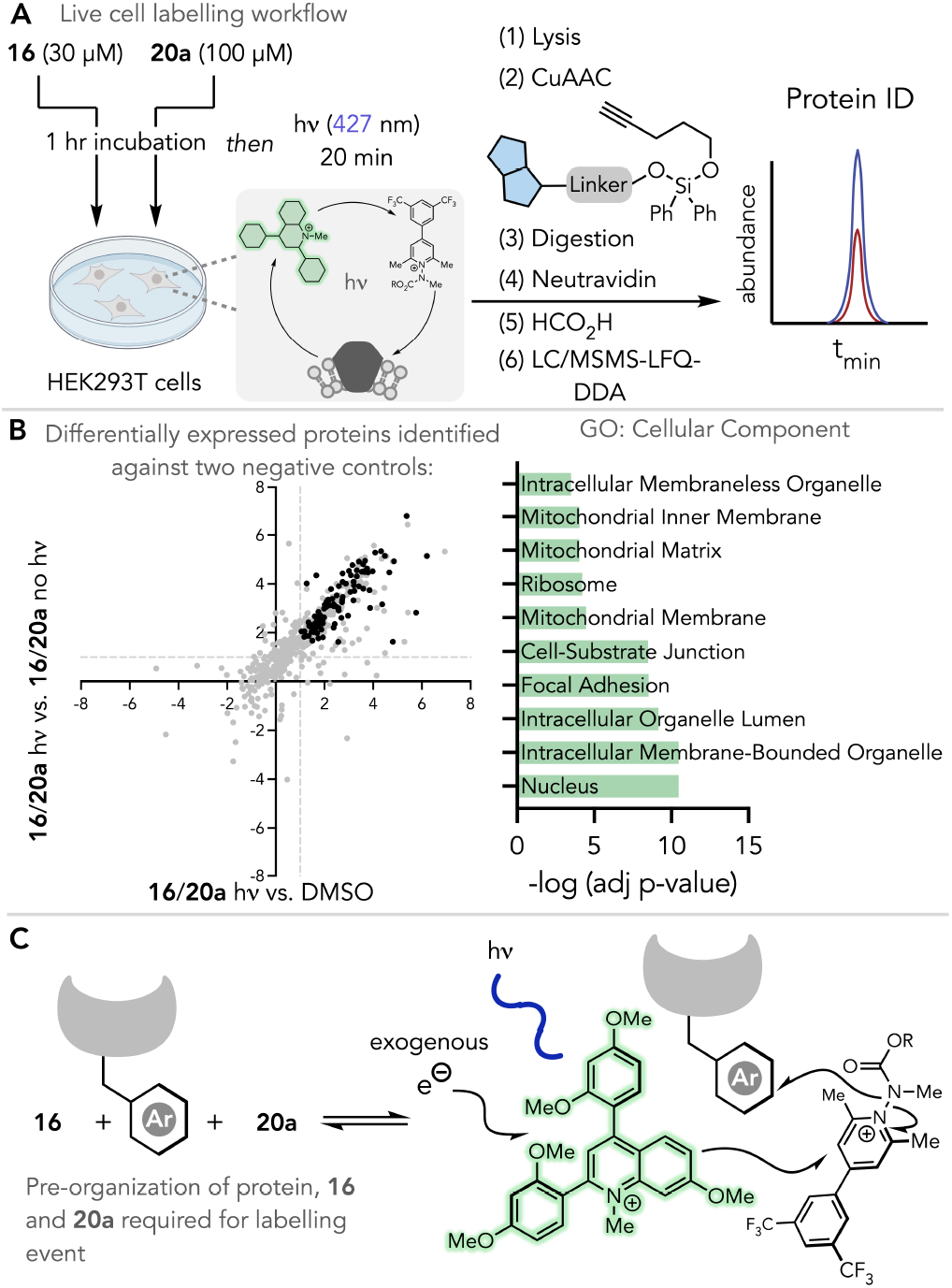
(**A**) Live cell protein labelling using 6/20a. (**B**) Differentially expressed proteins and GO cellular component analyses of identified proteins. (**C**) Additional mechanistic considerations for photosensitized labelling with **16/20a**.

In conclusion, we have developed a biocompatible photosensitization process using aromatic cations. The process features fast protein ligation under very mild conditions using diarylquinolinium sensitizer **16** paired with stoichiometric biarylpyridinium counterpart **20**. We show efficient photosensitized labelling of structurally variable biomolecules. Since the reagents are simple organic molecules that readily traverse cellular membranes, they are capable of sufficient intracellular colocalization to pre-complex with target proteins and produced labelling inside cells.

## Supporting information

Experimental Procedures

Computation Details

Proteomics Data Tables

## ASSOCIATED CONTENT

### Supporting Information

Supporting information consists experimental procedures for catalyst and pyridinium probe synthesis, protein labelling screening, protein modification experiments, mechanistic experiments, fixed cell imaging, proteome profiling, and small molecule NMR data. A second document contains computational details and procedures. A third spreadsheet contains proteomic profiling data tables for lysate- and live cell-level proteomics experiments.

The Supporting Information is available free of charge on the ACS Publications website.

## Notes

The authors declare no competing financial interest.

## ACKNOWLEDGMENT

We thank the National Institutes for General Medical Sciences (R35 GM143120) for supporting this work. We also thank the National Science Foundation, Division of Chemistry, Chemistry Life Processes Institute, for supporting of this work (CHE-2302483). We acknowledge the W.M. Keck Center for Nano-Scale Imaging (RRID:SCR_022884) and the Nuclear magnetic Resonance Facility (RRID:SCR_012716) at the University of Arizona Department of Chemistry & Biochemistry, as well as the Imaging Cores – Optical (RRID:SCR_023355) for instrumentation support. We also thank NSF (grant no.’s 1920234, 840336, 9214383, 9729350) for additional instrumentation support. We thank the University of Arizona for startup funds. We thank the Gianetti group for assistance with cyclic voltammetry experiments.

## Insert Table of Contents artwork here

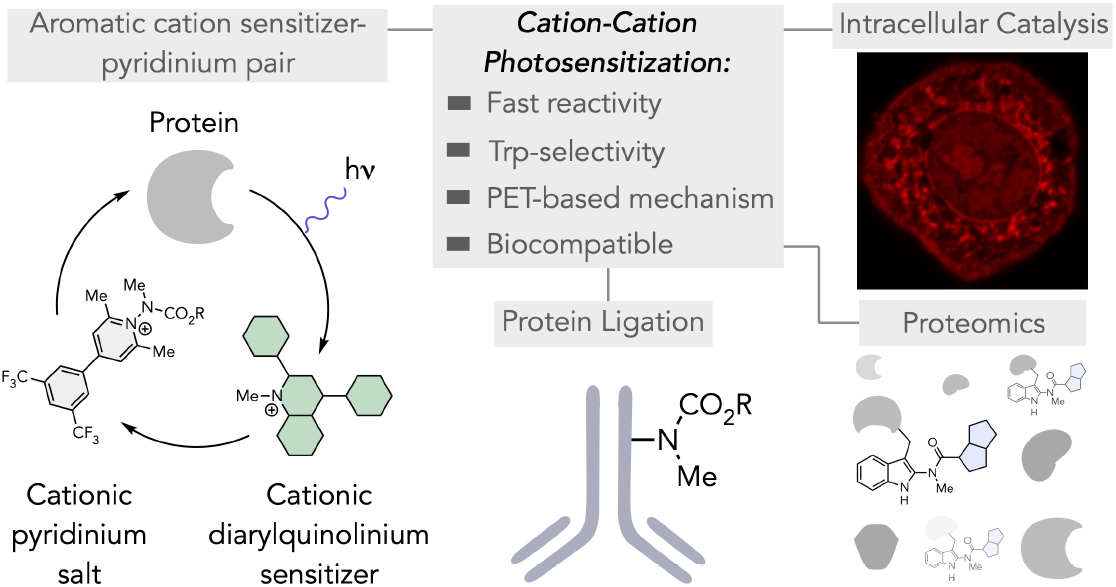

## Notes

### Competing Interest Statement

The authors have declared no competing interest.

