## Supplementary material for "Cation-Cation Photosensitization for Protein Ligation and Intracellular Catalysis": Experimental Procedures

#### Table of Contents


1. General considerations……………………………………………………………………...…S3
2. Analytical Methods…………………………………………………………………………S3-S4
3. Small molecule synthetic procedure…………………………………………………….S5-S45
4. Protein labelling:
   4.1 Observation of sensitization/commercial catalyst screen……………………….S46-S49
   4.2 Screening/optimization……………………………………………………………..S50-S52
   4.3 Labelling with reporter tagged pyridiniums………………………………………S53-S54
   4.4 Leuprorelin•OAc labelling……………………………......................................S55-S58
   4.5 Trastuzumab labelling………………………………………………………......S58-S60
5. Mechanistic studies………………………………………………………………………S61-S65
6. Biology……………………………………………………………………………………S66-S85
7. References……………………………………………………………………………………S86
8. NMR spectra……………………………………………………………………………S87-S172

#### 1. General considerations:

All commercially available compounds were used without further purification; chemicals were purchased from TCI, Sigma Aldrich, Synthonix, Ambeed and Combiblocks. Water was purified using a Millipore Milli–Q Integral Water Purification System. All solvents were obtained from Fisher and used as received. Dry solvents were obtained from Sigma-Aldrich. Reactions were monitored by thin layer chromatography (TLC) analysis. This was done with silica gel precoated glass silica gel plates (Millipore Sigma Ref. 1057150001) and visualized using UV light (254 nm). Column chromatography was performed under positive pressure of compressed air using SiliCycle SiliaFlash® P60 silica gel (230-400 mesh. Purification of Quinolinium salts were performed using an ISCO Combiflash $300^{+}$ or Thermo Fisher Dionex UltiMate 3000 semi-preparative HPLC. For synthesized compounds, Nuclear magnetic resonance (NMR) spectra were acquired at ambient temperature unless otherwise stated using either a Bruker Avance III 400 / NEO 500 or Avance NEO 600 spectrometer at the University of Arizona. Reactions were performed in heat-dried glassware using appropriate Schlenk techniques. Many of the pyridinium salts exist as rotamers. Where possible, NMR signals are assigned integration values equal to the number of protons associated with the given functional group of the given rotamer. The coupling constants J are given in Hz. Spin multiplicities are reported as s = singlet, d = doublet, t = triplet, q = quartet, dd = doublet of doublets, ddd = doublet of doublet of doublets, dd = doublet of doublets. Infrared (IR) spectra were collected on a Thermo Fisher Scientific Nicolet IS50R FT-IR equipped with an ATR probe at the Keck center, University of Arizona. High-resolution mass spectra were at the analytical & biological mass spectrometry facility at the University of Arizona. Absorption spectra were acquired on a Jasco V-760 spectrophotometer. Emission spectra, Fluorescence lifetime and absolute quantum yield were acquired on a Horiba FluoroMax Plus fluorescence spectrophotometer. Gels were visualized using a UVP ChemStudio Plus imaging instrument. The reagents purchased from commercial sources were used as received. Reverse-phased flash chromatography was performed on a Teledyne-Isco CombiFlash 300 instrument. Methyl 1-methylhydrazine-1-carboxylate[1], 5-((3a*S*,4S,6a*R*)-2-oxohexahydro-1*H*-thieno[3,4-d]imidazole-4-yl)pentyl 1methylhydrazine-1-carboxylate[2], 6-azidohexyl 1-methylhydrazine-1-carboxylate[2], Biotin-alkyne[3] was synthesized according to the previously reported procedure. Lysozyme (from chicken egg white, >40,000units/mg of protein) was purchased from Sigma Aldrich. Leuprolide acetate and Trastuzumab (anti-HER2) were purchased from Apexbio Technology LLC and used as received. Peptide, protein, and antibody were used without additional purification. Alexa Fluor 568 DBCO was purchased from Lumiprobe. Protein modification reactions were monitored and evaluated *via* LC/MS using a ThermoFisher Vanquish Flex LC coupled to a Thermo-Fisher LTQ XL ion trap mass spectrometer equipped with a heated ESI probe.

#### 2. Analytical Methods

##### 2.1 General HPLC method:

**Method A.** solvents: 0.1% $CF_3COOH$ in $H_2O$, B: 0.1% $CF_3COOH$ in $CH_3CN$; hold 10% B for 1 min, 10-45% B over 25 min, hold 45% B for 5 min, 45-95% B over 3 min, hold 95% B for 3 min, 95-10% B over 2 min, hold 10% B for 3 min, UV-Vis: 390, 365, 254 nm, column: Kinetex® 5µm C18 100 Å, LC Column 150 x 21.2 mm, flow rate: 10 mL/min.

#### 2.2 Reverse-phased flash chromatography methods

**Method A:** solvents: A: 0.1% $CF_3COOH$ in $H_2O$, B: 0.1% $CF_3COOH$ in $CH_3CN$; method: hold 5% B for 4 min, 5-12% B over 6 min, hold 12% B for 3 min, 12-35% B for 8 min, hold 35% for 2 min, 35-90% B over 7 min, hold 90% B for 2 min, 90-10% B over 1 min, hold 10% B for 2 min; UV-Vis: 280, 254 nm, column: RediSep Gold C18Aq 50g HP C18, 100 Å, Teledyne ISCO (part # 69-2203-335); flow rate: 50 mL/min, 300 psi max.

**Method B:** solvents: A: 0.1% $CF_3COOH$ in $H_2O$, B: 0.1% $CF_3COOH$ in $CH_3CN$; method: hold 5% B for 5.5 min, 5-10% B over 2.6 min, hold 10% B for 8.9 min, 10-20% B over 2 min, 20-30% B for 8 min, 30-50% B over 1 min, hold 50% for 5 min, 50-90% for 10 min, hold 90% for 2 min, 90-10% B over 2 min, hold 10% B for 2 min; UV-Vis: 280, 254 nm, column: RediSep Gold C18Aq 50g HP C18, 100 Å, Teledyne ISCO (part # 69-2203-335); flow rate: 50 mL/min, 300 psi max.

**Method C: solvents:** A: 0.1% $CF_3COOH$ in $H_2O$, B: 0.1% $CF_3COOH$ in $CH_3CN$; method: hold 5% B for 5 min, 5-20% B over 4 min, hold 20% B for 5 min, 20-45% B over 6 min, hold 45% B for 2 min, 45-64% B over 4 min, hold 64% B for 5 min, 64-90% B over 6 min, hold 90% B for 5 min, 90-10% B over 2 min, hold 10% B for 2 min; UV-Vis: 284, 254 nm, column: RediSep Gold C18Aq 50g HP C18, 100 Å, Teledyne ISCO (part # 69-2203-335); flow rate: 50 mL/min, 300 psi max.

#### 2.3 LCMS Methods:

**LC-MS Method A**: solvents: A: 0.1% formic acid in $H_2O$ B: 0.1% formic acid in $CH_3CN$, method: 20-70% B over 5 min, 70-95% B over 1 min, hold 95% B for 2 min, 95-20% B over 0.5 min, then hold 20% B 1.5 min, column: Aeris™ 3.6 µm WIDEPORE XB-C18 20, LC Column 100 x 21 mm, m/z range: 500-2000, flow rate: 0.2 mL/min.

**LC-MS Method B.** solvents: A: 0.1% formic acid in $H_2O$ B: 0.1% formic acid in $CH_3CN$, method: hold 10% B for 0.2 min, 10-90% B over 2.8 min, hold 90% over 1 min, 90-10% B over 0.5 min, then hold 10% B 30 sec, m/z range: 200-800, column: Kinetex® 1.3µm C18 100 Å, LC Column 50 x 2.1 mm, flow rate: 0.3 mL/min.

**LC-MS Method C** solvents: A: 0.1% formic acid in $H_2O$ B: 0.1% formic acid in $CH_3CN$, method: hold 5% B for 0 min, 5-10% B over 1.5 min, 10-20% B over 3.2 min, hold 20% over 1 min, 20-30% B over 0.5 min, 30-40% B over 1.0 min, 40-50% B over 0.5 min, 50-60% B over 0.4 min, 60-70% B over 0.6 min, 70-75% B over 0.5 min, 75-80% B over 0.6 min, 80-85% B over 0.6 min, 85-90% B over 0.5 min, 90-95% B over 1.0 min, hold 95% B for 1 min, 95-10% B over 1.0 min, hold 10% B for 2 min, m/z range: 200-800, column: Kinetex® 1.3µm C18 100 Å, LC Column 50 x 2.1 mm, flow rate: 0.3 mL/min.

**LC-MS Method D** solvents: A: 0.1% formic acid in $H_2O$ B: 0.1% formic acid in $CH_3CN$, method: hold 5% B for 0 min, 5-10% B over 1.5 min, 10-20% B over 2.5 min, hold 20% over 1 min, 20-30% B over 0.5 min, 30-40% B over 1.0 min, 40-50% B over 0.5 min, 50-60% B over 0.4 min, 60-70% B over 0.6 min, 70-75% B over 0.5 min, 75-80% B over 0.4 min, 80-85% B over 0.6 min, 85-90% B over 1.0 min, 90-95% B over 1.0 min, hold 95% B for 1 min, 95-10% B over 1.0 min, hold 10% B for 2 min, m/z range: 200-800, column: Kinetex® 1.3µm C18 100 Å, LC Column 50 x 2.1 mm, flow rate: 0.3 mL/min.

#### 3. Small molecule synthetic procedures

**1-((methoxycarbonyl)(methyl)amino)-4-(6-methoxynapthalen-2-yl)-2,6-dimethylpyridin-1-ium trifluoroacetate (1):**

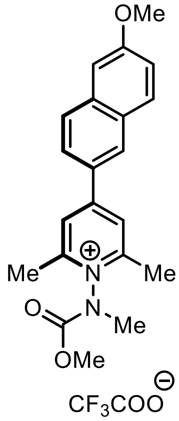


**1**

The synthesis of compound **1** was achieved by adapting our previously reported procedure,[2] with the added step that an aliquot of the tetrafluoroborate salt of **1** was purified by Teledyne ISCO Combiflash 300$^+$ method A to yield trifluoroacetate salt **1** as a yellow powder.

**$^1$H NMR (500 MHz, $CD_3CN$)**: δ: 8.55 – 8.53 (m, 1H), 8.40 (s, 2H), 8.03 – 7.94 (m, 3H), 7.34 (s, 1H), 7.29 – 7.23 (m, 1H), 3.99 (s, 1.7H), 3.96 (s, 3H), 3.82 (s, 1.3H), 3.61 (s, 1,7H), 3.57 (s, 1.3H), 2.81 (s, 6H) ppm;

**$^{13}$C NMR (126 MHz, $CD_3CN$):** δ: 161.8, 158.9, 155.2, 153.7, 138.6, 132.2, 130.9, 130.1, 129.7, 129.3, 125.5, 121.4, 106.9, 56.1, 55.6, 38.1, 37.1, 19.1 ppm;

**$^{19}$FNMR (471 MHz, $CD_3CN$):** δ = -77.24 (s, 3F) ppm;

**FT-IR ($v^{max}$ / $cm^{-1}$, neat):** 1732, 1689, 1611, 1562, 1493, 1457, 1408, 1376, 1351, 1327, 1266, 1193, 1168, 1051, 1022, 939, 853, 792, 759, 705;

**MP (**°C**):** 45-49

**2,6-Dimethyl-1-(methyl(((5-((3a*R*,4*R*,6a*S*)-2-oxohexahydro-1*H*-thieno[3,4-d]imidazol-4-yl)pentyl)oxy)carbonyl)amino)-4-phenylpyridin-1-ium 2,2,2-trifluoroacetate (2):**

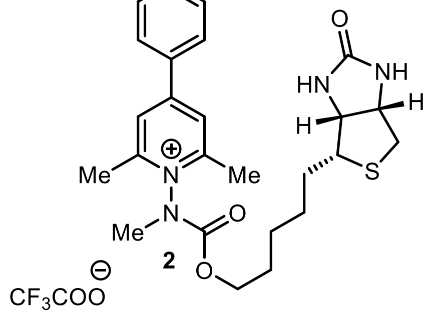


The synthesis of the compound **2** was achieved by adapting our previously reported procedure,[2] using pyrylium salt[4] **S22** (88 mg, 0.48 mmol, 1.0 eq.), added 5-((3a*S*,4S,6a*R*)-2-oxohexahydro-1*H*-thieno[3,4-d]imidazole-4-yl)pentyl 1methylhydrazine-1-carboxylate[2] (431 mg, 1.43 mmol, 3.0 eq.) with the added step that an aliquot of the tetrafluoroborate salt of **2** was purified by HPLC using Method A to yield trifluoroacetate salt **2** (51 mg, Yield: 23%) as a yellow oil.

**$^1$H NMR (600 MHz, $CD_3CN$):** δ 8.19 (d, *J* = 22.9 Hz, 2H), 8.01 – 7.92 (m, 2H), 7.75 – 7.60 (m, 3H), 5.62 (s, 2H), 4.43 (dd, *J* = 7.8, 4.9 Hz, 0.53 H), 4.35 – 4.24 (m, 2H), 4.15 (dtt, *J* = 19.1, 7.7, 4.4 Hz, 1.52 H), 3.53 (s, 1.44 H), 3.48 (s, 1.52 H), 3.19 (ddd, *J* = 8.9, 6.0, 4.4 Hz, 0.53 H), 3.01 (dt, *J* = 9.5, 5.2 Hz, 0.54 H), 2.77 (m, 0.58 H), 2.89 (dd, *J* = 12.7, 4.9 Hz, 0.64 H), 2.80 – 2.71 (m, 6H), 2.67 – 2.65 (m, 0.59 H), 2.57 (d, *J* = 12.7 Hz, 0.43 H), 1.81 – 1.66 (m, 1.53 H), 1.64 – 1.34 (m, 4.64 H), 1.27 – 1.09 (m, 1.69 H) ppm;

**$^{13}$C NMR (151 MHz, $CD_3CN$)**: δ 164.3, 160.8, 160.6, 160.4, 160.1, 159.2, 159.1, 159.1, 158.5, 158.5, 154.1, 152.7, 134.6, 134.4, 133.8, 133.7, 130.9, 130.8, 130.8, 129.4, 129.4, 126.1, 126.1, 126.0, 122.9, 120.6, 116.7, 114.8, 69.5, 69.2, 62.6, 62.6, 60.9, 60.8, 56.6, 56.4, 41.2, 41.1, 38.3, 37.2, 29.5, 29.4, 29.2, 29.2, 28.8, 26.5, 26.4, 19.6, 19.5 ppm;

**$^{19}$F NMR (471 MHz, $CD_3CN$):** δ -76.54 (s, 3F) ppm;

**FT-IR ($v^{max}$ / $cm^{-1}$, neat):** 1732, 1689, 1630, 1566, 1446, 1375, 1332, 1268, 1172, 1129, 1034, 1003, 919, 883, 799, 771, 706, 692

**Synthesis of Quinolinium Compounds and Salts:**

**Summary of synthetic route:**

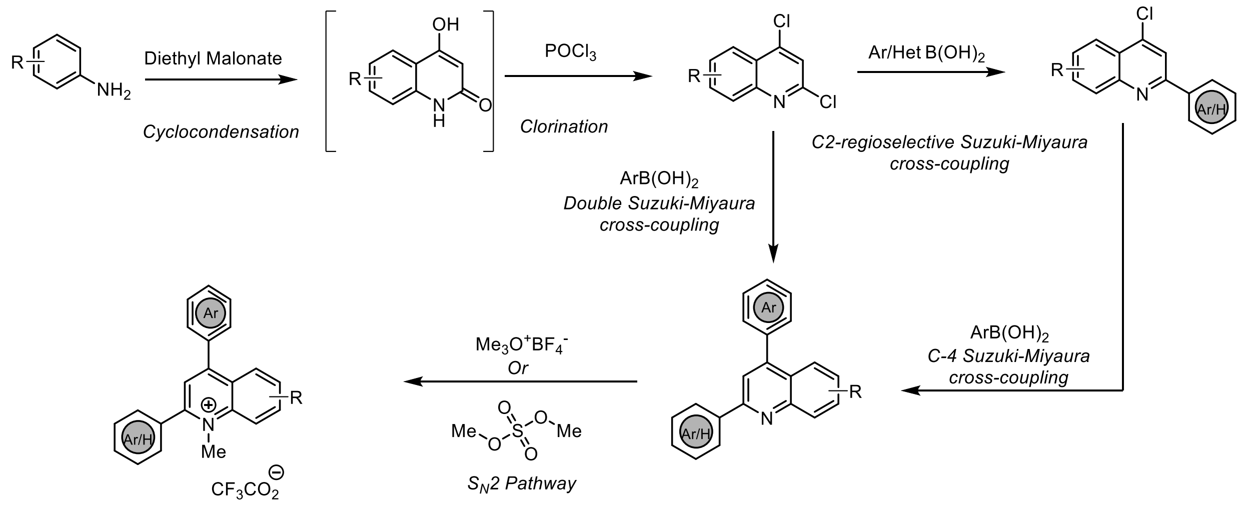


**General Synthetic Routes for 2,4-dichloro quinolinium precursor:**

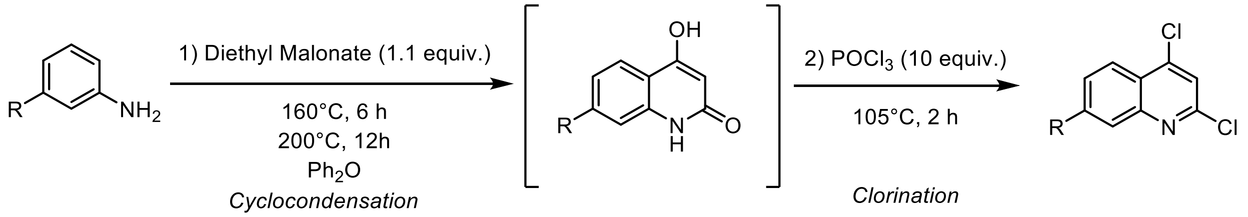


**General Procedure A:**

Synthesis 2,4-dichloro arylquinoline analogs was achieved by adapting the procedure from Chenoweth,[5] as follows: In a 20 mL vial Chemglass Life Sciences (CG-4920-02) equipped with a stir bar was dried under vacuum, allowed to cool to room temperature, and refilled with $N_2$, was charged with arylamine (1.0 eq.), diphenylether and the reaction mixture was stirred vigorously.

To the vial, diethyl malonate (1.1 eq.) was added, and capped with 20mm Aluminum seal with PTFE-faced Silicone septa to allow self-condensation on aluminum block. The reaction vial, and the reaction mixture was vigorously stirred and heated at 160 °C for 6 h until the solution turned solid. The reaction was then heated to 200 °C for approximately 12 h until TLC indicated the complete formation of 4-hydroxy arylquinolin-2(1*H*)-one. The reaction vial turned brown and cooled to room temperature. Without any wash or purification, phosphorous oxychloride (10 eq.) was added to the reaction vial. The reaction mixture was then stirred vigorously at 105 °C on aluminum block for 3 h. The reaction was cooled to 0 °C over the course of 30 min. The resulting mixture was poured into a larger Erlenmeyer flask and neutralized with neat $K_2CO_3$ and ice deionized water slowly. The reaction mixture was then poured into a separatory funnel and extracted with EtOAc (3 times). Finally, the organic layer was washed with brine, dried with $Na_2SO_4$, and concentrated using a rotary evaporator. Residual diphenylether was filtered out by silica column with 100% hexane. The crude mixture was purified by silica gel column chromatography (0% to 10% EtOAc in Hexane) to yield the desired 2,4-dichloro arylquinoloium compound.

**2,4-Dichloro-*N*,*N*-dimethylquinolin-7-amine (7-NMe$_2$-DCQ):**

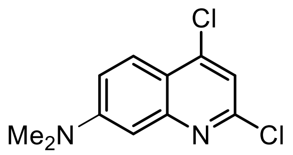


Following **General Procedure A** with *m*-anisidine (1.001 g, 10.75 mmol, 1.0 eq.), diethyl malonate (1.291 g, 11.82 mmol, 1.1 eq.) and diphenyl ether (2 mL) were heated at 160 °C for 6 h and further heated at 200ºC for 12 hours. After completion, the reaction mixture was allowed to cool to room temperature and $POCl_3$ (16.48 g, 107.5 mmol, 10.0 eq.) was added to the reaction mixture with vigorous stirring. The reaction was then heated at 105ºC for 2 hours. After completion of the reaction as monitored by thin layer chromatography (TLC) analysis, the mixture was quenched by excess solid $K_2CO_3$, and ice-cold water carefully followed by workup with EtOAc (3 x 200 mL). The organic layers were combined, washed with brine (50 mL), dried over anhydrous $Na_2SO_4$, filtered, and concentrated using a rotary evaporator. The crude mixture was purified by silica gel column chromatography to afford 2,4-dichloro-*N*,*N*-dimethylquinolin-7-amine as yellow solid in 68% yield (1.201 g).

**$^1$H NMR (500 MHz, CDCl$_3$):** δ 7.99 (d, *J* = 9.3 Hz, 1H), 7.20 (dd, *J* = 9.3, 2.6 Hz, 1H), 7.17 (s, 1H), 7.10 (d, *J* = 2.6 Hz, 1H), 3.13 (s, 6H) ppm;

**$^{13}$C NMR (126 MHz, CDCl$_3$):** δ 152.6, 149.6, 149.3, 144.8, 125.1, 117.3, 117.2, 117.0, 116.9, 104.7, 40.4, 30.3 ppm;

This compound is known, and characterization data is consistent with the literature report.[5]

**2,4-Dichloro-7-methoxyquinoline (7-OMe-DCQ):**

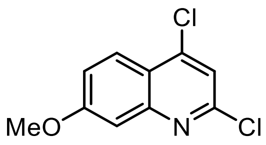


Following **General Procedure A** with 3-methoxyaniline (1.08 g, 8.77 mmol, 1.0 eq.), diethyl malonate (1.55 g, 9.65 mmol, 1.1 eq.) and diphenyl ether (2 mL) were heated at 160 °C for 6 h and further heated at 200ºC for 12 hours. After completion, the reaction mixture was allowed to cool to room temperature and $POCl_3$ (13.45 g, 87.69 mmol, 10.0 eq.) was added to the reaction mixture with vigorous stirring. The reaction was then heated at 105ºC for 2 hours. After completion of the reaction as monitored by thin layer chromatography (TLC) analysis, the mixture was quenched by solid $K_2CO_3$, and ice-cold water carefully followed by workup with EtOAc (3 x 200 mL). The organic layers were combined, washed with brine (50 mL), dried over anhydrous $Na_2SO_4$, filtered, and concentrated using a rotary evaporator. The crude mixture was purified by silica gel column chromatography to afford 2,4-dichloro-7-methoxyquinoline as white solid in 11 % yield (0.212 mg).

**$^1$H NMR (500 MHz, $CDCl_3$):** δ 8.06 (d, *J* = 9.2 Hz, 1H), 7.56 – 7.32 (m, 2H), 7.29 – 7.26 (m, 1H), 3.94 (s, 3H) ppm;

**$^{13}$C NMR (126 MHz, $CDCl_3$):** δ 162.4, 150.4, 150.2, 144.1, 125.4, 120.9, 120.3, 119.6, 55.9 ppm; This compound is known, and characterization data is consistent with the literature report.[6]

##### 3.3 Synthesis of symmetrical 2,4-diaryl quinolines *via* double Suzuki−Miyaura cross-coupling reaction

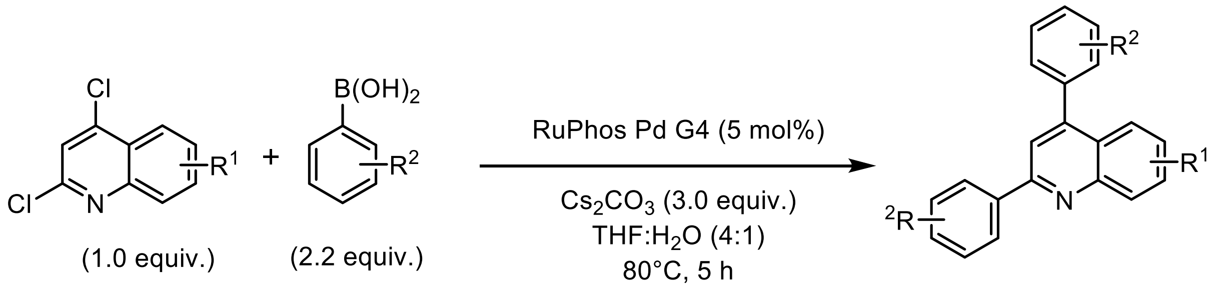


***Double Suzuki−Miyaura cross-coupling reaction***

**General Procedure B:**

A 20 mL vial Chemglass Life Sciences (CG-4920-02) equipped with a stir bar was dried under vacuum, allowed to cool to room temperature, and refilled with $N_2$. The vial was charged with 2,4-dichloro arylquinoline (1.0 eq.), degassed THF and stirred vigorously on aluminum block. To the vial, added RuPhos Pd G4 (5 mol%) followed by aryl boronic acid (2.2 eq.) while stirring. The vial was finally charged with a solution of $Cs_2CO_3$ (3.0 eq. in dd$H_2O$, 0.1 M). The vial was capped with 20mm Aluminum seal with PTFE-faced Silicone septa and the reaction mixture was stirred for 5 hours at 80 ˚C. After the completion of the reaction as monitored by TLC, the mixture was cooled to room temperature; after that, the solvent was evaporated using a rotary evaporator and diluted with ethyl acetate and water. The organic layer was separated, and the organic layer was washed with brine, dried over anhydrous $Na_2SO_4$, filtered, and concentrated using a rotary evaporator using a rotary evaporator. The crude mixture was purified by silica gel column chromatography to obtain the final product.

**2,4-Bis(4-methoxyphenyl)quinoline (S1):**

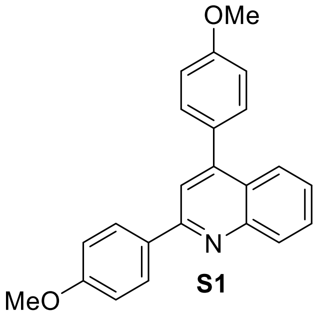

**S1**

Compound **S1** was synthesized using **General Procedure B:** 2,4-dichloroquinoline (100 mg, 505 μmol, 1.0 eq.) in THF (4.0 mL), 4-methoxyphenylboronic acid (169 mg, 1.11 mmol, 2.2 eq.), $Cs_2CO_3$ (494 mg, 1.51 mmol, 3.0 eq. suspended in 1.0 mL $H_2O$), and RuPhos Pd G4 (21 mg, 25 *μ*mol, 5.0 mol%), with heating to 80 °C for 5 hours on aluminum block. Purification by silica gel column chromatography using 20% ethyl acetate/hexane solvent mixture as an eluent, gave **S1** (121 mg, Yield: 70%) as a white solid.

**$^1$H-NMR (500 MHz, $CDCl_3$):** δ 8.27 – 8.13 (m, 3H), 7.93 (dd, *J* = 8.4, 1.4 Hz, 1H), 7.76 (s, 1H), 7.71 (ddd, *J* = 8.4, 6.8, 1.4 Hz, 1H), 7.59 – 7.48 (m, 2H), 7.45 (ddd, *J* = 8.2, 6.8, 1.3 Hz, 1H), 7.17 – 7.01 (m, 4H), 3.91 (s, 3H), 3.89 (s, 3H) ppm;

**$^{13}$C-NMR (126 MHz, $CDCl_3$):** δ 160.9, 159.9, 156.5, 148.8, 132.4, 130.9, 130.0, 129.5, 129.0, 125.9, 125.8, 125.7, 118.9, 114.3, 114.1, 55.5, 55.5 ppm;

**FT-IR ($v^{max}$ / $cm^{-1}$, neat):** 1710, 1605, 1590, 1542, 1514, 1495, 1456, 1440, 1425, 1357, 1289, 1238, 1169, 1110, 1028, 830, 790, 763, 732, 708

**HRMS-ESI (m/z):** calcd for $C_{23}H_{19}NO_2$ $[M + H]^+$ 342.1494, found 342.1488;

**M.P (°C):** 66-68

**2,4-Bis(2,4-dimethoxyphenyl)quinoline (S2):**

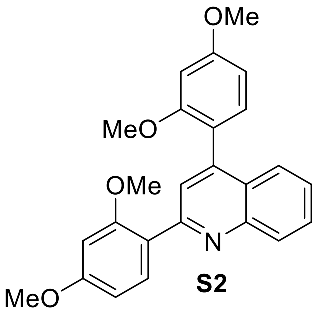

**S2**

Compound **S2** was synthesized using **General Procedure B:** 2,4-dichloroquinoline (0.151 g, 0.757 mmol, 1.0 eq.) in THF (6.1 mL), 2,4-dimethoxyphenylboronic acid (303 mg, 1.67 mmol, 2.2

eq.), $Cs_2CO_3$ (740 mg, 2.27 mmol, 3.0 eq. suspended in 1.5 mL $H_2O$), RuPhos Pd G4 (32 mg, 38 *μ*mol, 5.0 mol%) with heating to 80 °C for 5 hours on aluminum block. Purification by silica gel column chromatography using 30% ethyl acetate/hexane solvent mixture as an eluent, gave **S2** (0.261 g, Yield: 86%) as a white solid.

**$^{1}$H-NMR (500 MHz, $CDCl_3$):** δ 8.22 (s, 1H), 7.90 (d, *J* = 8.5 Hz, 1H), 7.81 (s, 1H), 7.72 – 7.61 (m, 2H), 7.40 (t, *J* = 7.4 Hz, 1H), 7.30 – 7.24 (m, 2H), 6.82 – 6.62 (m, 3H), 6.57 (d, *J* = 2.4 Hz, 1H), 3.91 (s, 3H), 3.88 (s, 3H), 3.83 (s, 3H), 3.72 (s, 3H) ppm;

**$^{13}$C-NMR (126 MHz, $CD_3CN$):** δ 162.9, 162.5, 159.6, 159.0, 157.2, 149.3, 145.7, 133.2, 132.7, 130.3, 129.9, 127.1, 126.7, 125.3, 123.0, 120.5, 106.6, 106.0, 99.7, 99.5, 56.4, 56.2, 56.2, 32.3 ppm;

**FT-IR ($v^{max}$ / $cm^{-1}$, neat):** 1710, 1605, 1577, 1543, 1509, 1494, 1454, 1437, 1418, 1353, 1301, 1281, 1261, 1205, 1157, 1128, 1115, 1028, 935, 831, 803, 765

**HRMS-ESI (m/z):** calcd for $C_{25}H_{24}NO_4$ $[M + H]^+$ 402.1705, found 402.1699;

**M.P (°C):** 88-90

**7-Methoxy-2,4-bis(4-methoxyphenyl)quinoline (S3):**

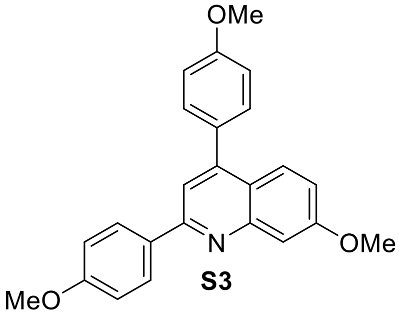


**S3**

Compound **S3** was synthesized using **General Procedure B:** 2,4-dichloro-7-methoxyquinoline (0.200 g, 0.877 mmol, 1.0 eq.) in THF (7.0 mL), 4-methoxyphenylboronic acid (147 mg, 1.93 mmol, 2.2 eq.), $Cs_2CO_3$ (857 mg, 2.63 mmol, 3.0 eq. suspended in 1.8 mL $H_2O$), RuPhos Pd G4 (37 mg, 44 *μ*mol, 5.0 mol%) with heating to 80 °C for 5 hours on aluminum block. Purification by silica gel column chromatography using 25% ethyl acetate/hexane solvent mixture as an eluent, gave **S3** (0.266 g, Yield: 82%) as a white solid.

**$^{1}$H-NMR (500 MHz, $CDCl_3$):** δ 8.14 (d, *J* = 8.8 Hz, 2H), 7.81 (d, *J* = 9.2 Hz, 1H), 7.60 (d, *J* = 5.7 Hz, 2H), 7.52 – 7.44 (m, 2H), 7.16 – 6.98 (m, 5H), 3.99 (s, 3H), 3.89 (d, *J* = 10.1 Hz, 6H) ppm;

**$^{13}$C-NMR (126 MHz, $CDCl_3$)** δ 160.9, 160.8, 159.9, 156.7, 150.4, 149.0, 132.1, 130.9, 130.8, 129.0, 126.9, 120.8, 119.0, 117.0, 114.3, 114.1, 107.6, 55.7, 55.5, 55.5 ppm;

**FT-IR ($v^{max}$ / $cm^{-1}$, neat):** 1709, 1618, 1606, 1545, 1514, 1495, 1460, 1440, 1396, 1358, 1290, 1266, 1220, 1173, 1111, 1028, 831, 788, 763, 735

**HRMS-ESI (m/z):** calcd for $C_{24}H_{22}NO_3$ $[M + H]^+$ 372.1599, found 372.1595;

**M.P (°C):** 106-108

**2,4-Bis(4-methoxyphenyl)-*N,N*-dimethylquinolin-7-amine (S4):**

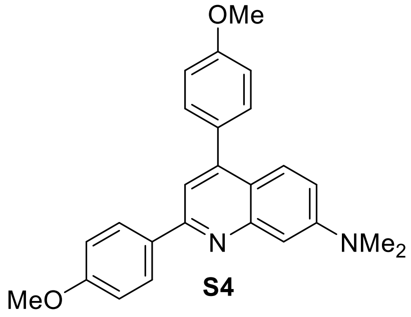

**S4**

Compound **S4** was synthesized using **General Procedure B:** 2,4-dichloro-*N*,*N*-dimethylquinolin-7-amine (301 mg, 1.24 mmol, 1.0 eq.) in THF (9.9 mL), 4-methoxyphenylboronic acid (416 mg, 2.74 mmol, 2.2 eq.), $Cs_2CO_3$ (1.22 g, 3.73 mmol, 3.0 eq. suspended in 2.5 mL $H_2O$), RuPhos Pd G4 (53 mg, 62 *µ*mol, 5.0 mol%) with heating to 80 °C for 5 hours on aluminum block. Purification by silica gel column chromatography using 25% ethyl acetate/hexane solvent mixture as an eluent, gave **S4** (0.312 g, Yield: 65%) as a yellow solid.

**$^1$H-NMR (500 MHz, CDCl$_3$)** δ 8.12 (d, *J* = 8.8 Hz, 2H), 7.75 (d, *J* = 9.3 Hz, 1H), 7.50 – 7.46 (m, 3H), 7.43 (s, 1H), 7.09 – 7.02 (m, 5H), 3.89 (d, *J* = 14.1 Hz, 6H), 3.12 (s, 6H) ppm;

**$^{13}$C-NMR (126 MHz, CDCl$_3$)** δ 160.7, 159.8, 156.8, 151.4, 142.1, 131.4, 130.8, 129.0, 126.5, 118.2, 115.7, 115.5, 114.2, 114.0, 60.5, 55.5, 55.5, 40.6, 21.2, 14.4 ppm;

**FT-IR ($v^{max}$ / cm$^{-1}$, neat):** 1683, 1605, 1516, 1502, 1466, 1431, 1363, 1294, 1242, 1199, 1174, 1127, 1026, 833, 800, 718

**HRMS-ESI (m/z):** calcd for $C_{25}H_{25}N_2O_2$ [M + H]$^+$ 385.1916, found 385.1901;

**M.P (°C):** 120-122

**4,4'-(7-(Dimethylamino)quinoline-2,4-diyl)bis(*N,N*-dimethylaniline) (S5):**

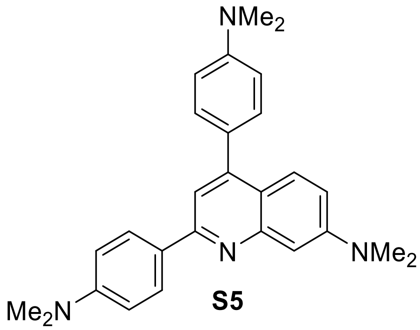

**S5**

Compound **S5** was synthesized using **General Procedure B:** 2,4-dichloro-*N*,*N*-dimethylquinolin-7-amine (0.204 g, 0.846 mmol, 1.0 eq.) in THF (6.8 mL), 4-(dimethylamino)phenylboronic acid (307 mg, 1.86 mmol, 2.2 eq.), $Cs_2CO_3$ (827 mg, 2.54 mmol, 3.0 eq. suspended in 1.7 mL $H_2O$), RuPhos Pd G4 (36 mg, 42 *µ*mol, 5.0 mol%) with heating to 80 °C for 5 hours on aluminum block. Purification by silica gel column chromatography using 40% ethyl acetate/hexane solvent mixture as an eluent, gave **S5** (0.265 g, Yield: 76%) as a yellow solid.

**$^1$H-NMR (500 MHz, $CDCl_3$):** δ 8.04 (d, *J* = 8.6 Hz, 2H), 7.84 (d, *J* = 9.4 Hz, 1H), 7.65 (s, 1H), 7.48 (d, *J* = 8.2 Hz, 2H), 7.31 (s, 1H), 7.03 (dd, *J* = 9.5, 2.4 Hz, 1H), 6.91 (d, *J* = 8.3 Hz, 2H), 6.77 (d, *J* = 8.4 Hz, 2H), 3.17 (s, 6H), 3.10 (s, 6H), 3.06 (s, 6H) ppm;

**$^{13}$C-NMR (126 MHz, $CDCl_3$)** δ 155.6, 153.2, 152.9, 151.9, 151.2, 142.5, 130.8, 129.9, 128.1, 117.8, 117.0, 115.7, 113.4, 112.6, 112.2, 98.0, 40.7, 40.2, 40.0 ppm;

This compound is known, and characterization data is consistent with the literature report.[5]

**6-Methoxy-2,4-bis(4-methoxyphenyl)quinoline (S6):**

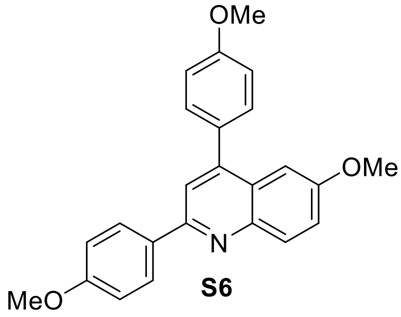


**S6**

Compound **S6** was synthesized using **General Procedure B:** 2,4-dichloro-6-methoxyquinoline (0.100 g, 0.439 mmol, 1.0 eq.) in THF (3.5 mL), 4-methoxyphenylboronic acid (0.147 g, 0.965 mmol, 2.2 eq.), $Cs_2CO_3$ (429 mg, 1.32 mmol, 3.0 eq. suspended in 0.9 mL $H_2O$), RuPhos Pd G4 (19 mg, 22 *µ*mol, 5.0 mol%) with heating to 80 °C for 5 hours on aluminum block. Purification by silica gel column chromatography using 25% ethyl acetate/hexane solvent mixture as an eluent, gave **S6** (0.121 g, Yield: 74%) as a white solid.

**$^1$H-NMR (500 MHz, $CDCl_3$):** δ 8.21 – 8.08 (m, 3H), 7.70 (s, 1H), 7.55 – 7.49 (m, 2H), 7.38 (dd, *J* = 9.1, 2.8 Hz, 1H), 7.22 (d, *J* = 2.8 Hz, 1H), 7.11 – 6.99 (m, 4H), 3.91 (s, 3H), 3.88 (s, 3H), 3.81 (s, 3H) ppm;

**$^{13}$C-NMR (126 MHz, $CDCl_3$):** δ 160.7, 159.8, 157.6, 154.2, 147.8, 144.6, 132.1, 131.1, 131.0, 130.6, 128.8, 126.6, 121.8, 119.3, 114.3, 114.2, 103.9, 55.5, 55.5, 55.5 ppm;

**FT-IR ($v^{max}$ / $cm^{-1}$, neat):** 1606, 1587, 1515, 1501, 1450, 1426, 1401, 1362, 1291, 1246, 1213, 1177, 1121, 1031, 832

**HRMS-ESI (m/z):** calcd for $C_{24}H_{22}NO_3$ $[M + H]^+$ 372.1599, found 372.1594;

**M.P (°C):** 98-100

**2,4-Bis(2,4-dimethoxyphenyl)-7-methoxyquinoline (S7):**

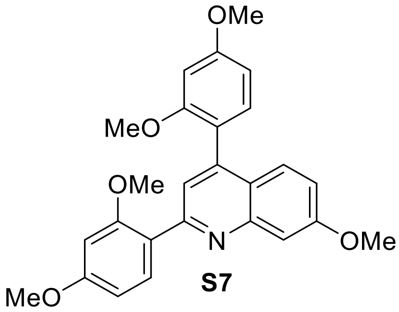

**S7**

Compound **S7** was synthesized using **General Procedure B:** 2,4-dichloro-7-methoxyquinoline (251 mg, 1.10 mmol, 1.0 eq.) in THF (8.8 mL), 2,4-dimethoxyphenylboronic acid (439 mg, 2.41 mmol, 2.2 eq.), $Cs_2CO_3$ (1.07 g, 3.29 mmol, 3.0 eq. suspended in 2.2 mL $H_2O$), RuPhos Pd G4 (46 mg, 55 $\mu$mol, 5.0 mol%) with heating to 80 °C for 5 hours on aluminum block. Purification by silica gel column chromatography using 40% ethyl acetate/hexane solvent mixture as an eluent, gave **S7** (0.418 g, Yield: 88%) as a white solid.

**$^1$H-NMR (500 MHz, $CDCl_3$):** δ 7.85 (d, *J* = 8.5 Hz, 1H), 7.66 (s, 1H), 7.60 – 7.47 (m, 2H), 7.25 (d, *J* = 12.7 Hz, 3H), 7.05 (dd, *J* = 9.2, 2.6 Hz, 1H), 6.75 – 6.62 (m, 3H), 6.57 (d, *J* = 2.3 Hz, 1H), 3.96 (s, 3H), 3.90 (s, 3H), 3.87 (s, 3H), 3.83 (s, 3H), 3.71 (s, 3H) ppm;

**$^{13}$C-NMR (126 MHz, $CDCl_3$):** δ 161.5, 161.2, 160.2, 158.5, 158.1, 156.7, 150.2, 144.4, 132.5, 132.1, 127.4, 123.2, 122.7, 121.5, 120.4, 118.5, 107.7, 105.3, 104.5, 99.0, 98.9, 55.8, 55.6, 55.6 ppm;

**FT-IR ($v^{max}$ / $cm^{-1}$, neat):** 1710, 1605, 1579, 1498, 1449, 1435, 1417, 1354, 1300, 1280, 1255, 1205, 1156, 1130, 1112, 1027, 934, 827, 803

**HRMS-ESI (m/z):** calcd for $C_{26}H_{26}NO_5$ $[M + H]^+$ 432.1811, found 432.1805;.

**M.P (°C):** 102-104

**2,4-Bis(2,4-dimethoxyphenyl)-6,7-dimethoxyquinoline (S8):**

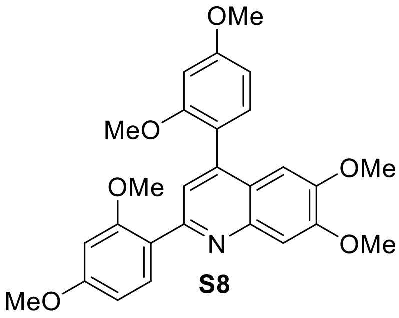

**S8**

Compound **S8** was synthesized using **General Procedure B:** 2,4-dichloro-6,7-dimethoxyquinoline (0.100 g, 0.388 mmol, 1.0 eq.) in THF (3.2 mL), 2,4-dimethoxyphenylboronic acid (0.155 g, 0.852 mmol, 2.2 eq.), $Cs_2CO_3$ (379 mg, 1.16 mmol, 3.0 eq. suspended in 1.0 mL $H_2O$), RuPhos Pd G4 (16 mg, 19 $\mu$mol, 5.0 mol%), with heating to 80 °C for 5 hours on aluminum block. Purification by silica gel column chromatography using 60% ethyl acetate/hexane solvent mixture as an eluent, gave **S8** (0.139 g, Yield: 78%) as a white solid.

**$^{1}$H-NMR (500 MHz, $CDCl_3$):** δ 7.84 (d, *J* = 8.4 Hz, 1H), 7.66 (s, 2H), 7.34 – 7.23 (m, 1H), 6.88 (s, 1H), 6.70 – 6.62 (m, 3H), 6.56 (d, *J* = 2.3 Hz, 1H), 4.05 (s, 3H), 3.91 (s, 3H), 3.87 (s, 3H), 3.83 (s, 6H), 3.74 (s, 3H) ppm;

**$^{13}$C-NMR (126 MHz, $CDCl_3$)** δ 161.3, 158.4, 157.9, 152.1, 149.2, 132.4, 132.08 125.7, 123.1, 121.7, 120.4, 118.9, 108.5, 105.3, 104.7, 104.1, 99.0, 99.0, 56.3, 55.9, 55.8, 55.6 ppm;

**FT-IR ($v^{max}$ / $cm^{-1}$, neat):** 1709, 1605, 1577, 1497, 1481, 1460, 1431, 1407, 1354, 1297, 1279, 1241, 1204, 1156, 1132, 1112, 1070, 1027, 1008, 934, 923, 869, 830, 808, 752, 727

**HRMS-ESI (m/z):** calcd for $C_{27}H_{28}NO_6$ $[M + H]^+$ 462.1916, found 462.1911;

**M.P (°C):** 130-132

**Synthesis of C-2 arylated quinolinium compound *via* regioselective Suzuki−Miyaura cross-coupling reaction**

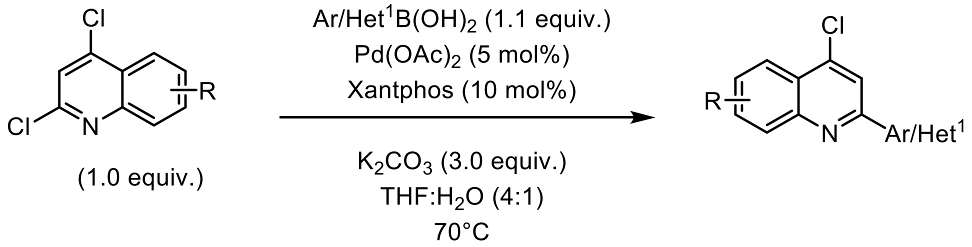


*C-2 Regioselective Suzuki−Miyaura cross-coupling reaction*

**General Procedure C:**

Synthesis of C-2 substituted quinolinium compound was achieved by adapting the procedure from Chenoweth,[5] as follows:

A 20 mL vial Chemglass Life Sciences (CG-4920-02) equipped with a stir bar was dried under vacuum, allowed to cool to room temperature, and refilled with $N_2$. The vial was charged with 2,4-dichloro arylquinoline (1.0 eq.) and then dissolved with dry degassed THF and stirred vigorously. The vial was then added with $Pd(OAc)_2$ (5 mol %) and the Xantphos ligand (10 mol %) was then added to the vial while stirring. To the vial, $K_2CO_3$ (3.0 eq.) dissolved in 2 mL of $ddH_2O$ was finally added. The vial was capped with 20mm Aluminum seal with PTFE-faced Silicone septa. The reaction mixture was stirred for 12 hours at 70 ˚C. After completion, the solvent was evaporated using rotary evaporator and the residue was extracted EtOAc (3 x 20 mL). The organic layers were combined, washed with brine, dried over anhydrous $Na_2SO_4$, filtered, and concentrated using a rotary evaporator. The crude mixture was purified by silica gel column chromatography.

**4-Chloro-2-(4-methoxyphenyl)quinoline (S9):**

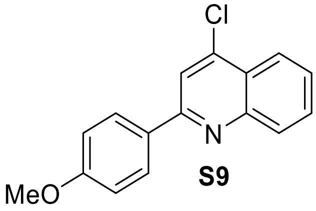


**S9**

Compound **S9** was synthesized using **General Procedure C:** 2,4-dichloroquinoline (200 mg, 1.01 mmol, 1.0 eq.) in THF (8.1 mL), 4-methoxyphenylboronic acid (169 mg, 1.11 mmol, 1.1 eq.), $Pd(OAc)_2$ (11 mg, 51 $\mu$mol, 5.0 mol%), Xantphos (58 mg, 0.11 mmol, 10.0 mol%), potassium carbonate (419 mg, 3.03 mmol, 3.0 eq. suspended in 2.0 mL $H_2O$) with heating to 70 °C for 12 hours on aluminum block. Purification by silica gel column chromatography using 15% ethyl acetate/hexane solvent mixture as an eluent, gave **S9** (0.216 g, Yield: 79%) as a white solid.

**$^1$H NMR-(500 MHz, $CDCl_3$):** δ 8.22 – 8.15 (m, 2H), 8.15 – 8.08 (m, 2H), 7.93 (s, 1H), 7.76 (ddd, *J* = 8.4, 6.9, 1.4 Hz, 1H), 7.59 (ddd, *J* = 8.2, 6.9, 1.2 Hz, 1H), 7.13 – 7.01 (m, 2H), 3.89 (s, 3H) ppm;

**$^{13}$C NMR-(126 MHz, $CDCl_3$):** δ 161.4, 156.8, 148.9, 143.3, 130.7, 129.7, 129.1, 129.0, 127.0, 125.1, 124.0, 118.7, 114.4, 55.5 ppm;

This compound is known, and characterization data is consistent with the literature report.[6]

**4-Chloro-2-(4-(trifluoromethyl)phenyl)quinoline (S10):**

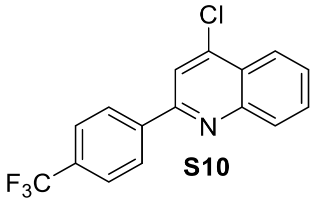


**S10**

Compound **S10** was synthesized using **General Procedure C:** 2,4-dichloroquinoline (251 mg, 1.26 mmol, 1.0 eq.) in THF (10 mL), 4-(trifluoromethyl)phenylboronic acid (264 mg, 1.39 mmol, 1.1 eq.), $Pd(OAc)_2$ (14 mg, 63 $\mu$mol, 5.0 mol%), Xantphos (73 mg, 0.13 mmol, 10.0 mol%), $K_2CO_3$ (523 mg, 3.79 mmol, 3.0 eq. suspended in 2.5 mL $H_2O$) with heating to 70 °C for 12 hours on aluminum block. Purification by silica gel column chromatography using 10% ethyl acetate/hexane solvent mixture as an eluent, gave **S10** (0.238 g, Yield: 61%) as a white solid.

**$^1$H-NMR (400 MHz, $CDCl_3$):** δ 8.10 (dd, *J* = 8.9, 1.3 Hz, 1H), 7.91 – 7.72 (m, 4H), 7.62 (d, *J* = 8.0 Hz, 2H), 7.54 (ddd, *J* = 8.2, 6.9, 1.3 Hz, 1H), 7.34 (s, 1H) ppm;

**$^{13}$C-NMR (126 MHz, $CDCl_3$)** δ 150.3, 150.1, 148.4, 140.4, 131.6, 131.4, 131.1, 130.9, 129.9, 129.2, 127.5, 125.9, 125.8, 125.8, 125.8, 125.5, 125.3, 122.2 ppm;

This compound is known, and characterization data are consistent with the literature report.[7]

**2-(Benzofuran-2-yl)-4-chloroquinoline (S11):**

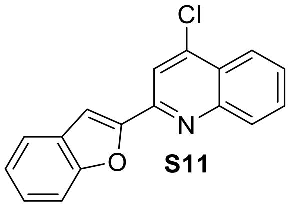


**S11**

Compound **S11** was synthesized using **General Procedure C:** 2,4-dichloroquinoline (504 mg, 2.52 mmol, 1.0 eq.) in THF (20.2 mL), benzofuran-2-boronic acid (450 mg, 2.78 mmol, 1.1 eq.), $Pd(OAc)_2$ (28 mg, 13 $\mu$mol, 5.0 mol%), Xantphos (146 mg, 253 $\mu$mol, 10.0 mol%), $K_2CO_3$ (1.05 g,

7.57 mmol, 3.0 eq. suspended in 5.1 mL $H_2O$) with heating to 70 °C for 12 hours on aluminum block. Purification by silica gel column chromatography using 15% ethyl acetate/hexane solvent mixture as an eluent, gave **S11** (0.397 g, Yield: 56%) as a yellow solid.

**$^1$H-NMR (500 MHz, $CDCl_3$):** δ 8.27 – 8.18 (m, 2H), 8.15 (d, *J* = 1.2 Hz, 1H), 7.80 (ddd, *J* = 8.4, 7.0, 1.4 Hz, 1H), 7.70 (dd, *J* = 8.0, 1.2 Hz, 1H), 7.67 – 7.58 (m, 2H), 7.40 (ddd, *J* = 8.4, 7.2, 1.3 Hz, 1H), 7.30 (ddd, *J* = 7.9, 7.3, 0.9 Hz, 1H) ppm;

**$^{13}$C-NMR (126 MHz, $CDCl_3$):** δ 155.8, 151.3, 148.9, 143.6, 131.1, 129.8, 128.7, 127.8, 126.1, 125.9, 124.2, 123.6, 122.2, 118.3, 111.9 ppm;

**FT-IR ($v^{max}$ / $cm^{-1}$, neat):** 1582, 1543, 1491, 1448, 1402, 1310, 1247, 1193, 1161, 1146, 1059, 976, 837, 820, 754, 737, 699

**HRMS-ESI (m/z):** calcd for $C_{17}H_{10}ClNO$ $[M + H]^+$ 280.0529, found 280.0522;

**M.P (°C):** 92-94

**4-Chloro-2-(thiophen-2-yl)quinoline (S12):**

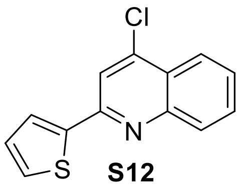


**S12**

Compound **S12** was synthesized using **General Procedure C:** 2,4-dichloroquinoline (210 mg, 1.06 mmol, 1.0 eq.) in THF (8.5 mL), thiophen-2-ylboronic acid (149 mg, 1.17 mmol, 1.1 eq.), $Pd(OAc)_2$ (12 mg, 53 $\mu$mol, 5.0 mol%), Xantphos (61 mg, 0.17 mmol, 10.0 mol%), $K_2CO_3$ (440 mg, 3.18 mmol, 3.0 eq. suspended in 2.1 mL $H_2O$) with heating to 70 °C for 12 hours on aluminum block. Purification by silica gel column chromatography using 10% ethyl acetate/hexane solvent mixture as an eluent, gave **S12** (0.167 g, Yield: 64%) as a yellow solid.

**$^1$H-NMR (400 MHz, $CDCl_3$):** δ 8.21 (dd, *J* = 8.3, 1.4 Hz, 1H), 8.05 (dt, *J* = 8.6, 0.8 Hz, 1H), 7.83 – 7.78 (m, 1H), 7.67 (ddd, *J* = 8.3, 7.0, 1.2 Hz, 1H), 7.53 (s, 1H), 7.26 (s, 3H) ppm;

**$^{13}$C-NMR (126 MHz, $CDCl_3$):** δ 150.0, 148.3, 144.5, 131.7, 129.1, 128.0, 125.3, 124.3, 122.1 ppm;

This compound is known, and characterization data is consistent with the literature report.[7]

**4-Chloro-2-(1-methyl-1*H*-pyrazol-4-yl)quinoline (S13):**

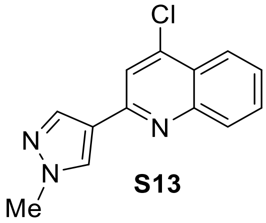


**S13**

Compound **S13** was synthesized using **General Procedure C:** 2,4-dichloroquinoline (309 mg, 1.56 mmol, 1.0 eq.) in THF (8.5 mL),1-methylpyrazole-4-boronic acid pinacol ester (357 mg, 1.72

mmol, 1.1 eq.), $Pd(OAc)_2$ (18 mg, 78 *µ*mol, 5.0 mol%), Xantphos (90 mg, 0.16 mmol, 10.0 mol%), $K_2CO_3$ (647 mg, 4.68 mmol, 3.0 eq. suspended in 2.1 mL $H_2O$) with heating to 70 °C for 12 hours on aluminum block. Purification by silica gel column chromatography using 10% ethyl acetate/hexane solvent mixture as an eluent, gave **S13** (0.226 g, Yield: 59%) as a yellow solid.

**$^1$H-NMR (500 MHz, $CDCl_3$):** δ 8.17 (ddd, *J* = 8.4, 1.5, 0.6 Hz, 2H), 8.08 (d, *J* = 0.8 Hz, 2H), 7.74 (ddd, *J* = 8.4, 6.9, 1.5 Hz, 1H), 7.69 (s, 1H), 7.56 (ddd, *J* = 8.2, 6.9, 1.2 Hz, 1H), 4.00 (s, 3H) ppm;

**$^{13}$C-NMR (126 MHz, $CDCl_3$):** δ 151.8, 149.1, 142.9, 138.3, 130.6, 129.8, 129.2, 126.6, 125.0, 124.0, 123.1, 118.7, 39.4, 24.9 ppm;

This compound is known, and characterization data is consistent with the literature report.[7]

**4-Chloro-2-(4-(trifluoromethyl)phenyl)quinoline (S14):**

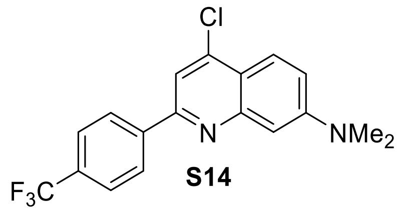


**S14**

Compound **S14** was synthesized using **General Procedure C:** 2,4-dichloro-*N*,*N*-dimethylquinolin-7-amine (121 mg, 498 µmol, 1.0 eq.) in THF (3.8 mL), 4-(trifluoromethyl)phenylboronic acid (104 mg, 547 µmol, 1.1 eq.), $Pd(OAc)_2$ (5 mg, 3 *µ*mol, 5.0 mol%), Xantphos (27 mg, 47 *µ*mol, 10.0 mol%), $K_2CO_3$ (195 mg, 1.41 mmol, 3.0 eq. suspended in 1.0 mL $H_2O$) with heating to 70 °C for 12 hours on aluminum block. Purification by silica gel column chromatography using 10% ethyl acetate/hexane solvent mixture as an eluent, gave **S14** (157 mg, Yield: 90%) as a red solid.

**$^1$H-NMR (500 MHz, $CDCl_3$):** δ 8.25 (d, *J* = 8.1 Hz, 2H), 8.12 (d, *J* = 9.6 Hz, 1H), 7.97 (s, 1H), 7.87 (d, *J* = 8.1 Hz, 2H), 7.51 (s, 1H), 7.35 (dd, *J* = 9.6, 2.5 Hz, 1H), 3.28 (s, 6H) ppm;

**$^{13}$C-NMR (126 MHz, $CDCl_3$):** δ 154.6, 152.0, 150.4, 143.2, 134.1, 129.6, 128.0, 126.7, 126.6, 126.2, 119.3, 119.0, 114.9, 97.9, 43.7, 40.6 ppm;

This compound is known, and characterization data is consistent with the literature report.[5]

**Synthesis of C-4 substituted quinolinium compound *via* regioselective Suzuki−Miyaura cross-coupling reaction.**

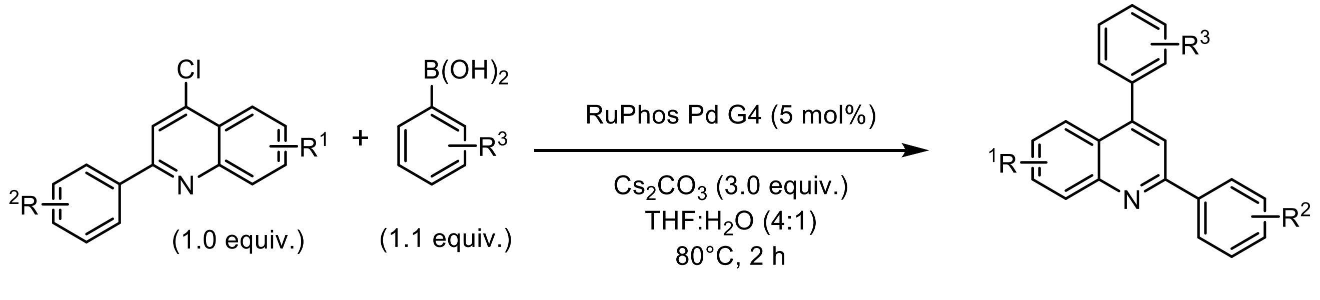


***C-4 substituted Suzuki−Miyaura cross-coupling reaction***

**General Procedure D:**

Synthesis of C-4 substituted quinolinium compound was achieved as follows:

A 20 mL vial Chemglass Life Sciences (CG-4920-02) equipped with a stir bar was dried under vacuum, allowed to cool to room temperature, and refilled with $N_2$. The vial was charged with C-2 substituted arylquinoline (**S9-S14**, 1.0 eq.) and then dissolved with dry degassed THF and stirred vigorously. The vial was then added with RuPhos Pd G4 (5 mol%) and boronic acid (1.1 eq.) was then added to the vial while stirring. To the vial, $Cs_2CO_3$ (3.0 eq.) dissolved in 1 mL of dd$H_2O$ was finally added. The vial was capped with 20mm Aluminum seal with PTFE-faced Silicone septa. The reaction mixture was stirred for 2 hours at 80 ˚C. After completion, the solvent was evaporated using a rotary evaporator and the residue was extracted EtOAc (3 x 20 mL). The organic layers were combined, washed with brine (50 mL), dried over anhydrous $Na_2SO_4$, filtered, and concentrated using a rotary evaporator. The crude mixture was purified by silica gel column chromatography. This method enabled the preparation of sufficient amounts of C-2 and C-4 substituted quinolinium compounds bearing acceptor and donor properties for *N*-alkylation.

**2-(4-Methoxyphenyl)-4-(4-(trifluoromethyl)phenyl)quinoline (S15):**

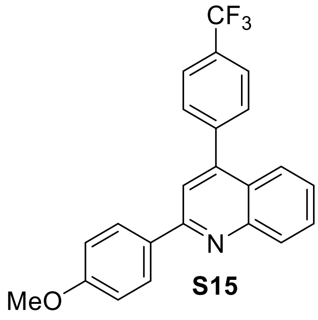


**S15**

Compound **S15** was synthesized using **General Procedure D:** 4-chloro-2-(4-methoxyphenyl)quinoline **S9** (0.156 g, 0.578 mmol, 1.0 eq.) in THF (4.6 mL), 4-(trifluoromethyl)phenylboronic acid (0.121 g, 0.636 mmol, 1.1 eq.), $Cs_2CO_3$ (565 mg, 1.74 mmol, 3.0 eq. suspended in 1.2 mL $H_2O$), RuPhos Pd G4 (25 mg, 29 $\mu$mol, 5.0 mol%,) with heating to 80 °C for 2 hours on aluminum block. Purification by silica gel column chromatography using 15% ethyl acetate/hexane solvent mixture as an eluent, gave **S15** (0.191 g, Yield: 87%) as a white solid.

**$^1$H-NMR (500 MHz, $CDCl_3$):** δ 8.19 – 8.17 (m, 3H), 7.83 – 7.49 (m, 7H), 7.48 (t, *J* = 7.6 Hz, 1H), 7.06 (d, *J* = 8.8 Hz, 2H), 3.90 (s, 3H) ppm;

**$^{13}$C-NMR (126 MHz, $CD_3CN$):** δ 162.5, 156.7, 149.5, 148.4, 142.9, 131.4, 131.3, 131.1, 130.8, 130.2, 129.7, 127.9, 126.6, 126.6, 126.5, 126.5, 126.3, 126.0, 124.4, 120.1, 115.3, 56.2 ppm;

**$^{19}$F-NMR (471 MHz, $CD_3CN$):** δ -63.09 (s, 3F) ppm;

**FT-IR ($v^{max}$ / $cm^{-1}$, neat):** 2917, 2849, 1598, 1501, 1464, 1358, 1323, 1248, 1198, 1172, 1126, 1109, 1066, 1031, 1018, 836, 765

**HRMS-ESI (m/z):** calcd for $C_{23}H_{17}F_3NO$ $[M + H]^+$ 380.1262, found 380.1257;

**M.P (°C):** 108-110

**4-(4-methoxyphenyl)-2-(4-(trifluoromethyl)phenyl)quinoline (S16):**

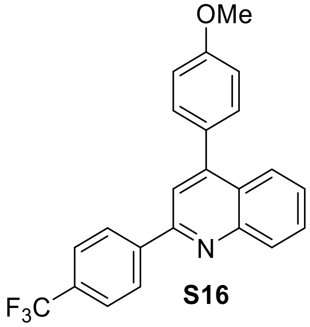

**S16**

Compound **S16** was synthesized using **General Procedure D:** 4-chloro-2-(4-(trifluoromethyl)phenyl)quinoline **S10** (0.200 g, 0.742 mmol, 1.0 eq.) in THF (5.2 mL), 4-methoxyphenylboronic acid (0.155 mg, 0.816 mmol, 1.1 eq.), $Cs_2CO_3$ (635 mg, 1.95 mmol, 3.0 eq. suspended in 1.3 mL $H_2O$), RuPhos Pd G4 (28 mg, 33 $\mu$mol, 5.0 mol%) with heating to 80 °C for 2 hours on aluminum block. Purification by silica gel column chromatography using 20% ethyl acetate/hexane solvent mixture as an eluent, gave **S16** (0.205 g, Yield: 73%) as a white solid.

**$^{1}$H-NMR (500 MHz, $CDCl_3$):** δ 8.31 – 8.09 (m, 3H), 7.84 – 7.56 (m, 7H), 7.47 (t, *J* = 7.5 Hz, 1H), 7.15 – 6.93 (m, 2H), 3.89 (d, *J* = 1.7 Hz, 3H) ppm;

**$^{13}$C-NMR (126 MHz, $CDCl_3$):** δ 161.0, 156.4, 148.8, 147.5, 142.2, 131.9, 130.7, 130.5, 130.1, 130.0, 129.8, 126.4, 125.7, 125.6, 125.6, 125.6, 125.1, 125.1, 118.8, 114.3, 55.4 ppm;

**$^{19}$F-NMR (471 MHz, $CD_3CN$):** δ -62.46 (s, 3F) ppm;

**FT-IR ($v^{max}$ / $cm^{-1}$, neat):** 1594, 1546, 1517, 1499, 1358, 1323, 1289, 1251, 1170, 1125, 1067, 1031, 1018, 834, 764

**HRMS-ESI (m/z):** calcd for $C_{23}H_{17}F_3NO$ $[M + H]^+$ 380.1262, found 380.1257;

**M.P (°C):** 110-112

**9-(2-(4-(Trifluoromethyl)phenyl)quinolin-4-yl)-2,3,6,7-tetrahydro-1*H*,5*H*-pyrido[3,2,1-ij]quinoline (S17):**

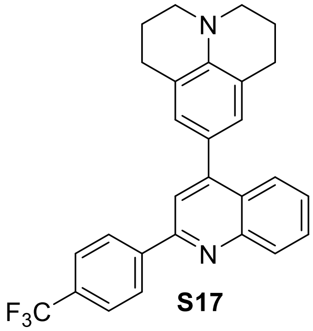

**S17**

Compound **S17** was synthesized using **General Procedure D:** 4-Chloro-2-(4-(trifluoromethyl)phenyl)quinoline **S10** (98 mg, 0.32 mmol, 1.0 eq.) in THF (2.6 mL), 4-julolidine boronic acid (0.103 g, 0.350 mmol, 1.1 eq.), $Cs_2CO_3$ (0.311 g, 0.596 mmol, 3.0 eq. suspended in 1 mL $H_2O$), RuPhos Pd G4 (14 mg, 16 $\mu$mol, 5.0 mol%,) with heating to 80 °C for 2 hours on aluminum block. Purification by silica gel column chromatography using 30% ethyl acetate/hexane solvent mixture as an eluent, gave **S17** (0.108 g, 76%) as a yellow solid.

**$^{1}$H-NMR (400 MHz, $CDCl_3$):** δ 8.18 (s, 1H), 7.81 (d, *J* = 8.0 Hz, 2H), 7.76 – 7.62 (m, 7H), 7.40 (t, *J* = 7.6 Hz, 1H), 3.24 (t, *J* = 5.7 Hz, 4H), 2.86 (t, *J* = 6.4 Hz, 4H), 2.11 – 1.93 (m, 4H) ppm;

**$^{13}$C-NMR (151 MHz, $CD_3CN$):** δ 158.3, 150.3, 148.3, 146.0, 144.2, 131.9, 131.2, 130.9, 127.5, 127.2, 127.0, 127.0, 127.0, 127.0, 126.6, 126.1, 122.8, 119.8, 51.1, 29.1, 23.2 ppm;

**$^{19}$F-NMR (471 MHz, $CD_3CN$):** δ -62.97 (s, 3F) ppm;

**FT-IR ($v^{max}$ / $cm^{-1}$, neat):** 2924, 2853, 1682, 1591, 1544, 1515, 1435, 1355, 1324, 1271, 1182, 1167, 1127, 1066, 846, 764

**HRMS-ESI (m/z):** calcd for $C_{28}H_{24}F_3N_2$ $[M + H]^+$ 445.1891, found 445.1885;

**M.P (°C):** 84-86

**2-(Benzofuran-2-yl)-4-(2,4-dimethoxyphenyl)quinoline (S18)**

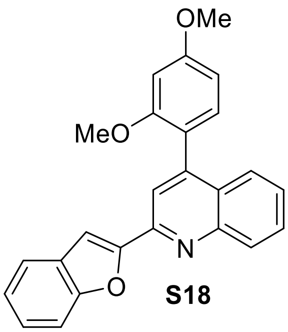


**S18**

Compound **S18** was synthesized using **General Procedure D:** 2-(benzofuran-2-yl)-4-chloroquinoline **S11** (0.120 g, 0.429 mmol, 1.0 eq.) in THF (3.4 mL), 2,4-dimethoxyphenylboronic acid (86 mg, 0.47 mmol, 1.1 eq.), $Cs_2CO_3$ (419 mg, 1.29 mmol, 3.0 eq. suspended in 0.9 mL $H_2O$), RuPhos Pd G4 (18 mg, 22 *μ*mol, 5.0 mol%) with heating to 80 °C for 2 hours on aluminum block. Purification by silica gel column chromatography using 25% ethyl acetate/hexane solvent mixture as an eluent, gave **S18** (0.134 g, Yield: 82%) as a yellow solid.

**$^{1}$H-NMR (500 MHz, $CDCl_3$):** δ 8.62 (d, *J* = 8.5 Hz, 1H), 8.30 (s, 1H), 8.15 (s, 1H), 7.88 (t, *J* = 7.7 Hz, 1H), 7.82 – 7.73 (m, 2H), 7.59 (dd, *J* = 8.0, 4.1 Hz, 2H), 7.45 (d, *J* = 7.7 Hz, 1H), 7.41 – 7.21 (m, 3H), 6.80 – 6.59 (m, 2H), 3.94 (s, 3H), 3.75 (s, 3H) ppm;

**$^{13}$C-NMR (126 MHz, $CDCl_3$):** δ 162.6, 157.9, 156.2, 153.1, 149.1, 144.8, 142.5, 132.9, 131.9, 128.5, 128.0, 127.7, 127.3, 127.2, 124.6, 124.1, 123.2, 119.9, 117.7, 112.5, 111.9, 105.2, 99.1, 55.8, 55.7 ppm;

**FT-IR ($v^{max}$ / $cm^{-1}$, neat):** 1669, 1638, 1596, 1506, 1465, 1439, 1409, 1376, 1337, 1305, 1271, 1241, 1207, 1159, 1143, 1081, 1027, 979, 942, 887, 828, 798, 752, 718, 705

**HRMS-ESI (m/z):** calcd for $C_{25}H_{19}NO_3$ $[M + H]^+$ 382.1443, found 382.1438;

**M.P (°C):** 114-116

**4-(2,4-Dimethoxyphenyl)-2-(1-methyl-1*H*-pyrazol-4-yl)quinoline (S19):**

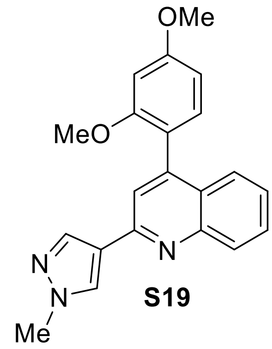


**S19**

Compound **S19** was synthesized using **General Procedure D:** 4-chloro-2-(1-methyl-1*H*-pyrazol-4-yl)quinoline **S13** (0.121 g, 0.492 mmol, 1.0 eq.) in THF (3.9 mL), 2,4-dimethoxyphenylboronic acid (99 mg, 0.54 mmol, 1.1 eq.), $Cs_2CO_3$ (481 mg, 1.48 mmol, 3.0 eq. suspended in 1.0 mL $H_2O$), RuPhos Pd G4 (21 mg, 25 *μ*mol, 5.0 mol%) with heating to 80 °C for 2 hours on aluminum block. Purification by silica gel column chromatography using 20% ethyl acetate/hexane solvent mixture as an eluent, gave **S19** (116 mg, Yield: 68%) as a white solid.

**$^1$H NMR (500 MHz, $CDCl_3$):** δ 8.22 – 7.99 (m, 3H), 7.72 – 7.42 (m, 3H), 7.36 (ddd, *J* = 8.2, 6.8, 1.3 Hz, 1H), 7.22 (d, *J* = 8.8 Hz, 1H), 6.77 – 6.45 (m, 2H), 3.99 (s, 3H), 3.91 (s, 3H), 3.71 (s, 3H) ppm;

**$^{13}$C NMR (126 MHz, $CDCl_3$):** δ 161.6, 158.1, 138.5, 131.9, 126.7, 126.4, 125.5, 120.2, 119.7, 104.8, 99.1, 60.6, 55.7, 55.7, 39.4, 21.2, 14.4 ppm;

**FT-IR ($v^{max}$ / $cm^{-1}$, neat):** 1598, 1564, 1511, 1495, 1463, 1438, 1413, 1304, 1281, 1255, 1207, 1158, 1133, 1116, 1031, 998, 830, 766

**HRMS-ESI (m/z):** calcd for $C_{21}H_{20}N_3O_2$ $[M + H]^+$ 346.1555, found 346.1549;

**M.P (°C):** 96-98

**4-(2,4-Dimethoxyphenyl)-2-(thiophen-2-yl)quinoline (S20)**

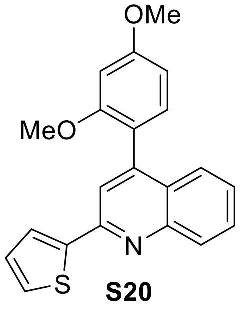


**S20**

Compound **S20** was synthesized using **General Procedure D:** 4-chloro-2-(thiophen-2-yl)quinoline **S12** (0.160 g, 0.651 mmol, 1.0 eq.) in THF (5.2 mL), 2,4-dimethoxyphenylboronic acid (0.130 mg, 0.716 mmol, 1.1 eq.), $Cs_2CO_3$ (636 mg, 1.95 mmol, 3.0 eq. suspended in 1.3 mL $H_2O$), RuPhos Pd G4 (28 mg, 33 *μ*mol, 5.0 mol%,) with heating to 80 °C for 2 hours on aluminum block.

Purification by silica gel column chromatography using 20% ethyl acetate/hexane solvent mixture as an eluent, gave **S20** (0.158 g, Yield: 70%) as a yellow solid.

**$^{1}$H-NMR (500 MHz, $CDCl_3$):** δ 7.98 – 8.00 (m, 1H), 7.65 – 7.55 (m, 3H), 7.39 – 7.40 (m, 1H), 7.26 – 7.11 (m, 3H), 6.57 - 6.59 (m, 3H), 3.84 (s, 3H), 3.65 (s, 3H) ppm;

**$^{13}$C-NMR (126 MHz, $CDCl_3$):** δ 161.8, 157.9, 150.4, 149.1, 148.1, 131.8, 130.2, 128.8, 126.7, 126.6, 126.5, 123.3, 118.3, 104.7, 99.0, 55.6, 55.6 ppm;

**FT-IR ($v^{max}$ / $cm^{-1}$, neat):** 1660, 1587, 1540, 1437, 1324, 1203, 1133, 837, 720

**HRMS-ESI (m/z):** calcd for $C_{21}H_{18}NO_2S$ $[M + H]^+$ 348.1058, found 348.1051;

**M.P (°C):** 80-82

**4-(4-(Dimethylamino)phenyl)-N,N-dimethyl-2-(4-(trifluoromethyl)phenyl)quinolin-7-amine (S21):**

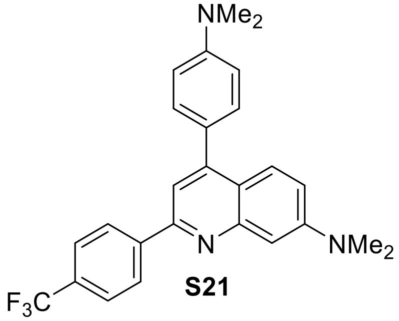

**S21**

Compound **S21** was synthesized using **General Procedure D:** 4-chloro-2-(4-(trifluoromethyl)phenyl)quinoline **S14** (0.150 g, 0.428 mmol, 1.0 eq.) in THF (3.4 mL), 4-(dimethylamino)phenylboronic acid (78 mg, 0.47 mmol, 1.1 eq.), $Cs_2CO_3$ (418 mg, 1.28 mmol, 3.0 eq. suspended in 0.9 mL $H_2O$), RuPhos Pd G4 (18 mg, 21 $\mu$mol, 5.0 mol%) with heating to 80 °C for 2 hours on aluminum block. Purification by silica gel column chromatography using 25% ethyl acetate/hexane solvent mixture as an eluent, gave **S21** (0.166 g, Yield: 89%) as a yellow solid.

**$^{1}$H-NMR (500 MHz, $CDCl_3$)** δ 8.12 (d, *J* = 8.8 Hz, 2H), 7.75 (d, *J* = 9.3 Hz, 1H), 7.50 – 7.46 (m, 3H), 7.43 (s, 1H), 7.09 – 7.02 (m, 5H), 3.89 (d, *J* = 14.1 Hz, 6H), 3.12 (s, 6H) ppm;

**$^{13}$C-NMR (126 MHz, $CDCl_3$)** δ 160.9, 159.9, 151.5, 131.4, 130.9, 129.1, 126.6, 118.3, 115.8, 115.5, 114.3, 114.1, 60.6, 55.6, 55.6, 40.7, 21.2, 14.4 ppm;

**FT-IR ($v^{max}$ / $cm^{-1}$, neat):** 2924, 1609, 1577, 1524, 1507, 1484, 1430, 1401, 1357, 1322, 1208, 1164, 1123, 1072, 1015, 846, 819

**HRMS-ESI (m/z):** calcd for $C_{25}H_{25}N_2O_2$ $[M + H]^+$ 385.1916, found 385.1901;

**M.P (°C):** 104-106

##### 3.6 Synthesis of *N*-methylated quinolinium salts *via* $S_N2$:

**General Procedure E:**

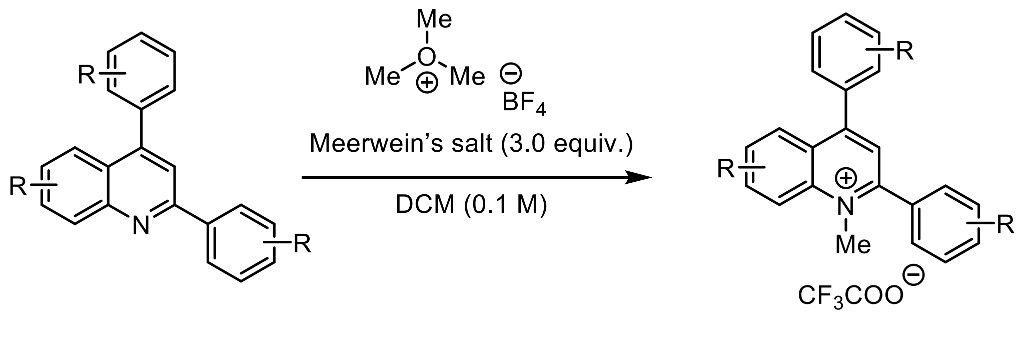


A round-bottom flask equipped with a stir bar and a side arm inlet adapter was heat dried under vacuum, allowed to cool to room temperature, and refilled with $N_2$. The flask was charged with trimethyloxonium tetrafluoroborate salt (3.0 eq.) and the flask was evacuated and refilled with $N_2$. The flask was chilled on an ice bath. To the flask was added dropwise a 0.1 M solution of the desired diarylquinoline compound (1.0 eq.) in dry DCM *via* syringe. The reaction mixture, which turns heterogenous shortly after addition of the quinoline, was allowed to stir at room temperature for 12 hours. After the completion of reaction, the solution was concentrated using a rotary evaporator. Diethyl ether (10 mL) was added to the residue and the suspension was stirred vigorously at room temperature for 30 min. The mixture was allowed to stand for 5 min, subsequently, the solvent was decanted carefully. In order to obtain an analytically pure compound, the residue was redissolved in acetonitrile and water (4:1). The residue was then purified with reverse-phased flash chromatography to yield 2,4-diaryl-*N*-methyl-quinolinium aromatic cation**.**

**General Procedure F:**

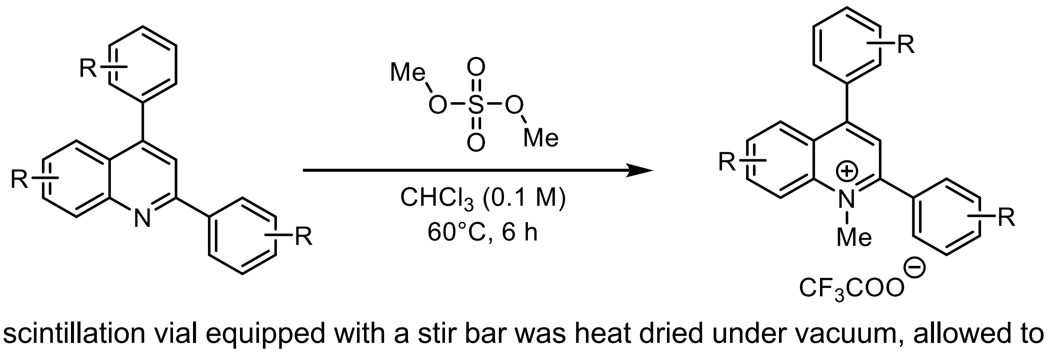


A 10 mL scintillation vial equipped with a stir bar was heat dried under vacuum, allowed to cool to room temperature, and refilled with $N_2$. The vial was charged with the desired quinoline compound (1.0 eq.) followed by addition of anhydrous chloroform (0.1 M) at room temperature and the resultant mixture was stirred. To the vial was added dimethyl sulfate (3.0 eq.) *via* syringe and the resulting solution was stirred for 30 minutes. The resulting mixture then stirred to 60ºC for 6 hrs on an aluminum heating block. After the completion of reaction, the solution was concentrated using a rotary evaporator before being redissolved in acetonitrile and water (4:1). The residue was then purified with reverse-phased flash chromatography to yield 2,4-diaryl-*N*-methyl-quinolinium aromatic cation**.**

#### Alkylation conditions used for each *N*-methylquinolinium sensitizer (3-17):

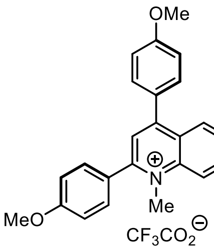

**3, 67%**

*From General Procedure E*

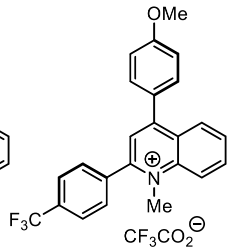

**4, 71%**

*From General Procedure E*

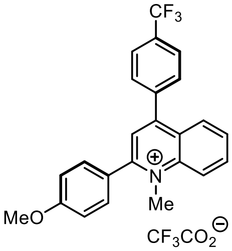

**5, 63%**

*From General Procedure E*

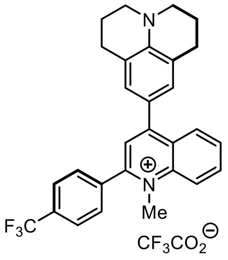

**6, 52%**

*From General Procedure F*

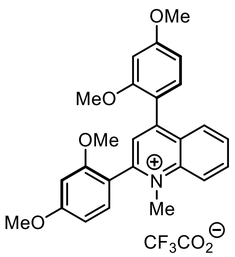

**7, 57%**

*From General Procedure E*

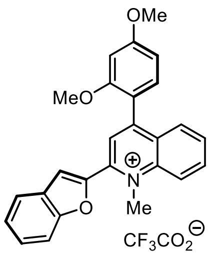

**8, 30%**

*From General Procedure F*

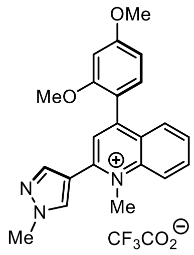

**9, 20%**

*From General Procedure F*

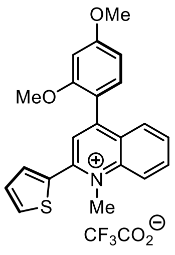

**10, 21%**

*From General Procedure F*

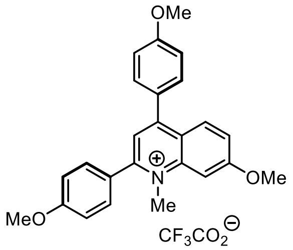

**11, 63%**

*From General Procedure E*

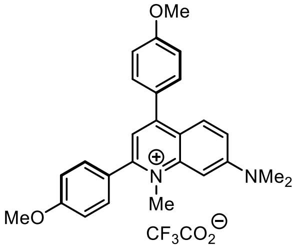

**12, 74%**

*From General Procedure E*

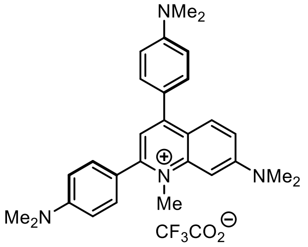

**13, 56%**

*From General Procedure F*

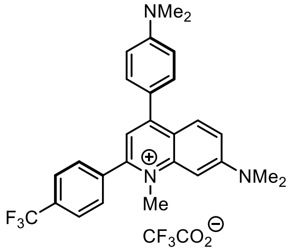

**14, 16%**

*From General Procedure F*

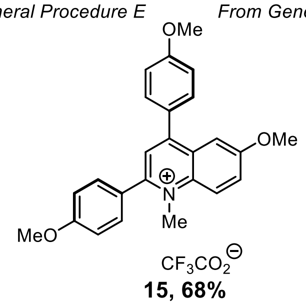

**15, 68%**

*From General Procedure E*

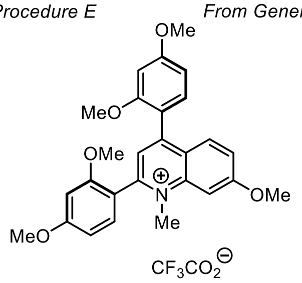

**16, 78%**

*From General Procedure E*

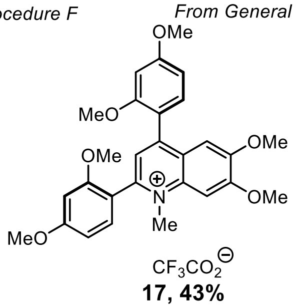

**17, 43%**

*From General Procedure E*

**2,4-Bis(4-methoxyphenyl)-1-methylquinolin-1-ium 2,2,2-trifluoroacetate (3):**

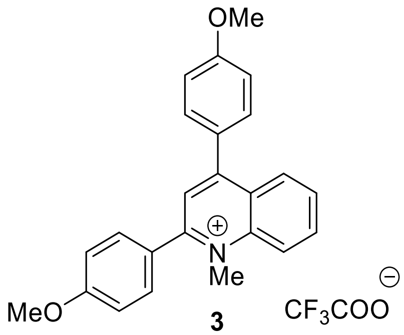

**3**

Salt **3** was synthesized using general procedure E using trimethyloxonium tetrafluoroborate (73 mg, 3.0 eq., 0.49 mmol) and solution of 2,4-bis(4-methoxyphenyl)quinoline **S1** in anhydrous DCM (56 mg, 0.16 mmol, 1.0 eq., 1.6 mL $CH_2Cl_2$). After completion of reaction, the resulting mixture was then concentrated using a rotary evaporator and the residue was washed with cold diethyl ether. The residue was redissolved in acetonitrile and water (4:1) and purified by reverse-phased flash chromatography Method A followed by lyophilized gave **3** (39 mg, Yield: 67%) as a yellow solid.

**$^1$H NMR (500 MHz, $CD_3CN$):** δ 8.44 (d, *J* = 9.0 Hz, 1H), 8.35 (dd, *J* = 8.5, 1.4 Hz, 1H), 8.24 (ddd, *J* = 8.8, 7.0, 1.5 Hz, 1H), 7.96 (ddd, *J* = 8.2, 7.0, 1.0 Hz, 1H), 7.91 (s, 1H), 7.69 (tt, *J* = 9.7, 2.5 Hz, 4H), 7.40 – 7.12 (m, 4H), 4.37 (s, 3H), 3.93 (d, *J* = 5.0 Hz, 6H) ppm;

**$^{13}$C NMR (126 MHz, $CDCl_3$):** δ 163.5, 163.0, 160.3, 159.9, 158.3, 141.7, 136.2, 133.0, 132.8, 130.5, 129.9, 128.1, 127.8, 126.3, 126.0, 120.7, 118.3, 115.8, 115.7, 56.6, 56.4, 43.4 ppm;

**$^{19}$F NMR (471 MHz, $CD_3CN$):** δ -76.62 (s, 3F) ppm;

**FT-IR ($\nu^{max}$ / $cm^{-1}$, neat):** 1735, 1593, 1507, 1461, 1436, 1388, 1361, 1297, 1245, 1171, 1129, 1020, 925, 835, 789, 764, 703

**HRMS-ESI (m/z):** calcd for $C_{24}H_{22}NO_2^+$ [M - $CF_3COO$]$^+$ 356.1645, found 356.1644;

**M.P (°C):** 42-44

**4-(4-Methoxyphenyl)-1-methyl-2-(4-(trifluoromethyl)phenyl)quinolin-1-ium 2,2,2-trifluoroacetate (4):**

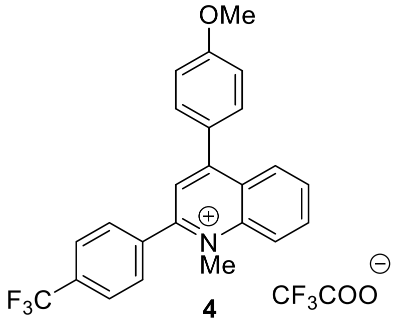

**4**

Salt **4** was synthesized using general procedure E using trimethyloxonium tetrafluoroborate (77 mg, 0.52 mmol, 3.0 eq.) and solution of 4-(4-methoxyphenyl)-2-(4-

(trifluoromethyl)phenyl)quinoline **S15** in anhydrous DCM (66 mg, 0.17 mmol, 1.0 eq., 1.7 mL $CH_2Cl_2$). After completion of reaction, the resulting mixture was then concentrated using a rotary evaporator and the residue was washed with cold diethyl ether. The residue was redissolved in acetonitrile and water (4:1) and purified by reverse-phased flash chromatography Method A followed by lyophilized gave **4** (49 mg, Yield: 71%) as a yellow solid.

**$^1$H NMR (500 MHz, $CD_3CN$):** δ 8.50 (dt, *J* = 9.0, 0.8 Hz, 1H), 8.28 (ddd, *J* = 8.8, 7.0, 1.5 Hz, 1H), 8.17 (ddd, *J* = 8.5, 1.5, 0.5 Hz, 1H), 8.03 – 7.93 (m, 4H), 7.85 (dt, *J* = 8.0, 0.8 Hz, 2H), 7.73 (d, *J* = 8.9 Hz, 2H), 7.24 (d, *J* = 8.9 Hz, 2H), 4.44 (s, 3H), 3.94 (s, 3H) ppm;

**$^{13}$C NMR (126 MHz, $CDCl_3$):** δ 163.8, 160.7, 160.5, 160.4, 156.8, 141.6, 139.9, 136.6, 133.0, 132.7, 132.4, 131.7, 131.0, 129.4, 127.8, 127.0, 127.0, 127.0, 126.9, 126.8, 126.2, 126.0, 124.1, 120.9, 115.9, 56.6, 43.8 ppm;

**$^{19}$F NMR (471 MHz, $CD_3CN$):** δ -76.60 (s, 3F), -63.31 (s, 3F) ppm;

**FT-IR ($v^{max}$ / $cm^{-1}$, neat):** 1675, 1601, 1567, 1510, 1462, 1417, 1389, 1324, 1247, 1175, 1128, 1066, 1037, 1018, 841, 767, 706

**HRMS-ESI (m/z):** calcd for $C_{24}H_{19}F_3NO^+$ $[M - CF_3COO]^+$ 394.1413, found 394.1413;

**M.P (°C):** 54-56

**2-(4-Methoxyphenyl)-1-methyl-4-(4-(trifluoromethyl)phenyl)quinolin-1-ium 2,2,2-trifluoroacetate (5):**

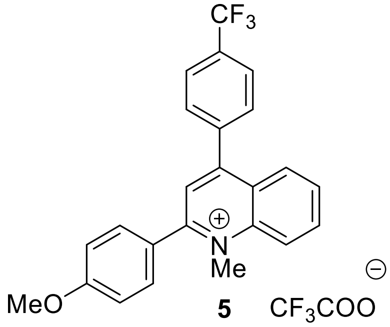


**5**

Salt **5** was synthesized using general procedure E using trimethyloxonium tetrafluoroborate (126 mg, 0.696 mmol, 3.0 eq.) and solution of 2-(4-methoxyphenyl)-4-(4-(trifluoromethyl)phenyl)quinoline **S14** in anhydrous DCM (88 mg, 0.23 mmol, 1.0 eq. 2.7 mL $CH_2Cl_2$). After completion of reaction, the resulting mixture was then concentrated using a rotary evaporator and the residue was washed with cold diethyl ether. The residue was redissolved in acetonitrile and water (4:1) and purified by reverse-phased flash chromatography Method A followed by lyophilized gave **5** (57 mg, Yield: 63%) as a yellow solid.

**$^1$H NMR (500 MHz, $CD_3CN$):** δ 8.50 (d, *J* = 9.0 Hz, 1H), 8.30 (t, *J* = 8.0 Hz, 1H), 8.19 (d, *J* = 8.4 Hz, 1H), 8.00 (d, *J* = 8.4 Hz, 4H), 7.86 (d, *J* = 8.0 Hz, 2H), 7.73 (d, *J* = 8.8 Hz, 2H), 7.26 (d, *J* = 8.8 Hz, 2H), 4.44 (s, 3H), 3.95 (s, 3H) ppm;

**$^{13}$C NMR (151 MHz, $CD_3CN$):** δ 162.7, 159.4, 155.7, 135.5, 131.9, 130.6, 129.9, 128.3, 126.7, 125.9, 125.9, 125.8, 124.9, 117.5, 117.3, 117.1, 55.6, 42.8, 29.0 ppm;

**$^{19}$F NMR (471 MHz, $CD_3CN$):** δ -75.86 (s, 3F), -63.31 (s, 3F) ppm;

**FT-IR ($v^{max}$ / $cm^{-1}$, neat):** 1683, 1599, 1565, 1510, 1463, 1433, 1417, 1389, 1362, 1322, 1247, 1167, 1121, 1065, 1036, 1017, 928, 838, 797, 765, 705

**HRMS-ESI (m/z):** calcd for $C_{24}H_{19}F_3NO^+$ [M - $CF_3COO]^+$ 394.1413, found 394.1413;

**M.P (°C):** 50-52

**1-Methyl-4-(2,3,6,7-tetrahydro-1*H*,5*H*-pyrido[3,2,1-ij]quinolin-9-yl)-2-(4-(trifluoromethyl)phenyl)quinolin-1-ium 2,2,2-trifluoroacetate (6):**

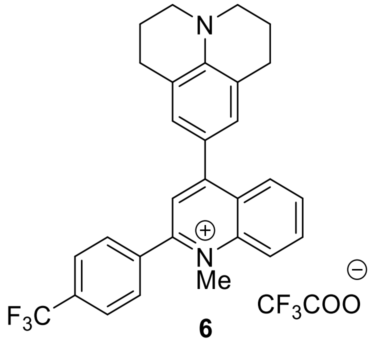


**6**

Salt **6** was synthesized using General procedure F using 9-(2-(4-(trifluoromethyl)phenyl)quinolin-4-yl)-2,3,6,7-tetrahydro-1*H*,5*H*-pyrido[3,2,1-ij]quinoline **S20** (85 mg, 0.19 mmol, 1.0 eq.), anhydrous chloroform (1.9 mL) and dimethyl sulfate (54 µL, 0.57 mmol, 3.0 eq.). The resulting mixture was then concentrated using a rotary evaporator and the residue was washed with cold diethyl ether. The residue was redissolved in acetonitrile and water (4:1) and purified by reverse-phased flash chromatography Method C followed by lyophilized gave **6** (46 mg, Yield: 52%) as a purple solid.

**$^1$H NMR (600 MHz, $CD_3CN$):** δ 8.32 (d, *J* = 8.9 Hz, 1H), 8.14 (ddd, *J* = 8.7, 7.0, 1.5 Hz, 1H), 8.00 (dd, *J* = 8.3, 1.4 Hz, 1H), 7.96 (d, *J* = 8.1 Hz, 2H), 7.90 (s, 1H), 7.86 – 7.79 (m, 3H), 7.25 (s, 2H), 4.41 (s, 3H), 3.45 – 3.08 (m, 4H), 2.80 (t, *J* = 6.3 Hz, 4H), 1.99 – 1.95 (m, 4H) ppm;

**$^{13}$C NMR (151 MHz, $CD_3CN$):** δ 160.9, 154.1, 147.8, 141.9, 140.4, 135.5, 132.3, 132.1, 131.5, 131.1, 129.7, 128.9, 126.9, 126.9, 126.9, 126.8, 126.7, 126.6, 126.1, 122.5, 120.5, 50.6, 44.5, 28.3, 21.8 ppm;

**$^{19}$F NMR (471 MHz, $CD_3CN$):** δ -63.23, -63.24, -76.24 ppm;

**FT-IR ($v^{max}$ / $cm^{-1}$, neat):** 2924, 2853, 1682, 1591, 1544, 1515, 1435, 1355, 1324, 1271, 1182, 1167, 1127, 1066, 846, 764

**HRMS-ESI (m/z):** calcd for $C_{29}H_{26}F_3N_2^+$ [M - $CF_3COO]^+$ 459.2043, found 459.2037;

**M.P (°C):** 56-58

**2,4-Bis(2,4-dimethoxyphenyl)-1-methylquinolin-1-ium 2,2,2-trifluoroacetate (7):**

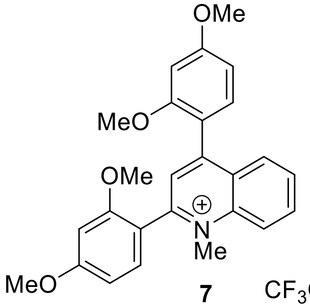

**7** $CF_3COO^{\ominus}$

Salt **7** was synthesized using General procedure E using trimethyloxonium tetrafluoroborate (81 mg, 0.55 mmol, 3.0 eq.) and solution of 2,4-bis(2,4-dimethoxyphenyl)quinoline **S2** in anhydrous DCM (73 mg, 0.18 mmol, 1.0 eq., 1.8 mL $CH_2Cl_2$). After completion of reaction, the resulting mixture was then concentrated using a rotary evaporator and the residue was washed with cold diethyl ether. The residue was redissolved in acetonitrile and water (4:1) and purified by reverse-phased flash chromatography Method A followed by lyophilized gave **7** (43 mg, Yield: 57%) as a yellow solid.

**$^1$H NMR (500 MHz, $CD_3CN$):** δ 8.41 (d, *J* = 9.0 Hz, 1H), 8.19 (ddd, *J* = 8.8, 7.0, 1.5 Hz, 1H), 8.04 (dd, *J* = 8.5, 1.5 Hz, 1H), 7.94 – 7.81 (m, 3H), 7.47 (d, *J* = 8.5 Hz, 1H), 7.35 (d, *J* = 8.4 Hz, 1H), 6.85 – 6.74 (m, 4H), 4.28 (s, 3H), 3.92 (d, *J* = 7.4 Hz, 6H), 3.88 (s, 3H), 3.74 (d, *J* = 2.4 Hz, 3H) ppm;

**$^{13}$C NMR (151 MHz, $CD_3CN$):** δ 165.6, 164.4, 160.6, 159.4, 159.1, 157.7, 156.7, 140.8, 136.0, 133.5, 130.3, 130.1, 127.9, 120.4, 107.5, 107.0, 100.0, 99.7, 56.9, 56.7, 56.5, 56.5, 42.5 ppm;

**FT-IR ($v^{max}$ / $cm^{-1}$, neat):** 1688, 1606, 1505, 1461, 1420, 1386, 1304, 1266, 1210, 1163, 1128, 1024, 834, 800, 767, 718

**$^{19}$F NMR (471 MHz, $CD_3CN$):** δ -76.44 (s, 3F) ppm;

**HRMS-ESI (m/z):** calcd for $C_{26}H_{26}NO_4^+$ $[M - CF_3COO]^+$ 416.1856, found 416.1856;

**M.P (°C):** 46-48

**2-(Benzofuran-2-yl)-4-(2,4-dimethoxyphenyl)-1-methylquinolin-1-ium 2,2,2-trifluoroacetate (8):**

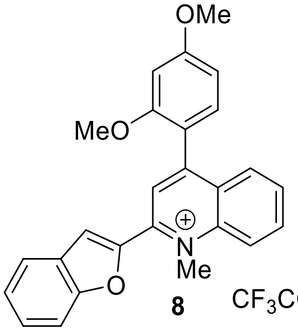

**8** $CF_3COO^{\ominus}$

Salt **8** was synthesized using General procedure F using 2-(benzofuran-2-yl)-4-(2,4-dimethoxyphenyl)quinoline **S17** (58 mg, 0.15 mmol, 1.0 eq.), anhydrous chloroform (1.5 mL) and dimethyl sulfate (43 μL, 0.46 mmol, 3.0 eq.). The resulting mixture was then concentrated using a rotary evaporator and the residue was washed with cold diethyl ether. The residue was redissolved in acetonitrile and water (4:1) and purified by reverse-phased flash chromatography Method C followed by lyophilized gave **8** (18 mg, Yield: 30%) as a yellow solid.

**$^1$H NMR (500 MHz, $CD_3CN$):** δ 8.46 (d, *J* = 9.0 Hz, 1H), 8.35 (s, 1H), 8.23 (ddd, *J* = 8.9, 7.0, 1.5 Hz, 1H), 8.05 (dd, *J* = 8.5, 1.4 Hz, 1H), 7.99 – 7.84 (m, 3H), 7.79 – 7.71 (m, 1H), 7.63 (ddd, *J* = 8.5, 7.2, 1.3 Hz, 1H), 7.48 (ddd, *J* = 8.0, 7.2, 0.9 Hz, 1H), 7.42 – 7.34 (m, 1H), 6.81 (d, *J* = 7.7 Hz, 2H), 4.68 (s, 3H), 3.93 (s, 3H), 3.76 (s, 3H) ppm;

**$^{13}$C NMR (151 MHz, $CD_3CN$):** δ 163.6, 156.4, 135.5, 132.4, 129.5, 129.3, 128.9, 127.9, 124.7, 124.6, 123.3, 119.4, 117.3, 112.1, 106.1, 99.0, 55.6, 55.5, 42.7 ppm;

**$^{19}$F NMR (471 MHz, $CD_3CN$):** δ -76.39 (s, 3F) ppm;

**FT-IR ($v^{max}$ / $cm^{-1}$, neat):** 1685, 1595, 1574, 1506, 1464, 1430, 1365, 1304, 1256, 1207, 1176, 1131, 1027, 838, 801, 762, 721

**HRMS-ESI (m/z):** calcd for $C_{26}H_{22}NO_3^+$ [M - $CF_3COO$]$^+$ 396.1594, found 396.1596;

**M.P (°C):** 52-54

**4-(2,4-Dimethoxyphenyl)-1-methyl-2-(1-methyl-1*H*-pyrazol-4-yl)quinolin-1-ium 2,2,2-trifluoroacetate (9):**

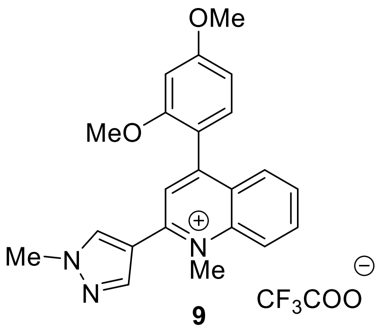


**9**

Salt **9** was synthesized using General procedure F using 4-(2,4-dimethoxyphenyl)-2-(1-methyl-1*H*-pyrazol-4-yl)quinoline **S19** (0.133 g, 0.385 mmol, 1.0 eq.), anhydrous chloroform (3.9 mL) and dimethyl sulfate (110 µL, 1.16 mmol, 3.0 eq.). The resulting mixture was then concentrated using a rotary evaporator and the residue was washed with cold diethyl ether. The residue was redissolved in acetonitrile and water (4:1) and purified by reverse-phased flash chromatography Method C followed by lyophilized gave **9** (28 mg, Yield: 20%) as a yellow solid.

**$^1$H NMR (500 MHz, $CD_3CN$):** δ 8.73 (d, *J* = 2.9 Hz, 2H), 8.08 (ddd, *J* = 8.5, 1.3, 0.7 Hz, 1H), 7.79 (ddd, *J* = 8.4, 6.8, 1.5 Hz, 1H), 7.73 – 7.59 (m, 2H), 7.54 (ddd, *J* = 8.3, 6.8, 1.3 Hz, 1H), 7.28 (d, *J* = 8.2 Hz, 1H), 6.88 – 6.60 (m, 2H), 4.12 (d, *J* = 0.9 Hz, 6H), 3.92 (s, 3H), 3.71 (s, 3H) ppm;

**$^{13}$C NMR (126 MHz, $CD_3CN$):** δ 163.0, 159.0, 149.0, 148.8, 148.1, 136.6, 132.7, 131.2, 130.2, 128.1, 127.9, 127.5, 124.9, 120.4, 118.3, 106.33, 99.8, 56.3, 38.3 ppm;

**$^{19}$F NMR (471 MHz, $CD_3CN$):** δ -76.44 (s, 3F) ppm;

**FT-IR ($v^{max}$ / $cm^{-1}$, neat):**

**HRMS-ESI (m/z):** calcd for $C_{22}H_{22}N_3O_2^+$ [M - $CF_3COO$]$^+$ 360.1707, found 360.1706;

**M.P (°C):** 124 – 126

**4-(2,4-Dimethoxyphenyl)-1-methyl-2-(thiophen-2-yl)quinolin-1-ium 2,2,2-trifluoroacetate (10):**

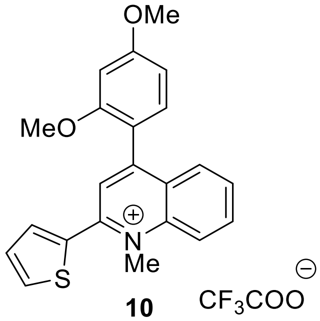

**10** $CF_3COO^{\ominus}$

Salt **10** was synthesized using general procedure F using 4-(2,4-dimethoxyphenyl)-2-(thiophen-2-yl)quinoline **S18** (76 mg, 0.22 mmol, 1.0 eq.), anhydrous chloroform (2.2 mL) and dimethyl sulfate (62 µL, 0.66 mmol, 3.0 eq.). The resulting mixture was then concentrated using a rotary evaporator and the residue was washed with cold diethyl ether. The residue was redissolved in acetonitrile and water (4:1) and purified by reverse-phased flash chromatography Method C followed by lyophilized gave **10** (17 mg, Yield: 21%) as a yellow solid.

**$^1$H NMR (500 MHz, $CD_3CN$):** δ 8.42 (d, *J* = 9.0 Hz, 1H), 8.23 (ddd, *J* = 8.9, 7.0, 1.5 Hz, 1H), 8.14 – 7.99 (m, 3H), 7.91 (ddd, *J* = 8.2, 7.0, 1.0 Hz, 1H), 7.78 (dd, *J* = 3.8, 1.2 Hz, 1H), 7.50 – 7.21 (m, 2H), 6.91 – 6.37 (m, 2H), 4.52 (s, 3H), 3.94 (s, 3H), 3.76 (s, 3H) ppm;

**$^{13}$C NMR (151 MHz, $CD_3CN$):** δ 166.1, 159.5, 158.8, 152.4, 141.3, 137.4, 133.4, 132.2, 132.0, 131.7, 131.5, 127.6, 127.5, 127.3, 121.1, 114.3, 107.8, 99.8, 99.7, 57.0, 56.8 ppm;

**$^{19}$F NMR (471 MHz, $CD_3CN$):** δ -76.63 (s, 3F) ppm;

**FT-IR ($v^{max}$ / $cm^{-1}$, neat):** 1737, 1686, 1607, 1592, 1507, 1461, 1439, 1419, 1351, 1305, 1286, 1266, 1210, 1163, 1136, 1022, 946, 858, 839, 799, 764, 717

**HRMS-ESI (m/z):** calcd for $C_{22}H_{20}NO_2S^+$ $[M - CF_3COO]^+$ 362.1209, found 362.0211;

**M.P (°C):** 48-50

**7-Methoxy-2,4-bis(4-methoxyphenyl)-1-methylquinolin-1-ium 2,2,2-trifluoroacetate (11):**

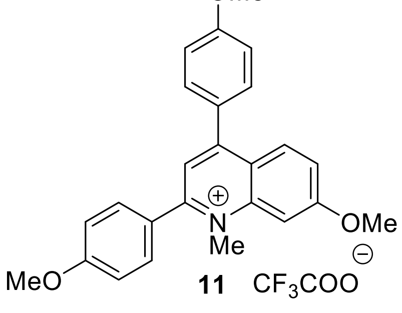

**11** $CF_3COO^{\ominus}$

Salt **11** was synthesized using general procedure E using trimethyloxonium tetrafluoroborate (122 mg, 3.0 eq., 824 µmol) and solution of 6-methoxy-2,4-bis(4-methoxyphenyl)quinoline **S4** in anhydrous DCM (102 g, 275 µmol, 1.0 eq., 2.8 mL $CH_2Cl_2$). After completion of reaction, the resulting mixture was then concentrated using a rotary evaporator and the residue was washed with cold diethyl ether. The residue was redissolved in acetonitrile and water (4:1) and purified by reverse-phased flash chromatography Method A followed by lyophilized gave **11** (67 mg, Yield: 63%) as a yellow solid.

**$^1$H NMR (500 MHz, $CD_3CN$):** δ 8.39 (d, *J* = 9.7 Hz, 1H), 7.88 (q, *J* = 3.3 Hz, 2H), 7.75 – 7.65 (m, 5H), 7.26 – 7.23 (m, 4H), 4.36 (s, 3H), 3.96 (d, *J* = 2.4 Hz, 9H) ppm;

**$^{13}$C NMR (126 MHz, $CDCl_3$):** δ 163.2, 162.7, 160.6, 157.3, 156.5, 137.0, 132.7, 132.5, 129.6, 128.4, 127.5, 126.3, 126.3, 122.4, 115.8, 115.6, 107.9, 63.0, 56.9, 56.5, 56.4, 43.4 ppm;

**FT-IR ($v^{max}$ / $cm^{-1}$, neat):** 1736, 1601, 1566, 1508, 1462, 1441, 1389, 1364, 1291, 1245, 1175, 1134, 1086, 1025, 837, 791, 763, 705

**$^{19}$F NMR (471 MHz, $CD_3CN$):** δ -75.63 (s, 3F) ppm;

**HRMS-ESI (m/z):** calcd for $C_{25}H_{24}NO_3^+$ [M - $CF_3COO$]$^+$ 386.1751, found 386.1750;

**M.P (°C):** 46-48

**7-(Dimethylamino)-2,4-bis(4-methoxyphenyl)-1-methylquinolin-1-ium 2,2,2-trifluoroacetate (12):**

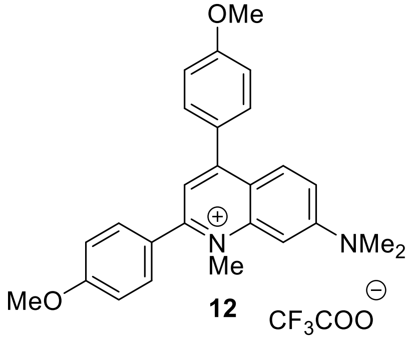


**12**

Salt **12** was synthesized using general procedure E using trimethyloxonium tetrafluoroborate (102 mg, 3.0 eq., 687 µmol) and solution of 2,4-bis(4-methoxyphenyl)-*N*,*N*-dimethylquinolin-7-amine **S7** in anhydrous DCM (88 mg, 0.23 mmol, 1.0 eq., 2.3 mL $CH_2Cl_2$). After completion of reaction, the resulting mixture was then concentrated using a rotary evaporator and the residue was washed with cold diethyl ether. The residue was redissolved in acetonitrile and water (4:1) and purified by reverse-phased flash chromatography Method A followed by lyophilized gave **12** (68 mg, Yield: 74%) as a yellow solid.

**$^1$H NMR (500 MHz, $CD_3CN$):** δ 8.65 (d, *J* = 2.9 Hz, 1H), 8.30 – 8.08 (m, 3H), 8.04 (s, 1H), 7.86 (dd, *J* = 9.4, 2.9 Hz, 1H), 7.59 – 7.34 (m, 2H), 7.27 – 6.97 (m, 4H), 3.89 (d, *J* = 13.7 Hz, 6H), 3.69 (s, 9H) ppm;

**$^{13}$C NMR (126 MHz, $CD_3CN$)** δ 162.8, 161.5, 160.7, 160.4, 159.1, 150.1, 148.8, 148.0, 132.1, 131.3, 130.3, 130.1, 129.9, 126.7, 121.6, 121.5, 118.2, 115.6, 115.3, 58.0, 56.2, 30.9 ppm;

**$^{19}$F NMR (471 MHz, $CD_3CN$):** δ -76.58 (s, 3F) ppm;

**FT-IR ($v^{max}$ / $cm^{-1}$, neat):** 1686, 1606, 1595, 1547, 1517, 1502, 1466, 1431, 1363, 1294, 1251, 1200, 1175, 1127, 1027, 835, 800, 718

**HRMS-ESI (m/z):** calcd for $C_{26}H_{27}N_2O_2^+$ [M - $CF_3COO$]$^+$ 399.2067, found 399.2067;

**M.P (°C):** 44-46

**7-(Dimethylamino)-2,4-bis(4-(dimethylamino)phenyl)-1-methylquinolin-1-ium 2,2,2-trifluoroacetate (13):**

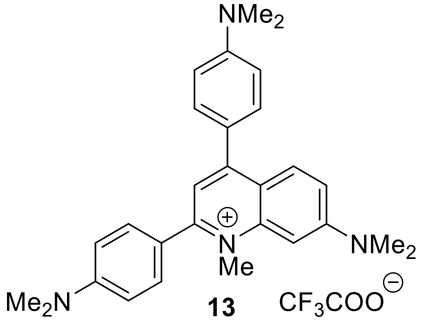

**13**

Salt **13** was synthesized using general procedure F using 2,4-bis(4-methoxyphenyl)-*N*,*N*-dimethylquinolin-7-amine **S8** (102 mg, 248 μmol, 1.0 eq.), anhydrous chloroform (2.5 mL) and dimethyl sulfate (71 μL, 3.0 eq., 0.75 mmol). The resulting mixture was then concentrated using a rotary evaporator and the residue was washed with cold diethyl ether. The residue was redissolved in acetonitrile and water (4:1) and purified by reverse-phased flash chromatography Method C followed by lyophilized gave **13** (59 mg, Yield: 56%) as a red solid.

**$^1$H NMR (500 MHz, $CD_3CN$):** δ 8.59 (s, 1H), 8.24 (d, J = 9.0 Hz, 1H), 8.01 (s, 1H), 7.78 (dd, J = 9.4, 2.9 Hz, 2H), 7.62 – 7.35 (m, 2H), 6.92 (dd, J = 34.0, 8.9 Hz, 0H), 3.67 (s, 1H), 3.06 (d, J = 1.7 Hz, 1H) ppm;

**$^{13}$C NMR (126 MHz, $CD_3CN$):** δ 158.9, 155.3, 153.3, 152.9, 148.9, 144.7, 139.1, 132.7, 132.4, 132.4, 132.3, 129.8, 122.3, 122.0, 120.8, 120.2, 119.3, 113.0, 112.6, 96.2, 95.3, 58.1, 43.1, 41.7, 40.9, 40.9, 40.4, 40.3, 40.3 ppm;

**$^{19}$F NMR (471 MHz, $CD_3CN$):** δ -76.36 (s, 3F) ppm;

**FT-IR ($v^{max}$ / $cm^{-1}$, neat):** 2980, 1686, 1628, 1604, 1578, 1511, 1382, 1328, 1254, 1200, 1173, 1127, 946, 824, 800, 718

**HRMS-ESI (m/z):** calcd for $C_{26}H_{33}N_4^+$ $[M - CF_3COO]^+$ 425.2700, found 425.2330;

**M.P (°C):** 42-44

**7-(Dimethylamino)-4-(4-(dimethylamino)phenyl)-1-methyl-2-(4-(trifluoromethyl)phenyl)quinolin-1-ium 2,2,2-trifluoroacetate (14):**

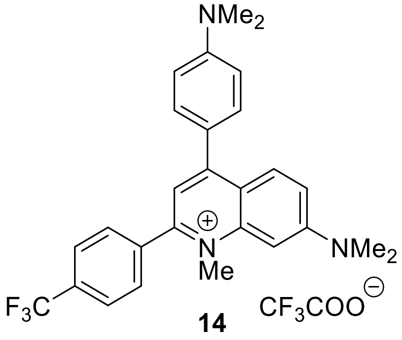

Salt **14** was synthesized using general procedure F using 4-(4-(dimethylamino)phenyl)-N,N-dimethyl-2-(4-(trifluoromethyl)phenyl)quinolin-7-amine **S16** (118 mg, 271 µmol, 1.0 eq.), anhydrous chloroform (2.7 mL) and dimethyl sulfate (77 µL, 0.81 mmol, 3.0 eq.). The resulting mixture was then concentrated using a rotary evaporator and the residue was washed with cold diethyl ether. The residue was redissolved in acetonitrile and water (4:1) and purified by reverse-phased flash chromatography Method C followed by lyophilized gave **14** (19 mg, Yield: 16%) as a red solid.

**$^1$H NMR (500 MHz, $CD_3CN$):** δ 8.38 (d, *J* = 8.1 Hz, 2H), 8.02 – 7.96 (m, 2H), 7.93 – 7.83 (m, 4H), 7.71 – 7.64 (m, 2H), 7.44 – 7.29 (m, 2H), 3.63 (d, *J* = 2.7 Hz, 9H), 3.16 (s, 6H) ppm;

**$^{13}$C NMR (151 MHz, $CD_3CN$):** δ 160.6, 160.4, 154.7, 153.5, 149.7, 149.3, 148.0, 142.0, 141.1, 132.4, 132.1, 131.9, 129.5, 129.5, 127.0, 126.8, 126.8, 126.8, 126.3, 124.5, 121.6, 121.5, 116.7, 116.4, 104.7, 58.2, 40.6 ppm;

**$^{19}$F NMR (471 MHz, $CD_3CN$):** δ -63.19 (s, 3F), -76.26 (s, 3F) ppm;

**FT-IR ($v^{max}$ / $cm^{-1}$, neat):** 2980, 1686, 1628, 1604, 1578, 1511, 1382, 1328, 1254, 1200, 1173, 1127, 946, 824, 800, 718

**HRMS-ESI (m/z):** calcd for $C_{27}H_{27}F_3N_3^+$ [M - $CF_3COO$]$^+$ 450.2152, found 450.2151;

**M.P (°C):** 78-80

**6-Methoxy-2,4-bis(4-methoxyphenyl)-1-methylquinolin-1-ium 2,2,2-trifluoroacetate (15):**

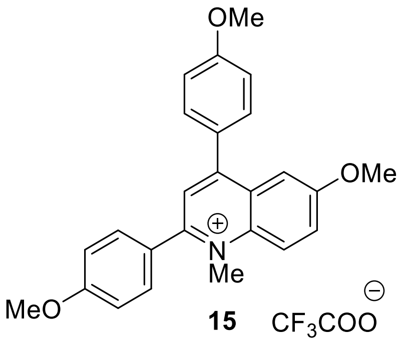

Salt **15** was synthesized using general procedure E using trimethyloxonium tetrafluoroborate (136 mg, 921 µmol, 3.0 eq.) and solution of 6-methoxy-2,4-bis(4-methoxyphenyl)quinoline **S5** in anhydrous DCM (114 mg, 307 µmol, 1.0 eq., 3.1 mL $CH_2Cl_2$). After completion of reaction, the resulting mixture was then concentrated using a rotary evaporator and the residue was washed with cold diethyl ether. The residue was redissolved in acetonitrile and water (4:1) and purified by reverse-phased flash chromatography Method A followed by lyophilized gave **15** (80 mg, Yield: 68%) as a yellow solid.

**$^1$H NMR (500 MHz, $CD_3CN$):** δ 8.41 (d, *J* = 9.7 Hz, 1H), 7.96 – 7.84 (m, 2H), 7.78 – 7.61 (m, 5H), 7.54 – 7.17 (m, 4H), 4.38 (s, 3H), 4.07 – 3.84 (m, 9H) ppm;

**$^{13}$C NMR (126 MHz, $CDCl_3$)** δ 162.9, 162.4, 160.5, 160.2, 159.9, 159.6, 156.9, 156.2, 136.7, 132.3, 132.2, 129.3, 128.0, 127.1, 125.9, 125.9, 122.0, 115.6, 115.5, 115.4, 115.3, 56.5, 56.1, 56.0, 43.0 ppm;

**$^{19}$F NMR (471 MHz, $CD_3CN$):** δ -76.60 (s, 3F) ppm;

**FT-IR ($v^{max}$ / $cm^{-1}$, neat):** 1681, 1602, 1510, 1462, 1421, 1390, 1365, 1291, 1246, 1202, 1177, 1128, 1026, 837, 800, 720

**HRMS-ESI (m/z):** calcd for $C_{25}H_{24}NO_3^+$ [M - $CF_3COO$]$^+$ 386.1751, found 386.1750;

**M.P (°C):** 46-48

**2,4-Bis(2,4-dimethoxyphenyl)-7-methoxy-1-methylquinolin-1-ium 2,2,2-trifluoroacetate (16):**

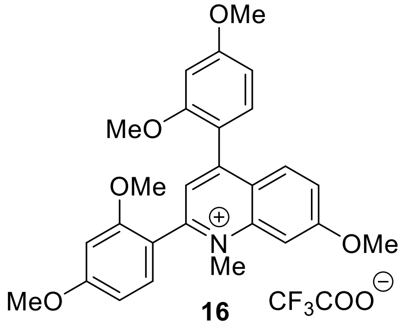

Salt **16** was synthesized using general procedure F using 2,4-bis(2,4-dimethoxyphenyl)-7-methoxyquinoline **S3** (202 mg, 468 µmol, 1.0 eq.), anhydrous chloroform (4.7 mL) and dimethyl sulfate (133 µL, 3.0 eq., 1.40 mmol). The resulting mixture was then concentrated using a rotary evaporator and the residue was washed with cold diethyl ether. The residue was redissolved in acetonitrile and water (4:1) and purified by reverse-phased flash chromatography Method C followed by lyophilized gave **16** (164 mg, Yield: 78%) as a yellow solid.

**$^1$H NMR (500 MHz, $CD_3CN$):** δ 7.92 (d, *J* = 9.3 Hz, 1H), 7.64 (s, 1H), 7.56 (d, *J* = 2.3 Hz, 1H), 7.48 – 7.34 (m, 2H), 7.32 (d, *J* = 8.4 Hz, 1H), 6.95 – 6.63 (m, 5H), 4.20 (s, 3H), 4.12 (s, 3H), 4.00 – 3.82 (m, 10H), 3.73 (s, 2H) ppm;

**$^{13}$C NMR (126 MHz, $CD_3CN$):** δ 165.6, 164.9, 163.8, 158.8, 158.5, 156.3, 155.4, 143.0, 132.9, 132.8, 132.6, 131.6, 124.9, 123.4, 121.5, 116.8, 115.2, 106.9, 106.4, 99.9, 99.5, 99.2, 57.2, 56.4, 56.2, 56.2, 56.1, 56.0, 56.0, 41.8 ppm;

**$^{19}$F NMR (471 MHz, $CD_3CN$):** δ -76.52 (s, 3F) ppm;

**FT-IR ($v^{max}$ / $cm^{-1}$, neat):** 1688, 1603, 1590, 1526, 1503, 1459, 1431, 1417, 1378, 1303, 1266, 1239, 1208, 1160, 1123, 1081, 1018, 936, 829, 797, 705

**HRMS-ESI (m/z):** calcd for $C_{27}H_{28}NO_5^+$ [M - $CF_3COO$]$^+$ 446.1962, found 446.1962;

**M.P (°C):** 54-56

**2,4-Bis(2,4-dimethoxyphenyl)-6,7-dimethoxy-1-methylquinolin-1-ium 2,2,2-trifluoroacetate (17):**

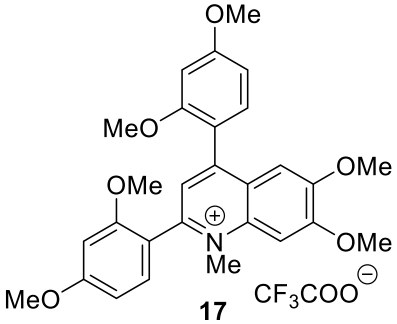


**17**

Salt **17** was synthesized using general procedure F using 2,4-bis(2,4-dimethoxyphenyl)-6,7-dimethoxyquinoline **S6** (98 mg, 0.21 mmol, 1.0 eq.), anhydrous chloroform (2.1 mL) and dimethyl sulfate (60 µL, 3.0 eq., 0.64 mmol). The resulting mixture was then concentrated using a rotary evaporator and the residue was washed with cold diethyl ether. The residue was redissolved in acetonitrile and water (4:1) and purified by reverse-phased flash chromatography Method C followed by lyophilized gave **17** (44 mg, Yield: 43%) as a yellow solid.

**$^1$H NMR (500 MHz, $CD_3CN$):** δ 7.61 (d, $J$ = 16.6 Hz, 1H), 7.53 (s, 1H), 7.43 (d, $J$ = 8.3 Hz, 1H), 7.35 (d, $J$ = 8.5 Hz, 1H), 7.18 (s, 1H), 6.89 – 6.70 (m, 4H), 4.18 (d, $J$ = 20.7 Hz, 6H), 3.91 (d, $J$ = 4.6 Hz, 6H), 3.86 (d, $J$ = 7.8 Hz, 6H), 3.77 (d, $J$ = 15.5 Hz, 3H) ppm;

**$^{13}$C NMR (151 MHz, $CD_3CN$)** δ 165.1, 164.1, 159.2, 158.8, 157.8, 153.8, 153.6, 152.1, 138.6, 133.3, 133.1, 133.1, 132.9, 125.8, 124.9, 124.8, 115.8, 107.4, 107.4, 107.2, 107.0, 100.1, 100.1, 100.0, 99.7, 57.9, 57.0, 56.8, 56.6, 56.5, 56.5, 42.3, 42.3 ppm;

**$^{19}$F NMR (471 MHz, $CD_3CN$):** δ -76.79 (s, 3F) ppm;

**FT-IR ($v^{max}$ / $cm^{-1}$, neat):** 1687, 1607, 1578, 1511, 1494, 1465, 1437, 1388, 1304, 1278, 1266, 1250, 1209, 1164, 1128, 1081, 1025, 831, 800, 719

**HRMS-ESI (m/z):** calcd for $C_{26}H_{30}NO_6^+$ [M - $CF_3COO$]$^+$ 476.2068, found 476.2066;

**M.P (°C):** 44-46

**Synthesis of Pyrylium and Pyridinium Salts:**

**Synthesis of Pyrylium Salts:**

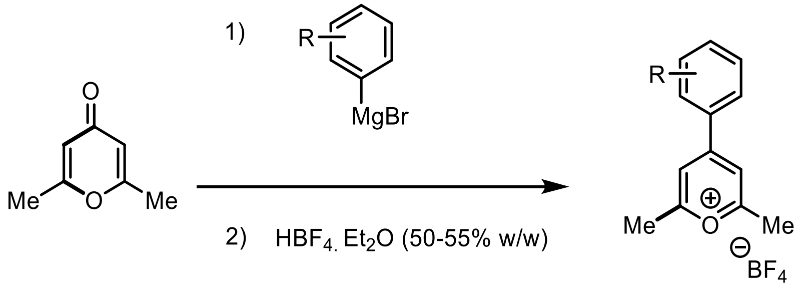


**General Procedure G:**

A round-bottom flask equipped with a side arm inlet adapter and stir bar was heat dried under vacuum, allowed to cool to room temperature, and refilled with $N_2$. The flask was charged with the 2,6-dimethyl-gamma-pyrone (1.0 eq.). The flask was cooled in an ice bath and stirred. To the stirring, Grignard reagent (2.0 eq.) was added dropwise. The resulting solution was stirred at 0°C for 30 minutes and then allowed to warm to room temperature while it was stirring. The solution was then stirred vigorously for 12 hours. The mixture was cooled down to 0°C, then added tetrafluoroboric acid (1.2 eq.) (50-55% w/w in $Et_2O$). The reaction was stirred for 1 h while warming to room temperature then precipitated with dropwise addition of cold diethyl ether. The precipitate was filtered and dissolved in ethanol then again precipitated out with cold diethyl ether. The resulting brown solid was finally washed with excess diethyl ether and dried under reduced pressure to yield the desired pyrylium salt (**S22-S24**).

**Summary of Pyrylium salts prepared here:**

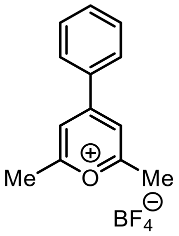


**S22, 79%**

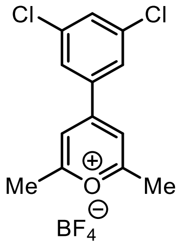


**S23, 73%**

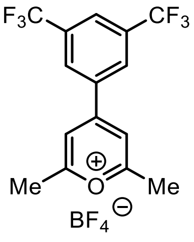


**S24, 74%**

**2,6-Dimethyl-4-phenylpyrylium tetrafluoroborate (S22):**

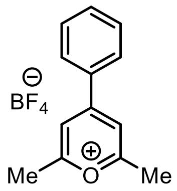

**S22**

Pyrylium salt **S22** was obtained by **General procedure G** using 2,6-dimethyl-gamma-pyrone (0.501 g, 4.03 mmol, 1.0 eq), and phenylmagnesium bromide solution (2.7 mL, 3.0 M in $Et_2O$, 8.06 mmol, 2.0 eq.). After the completion of reaction and quenching with tetrafluoroboric acid (50-55% w/w in $Et_2O$, 1.2 eq.) at 0°C, the reaction was additionally stirred for 1 h at room temperature. The resulting residue dissolved with ethanol using sonication, precipitated with cold diethyl ether, and filtered was washed with excess cold diethyl ether and dried under reduced pressure to yield **S22** (0.589 g, Yield: 79%) as a light brown solid.

**$^1$H NMR (500 MHz, $CD_3CN$)** δ 8.18 (s, 2H), 8.08 (dq, *J* = 8.5, 1.2 Hz, 2H), 7.84 – 7.78 (m, 1H), 7.74 – 7.67 (m, 2H), 2.88 (d, *J* = 1.2 Hz, 6H) ppm;

**$^{13}$C NMR (126 MHz, $CD_3CN$)** δ 179.5, 167.2, 136.4, 132.9, 131.2, 130.4, 119.5, 21.9 ppm;

This compound is known, and characterization data are consistent with the literature report.[5]

**4-(3,5-Dichlorophenyl)-2,6-dimethylpyrylium tetrafluoroborate (S23):**

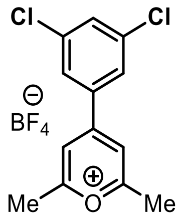

**S23**

Pyrylium salt **S23** was obtained *via* **General procedure G** using 2,6-dimethyl-gamma-pyrone (0.500 g, 4.03 mmol, 1.0 eq), and 3,5-dichlorophenylmagnesium bromide solution (2.1 mL, 0.5 M in THF, 8.06 mmol, 2.0 eq.). After quenching with tetrafluoroboric acid (50-55% w/w in $Et_2O$, 1.2 eq.) at 0°C and then stirred for 1 h at room temperature, the resulting residue dissolved with ethanol using sonication, precipitated with ether, and filtered was washed with excess diethyl ether and dried under reduced pressure to yield **S23** (0.743 g, Yield: 73%) as a dark brown solid.

**$^1$H NMR (500 MHz, $CD_3CN$):** δ 8.17 (s, 2H), 8.01 (d, *J* = 1.8 Hz, 2H), 7.87 (t, *J* = 1.8 Hz, 1H), 2.91 (s, 6H) ppm;

**$^{13}$C NMR (126 MHz, $CD_3CN$):** δ 180.6, 164.5, 137.4, 136.2, 135.0, 128.8, 22.1 ppm;

**$^{19}$F NMR (471 MHz, $CD_3CN$):** δ -149.34, -149.94, -149.99, -150.01, -150.52, -150.58, -151.9 ppm;

**FT-IR ($v^{max}$ / $cm^{-1}$, neat):** 1647, 1571, 1539, 1439, 1337, 1093, 1053, 1033, 957, 869, 808, 696

**HRMS (ESI) m/z:** [M - $BF_4$]$^+$ calcd for $C_{13}H_{11}Cl_2O^+$ 253.0181, found 253.0182;
**M.P (°C):** 130-132

**4-(3,5-Bis(trifluoromethyl)phenyl)-2,6-dimethylpyrylium tetrafluoroborate (S24):**

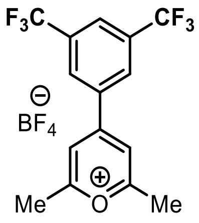


**S24**

Pyrylium salt **S24** was obtained *via* **General procedure G** using 2,6-dimethyl-gamma-pyrone (0.501 g, 4.03 mmol, 1.0 eq), and 3,5-bis(trifluoromethyl)phenyl-magnesium bromide (2.5 mL, 0.5 M in THF, 8.06 mmol, 2.0 eq.). After quenching with tetrafluoroboric acid (50-55% w/w in $Et_2O$, 1.2 eq.) at 0°C and then stirred for 1 h at room temperature, the resulting residue dissolved with ethanol using sonication, precipitated with ether, and filtered was washed with excess diethyl ether and dried under reduced pressure to yield **S24** (0.961 g, Yield: 74%) as a dark brown solid.

**$^1$H NMR (500 MHz, $CD_3CN$):** δ 8.62 (t, *J* = 2.0 Hz, 1H), 8.48 – 8.20 (m, 1H), 7.39 (s, 2H), 7.05 (s, 1H), 2.65 (s, 6H) ppm;

**$^{13}$C NMR (126 MHz, $CD_3CN$):** δ 181.0, 180.6, 180.5, 180.3, 177.9, 177.8, 176.5, 164.1, 135.6, 134.0, 133.8, 133.5, 133.2, 130.9, 128.7, 127.2, 125.1, 122.9, 121.0, 120.7, 112.2, 110.7, 21.1 ppm;

**$^{19}$F NMR (471 MHz, $CD_3CN$):** δ -63.45, -63.46, -63.47, -149.96, -149.98, -149.99, -151.63, -151.65, -151.67, -151.68 ppm;

**FT-IR ($v^{max}$ / cm$^{-1}$, neat):** 1709, 1641, 1538, 1489, 1423, 1362, 1322, 1282, 1222, 1180, 1139, 1092, 1057, 976, 958, 880, 700, 682

**HRMS (ESI) m/z:** [M - $BF_4$]$^+$ calcd for $C_{15}H_{11}F_6O^+$ 321.0709, found 321.0709;

**M.P (°C):** 120-122

#### 4.2 Synthesis of aryl substituted pyridinium salt:

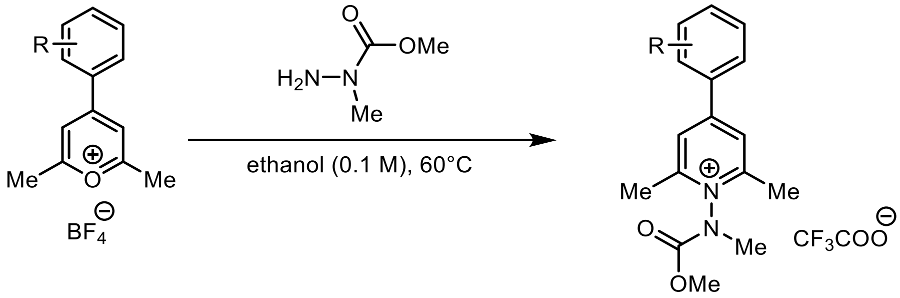


##### General Procedure H:

The synthesis of aryl substituted pyridinium salts was achieved by adapting the procedure of Taylor,[4] as follows:

A round-bottom flask equipped with a stir bar and a side arm inlet adapter was heat dried under vacuum, allowed to cool to room temperature, and refilled with $N_2$. The flask was charged with aryl pyrylium salt (**S22-S24**, 1.0 eq). To the flask was added ethanol *via* syringe, and the resulting mixture was stirred vigorously. To the stirring solution was added 1-methylhydrazine-1-carboxylate[1] (3.0 eq.), and the resulting solution was stirred vigorously at 60°C for 3 hours. The resultant mixture was then concentrated using a rotary evaporator before being redissolved in acetonitrile and then precipitated *via* dropwise addition of diethyl ether. The resulting solid was filtered, and the residue was washed with additional ether and purified using reverse-phased flash chromatography to yield the desired pyridinium salt (**18-20**).

##### Prepared pyridinium salt:

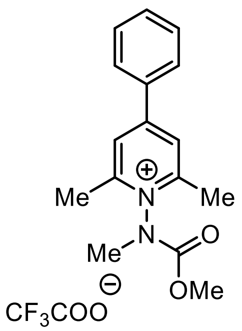

**18, 53%**

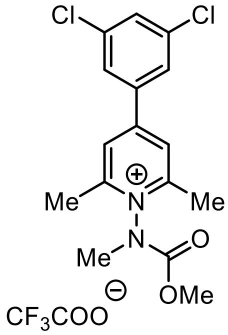

**19, 51%**

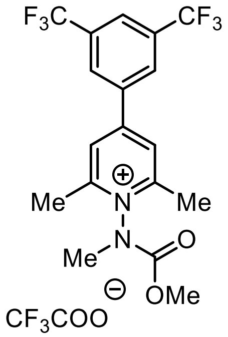

**20, 62%**

**1-((Methoxycarbonyl)(methyl)amino)-2,6-dimethyl-4-phenylpyridin-1-ium 2,2,2-trifluoroacetate (18):**

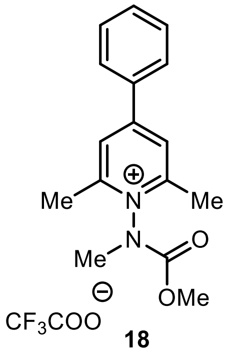

**18**

Phenyl pyridin-1-ium salt **18** was obtained *via* **General procedure H** using phenyl pyrilium[5] **S22** (0.462 g, 2.49 mmol, 1.0 eq), 1-methylhydrazine-1-carboxylate[1] (0.779 g, 7.48 mmol, 3.0 eq) and ethanol (25 mL). After the completion of reaction, the resulting solution was concentrated using a rotary evaporator. The dark residue was then dissolved with acetonitrile and water (4:1). The solution was purified by reverse-phased flash chromatography Method B followed by lyophilized to yield the desired pyridinium salt **16** (0.361 g, Yield: 53%) as white solid. The compound is a 1.2:1 mixture of rotamers at 298K.

**$^1$H NMR (500 MHz, $CD_3CN$):** δ 8.15 (s, 2H), 7.96 – 7.89 (m, 2H), 7.67 (dddt, *J* = 14.6, 8.6, 5.8, 2.2 Hz, 3H), 3.92 (s, 2H), 3.74 (s, 1H), 3.51 (s, 2H), 3.48 (s, 1H), 2.72 (d, *J* = 1.5 Hz, 6H) ppm;

**$^{13}$C NMR (126 MHz, $CD_3CN$):** δ 159.3, 159.1, 158.6, 158.5, 154.5, 153.1, 134.6, 134.5, 133.8, 133.8, 130.9, 130.9, 130.7, 129.3, 129.3, 126.1, 126.0, 55.8, 55.8, 38.3, 37.4, 19.5, 19.5 ppm;

**$^{19}$F NMR (471 MHz, $CD_3CN$):** δ -76.37 (s, 3F) ppm;

**FT-IR ($v^{max}$ / $cm^{-1}$, neat):** 1735, 1676, 1627, 1596, 1564, 1447, 1353, 1335, 1197, 1170, 1120, 1034, 1003, 983, 924, 880, 827, 799, 770, 717, 693

**HRMS (ESI) m/z:** $[M – CF_3COO^-]^+$ calcd for $C_{16}H_{19}N_2O_2^+$ 271.1441, found 271.1439;

**M.P (°C):** 48-50

**4-(3,5-Dichlorophenyl)-1-((methoxycarbonyl)(methyl)amino)-2,6-dimethylpyridin-1-ium 2,2,2-trifluoroacetate (19):**

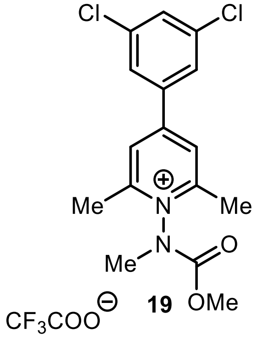


3,5-Dichloro phenyl pyridin-1-ium salt **19** was obtained *via* **General procedure H** using pyrylium **S23** (0.312 g, 1.23 mmol, 1.0 eq), 1-methylhydrazine-1-carboxylate[1] (0.383 g, 3.68 mmol, 3.0 eq) and ethanol (12 mL). After the completion of reaction, the resulting solution was concentrated using a rotary evaporator. The dark residue was then dissolved with acetonitrile and water (4:1). The solution was purified by reverse-phased flash chromatography Method B followed by lyophilized to yield the desired pyridinium salt **19** (0.212 g, Yield: 51%) as off white solid. The compound is a 1.2:1 mixture of rotamers at 298K.

**$^1$H NMR (500 MHz, $CD_3CN$):** δ 8.48 (d, *J* = 1.6 Hz, 2H), 8.36 – 8.14 (m, 3H), 3.93 (s, 2H), 3.75 (s, 1H), 3.52 (d, *J* = 19.0 Hz, 3H), 2.77 (d, *J* = 1.3 Hz, 6H) ppm;

**$^{13}$C NMR (126 MHz, $CD_3CN$):** δ 160.6, 160.2, 160.0, 155.3, 155.2, 154.4, 152.9, 137.1, 137.0, 133.8, 133.8, 133.6, 133.5, 133.3, 133.2, 133.0, 130.1, 127.4, 127.2, 127.1, 126.8, 126.7, 125.7, 125.2, 123.0, 120.9, 55.9, 55.8, 38.2, 37.3, 19.6, 19.5 ppm;

**$^{19}$F NMR (471 MHz, $CD_3CN$):** δ -76.50 (s, 3F) ppm;

**FT-IR ($v^{max}$ / $cm^{-1}$, neat):** 1734, 1629, 1560, 1454, 1380, 1350, 1331, 1271, 1134, 984, 935, 862, 803, 761, 705

**HRMS (ESI) m/z:** [M – $CF_3COO^-$]$^+$ calcd for $C_{16}H_{17}Cl_2N_2O_2^+$ 339.0662, found 339.0664

**M.P (°C):** 82-84

**4-(3,5-Bis(trifluoromethyl)phenyl)-1-((methoxycarbonyl)(methyl)amino)-2,6-dimethylpyridin-1-ium 2,2,2-trifluoroacetate (20):**

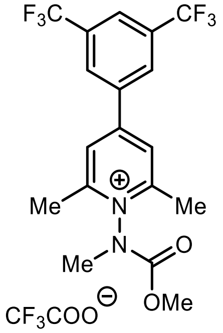

**20**

3,5-Bis(trifluoromethyl) phenyl pyridin-1-ium **20** was obtained *via* **General procedure H** using pyrylium **S24** (1.001 g, 3.121 mmol, 1.0 eq), 1-methylhydrazine-1-carboxylate[1] (0.973 g, 9.35 mmol, 3.0 eq) and ethanol (31 mL). After the completion of reaction, the resulting solution was concentrated using a rotary evaporator. The dark residue was then dissolved with acetonitrile and water (4:1). The solution was purified by reverse-phased flash chromatography Method B followed by lyophilized to yield the desired pyridinium salt **20** (0.788 g, Yield: 62%) as brown solid. The compound is a 1.2:1 mixture of rotamers at 298K.

**$^1$H NMR (500 MHz, $CD_3CN$):** δ 8.47 (s, 2H), 8.29 (d, *J* = 5.0 Hz, 3H), 3.93 (s, 2H), 3.75 (s, 1H), 3.51 (d, *J* = 16.8 Hz, 3H), 2.76 (d, *J* = 1.6 Hz, 6H) ppm;

**$^{13}$C NMR (126 MHz, $CD_3CN$):** δ 159.9, 159.7, 155.5, 154.4, 153.0, 137.8, 137.7, 137.1, 137.0, 132.8, 132.7, 128.0, 126.7, 55.9, 55.8, 38.2, 37.3, 19.6, 19.5 ppm;

**$^{19}$F NMR (471 MHz, $CD_3CN$):** δ -63.46, -63.47, -76.31, -76.79, -77.44 ppm;

**FT-IR ($v^{max}$ / $cm^{-1}$, neat):** 1738, 1633, 1572, 1460, 1381, 1351, 1318, 1280, 1172, 1132, 938, 907, 845, 797, 762, 706, 683

**HRMS (ESI) m/z:** $[M – CF_3COO^-]^+$ calcd for $C_{18}H_{17}F_6N_2O_2^+$ 407.1189, found 407.1188;

**M.P (°C):** 98-100

**Synthesis of arylated pyridinium salt with azide reporter tag:**

**1-((((6-Azidohexyl)oxy)carbonyl)(methyl)amino)-2,6-dimethyl-4-phenylpyridin-1-ium 2,2,2-trifluoroacetate (19a):**

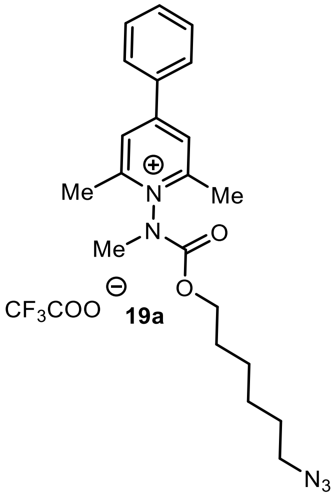


A round bottom flask equipped with a stir bar and a side arm inlet adapter was charged with 2,6-dimethyl-4-phenylpyrylium tetrafluoroborate **S22** (109 mg, 0.588 mmol, 1.0 eq.), 6-azidohexyl 1-methylhydrazine-1-carboxylate[2] **S25** (380 mg, 1.77 mmol, 3.0 eq.) and ethanol (6 mL) for 3 h. The reaction was stirred vigorously for 3 h at 60°C. The resulting mixture was concentrated using a rotary evaporator and purified by reverse-phased flash chromatography Method B followed by lyophilized gave **19a** (88 mg, 39%) as a yellow oil. The compound is a 1.2:1 mixture of rotamers at 293K.

**$^1$H NMR (500 MHz, $CD_3CN$):** δ 8.16 (d, *J* = 9.4 Hz, 2H), 8.05 – 7.88 (m, 2H), 7.66 (dtd, *J* = 14.2, 7.2, 5.1 Hz, 3H), 4.30 (t, *J* = 6.5 Hz, 1H), 4.16 (t, *J* = 6.4 Hz, 1H), 3.50 (d, *J* = 19.8 Hz, 3H), 3.37 – 3.24 (m, 2H), 3.15 (t, *J* = 6.9 Hz, 1H), 2.72 (d, *J* = 5.9 Hz, 6H), 1.76 (p, *J* = 6.6 Hz, 1H), 1.62 (t, *J* = 7.0 Hz, 1H), 1.57 – 1.28 (m, 4H), 1.25 – 1.16 (m, 1H), 1.12 (q, *J* = 7.4 Hz, 1H) ppm;

**$^{13}$C NMR (151 MHz, $CD_3CN$):** δ 161.1, 160.8, 160.5, 160.2, 159.2, 159.1, 158.6, 158.5, 154.1, 152.6, 134.6, 134.4, 133.9, 133.7, 130.9, 130.9, 130.8, 129.4, 129.3, 129.2, 126.1, 125.9, 120.6, 116.0, 113.7, 71.2, 69.4, 69.2, 52.1, 52.1, 51.8, 38.2, 37.2, 30.4, 29.5, 29.4, 29.2, 29.2, 28.9, 27.3, 27.0, 26.7, 26.5, 26.0, 25.9, 19.6, 19.5 ppm;

**$^{19}$F NMR (471 MHz, $CD_3CN$):** δ -76.55 (s, 3F) ppm;

**FT-IR ($v^{max}$ / $cm^{-1}$, neat):** 2097, 1734, 1630, 1566, 1446, 1394, 1333, 1268, 1140, 1035, 1003, 795, 770, 706

**HRMS (ESI) m/z:** $[M – CF_3COO^-]^+$ calcd for $C_{21}H_{28}F_6N_5O_2^+$ 382.2238, found 382.2240,

**1-((((6-Azidohexyl)oxy)carbonyl)(methyl)amino)-4-(3,5-bis(trifluoromethyl)phenyl)-2,6-dimethylpyridin-1-ium 2,2,2-trifluoroacetate (20a):**

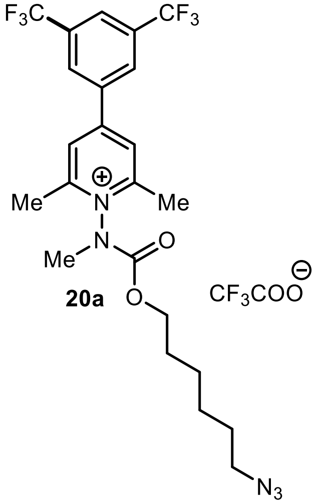


A round bottom flask equipped with a stir bar and a side arm inlet adapter was charged with 4-(3,5-bis(trifluoromethyl)phenyl)-2,6-dimethylpyrylium tetrafluoroborate **S24** (201 mg, 0.626 mmol, 1.0 eq.), 6-azidohexyl 1-methylhydrazine-1-carboxylate[2] **S25** (404 mg, 1.88 mmol, 3.0 eq.) and ethanol (6 mL) for 3 h. The reaction was stirred vigorously for 3 h at 60°C. The resulting mixture was concentrated using a rotary evaporator and the solution was purified by reverse-phased flash chromatography Method B followed by lyophilized gave **20a** (97 mg, Yield: 30%) as a brown oil. The compound is a 1.1:1 mixture of rotamers at 294K.

**$^{1}$H NMR (500 MHz, $CD_3CN$):** δ 8.49 (d, *J* = 9.1 Hz, 2H), 8.31 (d, *J* = 8.3 Hz, 3H), 4.31 (t, *J* = 6.5 Hz, 1H), 4.17 (t, *J* = 6.4 Hz, 1H), 4.07 (s, 5H), 3.51 (d, *J* = 18.7 Hz, 3H), 3.41 – 3.25 (m, 6H), 3.19 – 3.09 (m, 4H), 2.77 (s, 3H), 1.76 (q, *J* = 6.8 Hz, 1H), 1.67 – 1.30 (m, 18H), 1.16 (dq, *J* = 42.3, 8.0 Hz, 2H) ppm;

**$^{13}$C NMR (151 MHz, $CD_3CN$):** δ 161.0, 160.2, 160.0, 156.3, 155.3, 155.2, 154.0, 152.4, 137.1, 136.9, 133.7, 133.6, 133.5, 133.4, 133.3, 133.2, 133.1, 130.2, 130.1, 127.2, 127.1, 126.9, 126.8, 125.1, 123.3, 121.5, 117.7, 117.1, 115.8, 71.3, 71.2, 69.6, 69.3, 67.8, 67.3, 62.6, 52.1, 52.0, 51.8, 38.2, 37.1, 33.6, 30.9, 30.6, 30.4, 29.5, 29.4, 29.4, 29.3, 29.2, 28.9, 27.8, 27.7, 27.3, 27.1, 27.0, 26.9, 26.7, 26.6, 26.5, 26.1, 26.0, 25.9, 19.7, 19.6, 18.6 ppm;

**$^{19}$F NMR (471 MHz, $CD_3CN$):** δ -63.42, -63.43, -63.44, -75.93, -76.16 ppm;

**FT-IR ($v^{max}$ / $cm^{-1}$, neat):** 2096, 1715, 1633, 1461, 1396, 1338, 1317, 1281, 1136, 906, 845, 797, 762, 705, 682

**HRMS (ESI) m/z:** $[M – CF_3COO^-]^+$ calcd for $C_{23}H_{26}F_6N_5O_2^+$ 518.1985, found 518.1986.

**Synthesis of arylated pyridinium salt with Biotin reporter tag:**

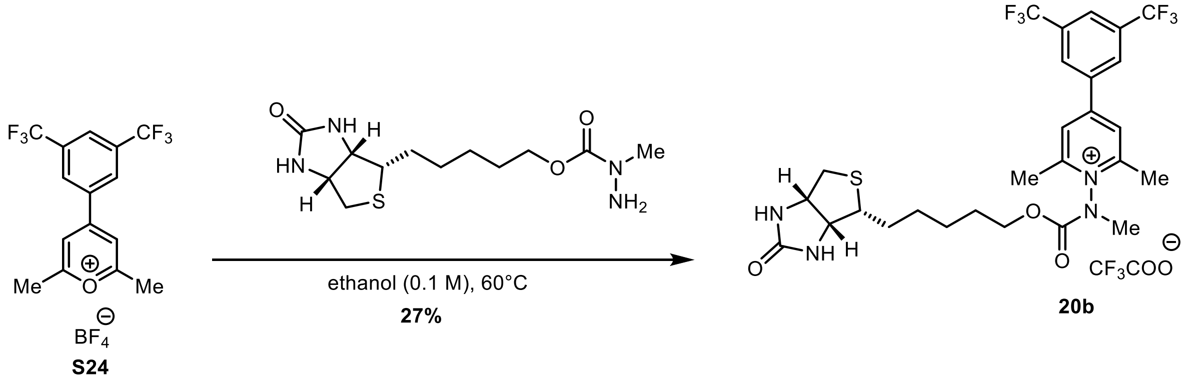


**Procedure:**

A round-bottom flask equipped with a stir bar and a side arm inlet adapter was heat dried under vacuum, allowed to cool to room temperature, and refilled with $N_2$. The flask was charged with the pyrylium salt **S24** (91 mg, 0.28 mmol, 1.0 eq.), added 5-((3a*S*,4S,6a*R*)-2-oxohexahydro-1*H*-thieno[3,4-d]imidazole-4-yl)pentyl 1methylhydrazine-1-carboxylate[2] (0.257 g, 3.0 eq., 0.850 mmol) and ethanol (3 mL) then stirred vigorously for 3 hours at 60°C. The resulting mixture was then concentrated using a rotary evaporator before being redissolved in acetonitrile and water (4:1). The residue was purified by HPLC Method A followed by lyophilized to give **20b** (46 mg, Yield: 27%) as a brown oil. The compound is a 1.2:1 mixture of rotamers at 298K.

**$^1$H NMR (600 MHz, $CD_3CN$)** δ 8.49 (d, *J* = 14.0 Hz, 2H), 8.36 – 8.27 (m, 3H), 5.58 (d, *J* = 309.3 Hz, 2H), 4.45 (s, 1H), 4.31 (t, *J* = 6.4 Hz, 2H), 4.17 (tt, J = 6.2, 3.2 Hz, 1H), 3.54 (s, 1.87 H), 3.49 (s, 1.56 H), 3.20 (s, 1H), 3.04 (s, 1H), 2.90 (dd, *J* = 12.7, 5.0 Hz, 1H), 2.85 – 2.74 (m, 7H), 2.71 – 2.63 (m, 1H), 2.57 (d, *J* = 12.7 Hz, 0.62 H), 1.74 (dp, *J* = 34.8, 6.8 Hz, 2H), 1.65 – 1.37 (m, 3H), 1.32 – 1.08 (m, 2H) ppm;

**$^{13}$C NMR (151 MHz, $CD_3CN$)** δ 164.1, 161.0, 160.7, 160.5, 160.2, 160.0, 160.0, 155.3, 155.2, 154.0, 152.5, 137.1, 137.0, 133.8, 133.7, 133.6, 133.6, 133.5, 133.4, 133.3, 133.1, 133.1, 130.2, 130.2, 130.2, 130.2, 130.2, 127.3, 127.2, 127.2, 127.1, 127.1, 126.9, 126.9, 126.9, 126.9, 126.8, 126.8, 126.8, 126.7, 126.7, 125.1, 123.3, 121.5, 116.9, 69.6, 69.4, 62.6, 61.0, 56.5, 56.3, 41.1, 40.9, 38.2, 37.2, 30.9, 29.5, 29.4, 29.2, 28.8, 26.4, 26.3, 20.2, 19.7, 19.6 ppm;

**$^{19}$F NMR (471 MHz, $CD_3CN$)**: δ -63.39, -63.44, -76.28, -76.53 ppm;

**FT-IR ($v^{max}$ / $cm^{-1}$, neat):** 1738, 1684, 1634, 1463, 1395, 1318, 1282, 1179, 1139, 938, 846, 801, 706, 683

**HRMS (ESI) m/z:** $[M – CF_3COO^-]^+$ calcd for $C_{27}H_{31}F_6N_4O_3S^+$ 605.2016, found 605.2016,

### 4. Protein Labelling

#### 4.1 Observation of photosensitization with 1 and commercial catalyst screening

**Attempted labelling of lysozyme with 2 using 427 nm light:**

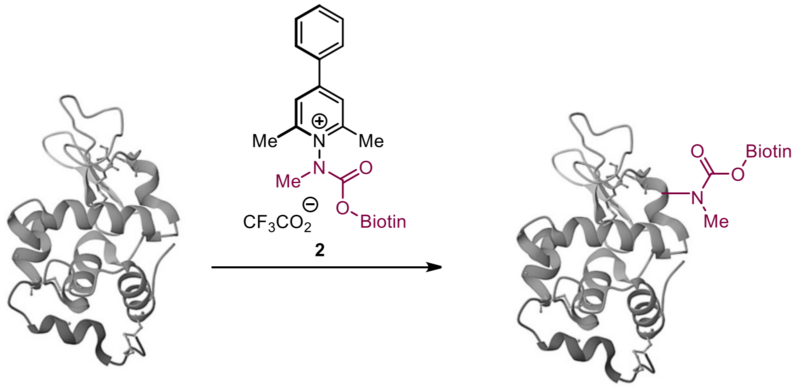


To a 2 mL Pyrex LC/MS vial (Thermo model # 03-391-39) containing Lysozyme (34 μL, 146 μM, 10 μM final concentration) were added solutions of $Na_2HPO_4$ (50 μL, 200 mM, pH 7.4, 20 mM final concentration), GSH (15 μL, 10 mM, 300 μM final concentration) and pyridinium salt **2** (7 μL, 100 μM, 7 mM final concentration) one after another. The solution vial was then placed into a HepatoChem photo-redox box (model # HCK1006-01-016) and then degassed using $N_2$ for 60 minutes. A 427 nm Kessil PR160L LED (100% intensity) was then used to irradiate the actively degassing solution for thirty minutes. The reaction mixture was analyzed directly by LC–MS Method A. The labeling outcomes were assessed from deconvoluted mass spectra. According to LC–MS Method A, the reaction was estimated to have proceeded with 0% conversion.

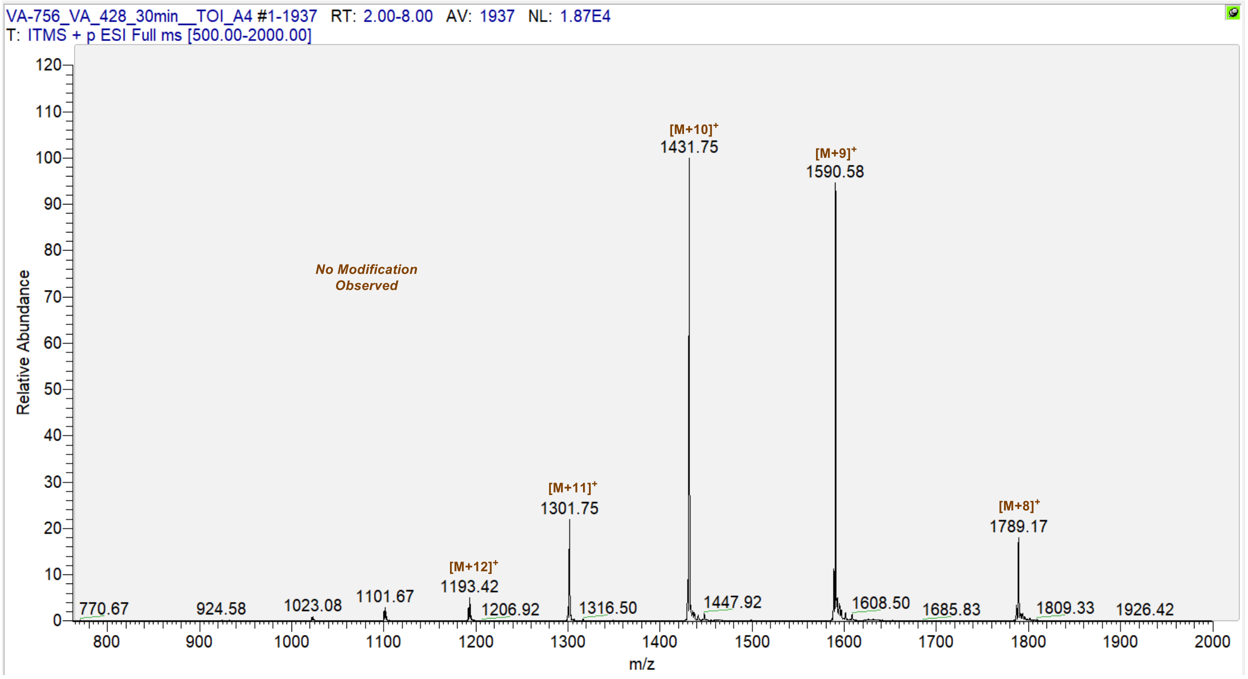


**Figure S1**: Mass spectrum of crude reaction mixture of lysozyme labelling with **2**.

**Photosensitization of 2 with 1 result in lysozyme labelling with both probes:**

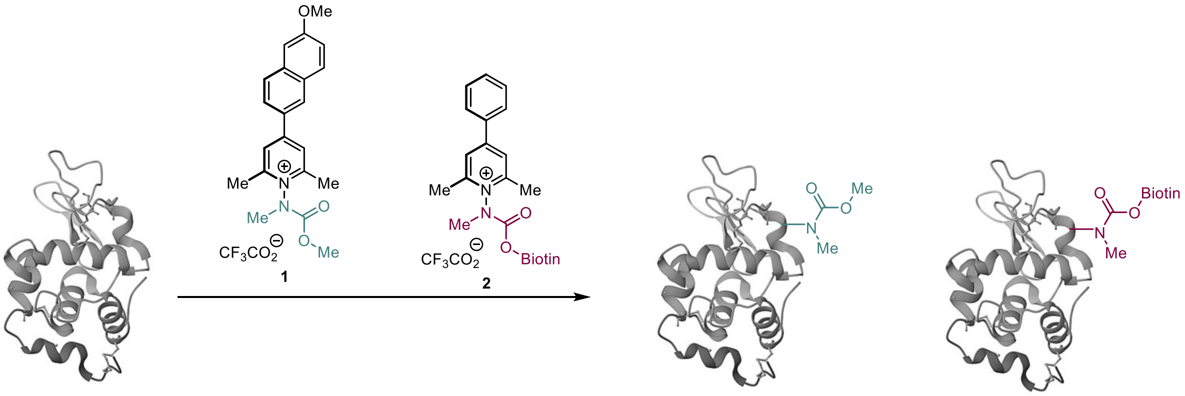


To a 2 mL Pyrex LC/MS vial (Thermo model # 03-391-39) containing a Lysozyme (34 µL, 146 µM, 10 µM final concentration) were added solutions of $Na_2HPO_4$ (50 µL, 200 mM, pH 7.4, 20 mM final concentration), GSH (15 µL, 10 mM, 300 µM final concentration), pyridinium salt **1** (5 µL, 100 µM, 10 mM final concentration) and pyridinium salt **2** (7 µL, 100 µM, 7 mM final concentration) one after another. The solution vial was then placed into a HepatoChem photo-redox box (model # HCK1006-01-016) and then degassed using $N_2$ for 60 minutes. A 427 nm Kessil PR160L LED (100% intensity) was then used to irradiate the actively degassing solution for thirty minutes. The reaction mixture was analyzed directly *via* LC/MS LC–MS Method A. The reaction mixture was directly analyzed with LC-MS Method A and was estimated to have proceeded with 64% conversion.

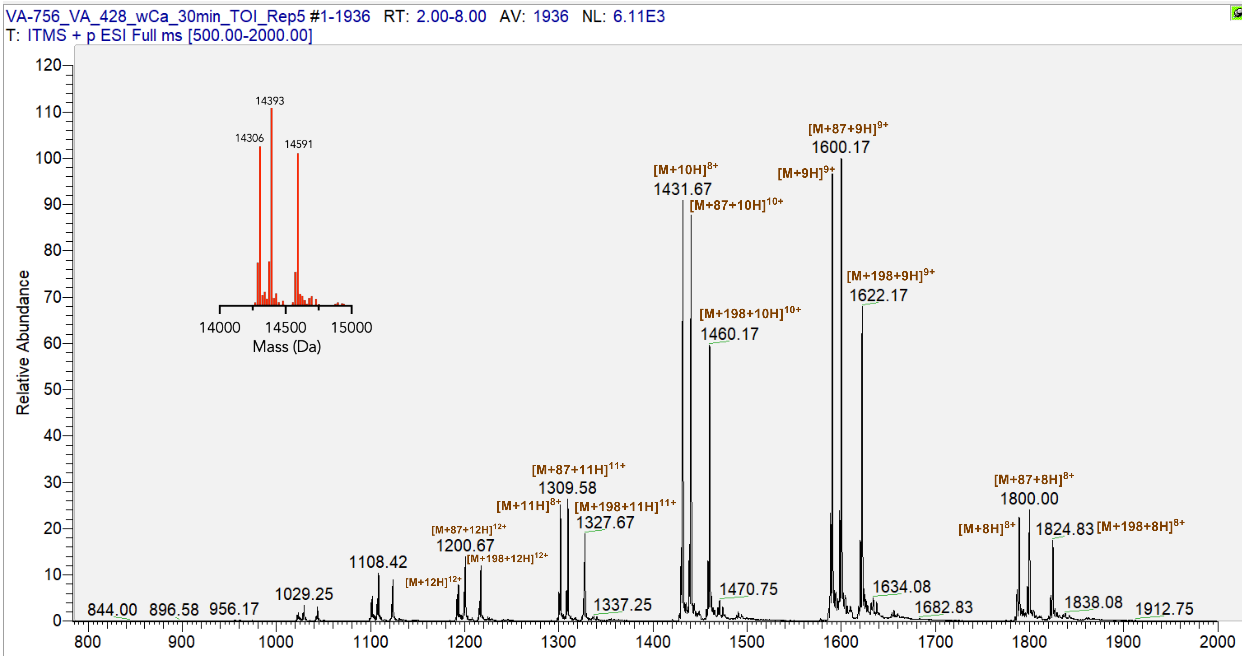


**Figure S2.** Mass spectrum of crude reaction mixtures of sensitized labelling with **1** and **2**.

#### Catalyst screening:

#### Commercial and known photocatalysts:

##### General Procedure:

To a 2 mL Pyrex LC/MS vial (Thermo model # 03-391-39) containing a 10 μM of lysozyme (from chicken egg) were added solutions of $Na_2HPO_4$ buffer (pH 7.4, 20 mM), L-Glutathione reduced (300 μM), pyridinium probe **18** (100 μM) and photocatalyst (Table S4, Entry 1-6, 10 μM) one after another. The resulting solution was diluted to a final volume of 500 μL by adding deionized water and sparged using Nitrogen for a minimum of 30 minutes. The solution vial was then placed into a HepatoChem photo-redox box (model # HCK1006-01-016). A 427 nm Kessil PR160L LED (100% intensity) was then used to irradiate the actively degassing solution for five minutes. The resultant reaction mixture was then analyzed directly via LC/MS. Labelling outcomes were analyzed using LC/MS (LC-MS Method A).

##### Commercial photocatalysts assayed:

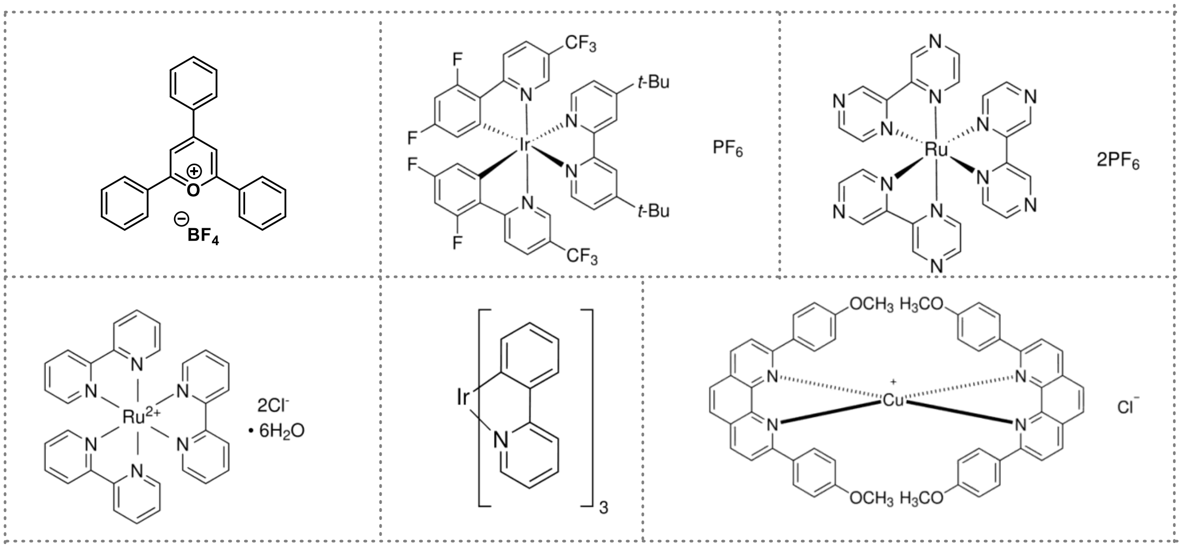


##### General Scheme:

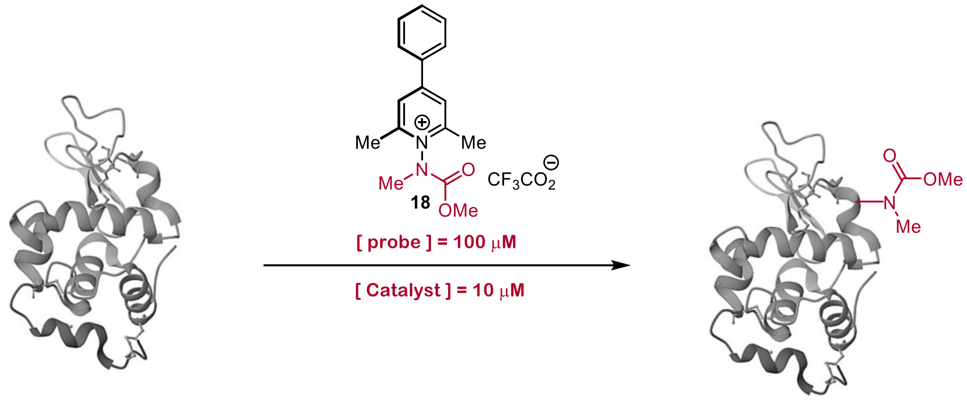

| Entry | Lysozyme [μM] | Probe 18 [μM] | Photocatalyst [10 μM] | pH 7.4 $PO_4$ [mM] | GSH [μM] | Time (min) | % mod* |
|---|---|---|---|---|---|---|---|
| **1** | 10 | 100 | Triphenyl Pyrylium $BF_4$[a] | 20 | 300 | 5 | - |
| **2** | 10 | 100 | $[Ir(dF(CF_3)ppy)_2(dtbpy)]$ $PF_6$[b] | 20 | 300 | 5 | 0 (4)[g] |
| **3** | 10 | 100 | $Ru(BPY)_3$ hexahydrate[c] | 20 | 300 | 5 | - |
| **4** | 10 | 100 | $[Ru(bpz)_3][PF_6]_2$[d] | 20 | 300 | 5 | - |
| **5** | 10 | 100 | $Ir(ppy)_3$[e] | 20 | 300 | 5 | - |
| **6** | 10 | 100 | $Cu(dap)Cl_2$[f] | 20 | 300 | 5 | - |

[a]Stock Solution in DMSO, [b]Stock Solution in MeCN:$H_2O$ (1:1), [c]Stock Solution in $H_2O$, [d]Stock Solution in $H_2O$, [e]Stock Solution in DMSO, [f]Stock Solution in MeCN:$H_2O$ (1:1), [g]Stock Solution in MeCN.

*Data are average values from independent photosensitizer screening experiments each performed with two technical replicates.

**Table S1:** Result for Screening commercial photocatalyst used for photosensitization study.

##### 4.2 Designer quinolinium screening/optimization:

To a 2 mL Pyrex LC/MS vial (Thermo model # 03-391-39) containing a 10 µM of lysozyme were added solutions of $Na_2HPO_4$ buffer (pH 7.4, 20 mM), L-Glutathione (300 µM), the desired pyridinium probes (**18-20**, 100 µM) and the desired photosensitizer (**3-17**, 5 – 200 µM). The resulting solution was diluted to a final volume of 500 µL with deionized water and the solutions sparged with $N_2$ for 60 minutes. The solution was then irradiated using a HepatoChem photo-redox box (model # HCK1006-01-016) equipped with a 427 nm Kessil PR160L LED (100% intensity) for five minutes with active degassing (See Figure S3 B). The resultant reaction mixture was then analyzed directly via LC/MS. Labelling outcomes were analyzed using LC/MS method A.

**General Scheme:**

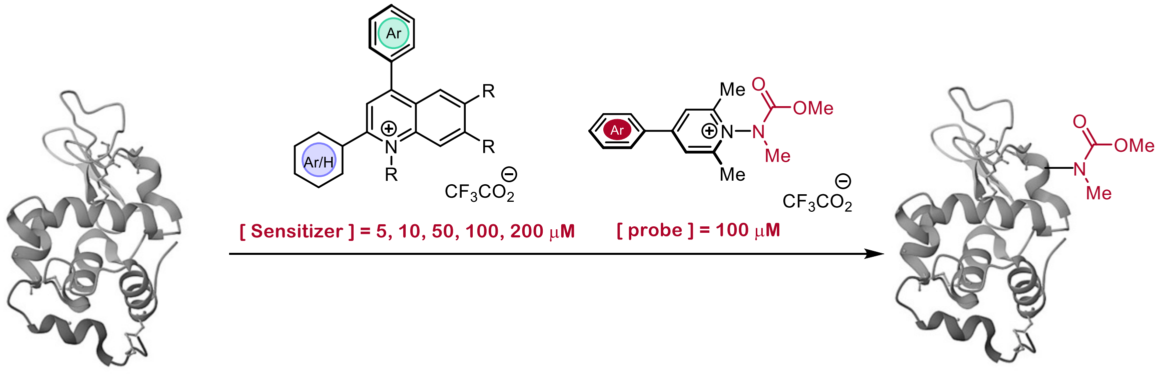


**Pyridinium salts screening used for photosensitization study:**

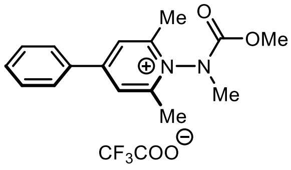

**18**

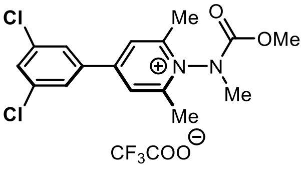

**19**

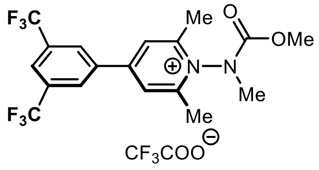

**20**

**Condition Screening for photosensitization:**

| *Condition A* | *Condition B* | *Condition C* | *Condition D* | *Condition E* |
|---|---|---|---|---|
| *10 µM Lysozyme* | *10 µM Lysozyme* | *10 µM Lysozyme* | *10 µM Lysozyme* | *10 µM Lysozyme* |
| *20 mM (pH 7.4)* | *20 mM (pH 7.4)* | *20 mM (pH 7.4)* | *20 mM (pH 7.4)* | *20 mM (pH 7.4)* |
| *5 µM Sensitizer* | *10 µM Sensitizer* | *50 µM Sensitizer* | *100 µM Sensitizer* | *200 µM Sensitizer* |
| *100 µM Probe* | *100 µM Probe* | *100 µM Probe* | *100 µM Probe* | *100 µM Probe* |
| *5 min Irradiation* | *5 min Irradiation* | *5 min Irradiation* | *5 min Irradiation* | *5 min Irradiation* |

**Synthesized diarylquinolinium aromatic cation sensitizer used for optimal photosensitizer screening:**

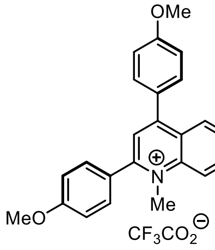

**3**

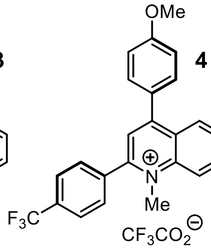

**4**

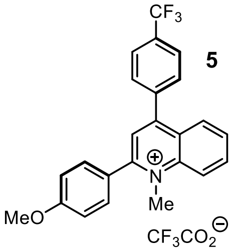

**5**

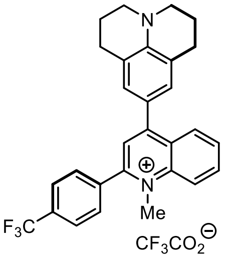

**6**

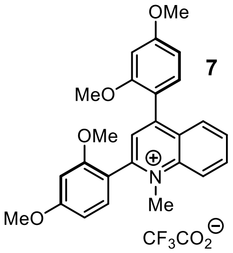

**7**

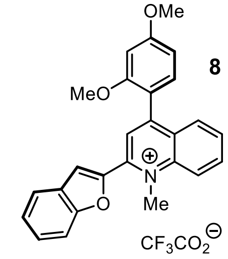

**8**

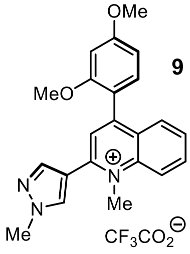

**9**

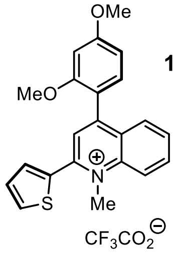

**10**

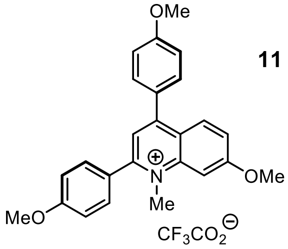

**11**

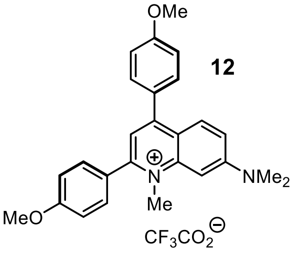

**12**

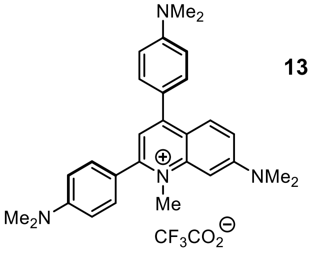

**13**

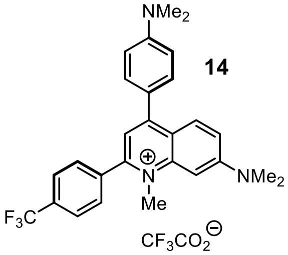

**14**

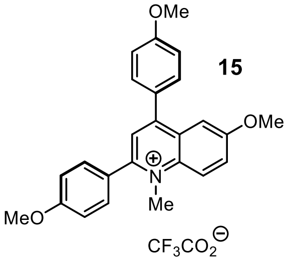

**15**

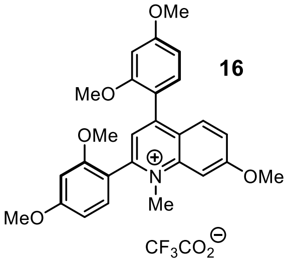

**16**

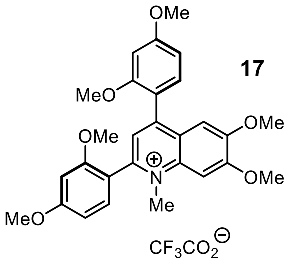

**17**

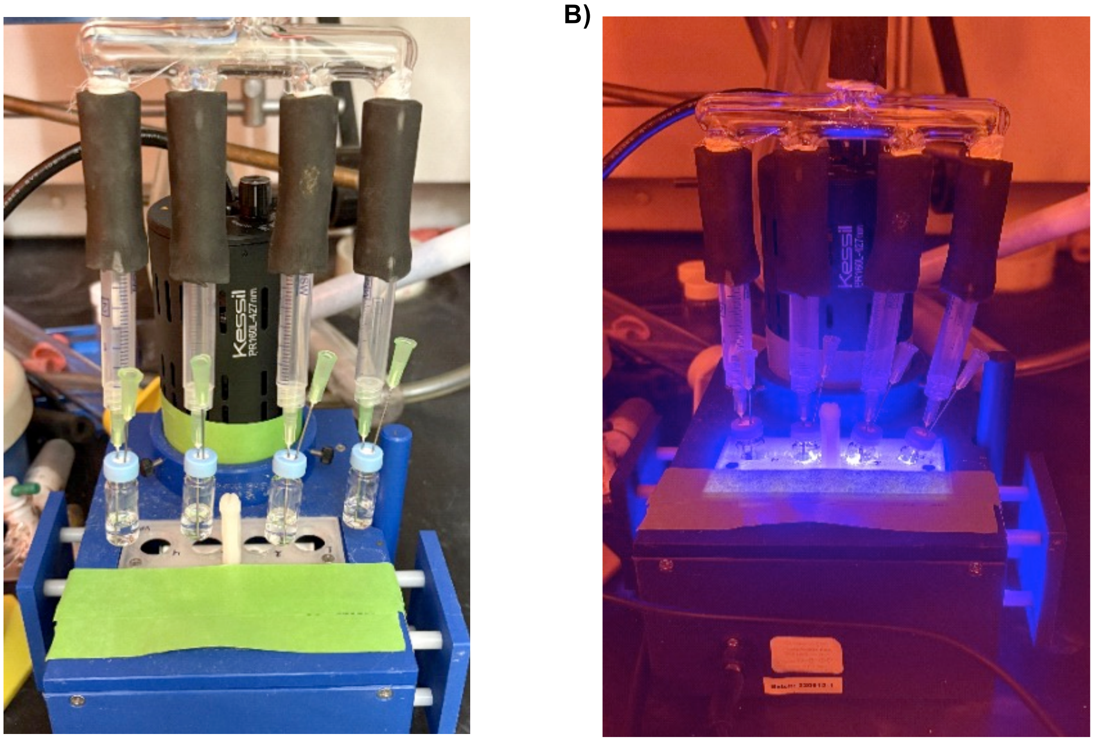


**Figure S3**: (A) Post-irradiation setup showing four reaction vials arranged in a holder for the photochemical labeling of lysozyme, each with a total reaction volume of 500 µL. (B) Sample during illumination with a 427 nm Kessil lamp, positioned in the photoredox box to ensure uniform light exposure.

**Summary of sensitizer screening:**

| Pyridinium Salt | [Sensitizer] (µM) | 3 | 4 | 5 | 6 | 7 | 8 | 9 | 10 | 11 | 12 | 13 | 14 | 15 | 16 | 17 |
|---|---|---|---|---|---|---|---|---|---|---|---|---|---|---|---|---|
| **18** | 200 | 0.25 | 0.30 | 0.14 | 0.00 | 0.27 | 0.00 | 0.00 | 0.00 | 0.43 | 0.00 | 0.00 | 0.00 | 0.30 | 0.28 | 0.29 |
| **18** | 100 | 0.22 | 0.34 | 0.07 | 0.00 | 0.27 | 0.00 | 0.00 | 0.00 | 0.24 | 0.00 | 0.00 | 0.00 | 0.00 | 0.23 | 0.26 |
| **18** | 50 | 0.26 | 0.07 | 0.00 | 0.00 | 0.25 | 0.00 | 0.00 | 0.00 | 0.19 | 0.00 | 0.00 | 0.00 | 0.00 | 0.14 | 0.19 |
| **18** | 10 | 0.00 | 0.00 | 0.00 | 0.00 | 0.04 | 0.00 | 0.00 | 0.00 | 0.00 | 0.00 | 0.00 | 0.00 | 0.00 | 0.02 | 0.00 |
| **18** | 5 | 0.00 | 0.00 | 0.00 | 0.00 | 0.00 | 0.00 | 0.00 | 0.00 | 0.00 | 0.00 | 0.00 | 0.00 | 0.00 | 0.00 | 0.00 |
| **19** | 200 | 0.34 | 0.18 | 0.32 | 0.00 | 0.32 | 0.04 | 0.00 | 0.00 | 0.34 | 0.00 | 0.00 | 0.00 | 0.16 | 0.28 | 0.29 |
| **19** | 100 | 0.23 | 0.17 | 0.21 | 0.00 | 0.18 | 0.03 | 0.00 | 0.00 | 0.20 | 0.00 | 0.00 | 0.00 | 0.00 | 0.24 | 0.22 |
| **19** | 50 | 0.16 | 0.16 | 0.16 | 0.00 | 0.26 | 0.03 | 0.00 | 0.00 | 0.12 | 0.00 | 0.00 | 0.00 | 0.00 | 0.19 | 0.16 |
| **19** | 10 | 0.11 | 0.00 | 0.00 | 0.00 | 0.05 | 0.00 | 0.00 | 0.00 | 0.00 | 0.00 | 0.00 | 0.00 | 0.00 | 0.14 | 0.00 |
| **19** | 5 | 0.00 | 0.00 | 0.00 | 0.00 | 0.02 | 0.00 | 0.00 | 0.00 | 0.00 | 0.00 | 0.00 | 0.00 | 0.00 | 0.00 | 0.00 |
| **20** | 200 | 0.35 | 0.29 | 0.21 | 0.00 | 0.32 | 0.00 | 0.21 | 0.00 | 0.31 | 0.00 | 0.00 | 0.00 | 0.37 | 0.35 | 0.39 |
| **20** | 100 | 0.25 | 0.23 | 0.26 | 0.00 | 0.26 | 0.05 | 0.08 | 0.00 | 0.34 | 0.00 | 0.00 | 0.00 | 0.32 | 0.25 | 0.36 |
| **20** | 50 | 0.27 | 0.12 | 0.25 | 0.00 | 0.25 | 0.04 | 0.04 | 0.00 | 0.29 | 0.00 | 0.00 | 0.00 | 0.23 | 0.26 | 0.25 |
| **20** | 10 | 0.04 | 0.00 | 0.00 | 0.00 | 0.14 | 0.00 | 0.00 | 0.00 | 0.06 | 0.00 | 0.00 | 0.00 | 0.07 | 0.19 | 0.00 |
| **20** | 5 | 0.00 | 0.00 | 0.00 | 0.00 | 0.04 | 0.00 | 0.00 | 0.00 | 0.02 | 0.00 | 0.00 | 0.00 | 0.00 | 0.12 | 0.00 |

Labelled/ Unlabelled Ratio: 0, 0.1, 0.2, 0.3, 0.4

**Figure S4:** Ratios of labelled:unlabelled lysozyme of all screened catalysts (**3-17**, heatmap X-axis) with different pyridinium salts (**18-20**) and differing catalyst concentration ranges (Y-axis). Data are average of two replicates.

#### 4.3 Photosensitization of Pyridium salts with reporter tag with optimal sensitizer

Photosensitized labelling of lysozyme with **16** and **20a**:

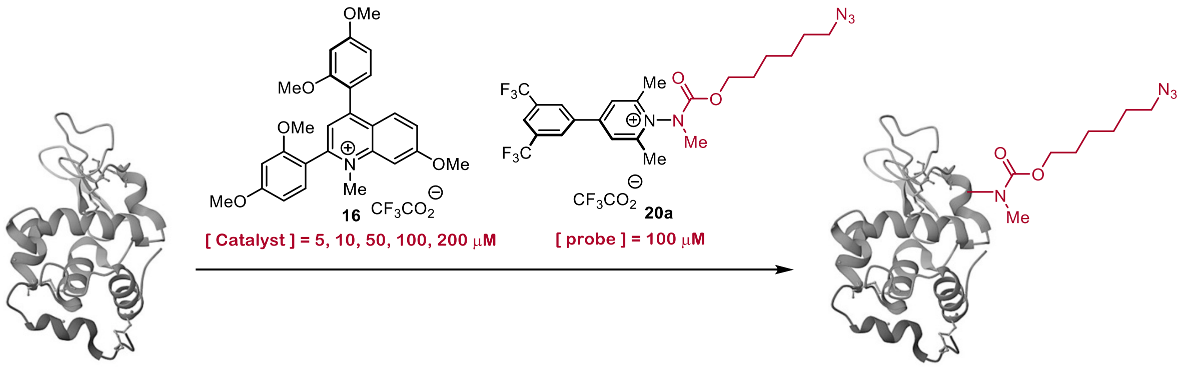


Photosensitized azidocarbamlyation of lysozyme with **16** was performed by following general procedure I. The reaction mixture was analyzed directly *via* LC/MS method A.

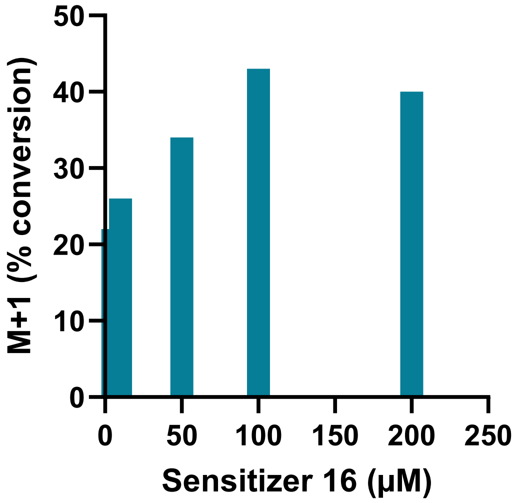


**Figure S5.** %Conversion of Lysozyme (from chicken egg white) labelling varying concentrations of **16** and 100 μM **20a**. Data are average of two replicates.

Photosensitized labelling of lysozyme with **16** and **20b**:

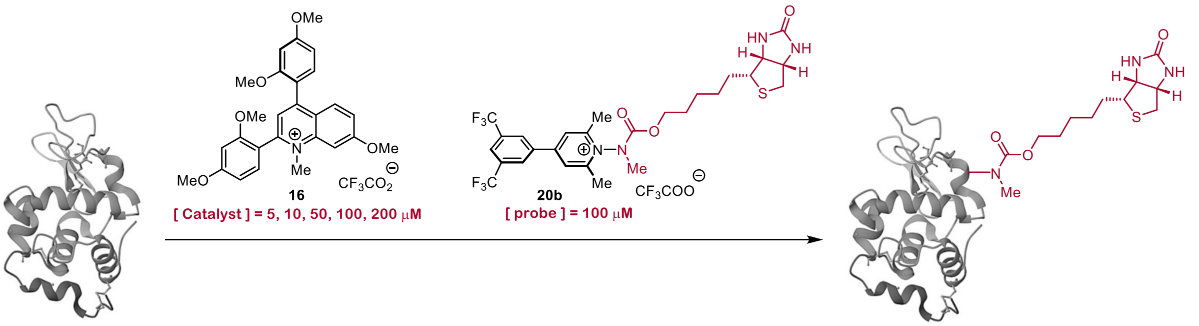


Photosensitized biotinylation of lysozyme with **16** was performed by following general procedure I. The reaction mixture was analyzed directly *via* LC/MS method A.

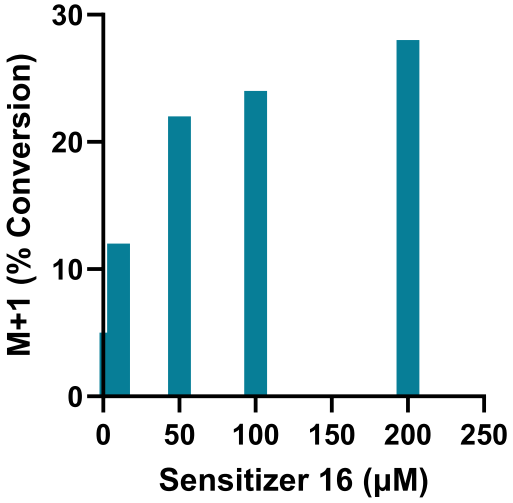


**Figure S6.** %Conversion of Lysozyme (from chicken egg white) labelling varying concentrations of **16** and 100 μM **20b**. Data are average of two replicates.

##### 4.4 Catalytic sensitization of Leuprorelin•OAc

To a 2 mL Pyrex LC/MS vial (Thermo model # 03-391-39) containing Leuprolide•OAc (as indicated in Table S1 and S2) was added solutions of buffer, L-Glutathione, **20** and **16** sequentially. The resulting solution was diluted to a final volume of 500 µL with DI-water and degassed with $N_2$ for 30 minutes prior to irradiation. The solution was then irradiated using a HepatoChem photo-redox box (model # HCK1006-01-016) equipped with a 427 nm Kessil PR160L LED (100% intensity) for five minutes with active degassing (See Figure S3 B). The resultant reaction mixture was then analyzed directly via LC/MS. Labelling outcomes were analyzed using LC/MS. The reaction mixture was analyzed directly *via* LC/MS method B.

**General Scheme:**

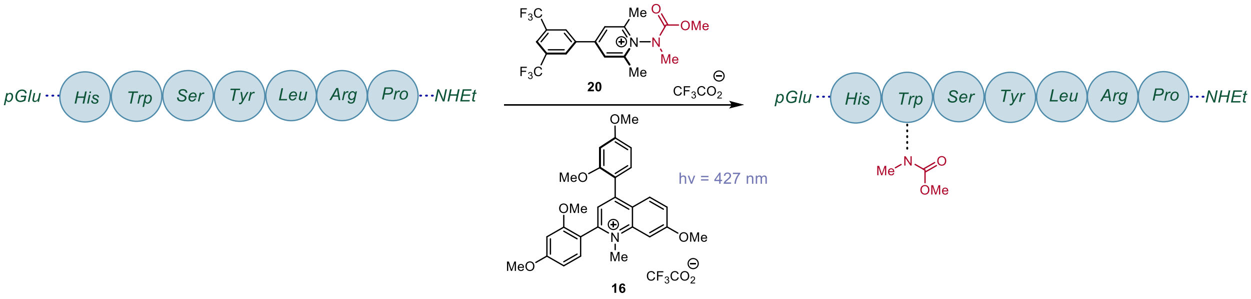


| S. No. | Peptide [µM] | Probe 20 [µM] | Sensitizer 16 [µM] | $Na_2HPO_4$ 7.4 [mM] | GSH µM | Time (min) | % conversion* |
|---|---|---|---|---|---|---|---|
| 1 | 100 | 100 | 10 | 20 | 300 | 5 | 18 |
| 2 | 100 | 100 | 20 | 20 | 300 | 5 | 17 |
| 3 | 100 | 100 | 30 | 20 | 300 | 5 | 20 |
| 4 | 100 | 100 | 40 | 20 | 300 | 5 | 18 |
| 5 | 100 | 200 | 10 | 20 | 300 | 5 | 23 |
| 6 | 100 | 300 | 10 | 20 | 300 | 5 | 19 |
| 7 | 100 | 500 | 10 | 20 | 300 | 5 | 30 |
| 8 | 100 | 1000 | 10 | 20 | 300 | 5 | 23 |
| 9 | 200 | 100 | 10 | 20 | 300 | 5 | 0 |
| 10 | 500 | 100 | 10 | 20 | 300 | 5 | 0 |
| 11 | 1000 | 100 | 10 | 20 | 300 | 5 | 0 |
| 12 | 100 | 400 | 10 | 20 | 300 | 5 | 33 |
| 13 | 100 | 400 | 10 | 20 | 300 | 10 | 37 |
| 14 | 100 | 400 | 10 | 20 | 300 | 20 | 43 |
| 15 | 100 | 400 | 10 | 20 | 300 | 40 | 46 |

* Average of two replicates.

**Table S2:** Attempted modification of Leuprolide Acetate Peptide for Optimal Photosensitizer **16** Turnover in presence of Phosphate (pH 7.4) Buffer. Conversions are averages of two replicates.

| S. No. | [Peptide] μM | [Probe 20] μM | [Sensitizer 16] μM | [$NH_4OAc$ 6.9] mM | GSH μM | Time (min) | % conversion* |
|---|---|---|---|---|---|---|---|
| 1 | 100 | 100 | 10 | 20 | 300 | 5 | 27 |
| 2 | 100 | 100 | 20 | 20 | 300 | 5 | 32 |
| 3 | 100 | 100 | 30 | 20 | 300 | 5 | 34 |
| 4 | 100 | 100 | 40 | 20 | 300 | 5 | 36 |
| 5 | 100 | 200 | 10 | 20 | 300 | 5 | 9 |
| 6 | 100 | 300 | 10 | 20 | 300 | 5 | 17 |
| 7 | 100 | 500 | 10 | 20 | 300 | 5 | 15 |
| 8 | 100 | 1000 | 10 | 20 | 300 | 5 | 14 |
| 9 | 200 | 100 | 10 | 20 | 300 | 5 | 15 |
| 10 | 500 | 100 | 10 | 20 | 300 | 5 | 0.4 |
| 11 | 1000 | 100 | 10 | 20 | 300 | 5 | 0 |
| 12 | 100 | 400 | 10 | 20 | 300 | 40 | **91** |
| 13 | 100 | 500 | 10 | 20 | 300 | 40 | 84 |
| 14 | 100 | 400 | 10 | 20 | 300 | 5 | 18 |
| 15 | 100 | 400 | 10 | 20 | 300 | 10 | 51 |
| 16 | 100 | 400 | 10 | 20 | 300 | 20 | 59 |
| 17 | 100 | 400 | 10 | 20 | 300 | 30 | 80 |
| 18 | 100 | 200 | 10 | 20 | 300 | 40 | 69 |
| 19 | 100 | 300 | 10 | 20 | 300 | 40 | 82 |

**Table S3:** Attempted modification of Leuprolide Acetate Peptide for Optimal Photosensitizer **16** Turnover in presence of Ammonium Acetate (pH 6.9) Buffer. Conversions are averages of two replicates.

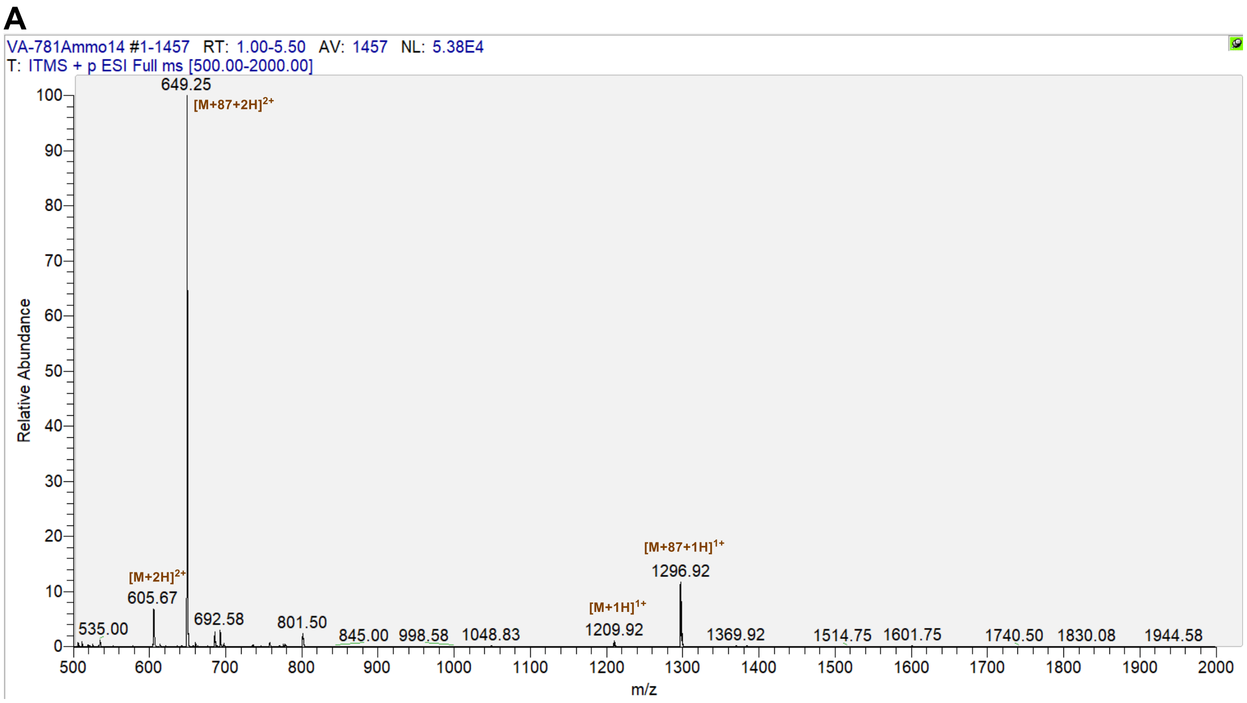


**Figure S7**. Mass spectrum of crude Leuprolide•OAc Acetate labelling from Table S2, entry 12.

**Tandem MS-MS Analysis of Leuprolide Acetate Peptide-*N*-methycarbamate Modification:**

Tandem MS data of Condition 12, Table S2 obtained from the crude Leuprolide Acetate Peptide-N-methycarbamate *via* LC/MS by selecting m/z = 1296.6 (+2 charge state) for collision induced dissociation (Voltage = 35 V).

Pyr—XH**W**SYLLRP—NHEt

| b+ | b2+ | # | Seq | # | y+ | y2+ |
|---|---|---|---|---|---|---|
| 112.0073 | 56.5073 | 1 | X | 9 | | |
| 249.0662 | 125.0367 | 2 | H | 8 | 1185.6054 | 593.3064 |
| **522.1775** | 261.5924 | 3 | **W** | 7 | **1048.5465** | 524.7769 |
| **609.2095** | 305.1084 | 4 | S | 6 | 775.4352 | 388.2212 |
| **772.2729** | 386.6401 | 5 | Y | 5 | **688.4032** | 344.7052 |
| **885.3569** | 443.1821 | 6 | L | 4 | 525.3398 | 263.1736 |
| **998.441** | 499.7241 | 7 | L | 3 | **412.2558** | 206.6315 |
| **1154.5421** | 577.7747 | 8 | R | 2 | 299.1717 | 150.0895 |
| | | 9 | P | 1 | 143.0706 | 72.0389 |

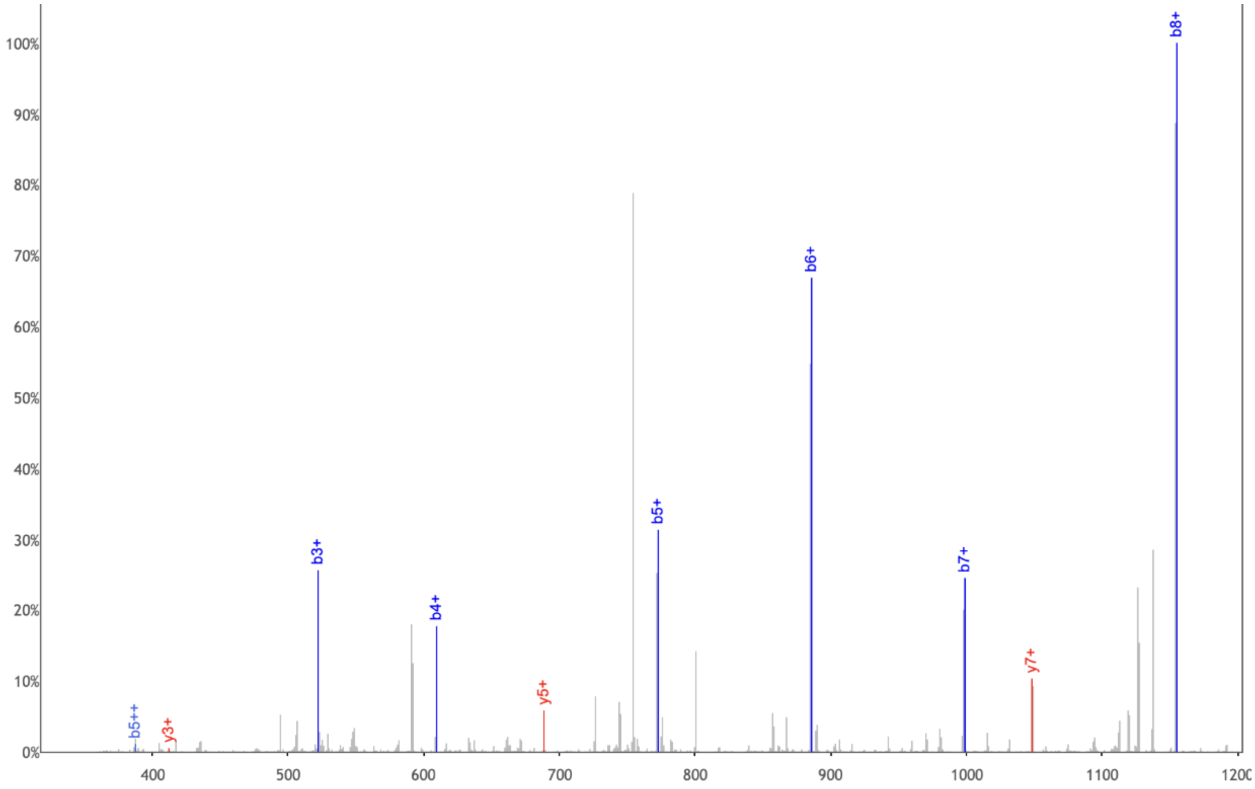


**Figure S8**. MS2 analysis of labelled Leuprolide from Table S2, entry 12.

##### 4.5 Trastuzumab labelling with 16 and 20a

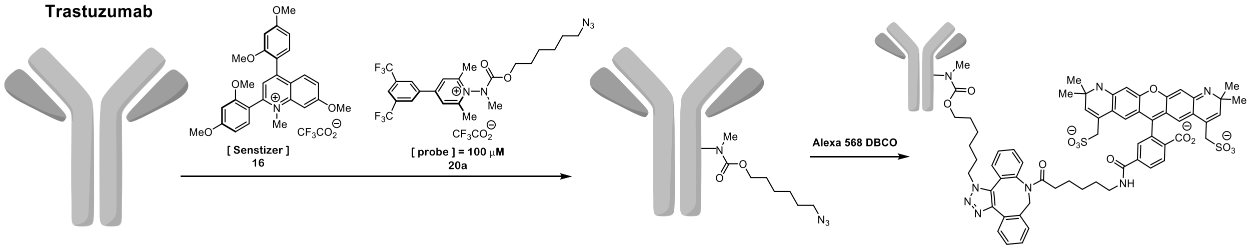


Procedure for Labelling: To a 200 μL microinsert (5x31mm Flat bottom vial, Fisher scientific, 13-622-208) containing a solution of Trastuzumab was added respective solutions of buffer (20 mM), probe **20a** (100 μM) and photosensitizer **16** (as indicated in Table S3) and nano pure water (final concentration of Trastuzumab = 7 μM, See Figure S8 A). The resulting solution was then irradiated using a HepatoChem photoredoxbox photoreactor equipped with a 427 nm Kessil PR160L LED (100% intensity) for 20 minutes (See Figure S8 C). At the end of irradiation, the solution was then dialyzed in a Slide-A-Lyzer Mini Dialysis Device (10K MWCO, Thermo Fisher Scientific) using 1x PBS (1.4 mL, pH 7.4) for 2 hr on an orbital shaker, replaced with fresh buffer 1x PBS (1.4 mL, pH 7.4), and dialyzed for an additional 12 hr on shaker. The purpose of the dialysis was to remove excess small molecules.

Procedure for Copper-Free Chemistry (SPAAC)[9-10]**:** All steps were performed in dark and over ice. In sequential order, labeled antibody (50 μL from dialysis conc. of mAb = 7 μM), pH 7.4 1x PBS buffer (20 mM), 10% DMF, and 50 μM of 2.5 mM DMSO stock solution of Alexa 568 DBCO were added to a 2 mL centrifuge and incubated on ice for 3 hr. At the end of incubation, an addition of 2.4 µL of 2.5 mM Alexa 568 DBCO was loaded into each tube, and the mixture was incubated overnight at 4 °C. At the end of irradiation, the solution in each tube was then dialyzed in a Slide-A-Lyzer Mini Dialysis Device (10K MWCO, Thermo Fisher Scientific) using 1x PBS (1.4 mL, pH 7.4) for 2 hr on an orbital shaker, replaced with fresh buffer 1x PBS (1.4 mL, pH 7.4), and dialyzed for an additional 12 hr on shaker.  The purpose of second dialysis was to remove unreacted dye.

**SDS-PAGE (Reductive Conditions):** Sodium dodecyl sulfate-polyacrylamide gel electrophoresis (SDS-PAGE) under reductive conditions was performed to identify the labeling site on either the heavy chains or light chains of Trastuzumab. 20 uL of a 7 µM of each DBCO-labelled trastuzumab were reduced with 20 µL Laemmli buffer (2x) containing 10 mM DTT and heated at 85 °C for 10 minutes. Reduced samples were loaded and separated on 4-20% Mini-Protean® TGM™ Precast Gels (Bio-Rad, Cat. No. 4561094) at 100V for 90 min. Precision Plus Protein Dual Color Standard (BioRad) was used for molecular weight comparison. Fluorescence images were recorded using using an Analytic Jena Chemstudio Plus geldoc. The band intensity of heavy chain and light chain analyzed using ImageJ.

| Lane | [mAb] | Probe (20a) | Sensitizer (16) | pH [20 mM] | Time (min) |
|---|---|---|---|---|---|
| 1 | 7 µM | - | - | - | - |
| 2 | 7 µM | 100 µM | 5 µM | $NH_4OAc$ 6.9 | 20 |
| 3 | 7 µM | 100 µM | 10 µM | $NH_4OAc$ 6.9 | 20 |
| 4 | 7 µM | 100 µM | 5 µM | $Na_2HPO_4$ 7.4 | 20 |
| 5 | 7 µM | 100 µM | 10 µM | $Na_2HPO_4$ 7.4 | 20 |

**Table S4:** Conditions assayed for Trastuzumab labelling.

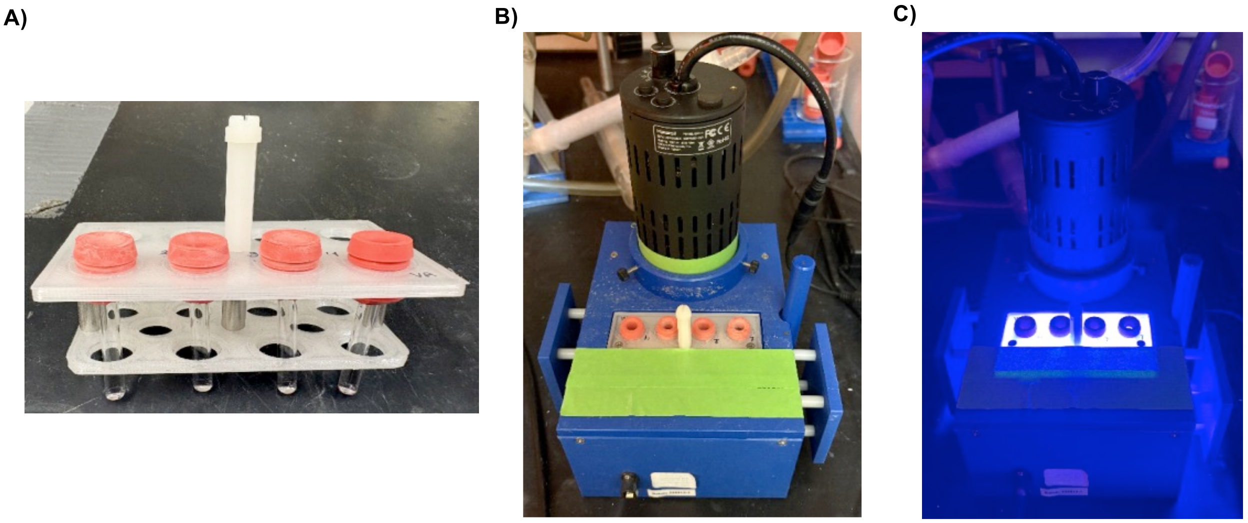

**Figure S9:** (A) Microvial arrangement in holder used for photochemical labeling of trastuzumab (anti-HER2) with a total reaction volume of 50 µL. (B) Sample holder setup in the photoredox box prior to irradiation. (C) Sample during illumination with a 427 nm Kessil lamp, positioned in the photoredox box to ensure uniform light exposure.

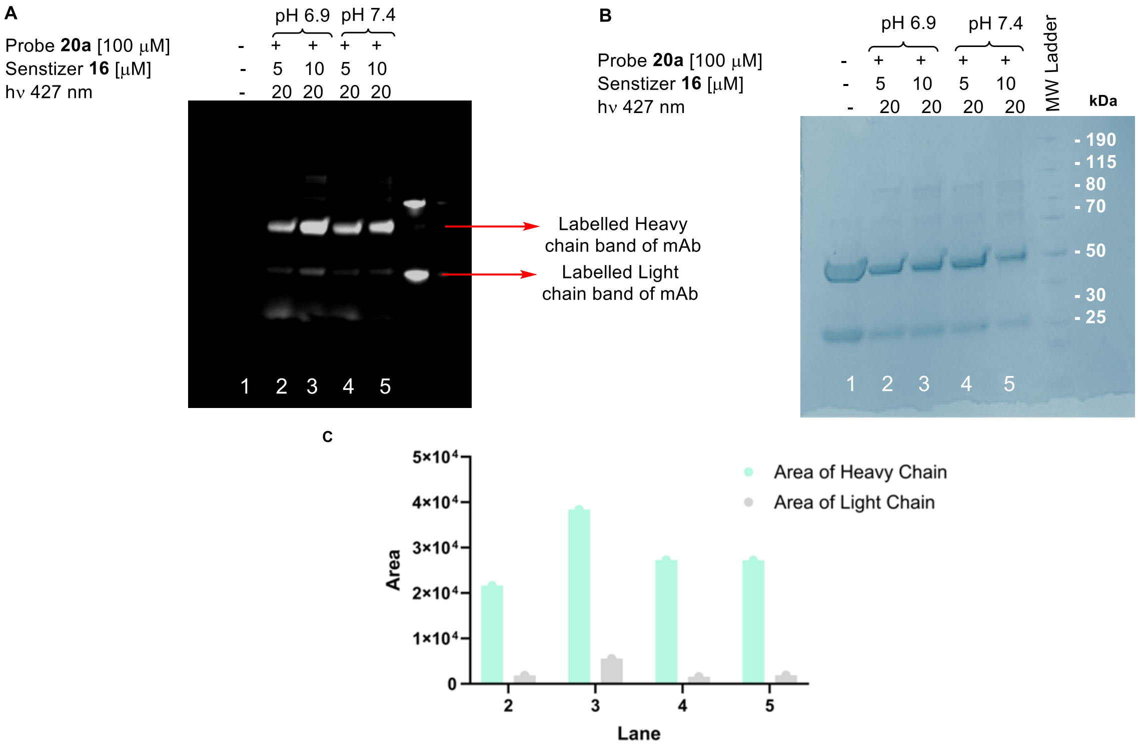


**Figure S10:** (A) SDS-PAGE gel displays direct comparison of bioconjugation of monoclonal antibody between ammonium acetate 6.9 vs phosphate buffer 7.4, (B) Corresponding Coomassie stain, (C) Relative band intensity of labelled heavy chain and light chain analyzed using ImageJ.

#### 5. Mechanistic studies

##### Electrochemistry:

**General methods and materials:** Electrochemical analyses were conducted inside an Argon-filled MBraun Unilab glovebox using a BioLogic SP-200 potentiostat/galvanostat and the EC-Lab® software (v11.33) from BioLogic Science Instruments. For convenience, potentials are expressed versus the internal reference electrode $AgNO_3/Ag$.

**Cyclic voltammetry:** Cyclic voltammograms (CV) and Differential pulse voltammetry (DPV) were measured in a three-electrode electrochemical cell, consisting of a platinum wire counter electrode (Ec), a $AgNO_3/Ag$ reference electrode ($E_{ref}$, 0.01 M $AgNO_3$ in 0.1 M $TBAPF_6$ in $CH_3CN$), and a glassy carbon working electrode ($E_w$, 0.071 cm², CH Instrument, Inc.). The working electrode was polished prior to each record using aluminum oxide on polishing paper and anhydrous $CH_3CN$ to remove residual particles.

**Measurement of cyclic voltammetry (CV) of compound optimal Pyridinium Probe (20):**

Solution for the cyclic voltammetry study was prepared with1.5 mM of the optimal pyridinium probe **20** 0.1 M $TBAPF_6$ in $CH_3CN$ at 298 K, recorded at 100 mV/s.

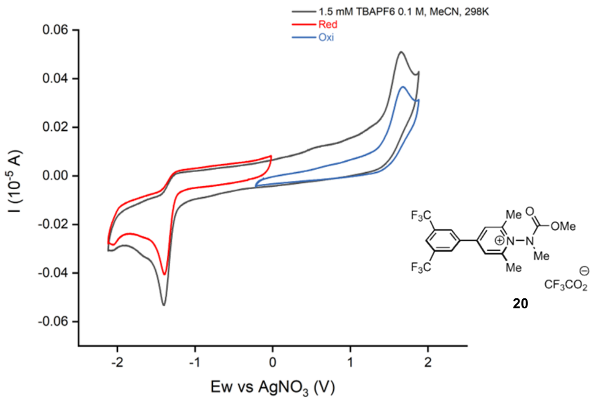


**Figure S11:** Cyclic voltammograms of optimal pyridinium probe **20** at a scan rate of 100 mV/s

**Measurement of cyclic voltammetry (CV) of selected diarylquinolinium aromatic cation sensitizer (16):**

Solution for the cyclic voltammetry study was prepared with1.1 mM of the optimal sensitizer **16** 0.1 M TBAPF6 in $CH_3CN$ at 298 K, recorded at 100 mV/s.

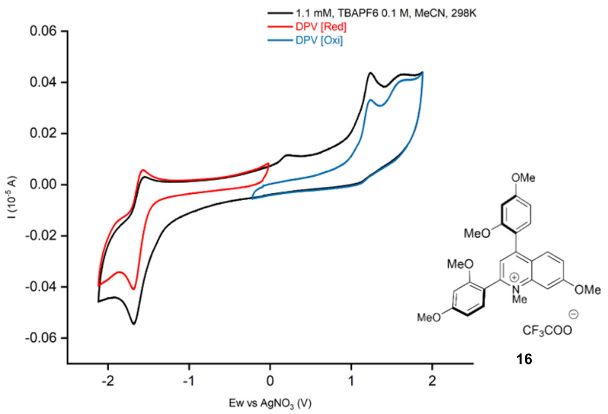


**Figure S12.** Cyclic voltammograms of diarylquinolinium aromatic cation sensitizer (**16**) at a scan rate of 100 mV/s

#### Photophysical properties of diarylquinolinium aromatic cation sensitizer (16):

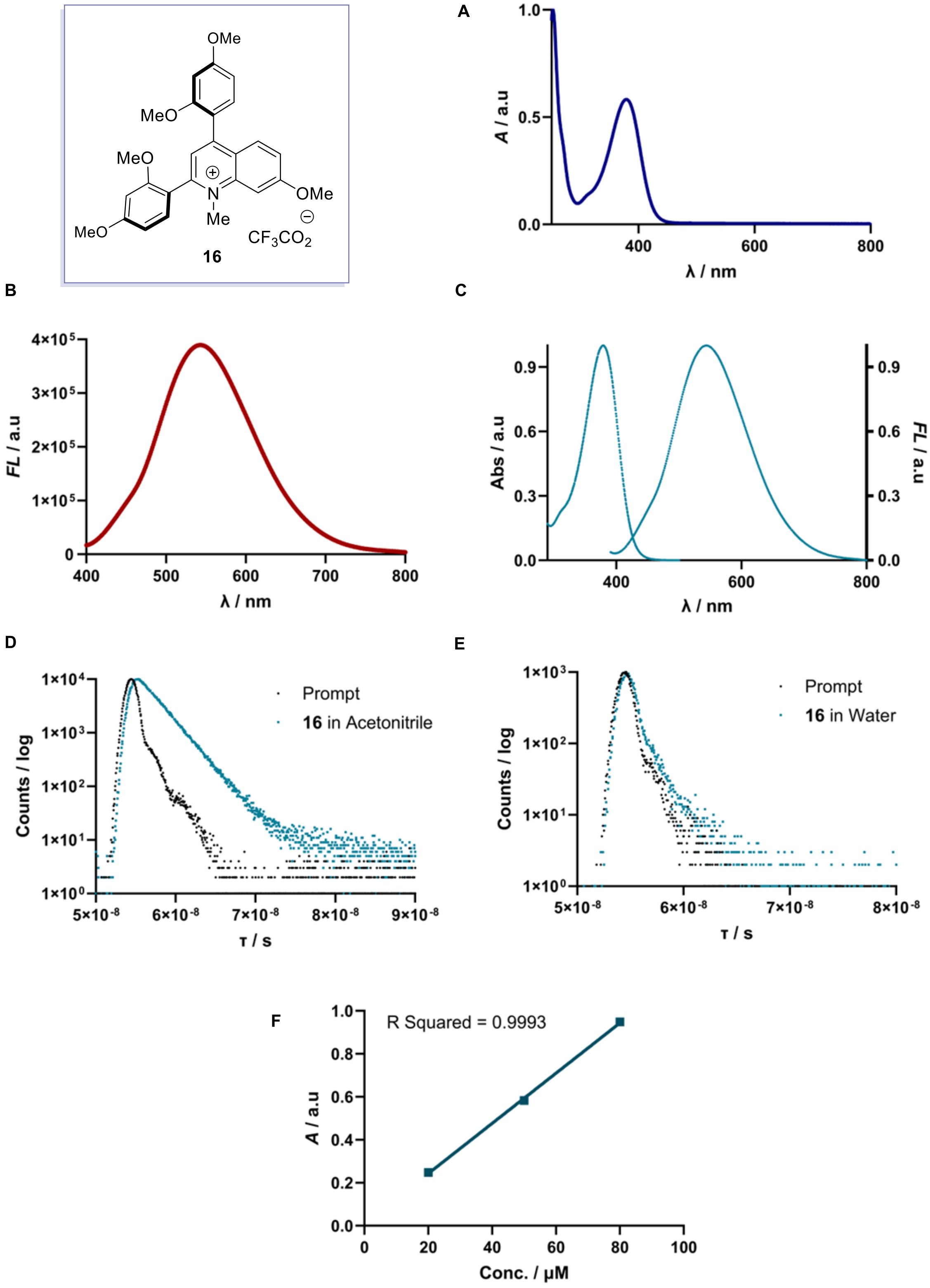


**Figure S13:** (A) Absorbance of compound **16** in 20 mM $Na_2HPO_4$ 7.4. (B) Emission of compound **16** in 20 mM $Na_2HPO_4$ 7.4. (C) Normalized Absorbance and Emission of compound **16** in 20 mM

$Na_2HPO_4$ 7.4. (D) Fluorescence lifetimes of **16** in Acetonitrile ($\tau_1$ = 2.23 ns $\chi^2$ = 1.08) was measured using NanoLED-370 (peak wavelength 367 nm, bandpass = 5 nm), IRF = Instrument Response Function; Fluorescence lifetimes of **16** in $H_2O$ ($\tau_1$ = 0.25 ns and $\tau_2$ = 4.54 ns, $\chi^2$ = 1.08) was measured using NanoLED-370 (peak wavelength 367 nm, bandpass = 5 nm), IRF = Instrument Response Function. (F) Linear fitting plot of the relationship between the concentration and absorbance of **16** (10, 20, and 50 μM in 20 mM $Na_2HPO_4$ 7.4).

#### Perturbation studies

A 2 mL Pyrex LC/MS vial containing the desired combination of lysozyme, $Na_2HPO_4$ buffer, L-Glutathione), (**20**, 100 μM) and **16** (10 μM) was diluted to a final volume of 500 μL with Di-$H_2O$ and the solutions sparged with $N_2$ for 60 minutes. The solution was then irradiated using a HepatoChem photo-redox box (model # HCK1006-01-016) equipped with a 427 nm Kessil PR160L LED (100% intensity) for 20 minutes with active degassing. The resultant reaction mixture was then analyzed directly via LC/MS. Labelling outcomes were analyzed using LC/MS. 10 μM of internal standard was then added and the resulting solution analyzed directly by LC/MS. An identical solution that was not irradiated was also analyzed by LC/MS to determine response factors.

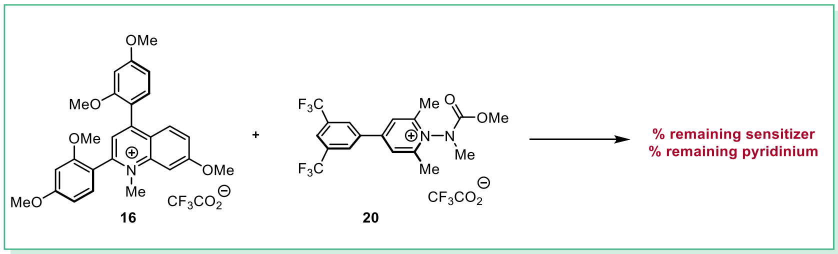


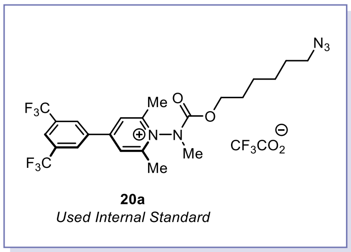


| Entry | [Protein] μM | [Pyridinium] μM | [sensitizer] μM | [GSH] μM | [pH 7.4 PO4] mM | Light? 20 min | % remaining Sensitizer | % remaining Probe |
|---|---|---|---|---|---|---|---|---|
| 1 | 10 | 100 | 10 | 300 | 20 | yes | 96 | 10 |
| 2 | 0 | 100 | 10 | 300 | 20 | yes | 75 | 9 |
| 3 | 0 | 100 | 10 | 0 | 20 | yes | 97 | 82 |
| 4 | 0 | 0 | 10 | 0 | 20 | yes | 98 | N/A |
| 5 | 0 | 0 | 10 | 0 | 0 | yes | 102 | N/A |
| 6 | 10 | 100 | 10 | 300 | 20 | no | 96 | 97 |
| 7 | 0 | 100 | 10 | 0 | 0 | yes | 96 | 87 |
| 8 | 0 | 0 | 10 | 300 | 0 | yes | 86 | NA |
| 9 | 10 | 100 | 10 | 0 | 20 | yes | 97 | 92 |

**Table S5:** Results for remaining percentage of sensitizer (**16**) and probe (**20**) after completion of irradiation using LC-MS Method C. Average of two replicates.

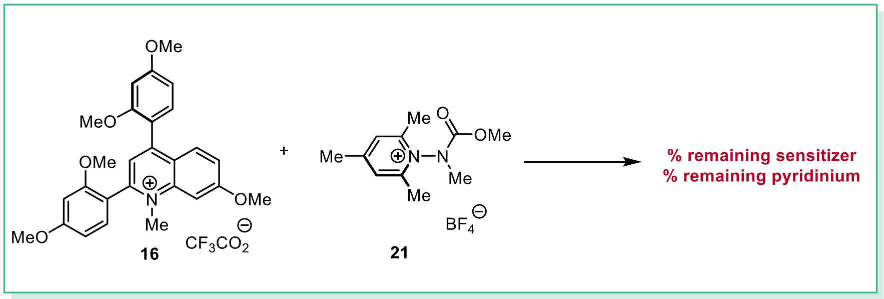


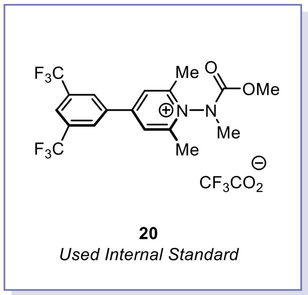


| *Entry* | *[Protein] µM* | *[probe] µM* | *[sensitizer] µM* | *[GSH] µM* | *[pH 7.4 $PO_4$] mM* | *Light? 20 min* | *% remaining probe* | *% remaining sensitizer* |
|---|---|---|---|---|---|---|---|---|
| 1 | 10 | 100 | 10 | 300 | 20 | yes | 0 | ≥95 |
| 2 | 0 | 100 | 10 | 300 | 20 | yes | 0 | ≥95 |
| 3 | 0 | 100 | 10 | 0 | 20 | yes | 96 | ≥95 |
| 4 | 0 | 100 | 10 | 0 | 0 | yes | 98 | 95 |

**Table S6:** Results for remaining percentage of sensitizer (**16**) and probe (**21**) after completion of irradiation using LC-MS Method D. Average of two replicates.

#### 6. Biology

##### Cell cultures

Adherent HEK 293T and HeLa cells were cultured at 37 °C under 5% $CO_2$ in complete growth DMEM (Dulbecco's Modified Eagle Medium (Genesee), high glucose and with L-glutamine, supplemented with 10% FBS (Genesee) and 100 units/mL (or 100 µg/mL) penicillin/streptomycin (Thermo Scientific) respectively). Cells were split at 80 – 90% confluency for up to 15 passages.

##### Cell lysate preparation for lysate-based studies

HEK293T cells were cultured in T75 flasks (Thermo Scientific) and harvested with 0.25% Trypsin-EDTA (Thermo Scientific) after reaching 80 – 90% confluency. Only cell batches with viability greater than 90% were selected for lysate generation. Cell pellets were washed once with D-PBS (Thermo Scientific), then flash-frozen with liquid $N_2$ and stored at - 80 °C until use. Cell pellets were resuspended in 50 mM HEPES, pH 7.5, 150 mM NaCl, 1.5 mM $MgCl_2$, 1% NP-40, and 1x Halt Protease Inhibitor (Thermo Scientific). The cell suspension was chilled on ice for 30 min, followed by sonication (QSonica) for 30 s (2 s on, 3 s off) at 20% amplitude on ice. Cell debris was removed via centrifugation at 20,000 g for 20 min at 4 °C. The supernatant was transferred to a clean tube, and the protein concentration was measured by Pierce™ 660nm Protein Assay Kit (Thermo Scientific)

##### Fixed cells imaging (dose-response effect)

30, 15, 6, 0 µL of 10 mM **20a** (in DMSO) and 45, 30, 15, 0 µL of 2 mM **16** (in nano-pure water) were diluted together in 2.925, 2.955, 2.979, 3 mL of Opti-MEM (Gibco) respectively to make three treating solutions containing 100 µM **20a** + 30 µM **16**, 50 µM **20a** + 20 µM **16**, and 20 µM **20a** + 10 µM **16**. $2.5\times10^5$ HeLa cells (viability > 90%) were seeded in 35 mm dishes (No. 1.5 coverslip, 14 mm glass diameter) (Mattek) for 24 h at 37 °C under 5% $CO_2$ in complete growth DMEM. Cells were washed with D-PBS (2 x 610 µL), then treated with 2 mL of **20a/16** solutions in Opti-MEM for 1 h at 37 °C. The treated cells were illuminated with 427 nm Kessil lamp for 20 min at 4 °C (Figure S14). After washing with 1x PBS (3 x 610 µL), cells were fixed with 3 mL of 4% paraformaldehyde in 1x PBS (Thermo Scientific) for 20 min at room temperature. Cells were washed once with 610 µL of 1x PBS, followed by permeabilization with 1.5 mL of 1x PBS, 0.1% Triton X-100 for 10 min at room temperature. After washing with 1x PBS (3 x 610 µL each), cells were incubated with 1.5 mL of 3 µM DBCO-AF568 (in nano-pure water) (CCT-1294-1, VectorLabs) in the dark for 1.5 h at room temperature. Cells were then washed several times with the following buffers: 1x PBS, 0.5% Triton X-100 (3 x 610 µL); 1x PBS, 0.1% Triton X-100 (3 x 610 µL); and 1x PBS (3 x 610 µL). The cells were further stained with 1.5 mL of 1 µg/mL DAPI (Invitrogen) in the dark for 10 min at room temperature, followed by 3 washes with 1x PBS, 0.1% Tween-20 (610 µL each) and 1x PBS (3 x 610 µL). Finally, cells were covered with 1 mL of 90% glycerol in 1x D-PBS containing 1% (w/v) sodium ascorbate and stored at 4 °C and protected from light. Cells were imaged within 24 h after preparation using Zeiss LSM880 Inverted Confocal Microscope. DAPI was excited using a 405 nm laser, and the emitted light was observed between 410 – 550 nm. Meanwhile, AF568 was excited using a 561 nm laser, and emission was captured between 580 – 690 nm. The images were acquired using Zen (Black edition) software with a 63x objective (oil immersion) and were processed with FIJI.

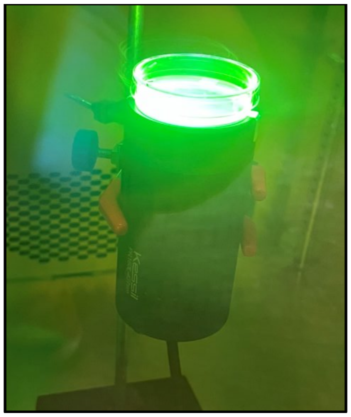


**Figure S14:** Photochemical setup for fixed cell imaging.

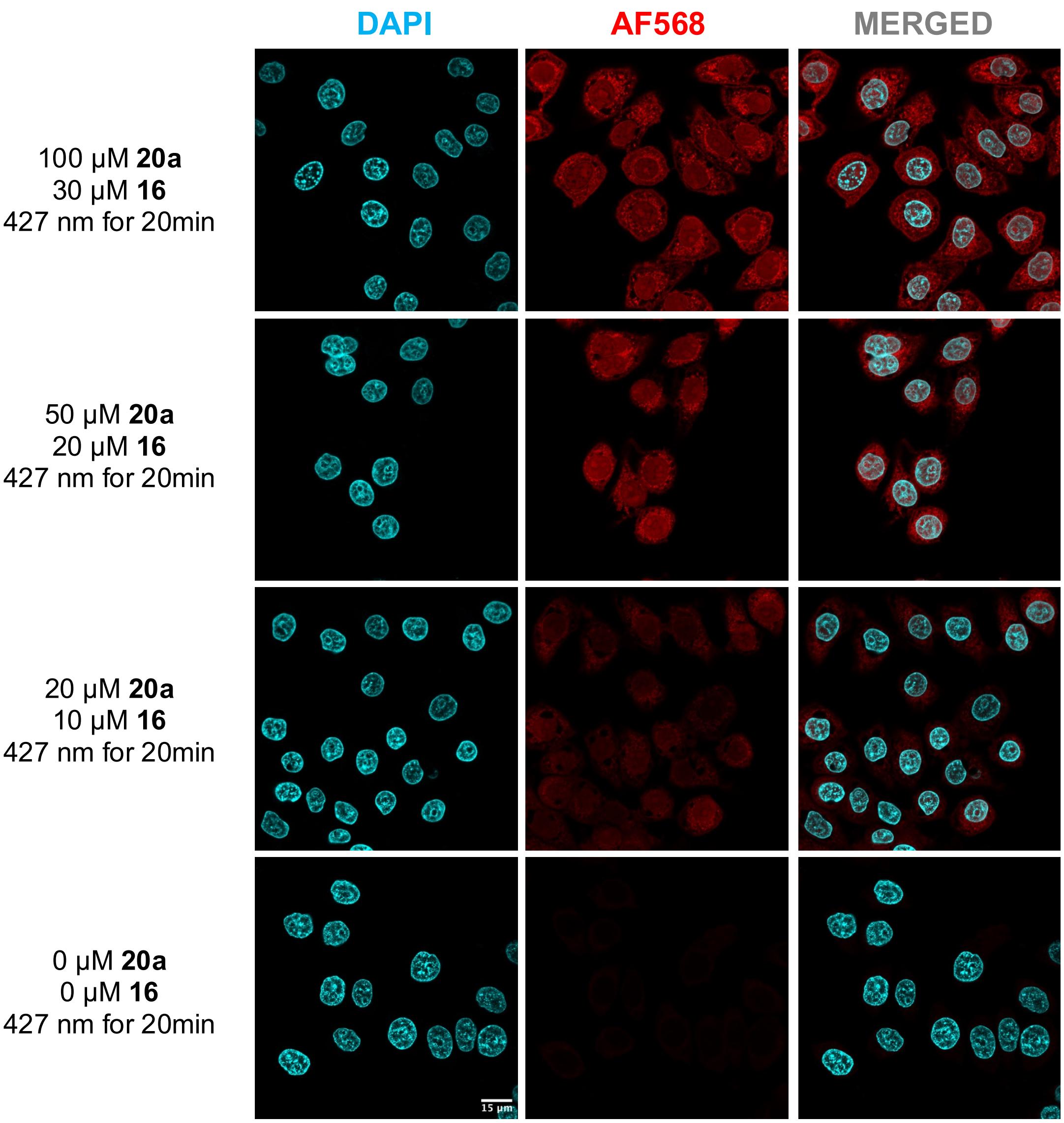


**Figure S15:** Dose-response effect of the photocatalysis chemistry visualized via fluorescent microscopy.

**Fixed cells imaging (comparison between light versus no light effect)**
30 µL of 10 mM **20a** (in DMSO) and 45 µL of 2 mM **16** (in nano-pure water) were diluted together in 2.925 mL of Opti-MEM (Gibco) to make a treating solution containing 100 µM **20a** and 30 µM **16**. A different treating solution only contained PBS, also prepared in Opti-MEM. $2.5\times10^5$ HeLa cells (viability > 90%) were seeded in 35 mm dishes (No. 1.5 coverslip, 14 mm glass diameter) (Mattek) for 24 h at 37 °C under 5% $CO_2$ in complete growth DMEM. Cells were washed with D-PBS (2 x 610 µL), then treated with 2 mL of **20a/16** solutions or PBS in Opti-MEM for 1 h at 37 °C. The treated cells were illuminated with 427 nm Kessil lamp for 20 min at 4 °C (else indicated) (Figure S14). After washing with 1x PBS (3 x 610 µL), cells were fixed with 3 mL of 4% paraformaldehyde in 1x PBS (Thermo Scientific) for 20 min at room temperature. Cells were washed once with 610 µL of 1x PBS, followed by permeabilization with 1.5 mL of 1x PBS, 0.1% Triton X-100 for 10 min at room temperature. After washing with 1x PBS (3 x 610 µL), cells were incubated with 1.5 mL of 3 µM DBCO-AF568 (in nano-pure water) (CCT-1294-1, VectorLabs) in the dark for 1.5 h at room temperature. Cells were then washed several times with the following buffers: 1x PBS, 0.5% Triton X-100 (3 x 610 µL); 1x PBS, 0.1% Triton X-100 (3 x 610 µL); and 1x PBS (3 x 610 µL). The cells were further stained with 1.5 mL of 1 µg/mL DAPI (Invitrogen) in the dark for 10 min at room temperature, followed by 3 washes with 1x PBS, 0.1% Tween-20 (610 µL each) and 1x PBS (3 x 610 µL). Finally, cells were covered with 1 mL of 90% glycerol in 1x D-PBS containing 1% (w/v) sodium ascorbate and stored at 4 °C and protected from light. Cells were imaged within 24 h after preparation using Zeiss LSM880 Inverted Confocal Microscope. DAPI was excited using a 405 nm laser, and the emitted light was observed between 410 – 550 nm. Meanwhile, AF568 was excited using a 561 nm laser, and emission was captured between 580 – 690 nm. The images were acquired using Zen (Black edition) software with a 63x objective (oil immersion) and were processed with FIJI.

**Fixed cell imaging attempts using CuAAC**
$1.0\times10^5$ HEK293T cells (viability > 90%) were seeded in 35 mm dishes (No. 1.5 coverslip, 14 mm glass diameter) for 24 h at 37 °C under 5% $CO_2$ in complete growth DMEM. Cells were washed with D-PBS (2 x 60 µL each), then treated with 200 µL of Opti-MEM without any sensitizer or pyridinium salts for 1 h at 37 °C. The treated cells were illuminated with 427 nm Kessil lamp for 20 min at 4 °C. After washing with 1x PBS (3 x 60 µL), cells were fixed with 150 µL of 4% paraformaldehyde in 1x PBS for 20 min at room temperature. Cells were washed once with 60 µL of 1x PBS, followed by permeabilization with 150 µL of 1x PBS, 0.1% Triton X-100 for 10 min at room temperature. After washing with 1x PBS (3 x 60 µL), cells were incubated twice (15 min each) with 100 µL of 100 µM N-methyl maleimide (Sigma Aldrich). Cells were washed with 1x PBS (3 x 60 µL). The click-chemistry cocktail was prepared by combining the following reagents: 500 µL of 200 mM phosphate buffer pH 7.5, 25 µL of 200 mM $CuSO_4$ (Sigma Aldrich), 6 µL of 500 µM AF568-alkyne (CCT-1293-1, VectorLabs), 285 µL of 100 mM sodium L-ascorbate (Sigma Aldrich), and 184 µL of nano-pure water. Cells were incubated with 100 µL of the click-chemistry cocktail for 5 min at room temperature in the dark. Cells were then washed several times with the following buffers: 1x PBS, 0.5% Triton X-100 (1 x 60 µL) and 1x PBS (3 x 60 µL each). Cells were further stained with 200 µL of 1 µg/mL DAPI in the dark for 10 min at room temperature, followed by 3 washes with 1x PBS (60 µL each). Finally, cells were covered with 50 µL of 90% glycerol in 1x D-PBS containing 1% (w/v) sodium ascorbate and stored at 4 °C and protected from light. Cells were imaged within 24 h after preparation using Zeiss LSM880 Inverted Confocal Microscope. DAPI was excited using a 405 nm laser, and the emitted light was observed between 410 – 550 nm. Meanwhile, AF568 was excited using a 561 nm laser, and emission was captured between 580 – 690 nm. The images were acquired using Zen (Black edition) software with a 63x objective (oil immersion, 2x zoom) and were processed with FIJI.

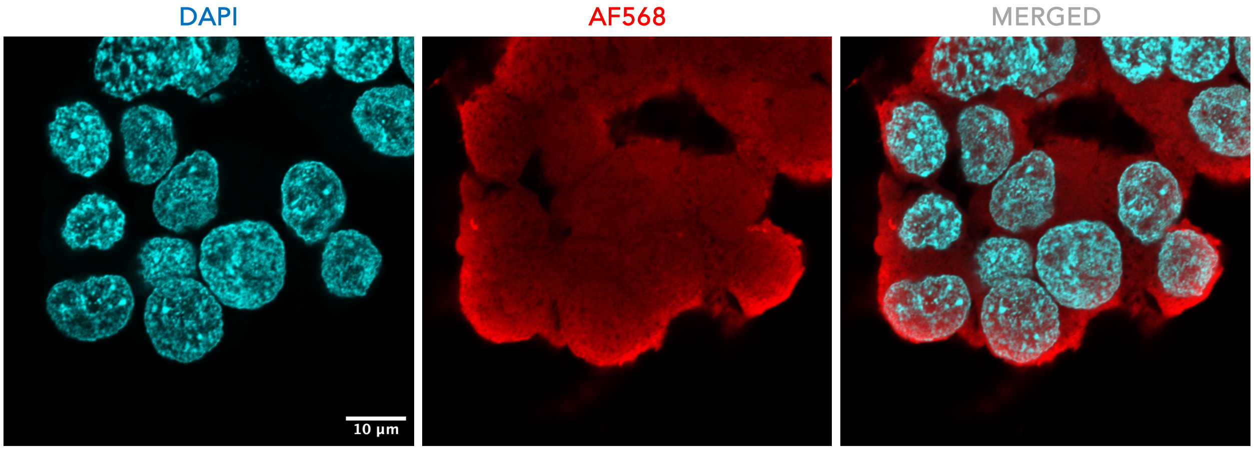


**Figure S16:** Non-specific copper-catalyzed click-chemistry in fixed cell imaging visualized with fluorescent microscopy

**Immunoblotting**

Protein samples were denatured at 95 °C for 10 min in Laemmli buffer containing 60 mM DTT and then separated by either 4 – 20% Mini-Protean® TGM™ Precast Gels or 4–20% Criterion™ TGX™ Precast Midi Protein Gels (Bio-Rad) at 100-130 V for 90 min then transferred onto 0.45 µm nitrocellulose membranes (Bio-Rad) at 100 V for 1 h using the Mini Trans-Blot® Cell (Bio-Rad). The membranes were washed with 1x PBST, 0.1% Tween-20 for 5 min. The membranes were blocked with EveryBlot Blocking Buffer (Bio-Rad) at 4 °C for 16 h. The membranes were incubated with HRP-linked Streptavidin (Abcam) at 4 °C for 16 h. After washing with 1x PBST, 0.1% Tween-20 (6 x 5 min), the membranes were visualized by incubating with the Clarity™ Western ECL Substrate for 2 min (Bio-Rad). The membrane was imaged with UVP ChemStudio PLUS Imaging Systems, Analytik Jena (Avantor®). FIJI was used to estimate the relative intensity of the protein bands.

**Cell lysate photolabeling with 16 and 20b (or 2)**

2 mg/mL (100 µg) of native HEK293T lysates in 50 mM HEPES, pH 7.5, 150 mM NaCl, 1.5 mM $MgCl_2$, 1% NP-40 were treated with 5, 10, 50 µM (else indicated) of **16** (2.5 mM stock solution in nano-pure water) and 100 µM (else indicated) of **20b** (or **2**) (2.5 mM stock solution in nano-pure water) with an addition of 20 mM $NaH_2PO_4/Na_2HPO_4$ buffer, pH 7.5. The reaction mixtures were mixed well in 1.5 mL Eppendorf tubes and then transferred to 200 µL flat-bottom insert vials (Fisher Scientific). The reaction mixture was illuminated with 427 nm Kessil lamp for 20 min (else indicated) at room temperature (Figure S17), then directly concentrated and exchanged with 20 mM $NaH_2PO_4/Na_2HPO_4$ buffer, pH 7.5 using Amicon® Ultra Centrifugal 3K spinning columns (Sigma Millipore).

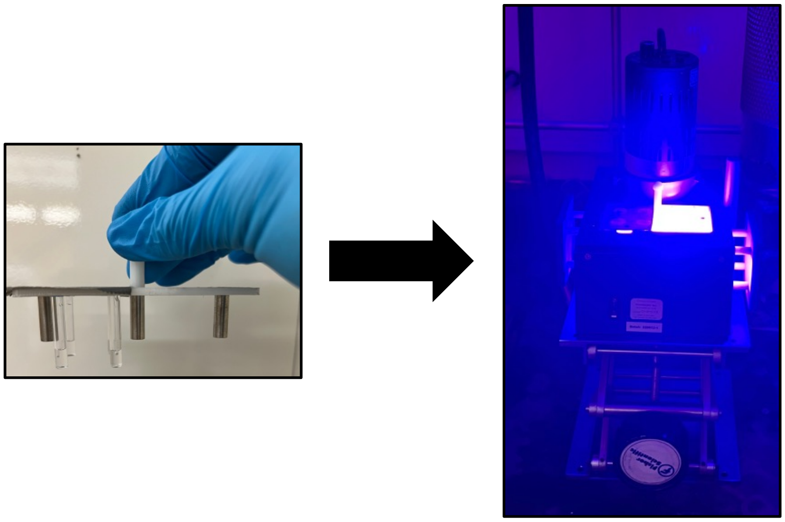

**Figure S17:** Photochemical setup for labeling with **16** and **20b** (or **2**) using 100 µg HEK293T lysate.

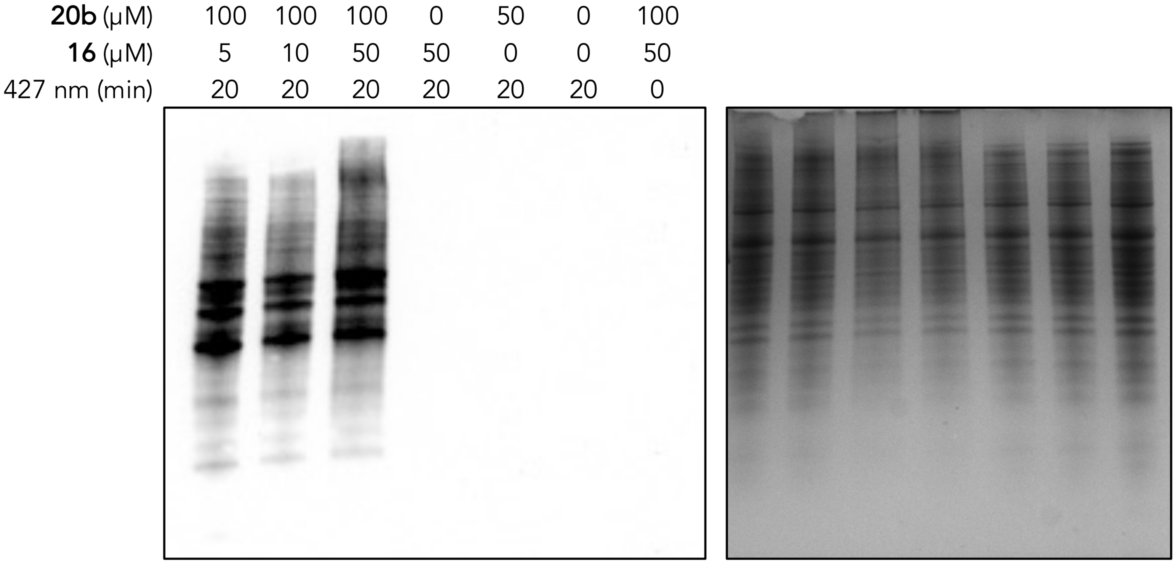


**Figure S18:** Photolabeling of 100 µg HEK293T lysate with **16** and **20b**. Western blot (left), Coomassie stain (right)

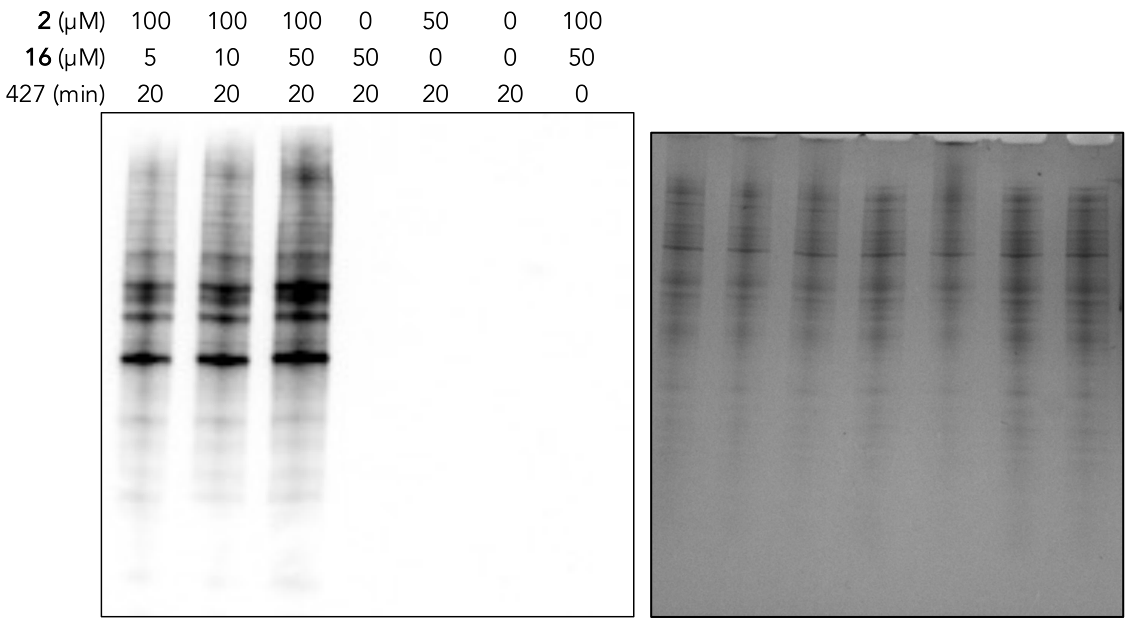


**Figure S19:** Photolabeling of 100 μg HEK293T lysate with **16** and **2**. Western blot (left), Coomassie stain (right)

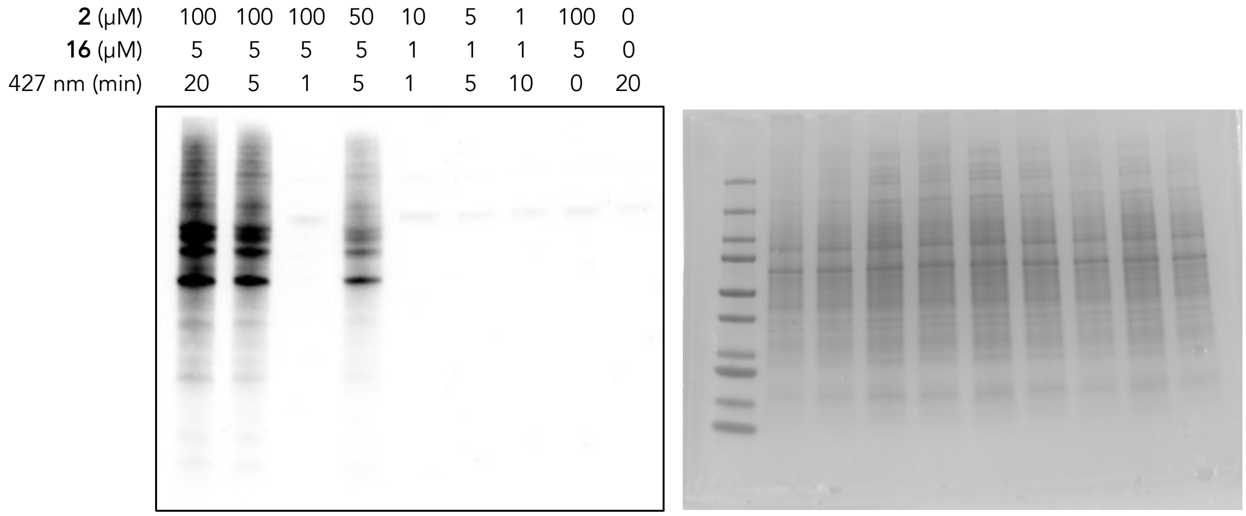


**Figure S20:** Photolabeling of 100 μg HEK293T lysate with varying concentrations of **16** and **2** and different irradiation times. Western blot (right), Coomassie stain (right)

**Cell lysate photolabeling with 16 and 20a (changing 20a:16 ratio)**

2 mg/mL (500 μg) of native HEK293T cell lysates were treated with 2.5, 5, 10, 20, 50 μM of **16** (else indicated) (100 μM stock solution in nano-pure water) and 5, 10, 20, 40, 50, 80, 100 μM of **20a** (else indicated) (500 μM stock solution in DMSO) in 50 mM HEPES pH 7.5, 150 mM NaCl, 1.5 mM $MgCl_2$, 1% NP-40. The reaction mixtures were mixed well in LC-MS vials (VWR) and incubated at room temperature for 30 min protected from light. The reaction mixture was illuminated with 427 nm Kessil lamp for 20 min (else indicated) at room temperature (Figure S21). Protein samples were precipitated with methanol:chloroform and resuspended in 235.7 μL of 1x PBS pH 7.4, 0.1% Triton. The protein solutions were combined with 3.125 μL of 20 mM $CuSO_4$, 2.5 μL of 50 mM BTTP (VectorLabs), 2.5 μL of 10 mM biotin-alkyne (Figure S22)[11], and 6.25 μL of 100 mM sodium L-ascorbate. The click reactions were agitated on a shaker for 60 min at room

temperature, then quenched with 25 µL of 50 mM EDTA, pH 8.0. Protein samples were precipitated with methanol:chloroform and resuspended in 2x Laemmli buffer containing 60 mM DTT (Thermo Scientific)

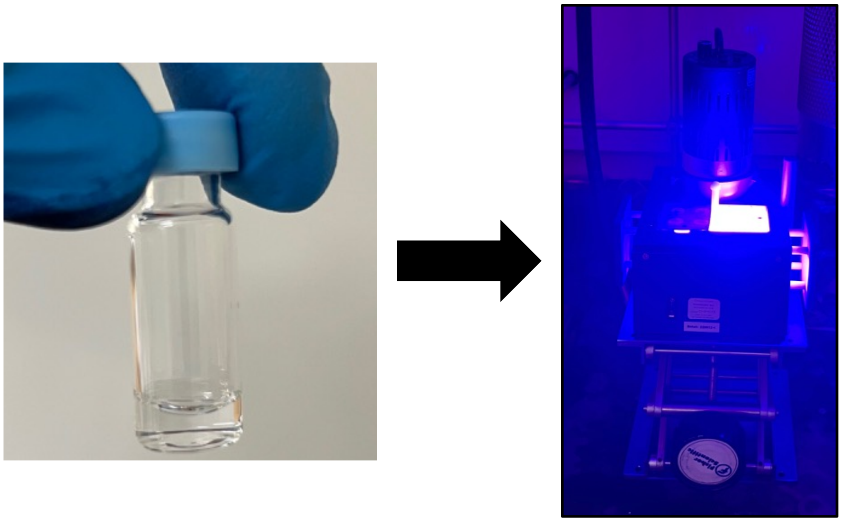

**Figure S21:** Photochemical setup for labeling with **16** and **20a** using 500 µg of HEK293T lysate.

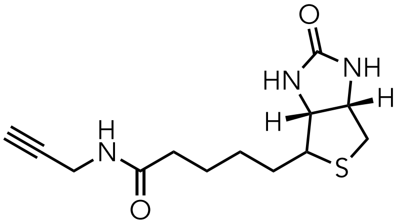

**Figure S22:** Biotin-alkyne structure used for *CuAAC* for Western analysis

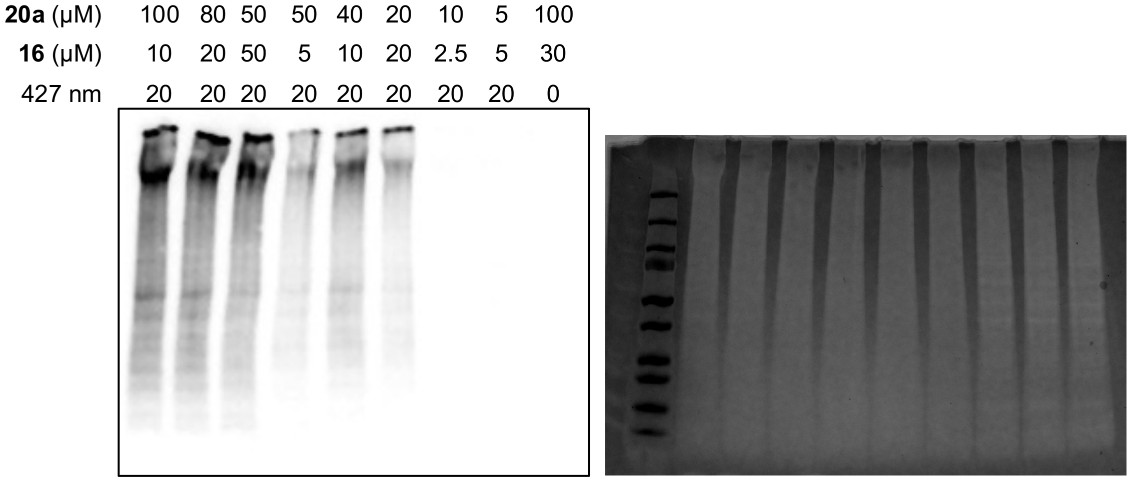


**Figure S23:** Different photolabeling outcomes when 500 µg of HEK293T lysate was exposed to varying **20a**:**16** ratios. Western blot (left), Zinc stain (right)

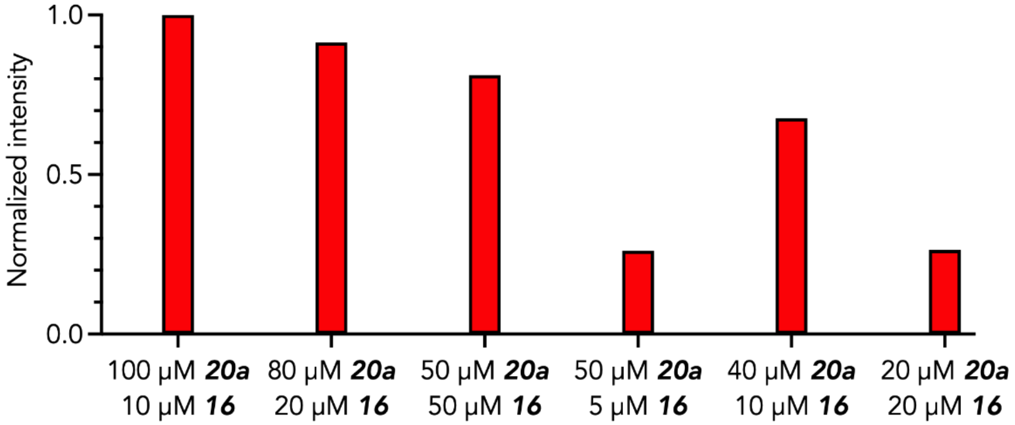


**Figure S24:** Relative band intensity for different labeling outcomes when 500 µg of HEK293T lysate was exposed to varying **20a**:**16** ratios

**Comparisons of 20b and 2 in live HEK293T cell photolabelling with 16**

$4.3\times10^6$ HEK293T cells (viability > 90%) were seeded in T25 flasks (Corning) for 24 h at 37 °C under 5% $CO_2$ in complete growth DMEM. Various probe-catalyst solutions, including 100 µM **20b** (or **2**) + 20 µM **16**, 100 µM **20b** (or **2**) + 30 µM **16**, 50 µM **20b** (or **2**) + 20 µM **16**, and 20 µM **20b** (or **2**) + 10 µM **16**, were prepared in Opti-MEM. Cells were washed with D-PBS (2 x 900 µL) then treated with 2.7 mL of **20b (or 2)/16** solutions for 1 h at 37 °C. The treated cells were illuminated with 427 nm Kessil lamp by placing the T25 flask on the photoreactor box for 0, 5, 10, or 20 min at 4 °C (Figure S25). Cells were washed once with D-PBS then collected with ice-cold D-PBS. Cell pellets were resuspended in 50 mM HEPES, pH 7.5, 150 mM NaCl, 1.5 mM $MgCl_2$, 1% NP-40, and 1x Halt Protease Inhibitor (Thermo Scientific). The cell suspension was sonicated (QSonica) for 30 s (2 s on, 3 s off) at 40% amplitudes. Cell debris was removed *via* centrifugation at 20,000 g for 20 min at 4 °C. The supernatant was transferred to a clean tube, and the protein concentration was determined using Pierce™ 660nm Protein Assay Kit.

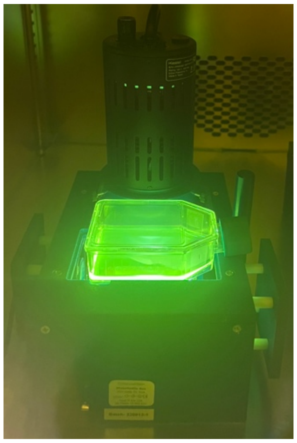

**Figure S25:** Photochemical setup for live cell labeling

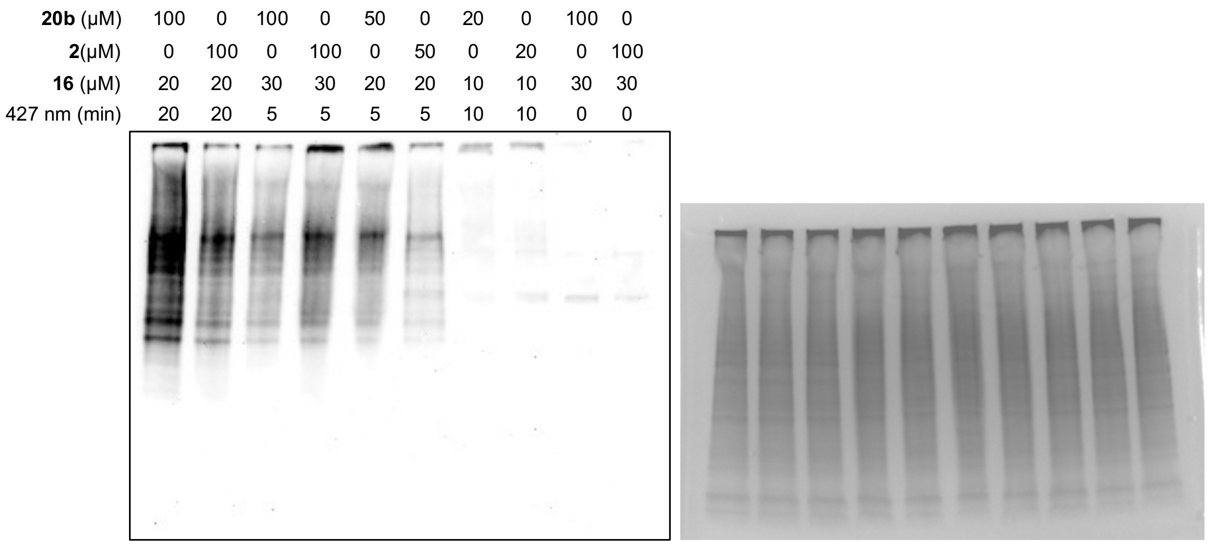


**Figure S26:** Photolabeling trend comparison between **20b** and **2**. Western blot (left), Coomassie stain (right)

**Live HEK293T cells photolabeling with 16 and 20a for Western analysis**
$4.0{\times}10^{6}$ HEK293T cells (viability > 90%) were seeded in T25 flasks (Corning) for 20 h at 37 °C under 5% $CO_2$ in complete growth DMEM. 2 identical culture flasks were prepared for each condition. Various **16** + **20a** solutions were prepared similarly in the above procedure (specific concentration would be indicated for each individual experiment). Cells were washed with D-PBS (2x900 μL), then treated with 2.7 mL of **16** + **20a** solutions for 1 h at 37 °C. The treated cells from both identical T25 flasks were illuminated simultaneously with 427 nm Kessil lamp by placing the flask on the photoreactor box for 0, 5, 10, or 20 min at 4 °C (Figure S23). Cells from both T25 flasks were washed once with D-PBS, then collected with ice-cold D-PBS and combined into an Eppendorf tube. Cell pellets were resuspended in 50 mM HEPES, pH 7.5, 150 mM NaCl, 1.5 mM $MgCl_2$, 1% NP-40, and 1x Halt Protease Inhibitor (Thermo Scientific). The cell suspension was sonicated (Qsonica) for 30 s (2 s on, 3 s off) at 40% amplitudes. Cell debris was removed via centrifugation at 20,000 g for 20 min at 4 °C. The supernatant was transferred to a clean tube, and the protein concentration was determined using Pierce™ 660nm Protein Assay Kit. 2 mg/mL (500 μg) of HEK293T cell lysates were precipitated with methanol:chloroform and resuspended in 235.7 μL of 1x PBS pH 7.4, 0.1% Triton. The protein solutions were combined with 3.125 μL of 20 mM $CuSO_4$, 2.5 μL of 50 mM BTTP, 2.5 μL of 10 mM biotin-alkyne (Figure S22)[11], and 6.25 μL of 100 mM sodium L-ascorbate. The click reactions were agitated on a shaker for 60 min at room temperature then quenched with 25 μL of 50 mM EDTA, pH 8.0. Protein samples were precipitated with methanol:chloroform and resuspended in 2x Laemmli buffer containing 60 mM DTT.

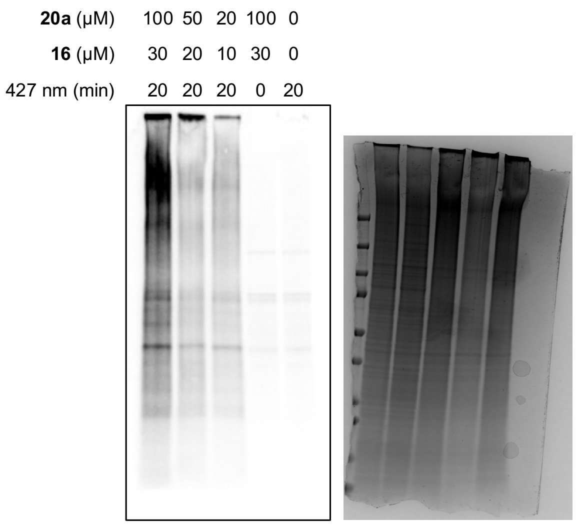


**Figure S27:** Live HEK 293T cells photolabeling with different **20a** and **16** concentrations and fixed illumination time. Western blot (left), Coomassie stain (right)

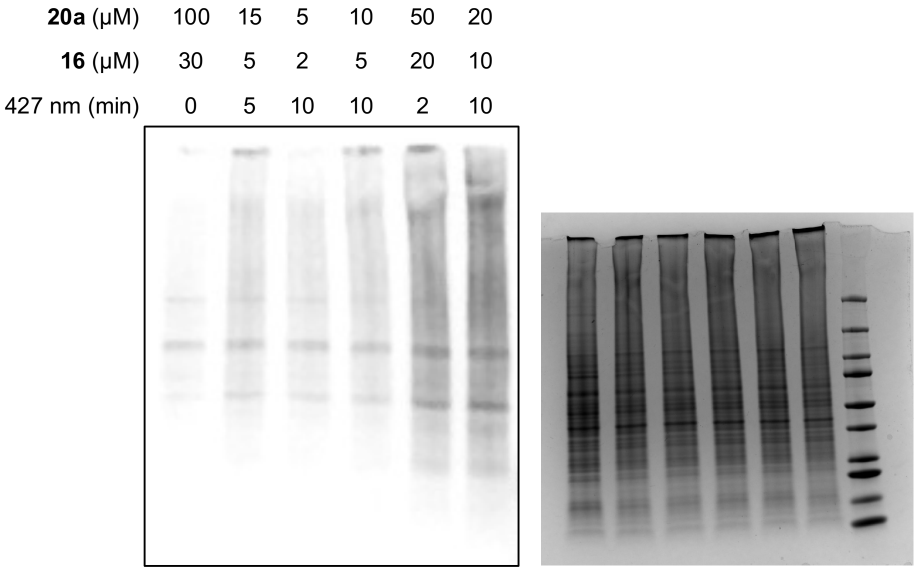


**Figure S28:** Live HEK 293T cells photolabeling with lower concentrations of **20a** and **16** and shorter illumination time. Western blot (left), Coomassie stain (right)

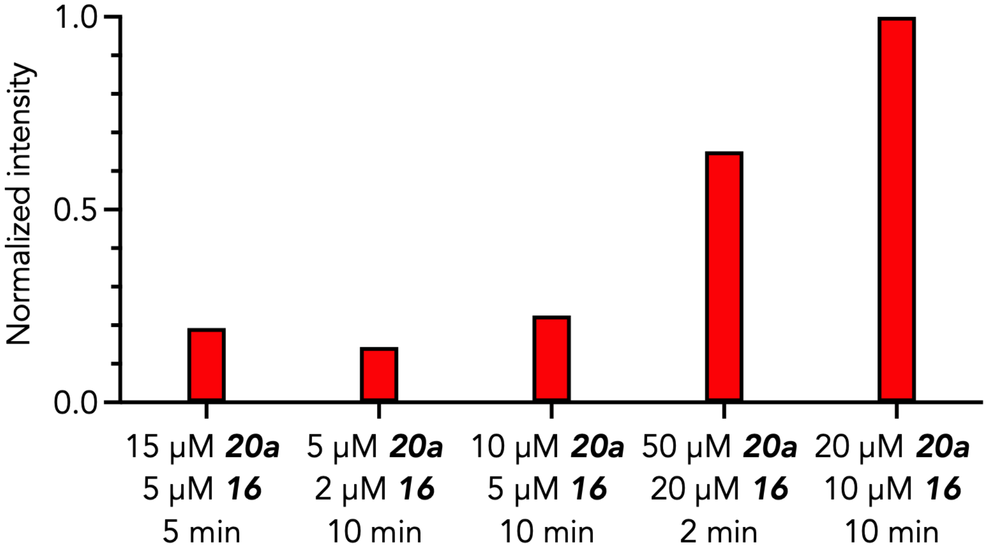


**Figure S29:** Relative band intensity for different labeling outcomes when HEK293T cells were treated with lower concentrations of **20a** and **16** and exposed to shorter illumination time

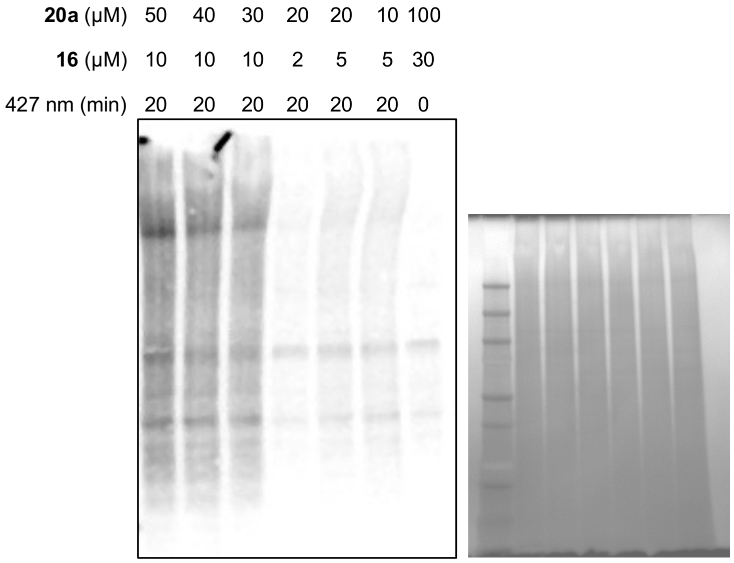


**Figure S30:** Live HEK 293T cells photolabeling with varying **20a**:**16** ratio. Western blot (left), Zinc stain (right)

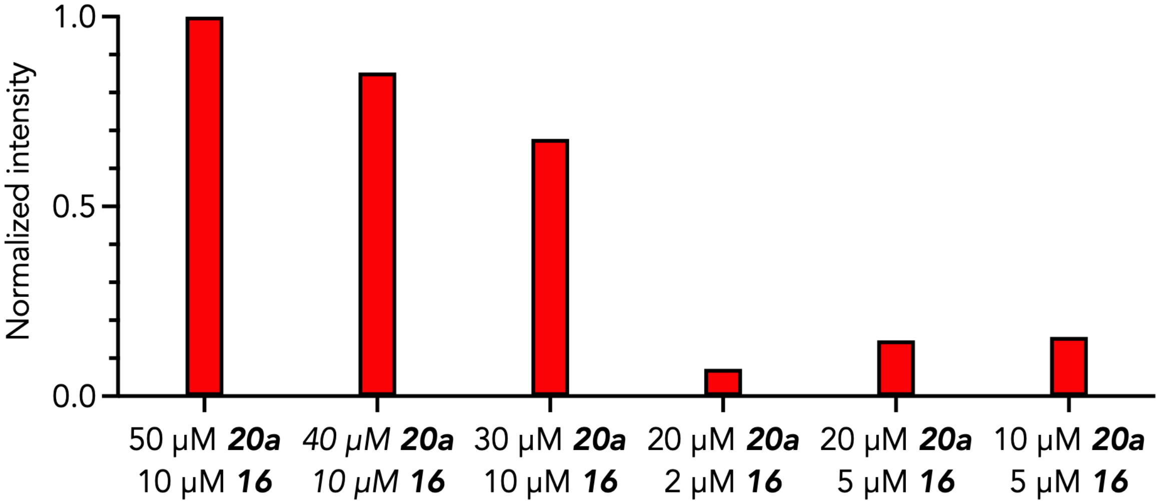


**Figure S31:** Relative band intensity for different labeling outcomes when HEK293T cells were exposed to varying **20a**:**16** ratios

**Lysate level photolabeling with 20a and 16 for mass spectrometry-based proteomics analysis**

**A. In-lysate photolabeling with 20a and 16.** 2 mg/mL (500 µg) of native HEK293T cell lysates were treated with 30 µM **16** and 100 µM **20a**, 2 mM and 10 mM stock solution in DMSO respectively, or DMSO, in 50 mM HEPES pH 7.5, 150 mM NaCl, 1.5 mM $MgCl_2$, 1% NP-40. A triplicate set was prepared for each condition. The reaction mixtures were mixed well in LC-MS vials and incubated at room temperature for 30 min protected from light. All reaction mixtures were illuminated with 427 nm Kessil lamp for 20 min at room temperature. Protein samples were precipitated with methanol:chloroform and resuspended in 235.7 µL of 1x PBS pH 7.4, 0.1% Triton.

**B. Click-chemistry and protein digestion.** The protein solutions were combined with 3.125 µL of 20 mM $CuSO_4$, 2.5 µL of 50 mM BTTP, 2.5 µL of 10 mM DADPS-biotin-alkyne (CCT-1331-1, VectorLab), and 6.25 µL of 100 mM sodium L-ascorbate. The click reactions were agitated on a shaker for 60 min at room temperature, then quenched with 25 µL of 50 mM EDTA, pH 8.0. Protein samples were precipitated with methanol:chloroform, then resuspended in 200 µL of 8 M Urea in 50 mM Tris-HCl, pH 8.0. The protein solution was sonicated for 2 min at 20% amplitude. 2 µL of 1 M DTT was added to the solution, $[DTT]_{final}$ = 10 mM, then incubated at room temperature for 1 h. 8 µL of 656 mM iodoacetamide (IAA) was further added to the mixture, with a final concentration of [IAA] = 25 mM, and the mixture was then incubated for 30 min at room temperature in the dark. Post incubation, 1740 µL of 50 mM Tris-HCl, pH 8.0, and 40 µL of 100 mM $CaCl_2$ (final concentration was 2 mM) were introduced to the denatured, alkylated protein solution. The mixture was vortexed at top speed for 1 min. 10 µL of 1 mg/mL Trypsin/LysC (Thermo Fisher Scientific) in nano-pure water (10 µg, 1:50 w/w protease:protein) was added to the mixture and then incubated at 37 °C for 16 h. After incubation, the solution was heated up to 95 °C for 10 min followed by vortexing for 1 min.

**C. Peptide-level enrichment.** In a 2 x 8 cm PD-10 column (17043501, Cytiva), 200 µL of 50% slurry Neutravidin Ultralink resin (Thermo Fisher Scientific) was added and washed with 1x PBS, pH 7.4 (4 x 2 mL). The digested peptide mixture was introduced to the pre-washed Neutravidin resin and incubated with mild rotation at 4 °C for 20 h. After the incubation, the resin was washed with 0.2% SDS in 1x PBS, pH 7.4 (4 x 2 mL), 1x PBS, pH 7.4 (4 x 2 mL), 2 M NaCl (2 x 2 mL), and nano-pure water (3 x 2 mL). The bound/modified peptides were eluted with 10% formic acid in LC-MS grade water for 30 min at room temperature with mild rotation (3 x 1 mL). The resin was washed once with 2 mL of 0.1% formic acid in LC-MS grade water for 5 min with mild rotation. All eluate fractions were combined, lyophilized, and desalted using Pierce™ Peptide Desalting Spin Column (Thermo Scientific). The desalted peptides were then concentrated using speed-vac then reconstituted in 40 µL of 0.1% formic acid in LC-MS grade water.

**D. Liquid chromatography-tandem mass spectrometry (nLC-MS/MS).** 5 µL of desalted peptides solution was separated on a 45 °C heated PepMap™ RSLC C18 (3 µm, 100 Â, 75 µm x 15 cm) (P/N ES900) facilitated by an online Vanquish-Neo Nano-LC system (Thermo Fisher Scientific) equipped with a trap column (P/N 174502). The liquid chromatography was operated at a flow rate of 300 nL/min using the following mobile phases: A = 0.1% formic acid in $H_2O$ and B = 0.1% formic acid in 80% $CH_3CN/H_2O$. The gradient started at 1 - 5% B over 0.2 min, 5 - 40% B over 120 min, and finally 40 - 90% B for 10 min. All data were acquired using data-dependent acquisition mode on an Orbitrap Exploris 240 with XCalibur (version 4.7) (Thermo Fisher Scientific) coupled to the above nano-LC system. The expected LC peak width was set to 12 s with a default charge state of 2, and EASY-IC™ was used for lock mass correction at the start of each run. The spray voltage was set at static and positive mode with a value of 2000 V. The ion transfer tube temperature was held constant at 280 °C. One full MS1 scan was taken with the following parameters: resolution = 180 K, scan range = 375 - 1500 m/z, RF lens = 70%, AGC target = 300%, maximum injection time = 25 ms, microscan = 1, data type = profile, polarity = positive. The monoisotopic peak determination was set to peptide mode, and the restrictions were relaxed if too few precursors were found. Peaks with charge states from 2 to 8 were selected in each MS1 scan for MS2 with an intensity threshold of 8E3; meanwhile, undetermined charge state peaks were disregarded. For the dynamic exclusion, the ion precursors were placed on the exclusion list for a duration of 30 s after being selected once with a mass tolerance of 10 ppm (low/high), isotope exclusion, and only performing dependent scan on single charge state per precursor. As many MS2 scans were obtained within the 2 s cycle time between MS1 and MS2 using the following parameters: isolation window = 2 ms, isolation offset = off, collision energy type = normalized, HCD collision energy = 30%, resolution = 30 K, scan range mode = define first mass (120 m/z), AGC target = 100, maximum injection time = auto, microscan = 1, data type = centroid.

**E. Label-free quantification proteomics for enriched digested peptides.** LC/MS data were analyzed on FragPipe (version 23.0) and searched with MS Fragger[12] (version 4.3) against the *homo sapiens* proteome database (40906 entries, UniProtKB) containing common contaminants and reversed sequences. Enriched peptides were further quantified with IonQuant[13] (version 1.11.11). The built-in workflow LFQ-MBR from FragPipe was used to identify and quantify the enriched peptides. Trypsin was specified as the proteolytic enzyme with a maximum of 2 missed cleavages. Cysteine alkylation (+57.02146) was set as a static modification. Methionine oxidation (+15.9949) and N-terminal acetylation (+42.0106) were set at dynamic modification. The enriched peptides were also searched for tryptophan, histidine, phenylalanine, tyrosine, and cysteine, which contained a cleaved tag modification (+282.1692). Precursor and fragment mass tolerance were set to 10 and 20 ppm, respectively. The false discovery rate (FDR) was input at 1% for peptide and protein identification. The MBR RT tolerance in the IonQuant was set at 10 min.

Data availability statement: proteomic datasets will be uploaded to a database upon acceptance of the manuscript for publication.

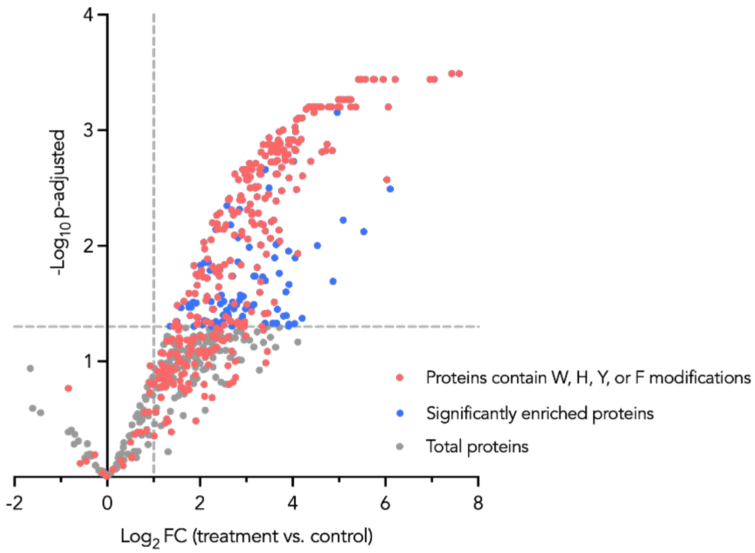


**Figure S32:** Volcano plot for enriched proteins resulted from photolabeling of 500 µg of HEK 293T lysate with **20a** and **16** for 20 min

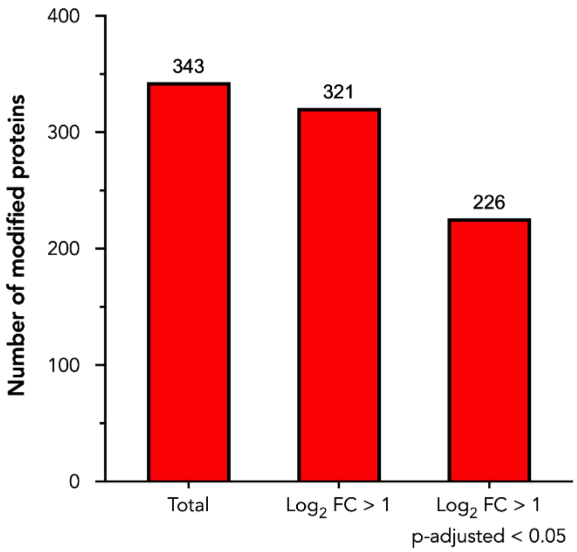


**Figure S33:** Bar graphs for the number of enriched proteins containing W, H, Y, or F modifications.

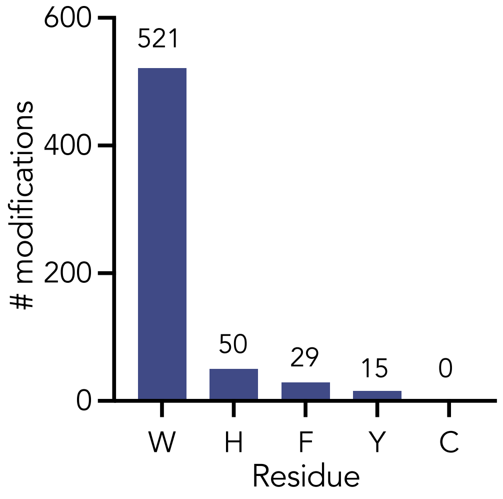


**Figure S34**: All detected modifications by residue.

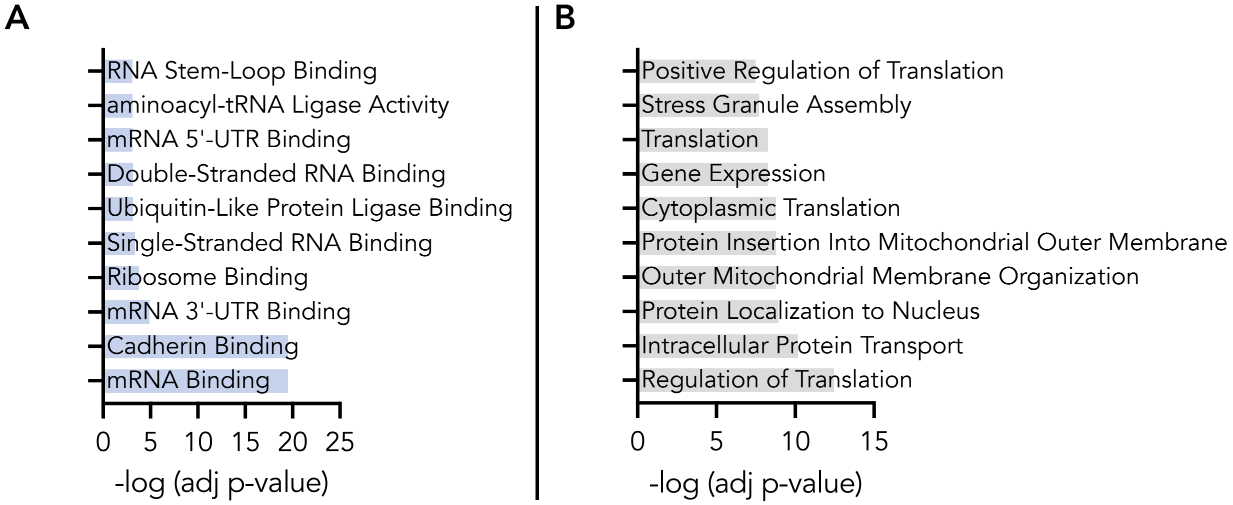


**Figure S35**: Gene ontology using Enrichr of lysate-enriched proteins for (A) Molecular functions and (B) biological processes.

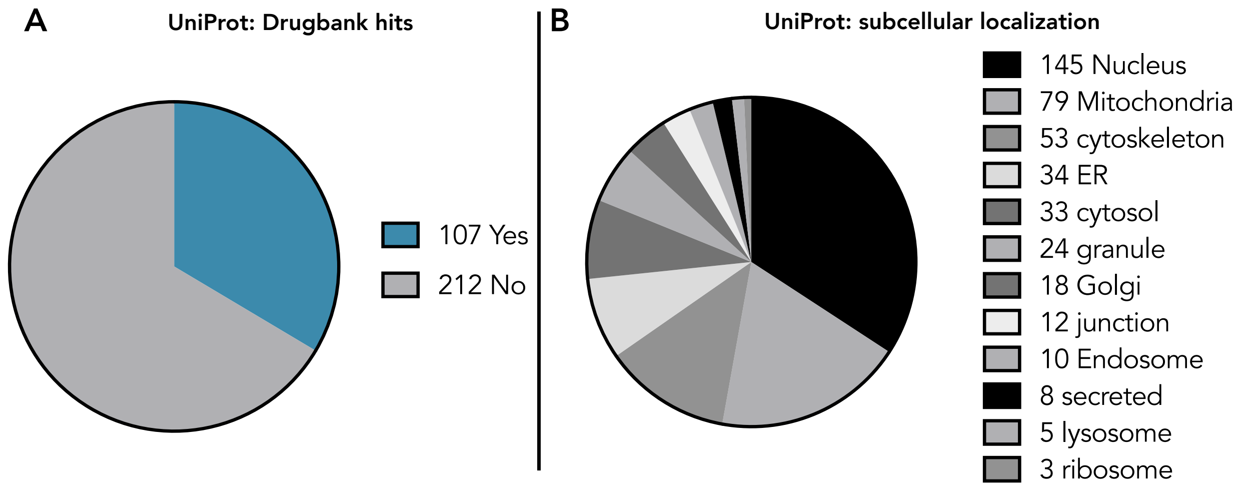


**Figure S36**: UniProt analyses of all lysate-enriched proteins for (A) hits in DrugBank and (B) annotated subcellular localization.

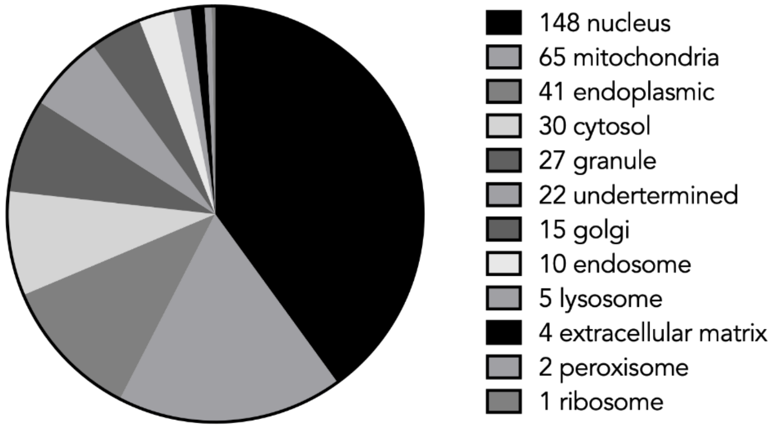


**Figure S37:** Cellular localization for W, H, Y, or F modified proteins with $\log_2$ FC > 1

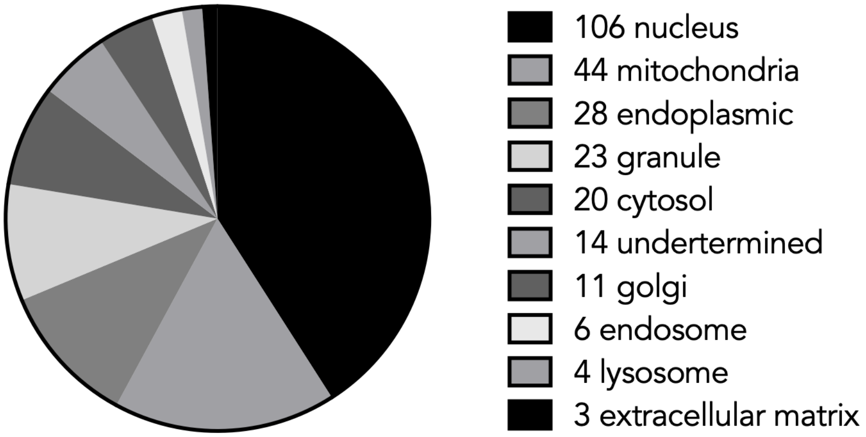


**Figure S38:** Cellular localization for W, H, Y, or F modified proteins with $\log_2$ FC > 1 and p-adjusted < 0.05

**Live HEK293T cells photolabeling with 20a and 16 for mass spectrometry-based proteomics analysis**

**A. Live HEK293T cells photolabeling with 20a and 16.** $4.0\times10^6$ HEK293T cells (viability = 98%) were seeded in T25 flasks for 20 h at 37 °C under 5% $CO_2$ in complete growth DMEM. 2 identical culture flasks were seeded for each condition. A treating solution containing 100 μM **20a** and 30 μM **16** was prepared in Opti-MEM (stock concentrations were 10 mM and 2 mM in DMSO, respectively). A different treating solution only contained DMSO, also prepared in Opti-MEM. Cells were washed with 1x PBS (2 x 500 μL), then treated with 2.7 mL of the treating solution for 1 h at 37 °C. The treated cells from both identical T25 flasks were illuminated simultaneously with 427 nm Kessil lamp for 0 or 20 min at 4 °C. Cells from both T25 flasks were washed once with 1x PBS, then harvested with ice-cold 1x PBS and combined into an Eppendorf tube. Cell pellets were resuspended in 50 mM HEPES, pH 7.5, 150 mM NaCl, 1.5 mM $MgCl_2$, 1% NP-40, and 1x Halt Protease Inhibitor. The cell suspension was sonicated for 30 s (2 s on, 3 s off) at 40% amplitudes. Cell debris was removed via centrifugation at 20,000 g for 20 min at 4 °C. The supernatant was transferred to a clean tube, and the protein concentration was determined using Pierce™ 660nm Protein Assay Kit. Experiments were performed in biological triplicate.

**B. Click-chemistry and protein digestion.** 2 mg/mL (1 mg) of HEK293T cell lysates were precipitated with methanol:chloroform and resuspended in 471.25 μL of 1x PBS pH 7.4, 0.1% Triton. The protein solutions were combined with 6.25 μL of 20 mM $CuSO_4$, 5 μL of 50 mM BTTP, 5 μL of 10 mM DADPS-biotin-alkyne (CCT-1331-1, VectorLab), and 12.5 μL of 100 mM sodium L-ascorbate. The click reactions were agitated on a shaker for 60 min at room temperature then quenched with 50 μL of 50 mM EDTA, pH 8.0. Protein samples were precipitated with methanol:chloroform then resuspended in 400 μL of 8 M Urea in 50 mM Tris-HCl, pH 8.0. The protein solution was sonicated for 2 min at 20% amplitude. 4 μL of 1 M DTT was added to the solution, $[DTT]_{final}$ = 10 mM, then incubated at room temperature for 1 h. 16 μL of 656 mM iodoacetamide (IAA) was further added to the mixture, with a final concentration of [IAA] = 25 mM, and the mixture was then incubated for 30 min at room temperature in the dark. Post incubation, 2716 μL of 50 mM Tris-HCl, pH 8.0, and 64 μL of 100 mM $CaCl_2$ (final concentration was 2 mM) were introduced to the denatured, alkylated protein solution. The mixture was vortexed at top speed for 1 min. 20 μL of 1 mg/mL Trypsin/LysC in nano-pure water (10 μg, 1:50 w/w protease:protein) was added to the mixture and then incubated at 37 °C for 16 h. After incubation, the solution was heated up to 95 °C for 10 min followed by vortexing for 1 min.

**C. Peptide-level enrichment.** In a 2 x 8 cm PD-10 column (17043501, Cytiva), 400 µL of 50% slurry Neutravidin Ultralink resin was added and washed with 1x PBS, pH 7.4 (4 x 4 mL). The digested peptide mixture was introduced to the pre-washed Neutravidin resin and incubated with mild rotation at 4 °C for 20 h. After the incubation, the resin was washed with 0.2% SDS in 1x PBS, pH 7.4 (4 x 4 mL), 1x PBS, pH 7.4 (4 x 4 mL), 2 M NaCl (2 x 4 mL), and nano-pure water (3 x 4 mL). The bound/modified peptides were eluted with 1% formic acid in LC-MS grade water for 30 min at room temperature with mild rotation (3 x 2 mL), followed by elution with 10% formic acid in LC-MS grade water for 30 min at room temperature with mild rotation (3 x 2 mL). The resin was washed once with 2 mL of 1% formic acid in LC-MS grade water for 5 min with mild rotation. All eluate fractions were combined, lyophilized, and desalted using Pierce™ Peptide Desalting Spin Column. The desalted peptides were then concentrated using speed-vac then reconstituted in 60 µL of 0.1% formic acid in LC-MS grade water.

**D. Liquid chromatography-tandem mass spectrometry (nLC-MS/MS).** 5 µL of desalted peptides solution were separated on a 45 °C heated PepMap™ RSLC C18 (3 µm, 100 Â, 75 µm x 15 cm) (P/N ES900) facilitated by an online Vanquish-Neo Nano-LC system (Thermo Fisher Scientific) equipped with a trap column (P/N 174502). The liquid chromatography was operated at a flow rate of 300 nL/min using the following mobile phases: A = 0.1% formic acid in $H_2O$, and B = 0.1% formic acid in 80% $CH_3CN/H_2O$. The gradient started at 1 - 5% B over 0.2 min, 5 - 40% B over 120 min, and finally 40 - 90% B for 10 min. All data were acquired using data-dependent acquisition mode on an Orbitrap Exploris 240 with XCalibur (version 4.7) (Thermo Fisher Scientific) coupled to the above nano-LC system. The expected LC peak width was set to 12 s with a default charge state of 2, and EASY-IC™ was used for lock mass correction at the start of each run. The spray voltage was set at static and positive mode with a value of 2000 V. The ion transfer tube temperature was held constant at 280 °C. One full MS1 scan was taken with the following parameters: resolution = 180 K, scan range = 375 - 1500 m/z, RF lens = 70%, AGC target = 300%, maximum injection time = 25 ms, microscan = 1, data type = profiles, polarity = positive. The monoisotopic peak determination was set to peptide mode, and the restrictions were relaxed if too few precursors were found. Peaks with charge states from 2 to 8 were selected in each MS1 scan for MS2 with an intensity threshold of 8E3; meanwhile, undetermined charge state peaks were disregarded. For the dynamic exclusion, the ion precursors were placed on the exclusion list for a duration of 30 s after being selected once with a mass tolerance of 10 ppm (low/high), isotope exclusion, and only performing dependent scan on single charge state per precursor.  As many MS2 scans were obtained within the 2 s cycle time between MS1 and MS2 using the following parameters: isolation window = 2 ms, isolation offset = off, collision energy type = normalized, HCD collision energy = 30%, resolution = 30 K, scan range mode = define first mass (120 m/z), AGC target = 100, maximum injection time = auto, microscan = 1, data type = centroid.

**E. Label-free quantification proteomics for enriched digested peptides.** LC/MS data were analyzed on FragePipe (version 23.0) and searched with MS Fragger[12] (version 4.3) against the *homo sapiens* proteome database (40906 entries, UniProtKB) containing common contaminants and reversed sequences. Enriched peptides were further quantified with IonQuant[13] (version 1.11.11). The built-in workflow LFQ-MBR from FragPipe was used to identify and quantify the enriched peptides. Trypsin was specified as the proteolytic enzyme with a maximum of 2 missed cleavages. Cysteine alkylation (+57.02146) was set as a static modification. Methionine oxidation (+15.9949) and N-terminal acetylation (+42.0106) were set at dynamic modification. The enriched peptides were also searched for tryptophan, histidine, phenylalanine, tyrosine, and cysteine, which contained a cleaved tag modification (+282.1692). Precursor and fragment mass tolerance were set to 10 and 20 ppm, respectively. The false discovery rate (FDR) was input at 1% for peptide and protein identification. The MBR RT tolerance in the IonQuant was set at 10 min.

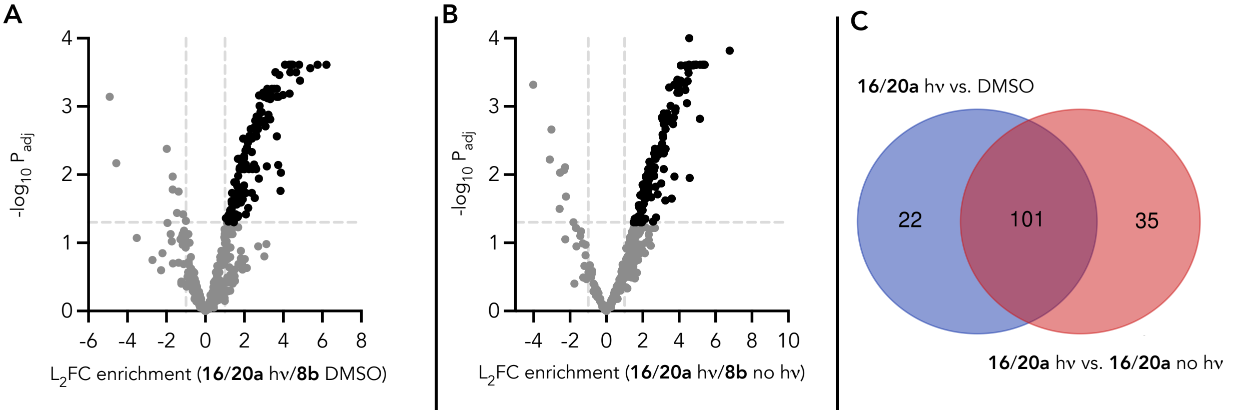


**Figure S39**. Volcano plots of (A) live-cell treatment with **16/20a** with light vs. DMSO and (B) live-cell treatment of **16/20a** with light vs. 16/20a no light. (C) Venn-Diagram depicting enriched proteins from live cell experiments using the following analyses: **16/20a** with light vs. DMSO (blue) and **16/20a** with light vs. **16/20a** no light (blue). The 101 overlapping proteins possess both L2F enrichment ≥1 and -log10($_{padj.}$≥1.3) and were used for further analysis.

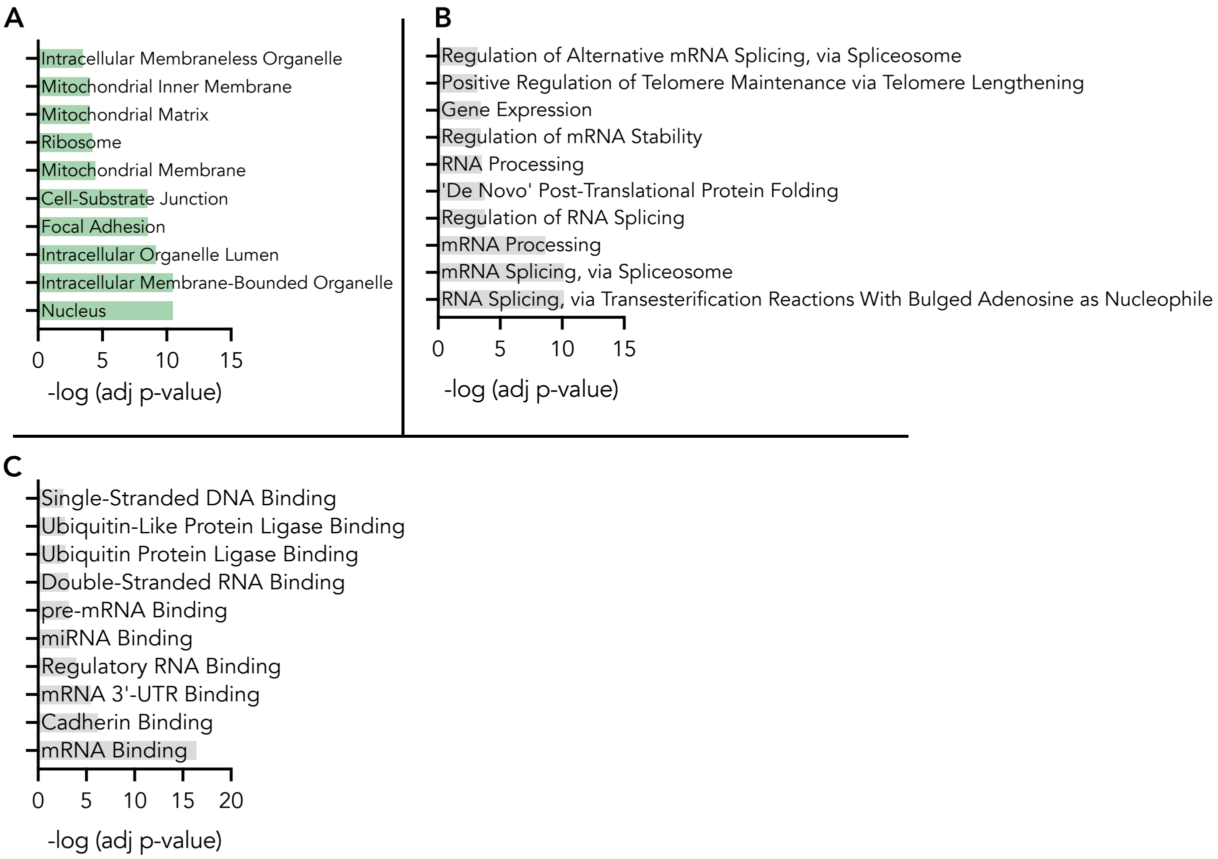

**Figure S40.** GO analysis using Enrichr of live cell-enriched proteins of (A) cellular component (B) biological process and (C) molecular function.

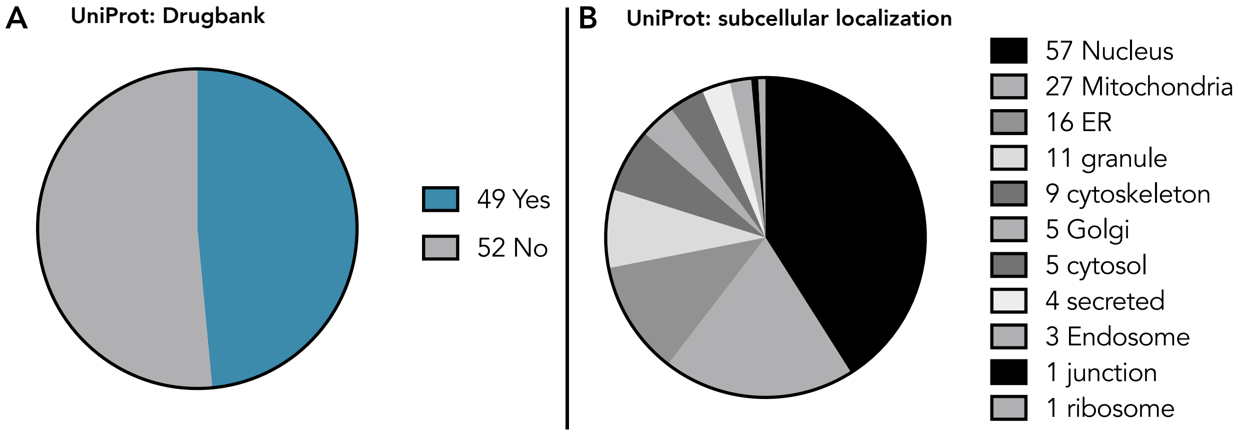


**Figure S41.** UniProt analyses of live cell-enriched proteins for (A) hits in DrugBank and (B) annotated subcellular localization.

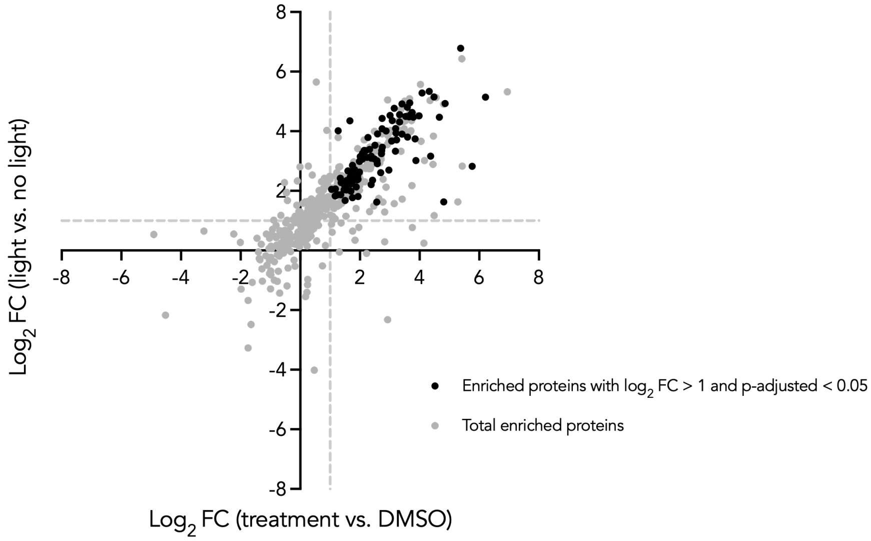


**Figure S42:** Volcano plot for enriched proteins resulted from photolabeling of live HEK 293T cells with **20a** and **16** for 20 min.

#### 8. NMR Spectra

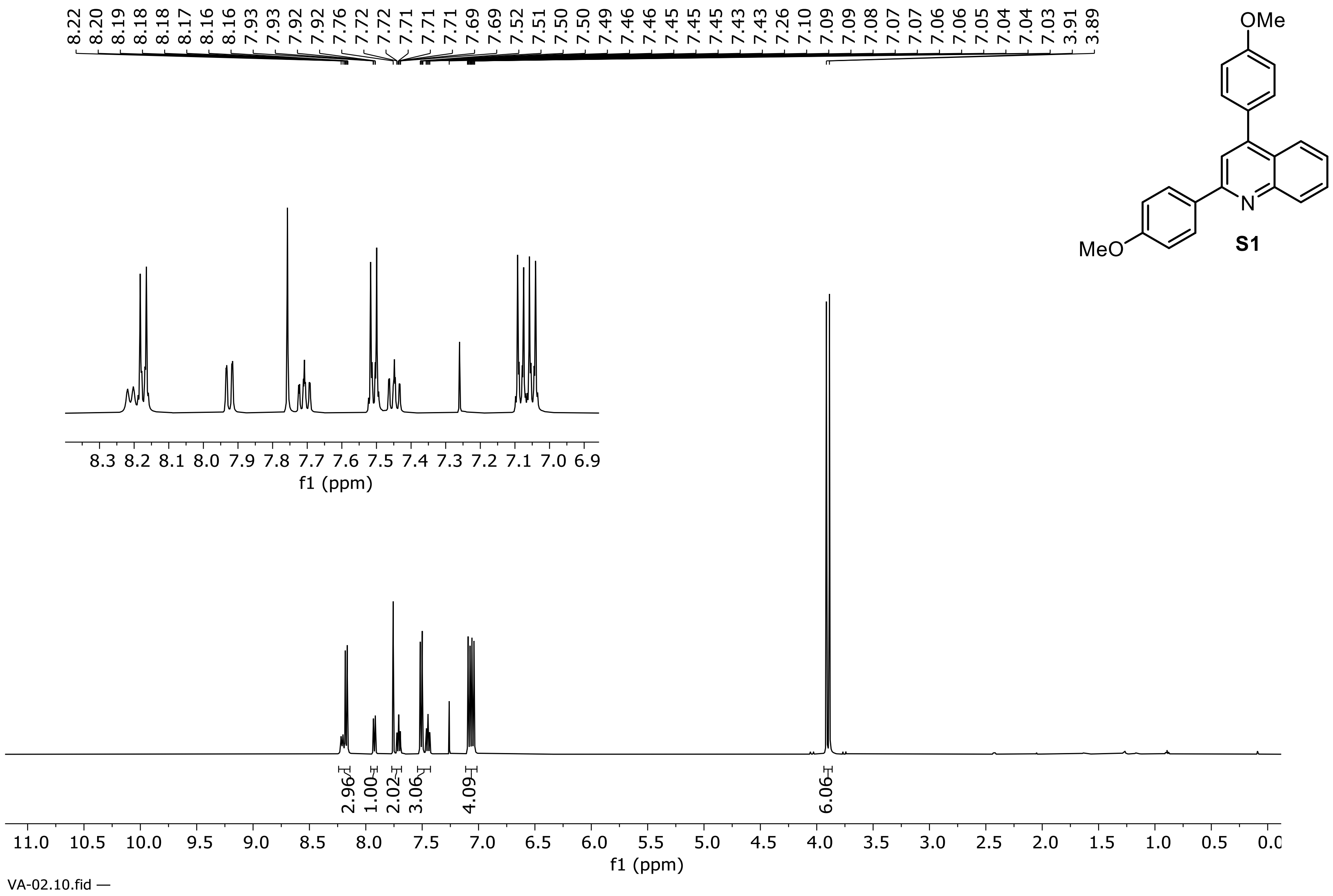


**$^1$H NMR Spectrum of compound S1 (500 MHz, $CDCl_3$)**

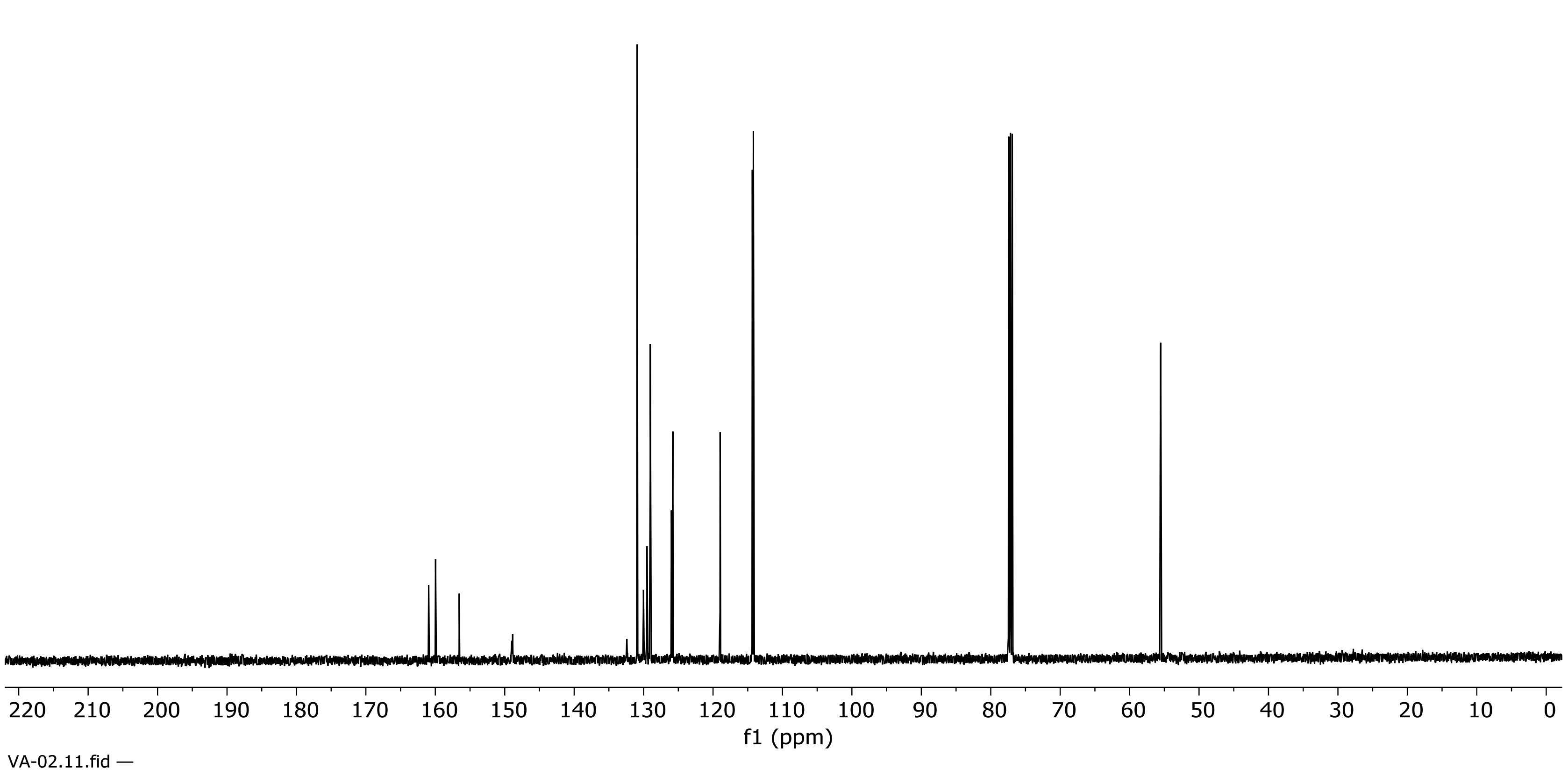


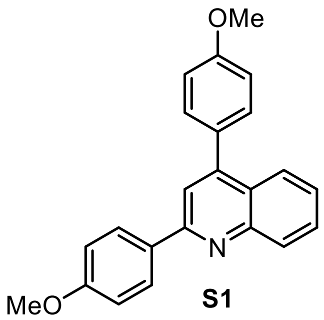


**S1**

**$^{13}C$ NMR Spectrum of compound S1 (126 MHz, $CDCl_3$)**

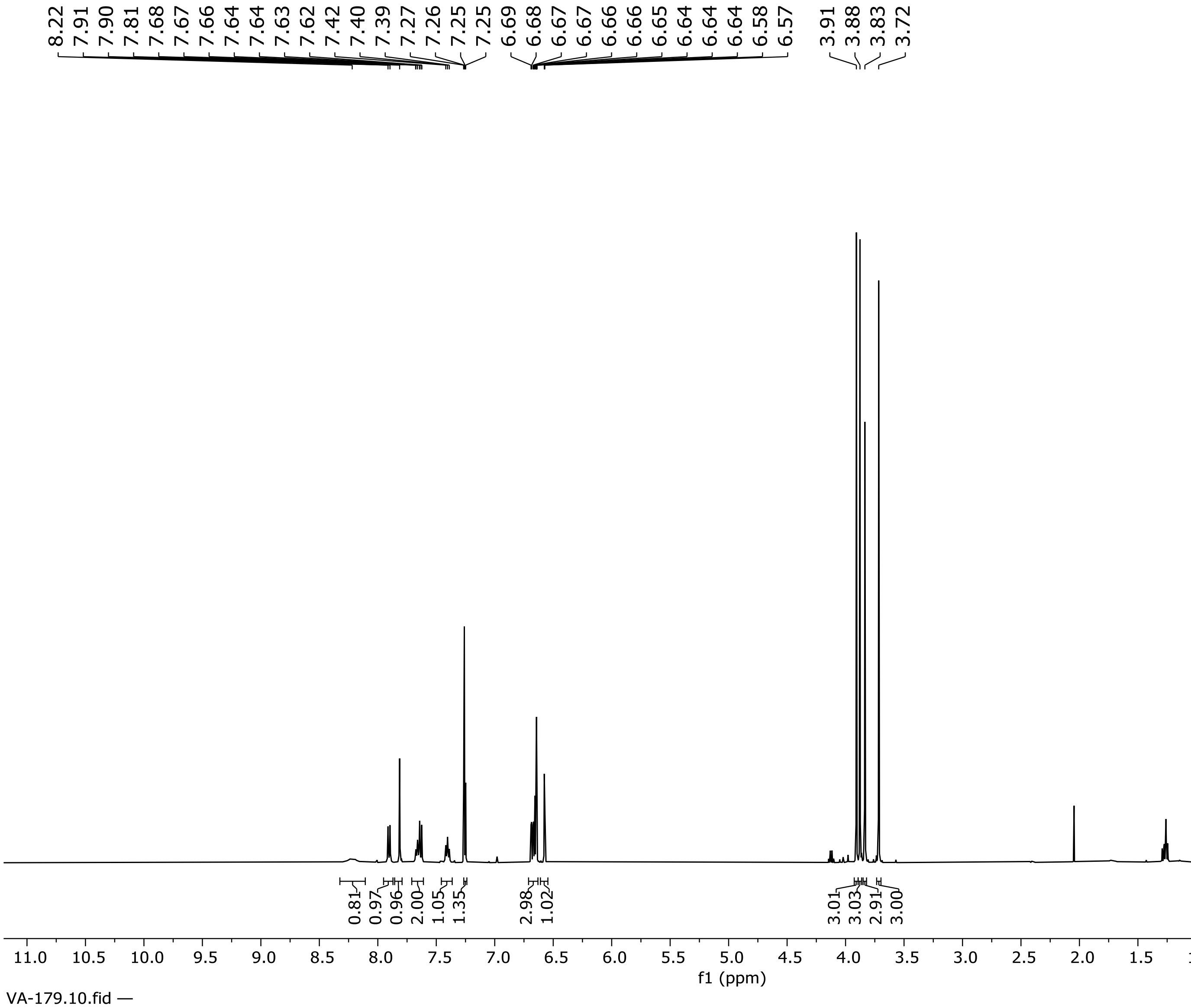


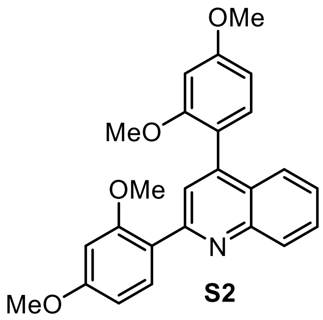


**S2**

**¹H NMR Spectrum of Compound S2 (500 MHz, $CDCl_3$)**

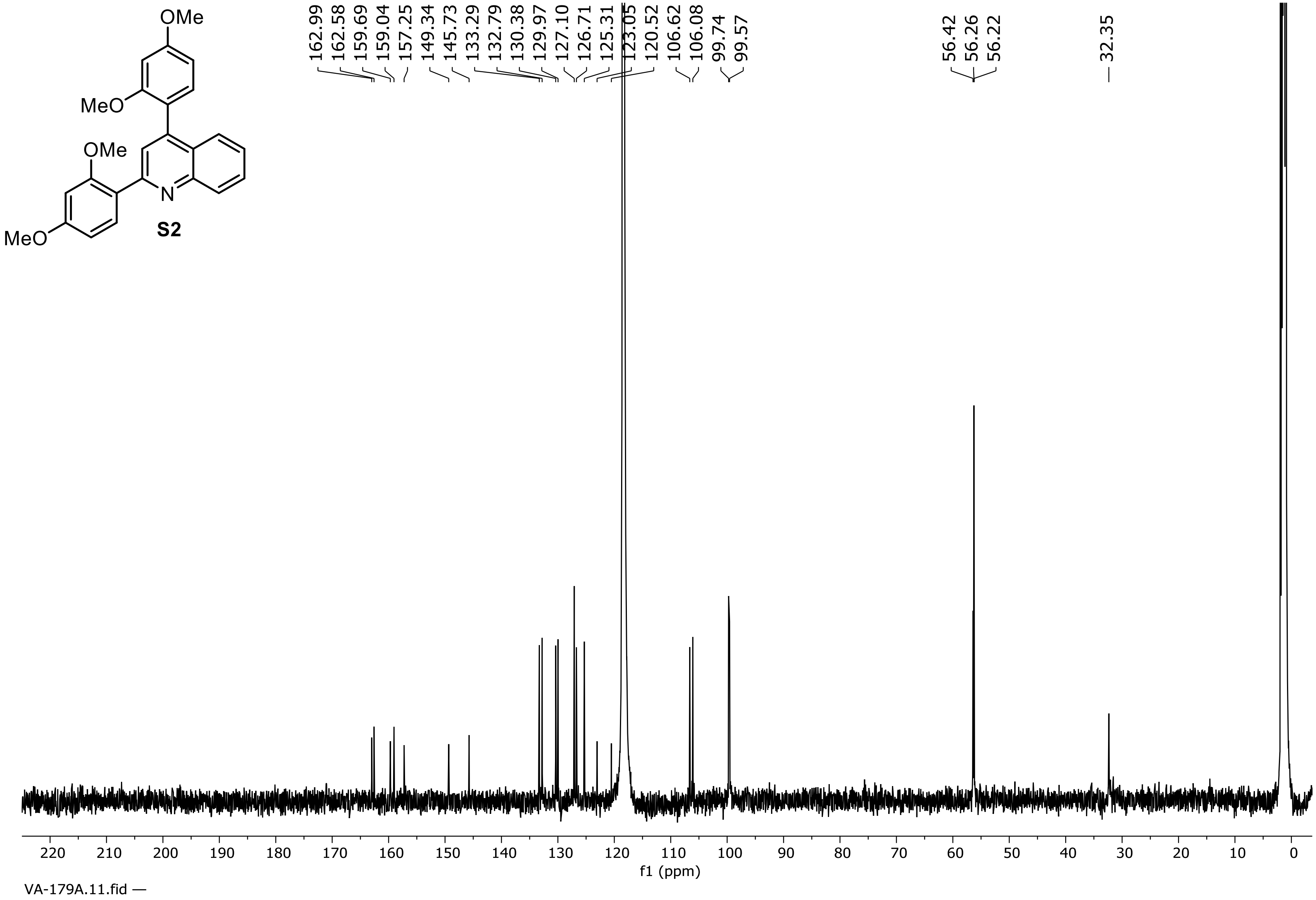


**$^{13}C$ NMR Spectrum of Compound S2 (126 MHz, $CDCl_3$)**

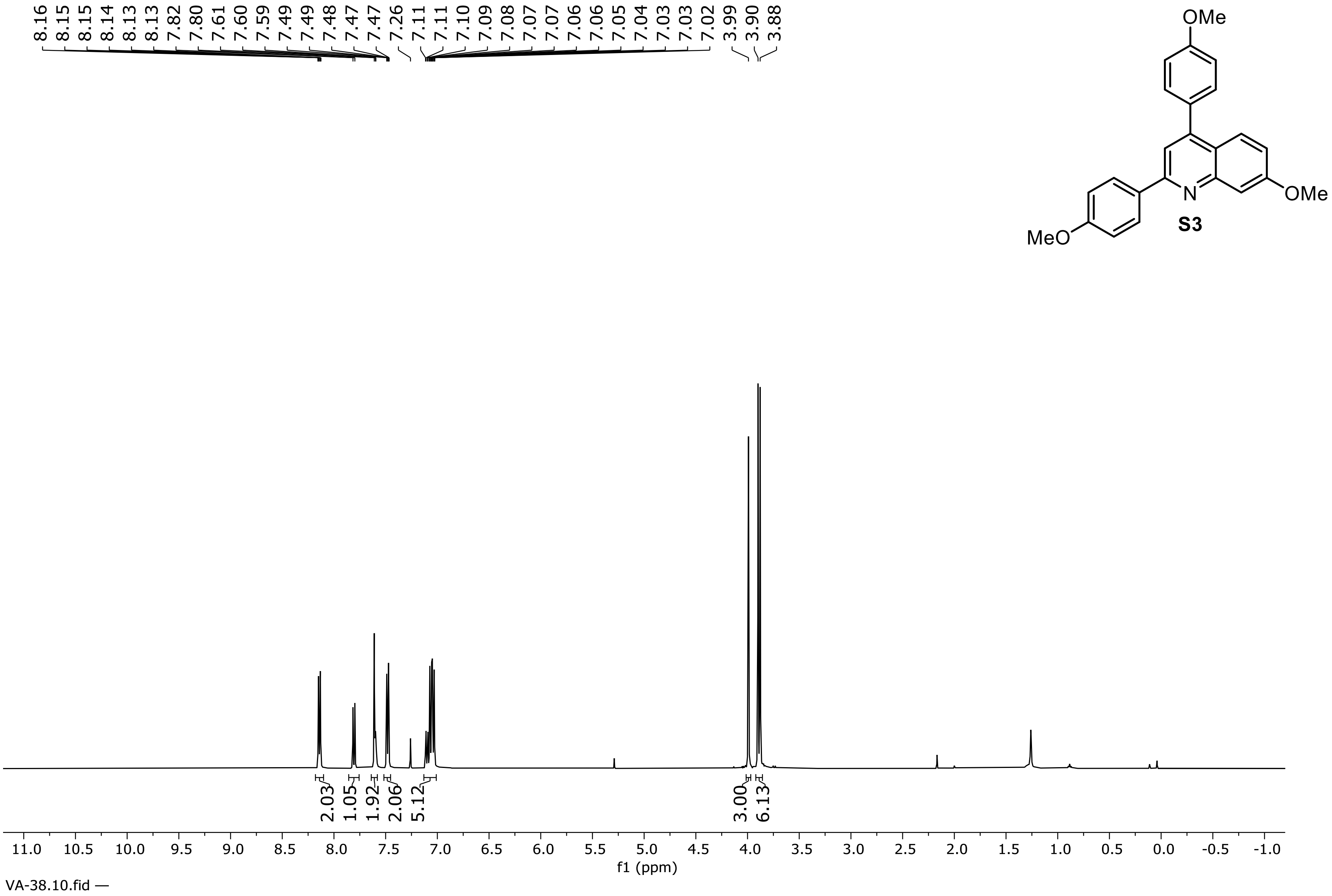


**$^1$H NMR Spectrum of Compound S3 (500 MHz, $CDCl_3$)**

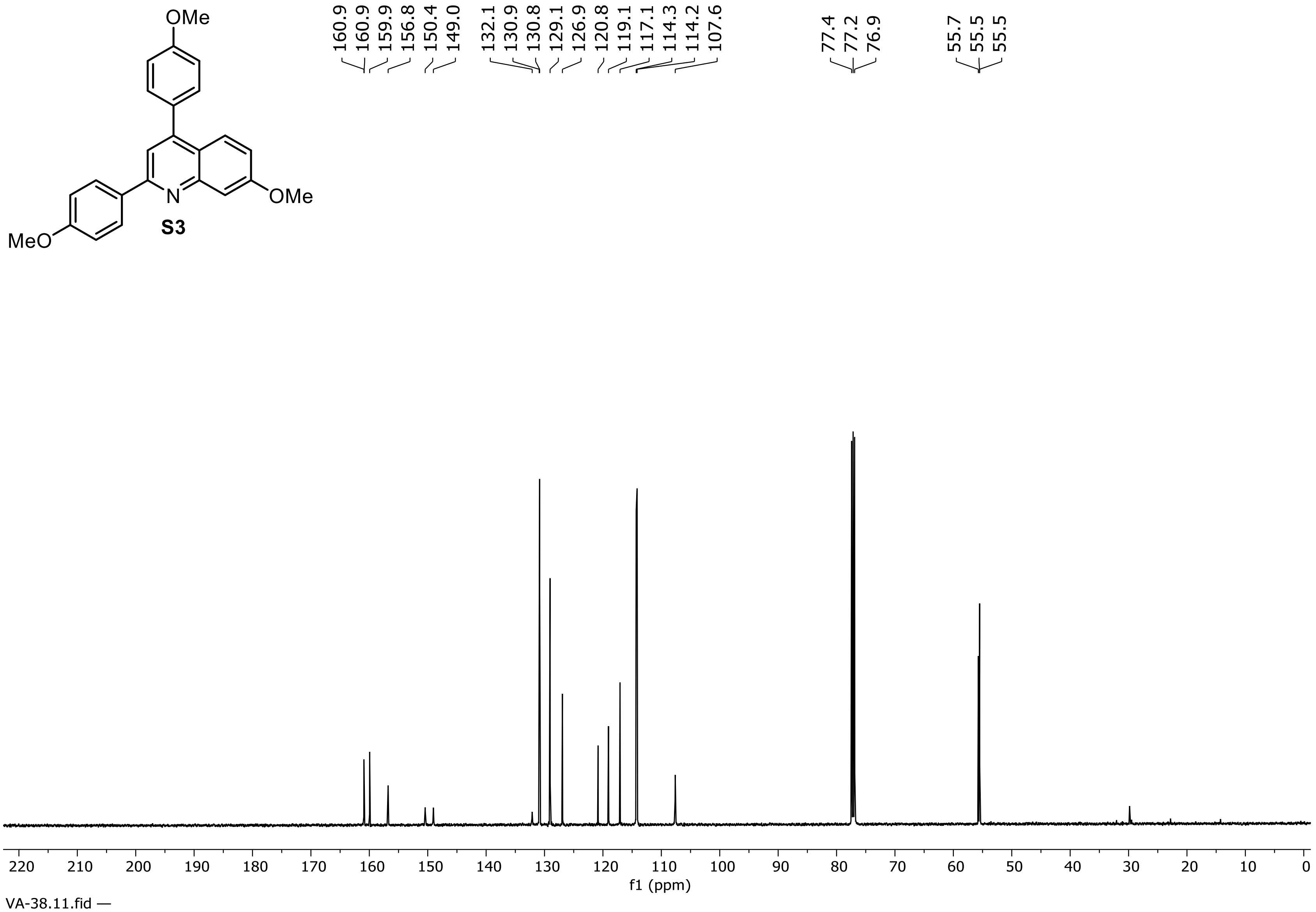


**$^{13}C$ NMR Spectrum of Compound S3 (126 MHz, $CDCl_3$)**

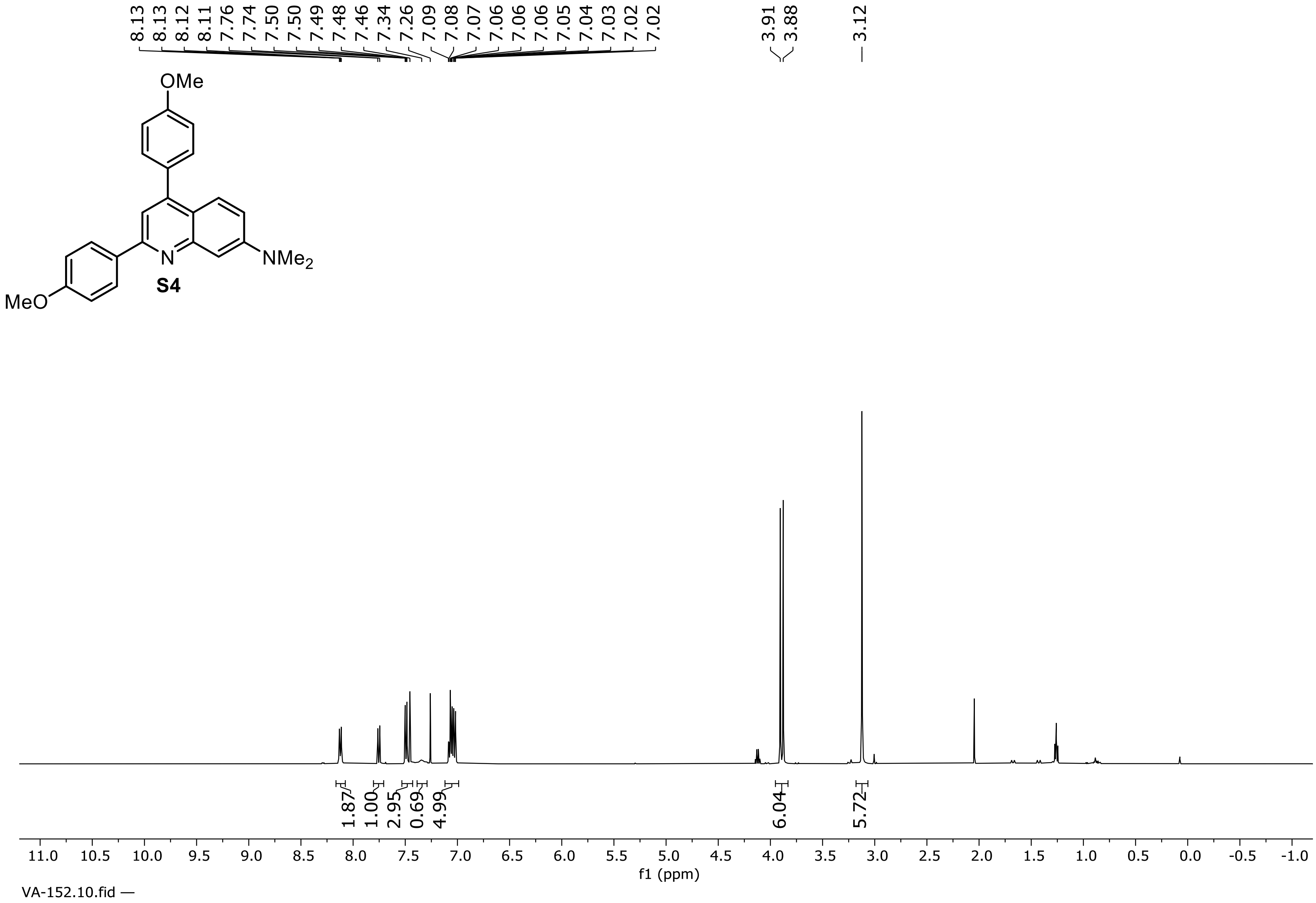


**$^1H$ NMR Spectrum of Compound S4 (500 MHz, $CDCl_3$)**

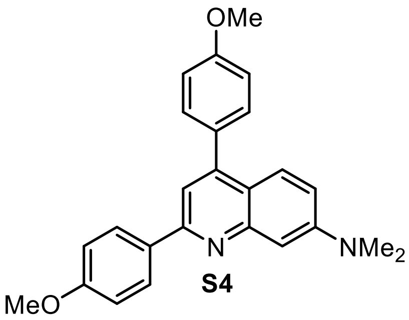

**S4**

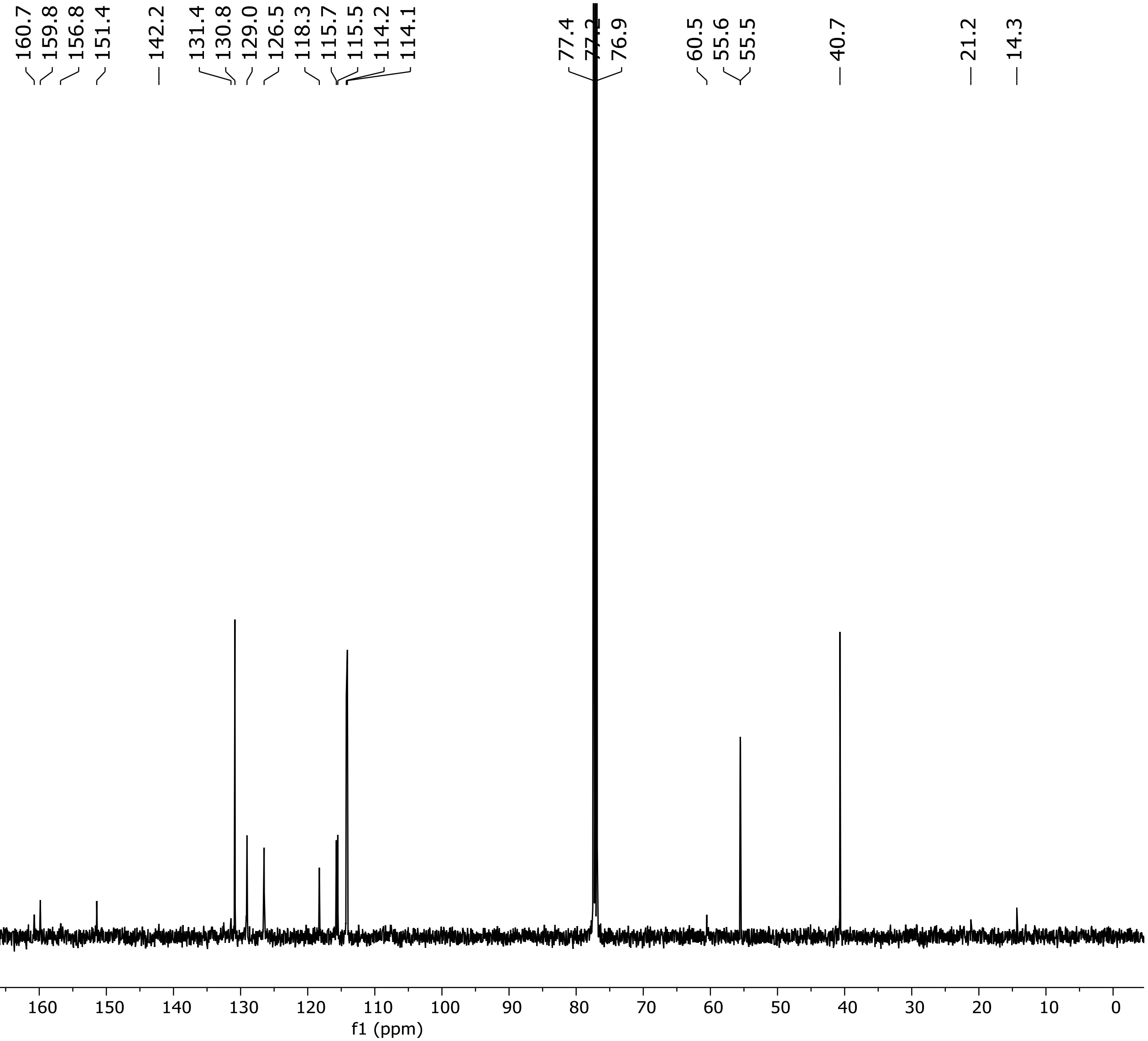


**$^{13}$C NMR Spectrum of Compound S4 (126 MHz, $CD_3CN$)**

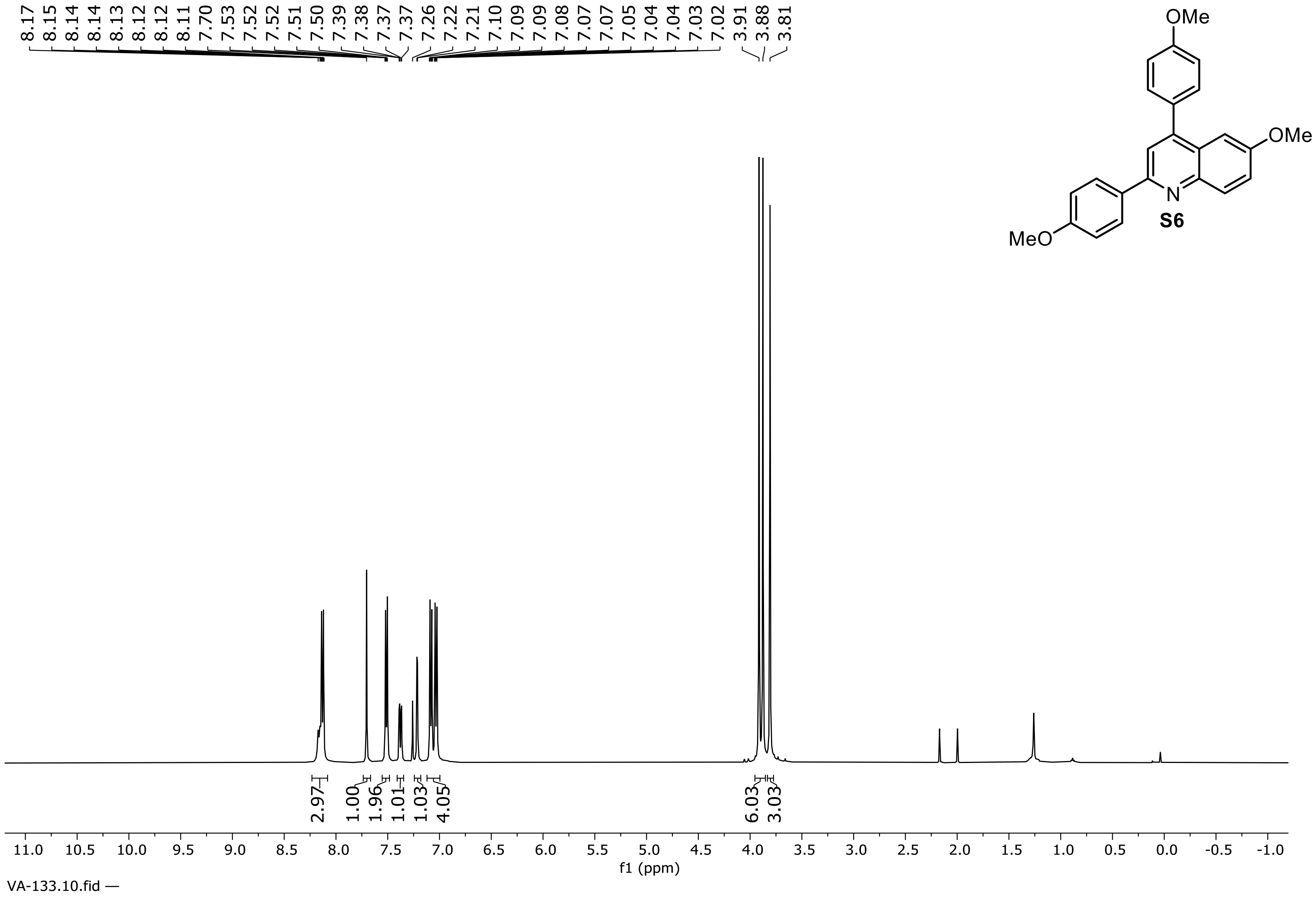


**$^1$H NMR Spectrum of Compound S6 (500 MHz, $CDCl_3$)**

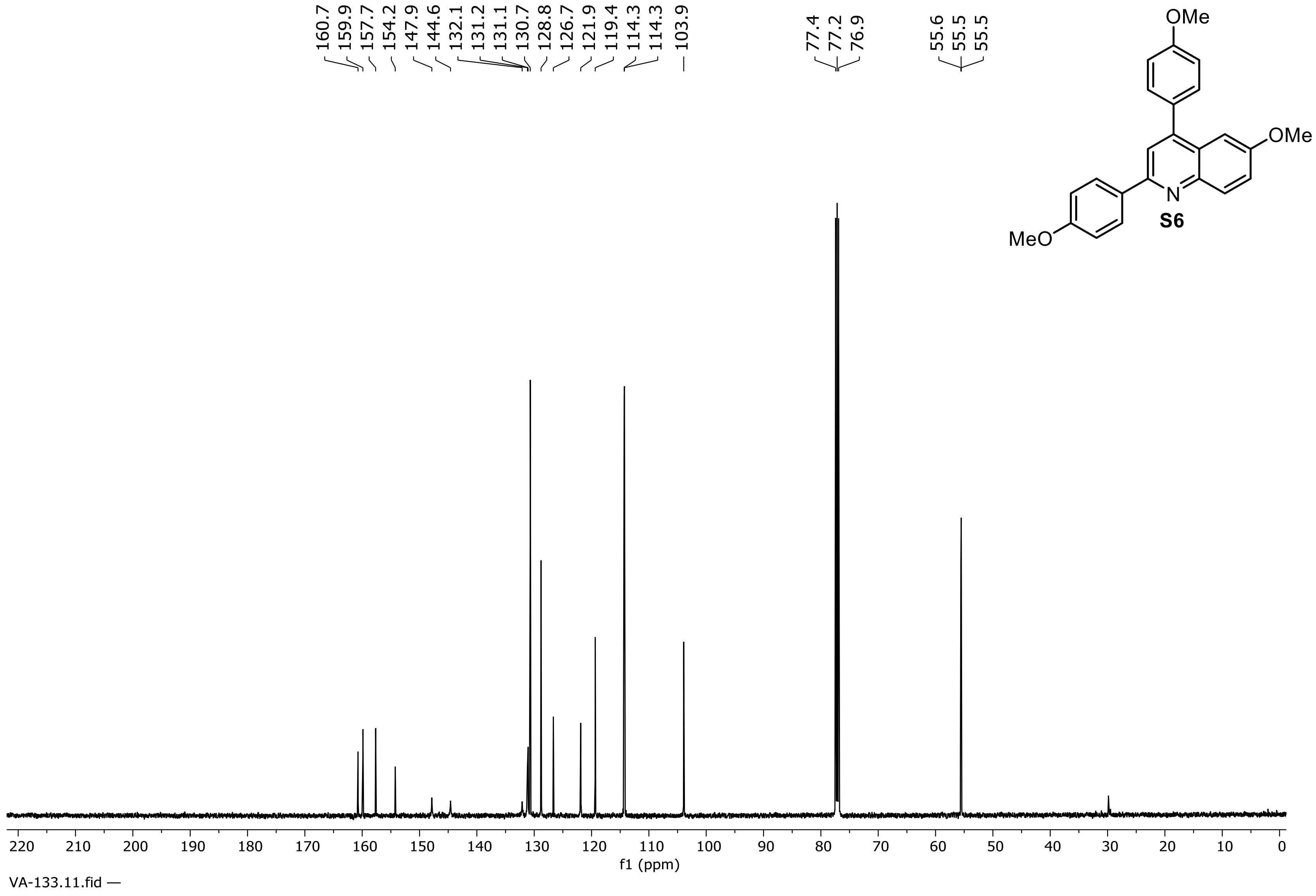


**$^{13}$C NMR Spectrum of Compound S6 (126 MHz, $CDCl_3$)**

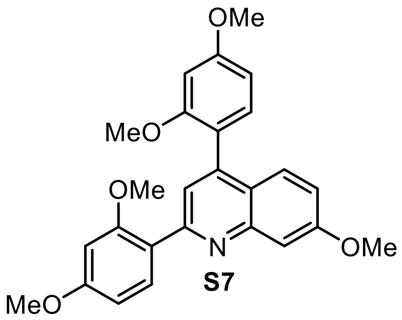

S7

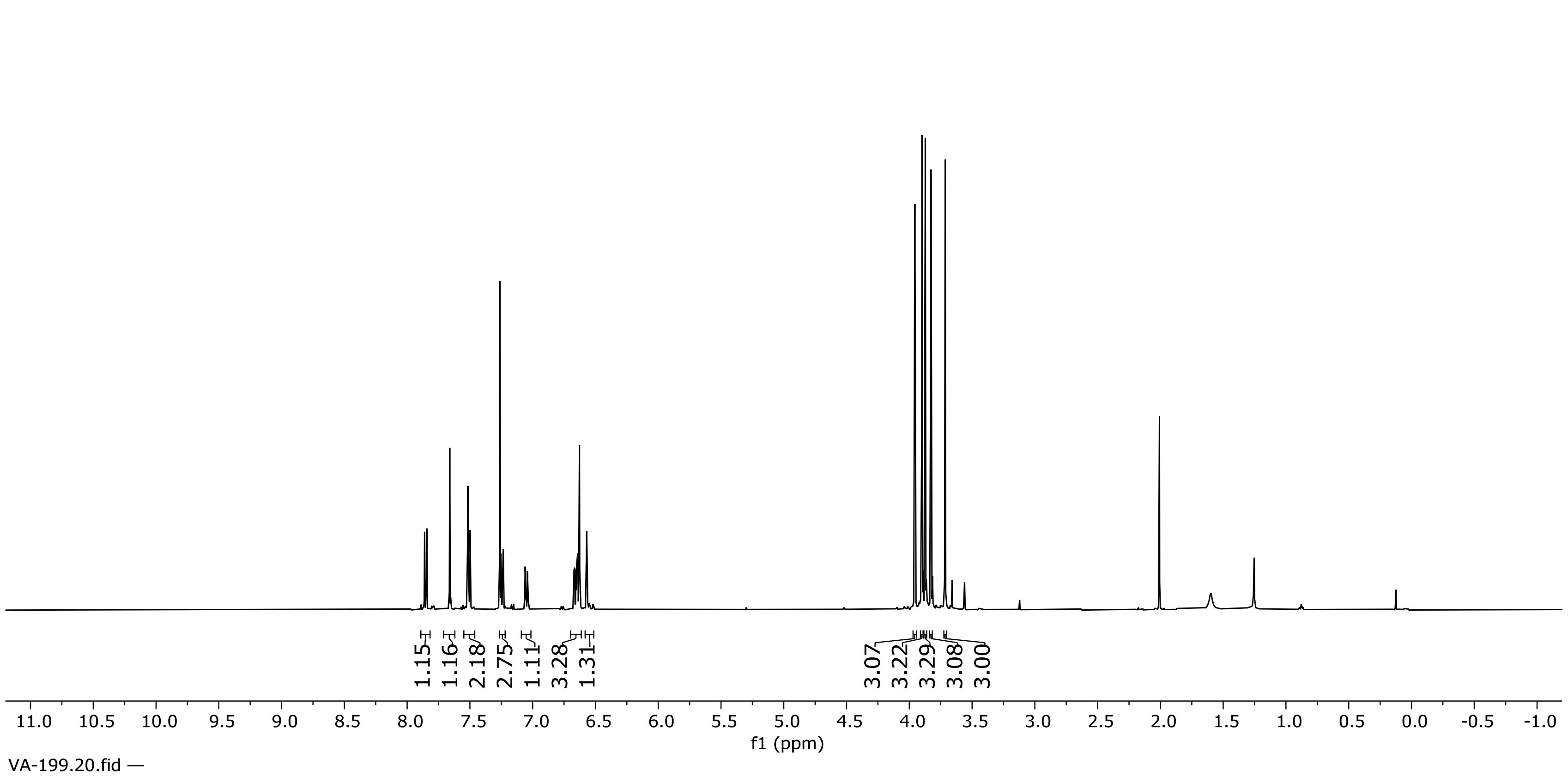


**[1]H NMR Spectrum of Compound S7 (500 MHz, $CDCl_3$)**

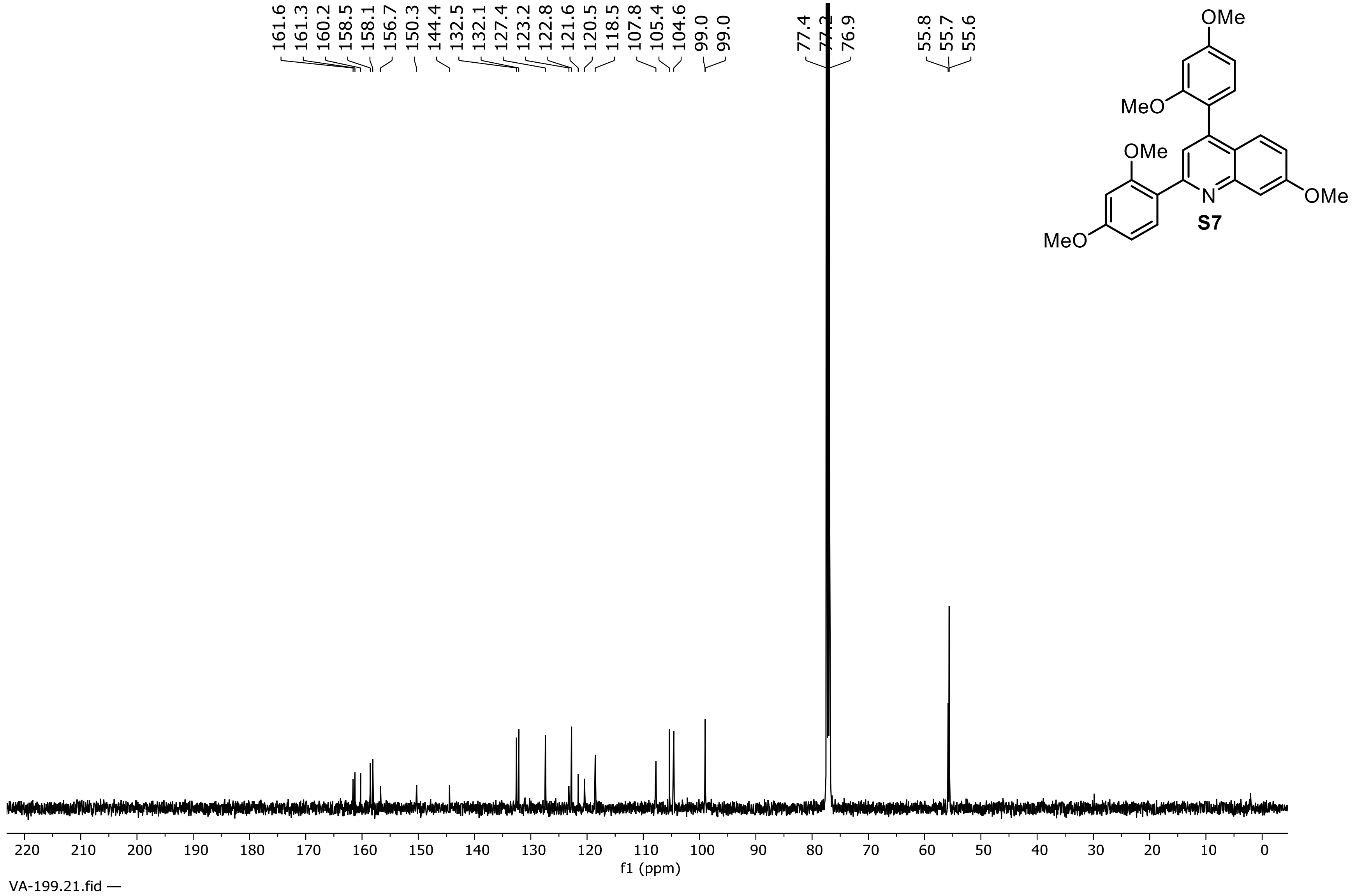


**$^{13}C$ NMR Spectrum of Compound S7 (126 MHz, $CDCl_3$)**

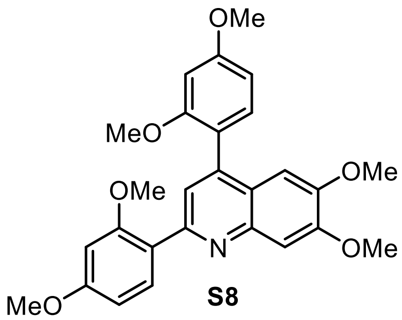


**S8**

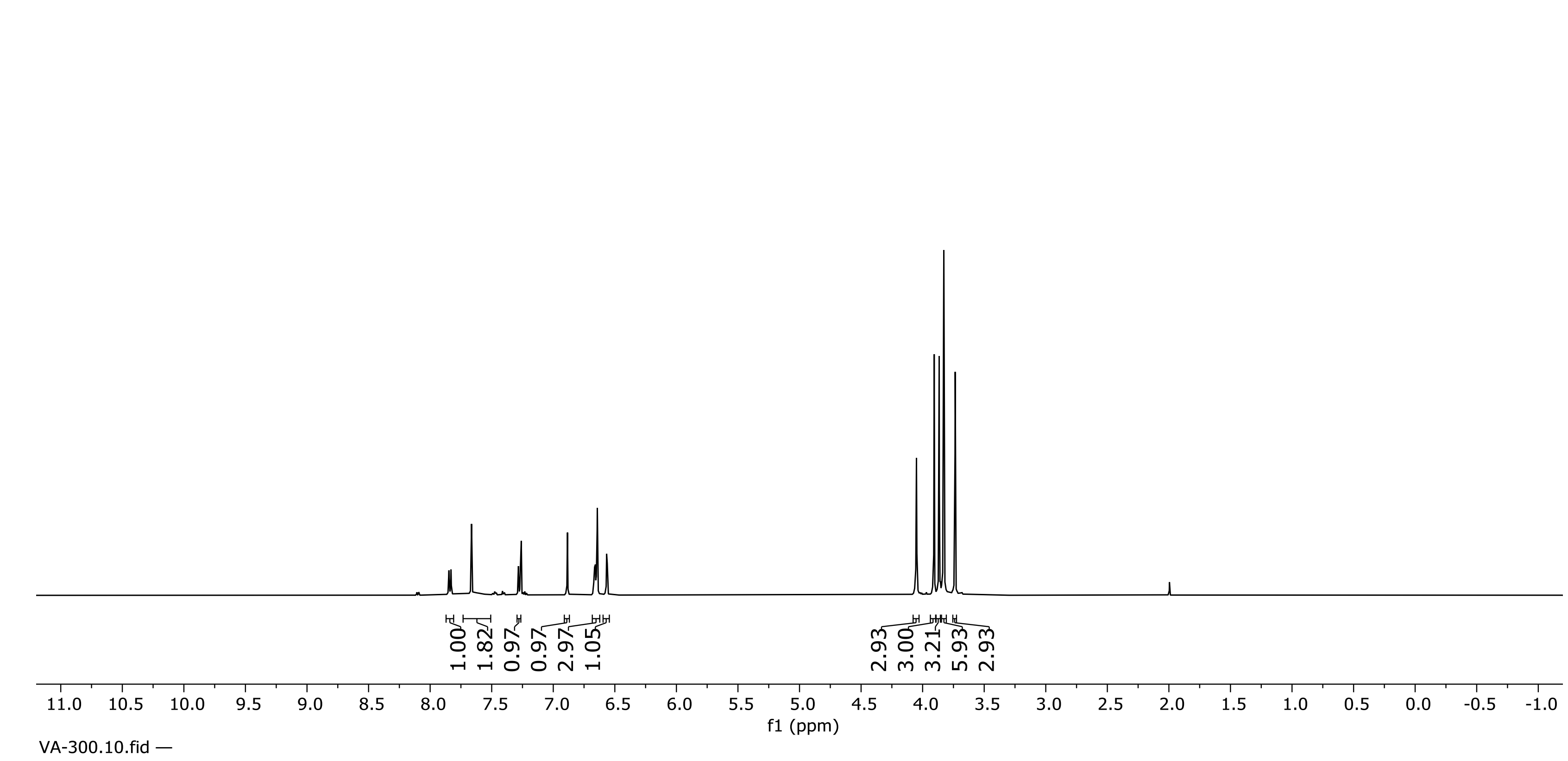


**[1]H NMR Spectrum of Compound S8 (500 MHz, $CDCl_3$)**

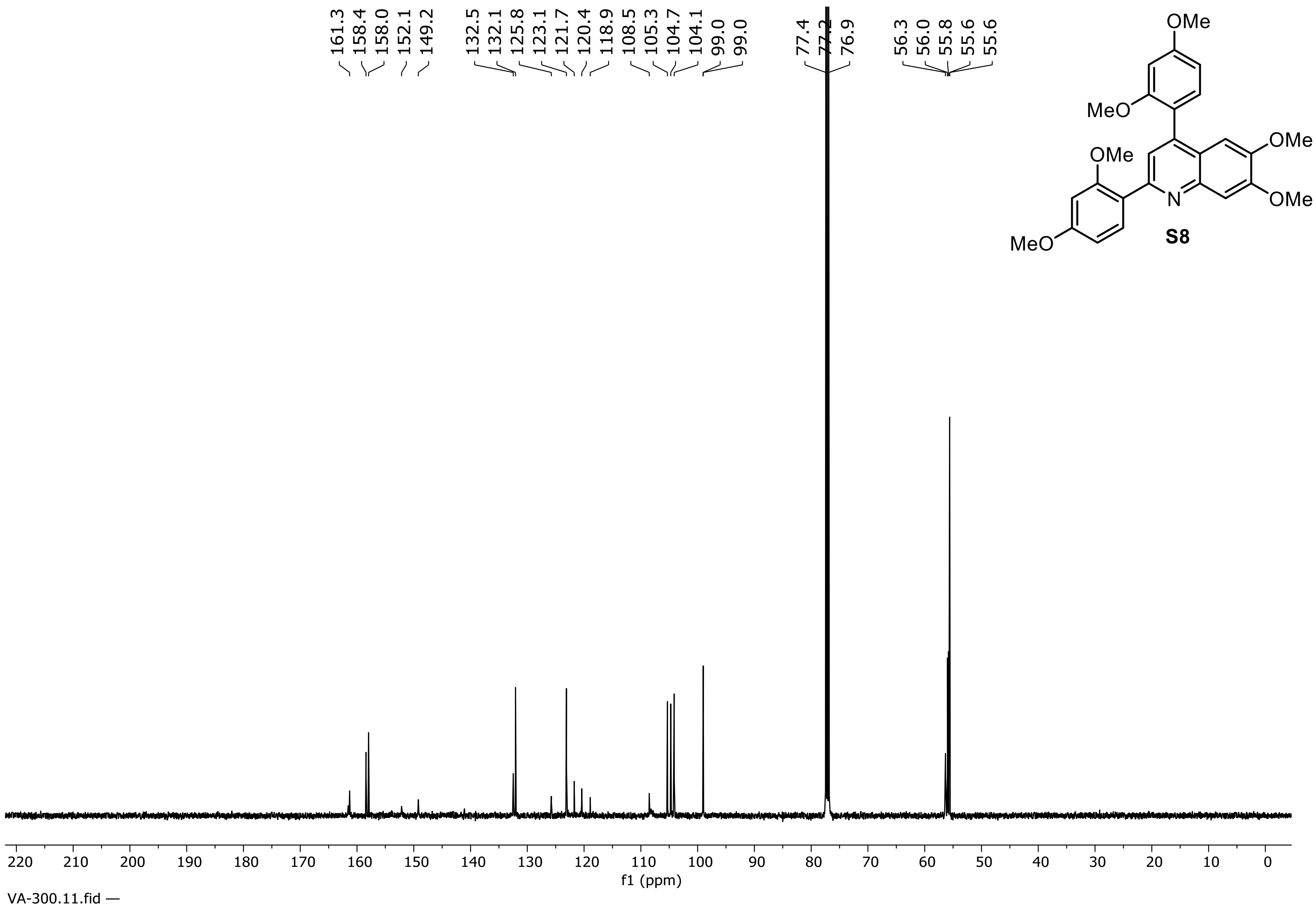


**S8**

**$^{13}$C NMR Spectrum of Compound S8 (126 MHz, $CDCl_3$)**

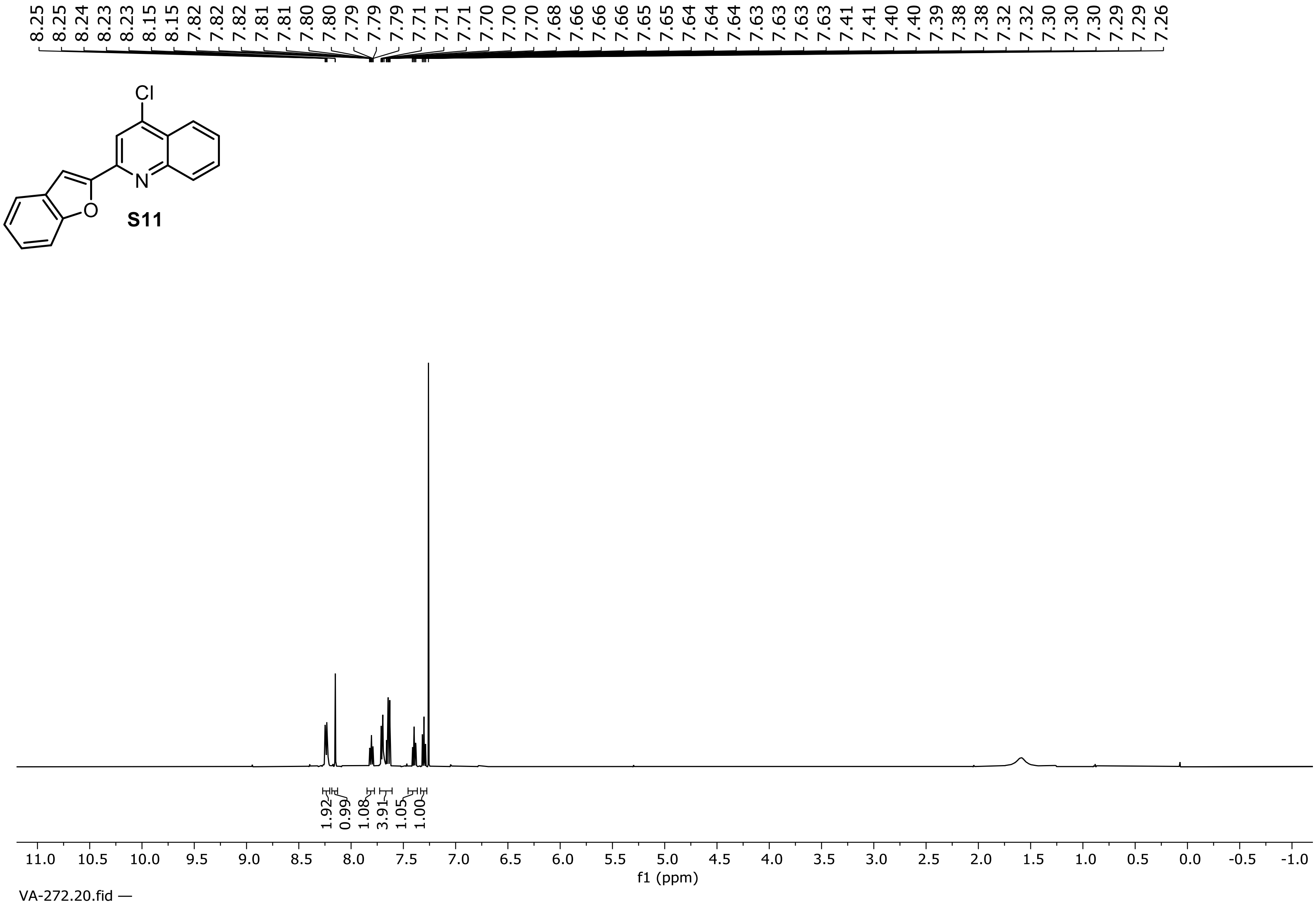


**$^1$H NMR Spectrum of Compound S11 (500 MHz, $CDCl_3$)**

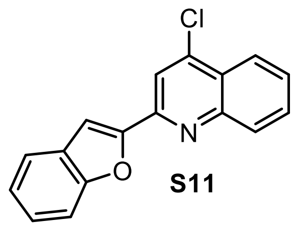


**S11**

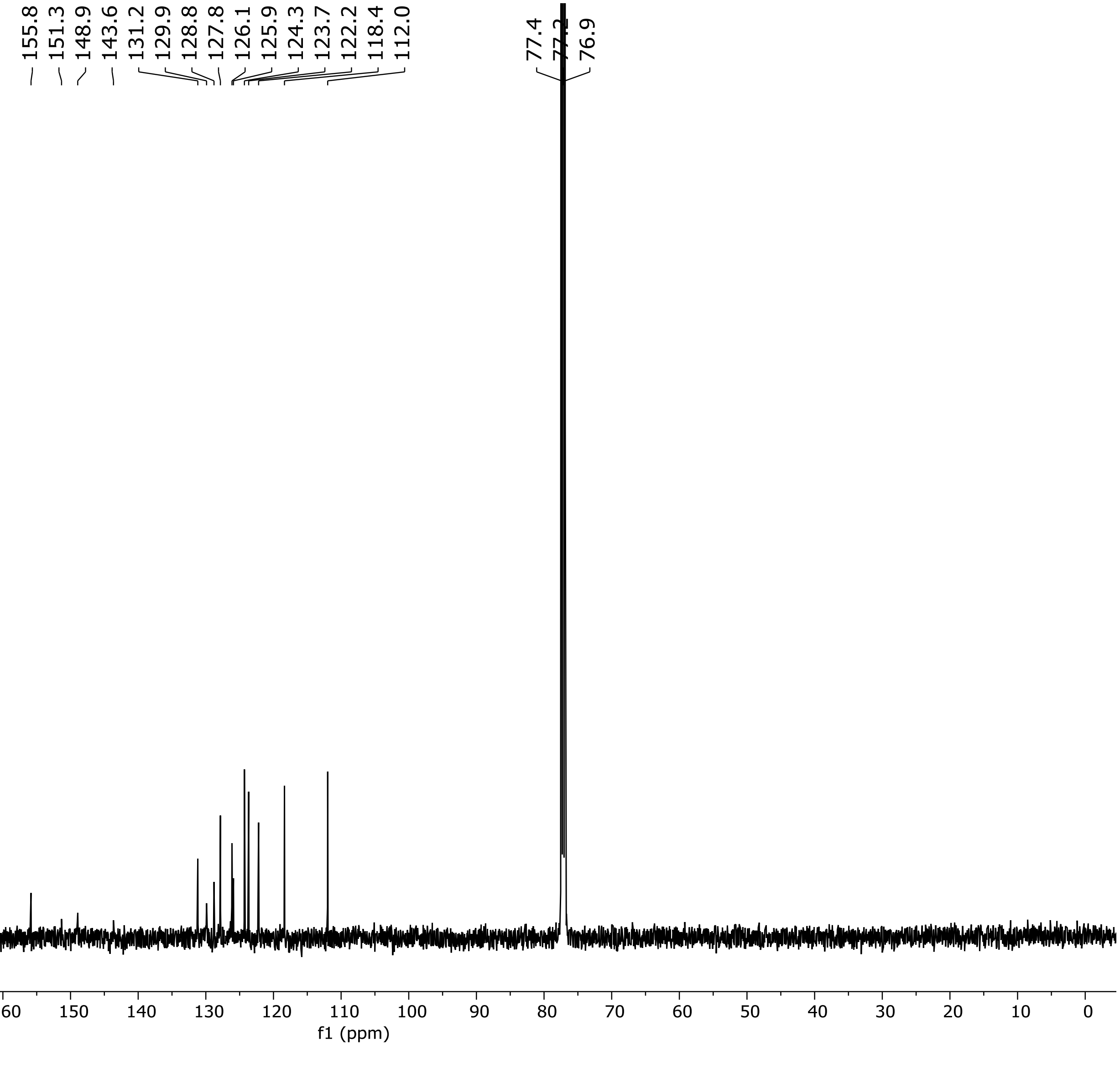


**$^{13}C$ NMR Spectrum of Compound S11 (126 MHz, $CDCl_3$)**

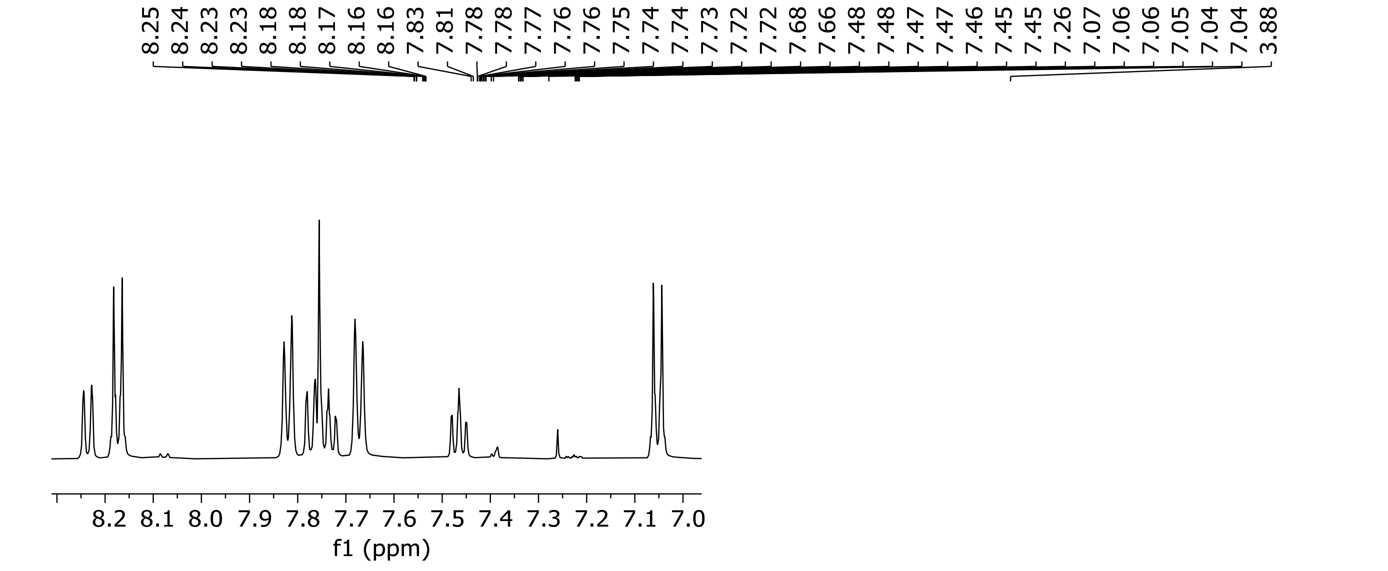


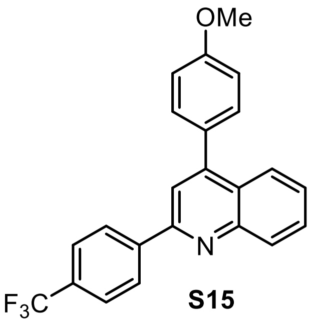


**S15**

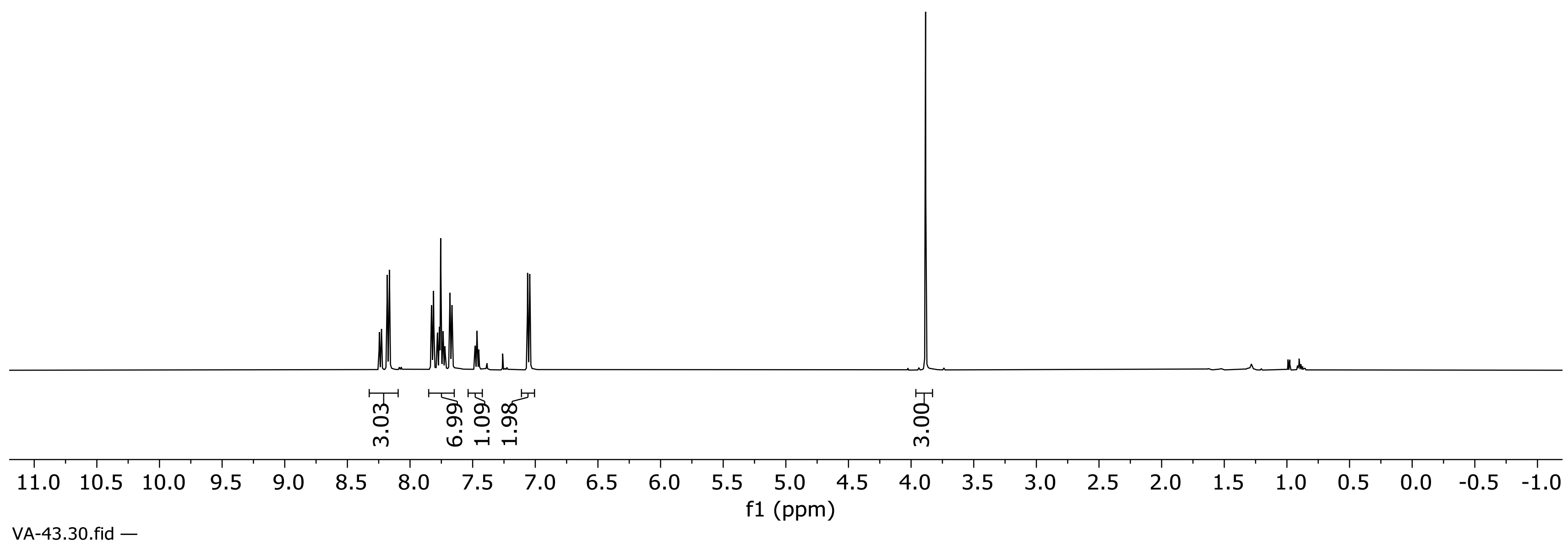


**$^1$H NMR Spectrum of Compound S15 (500 MHz, $CDCl_3$)**

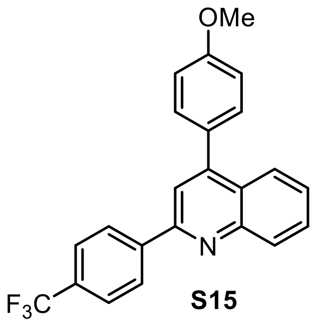


S15

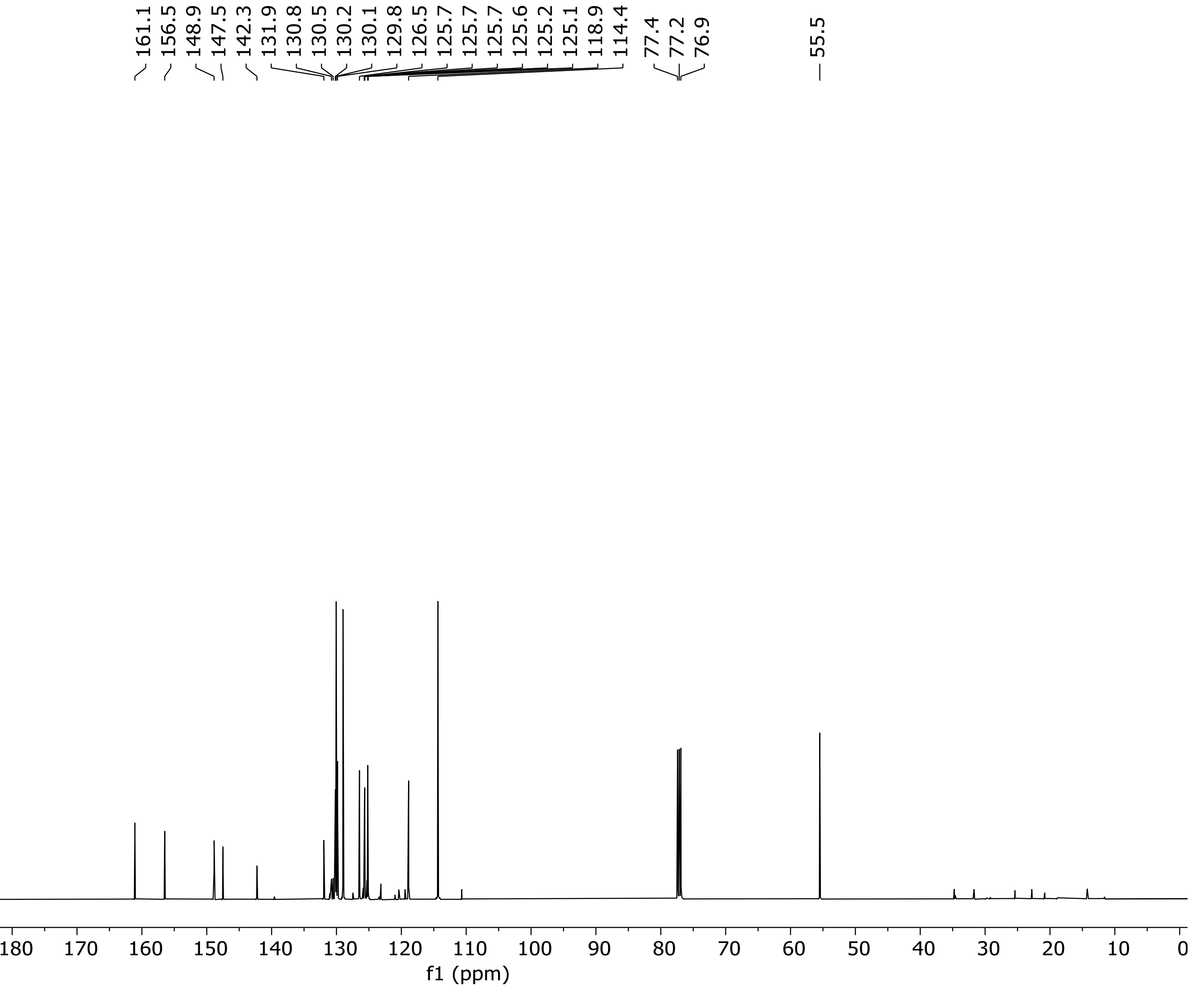


**$^{13}C$ NMR Spectrum of Compound S15 (126 MHz, $CDCl_3$)**

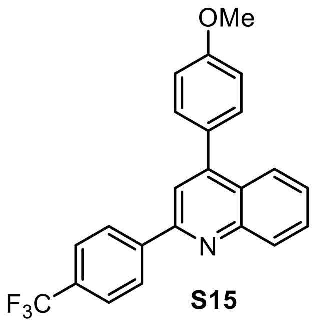


S15

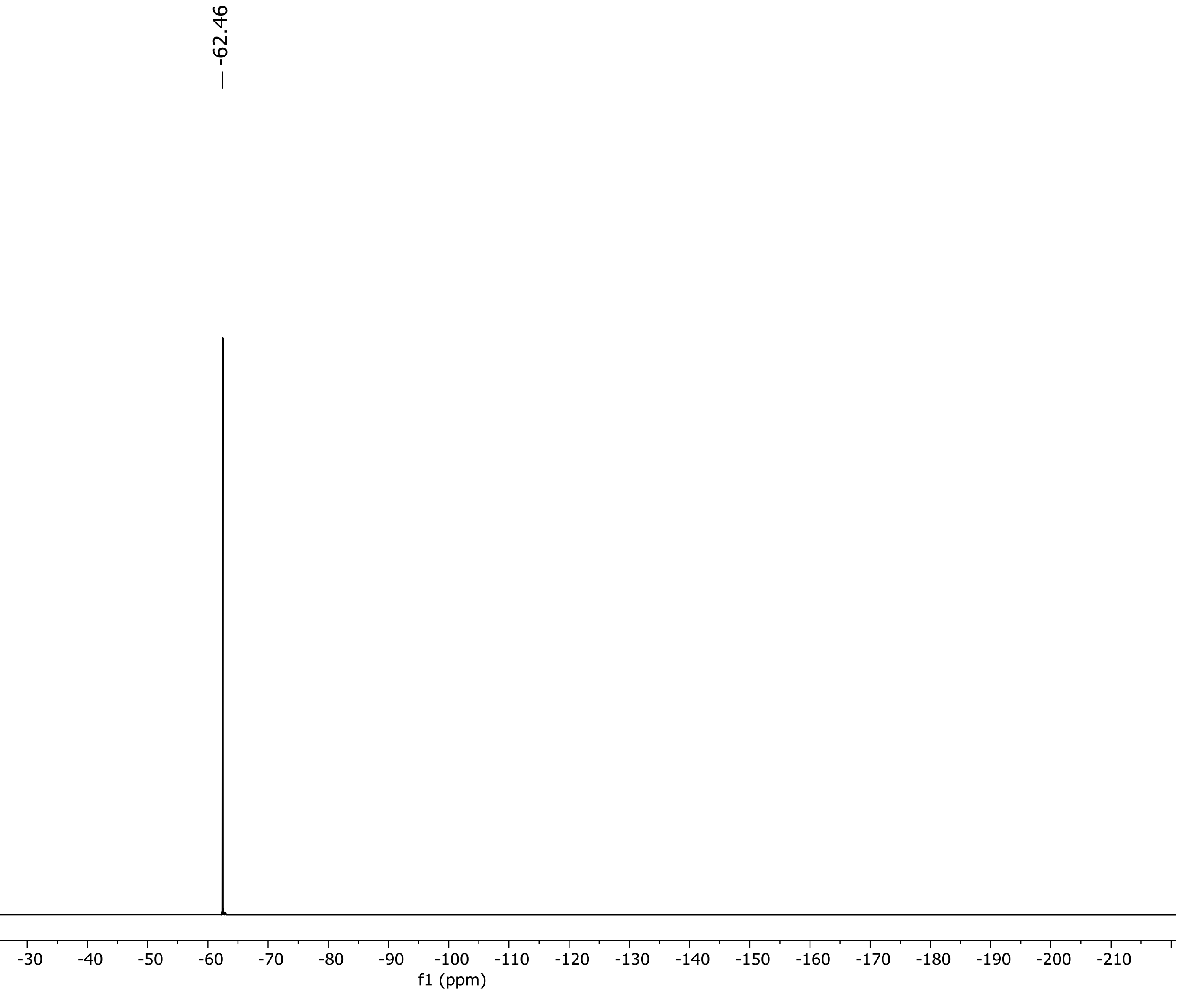


**$^{19}$F NMR Spectrum of Compound S15 (471 MHz, $CDCl_3$)**

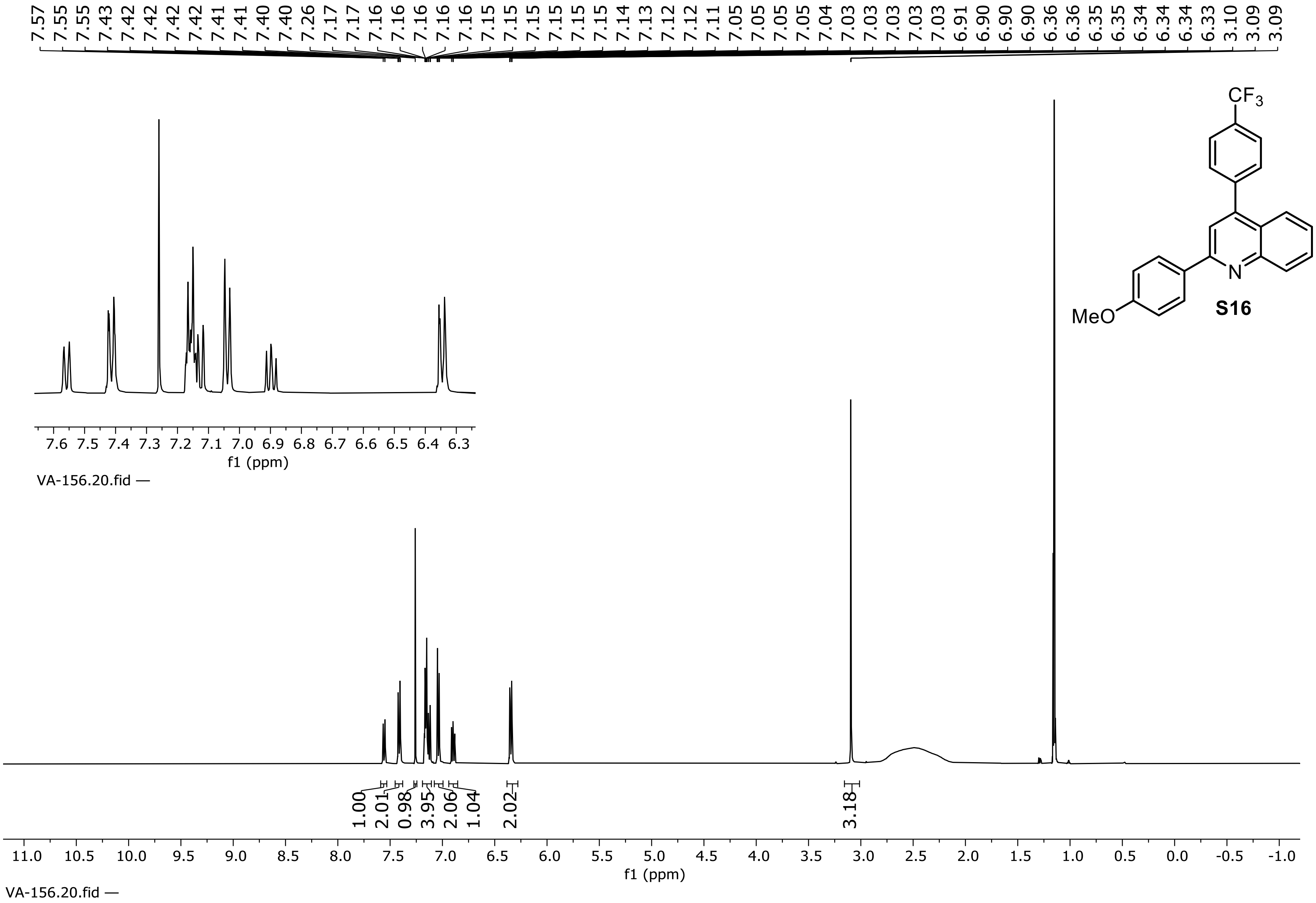


**$^{1}$H NMR Spectrum of Compound S16 (500 MHz, $CD_3CN$)**

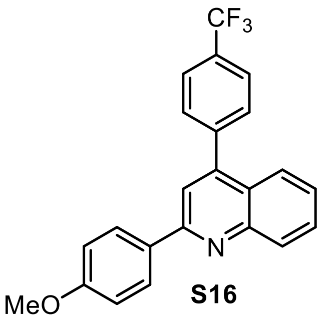

**S16**

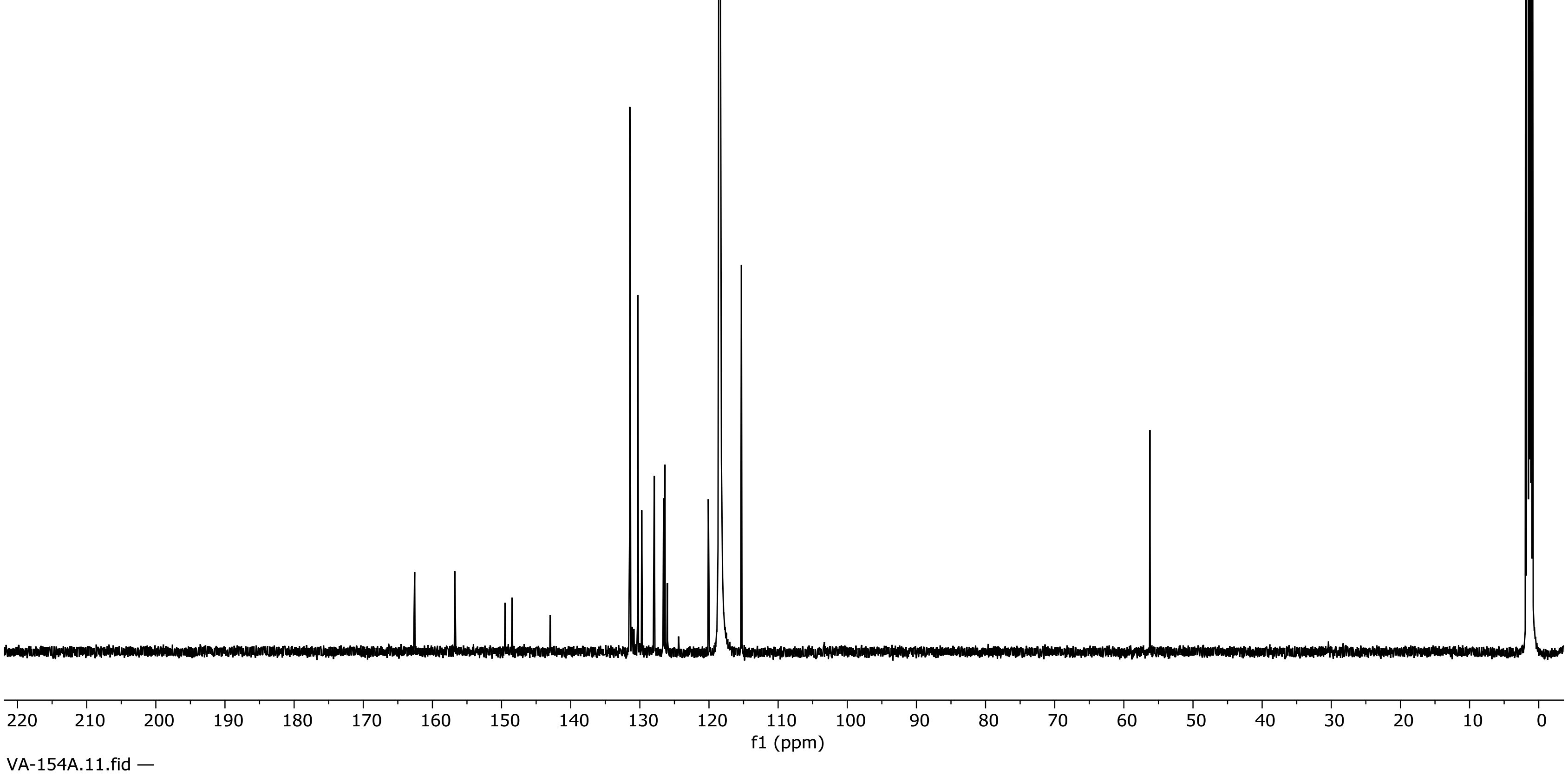


**[13]C NMR Spectrum of Compound S16 (126 MHz, $CD_3CN$)**

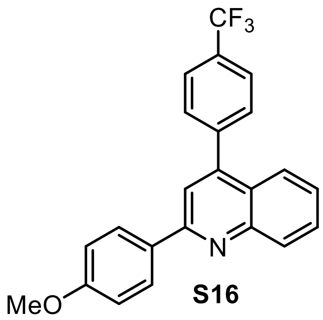


**S16**

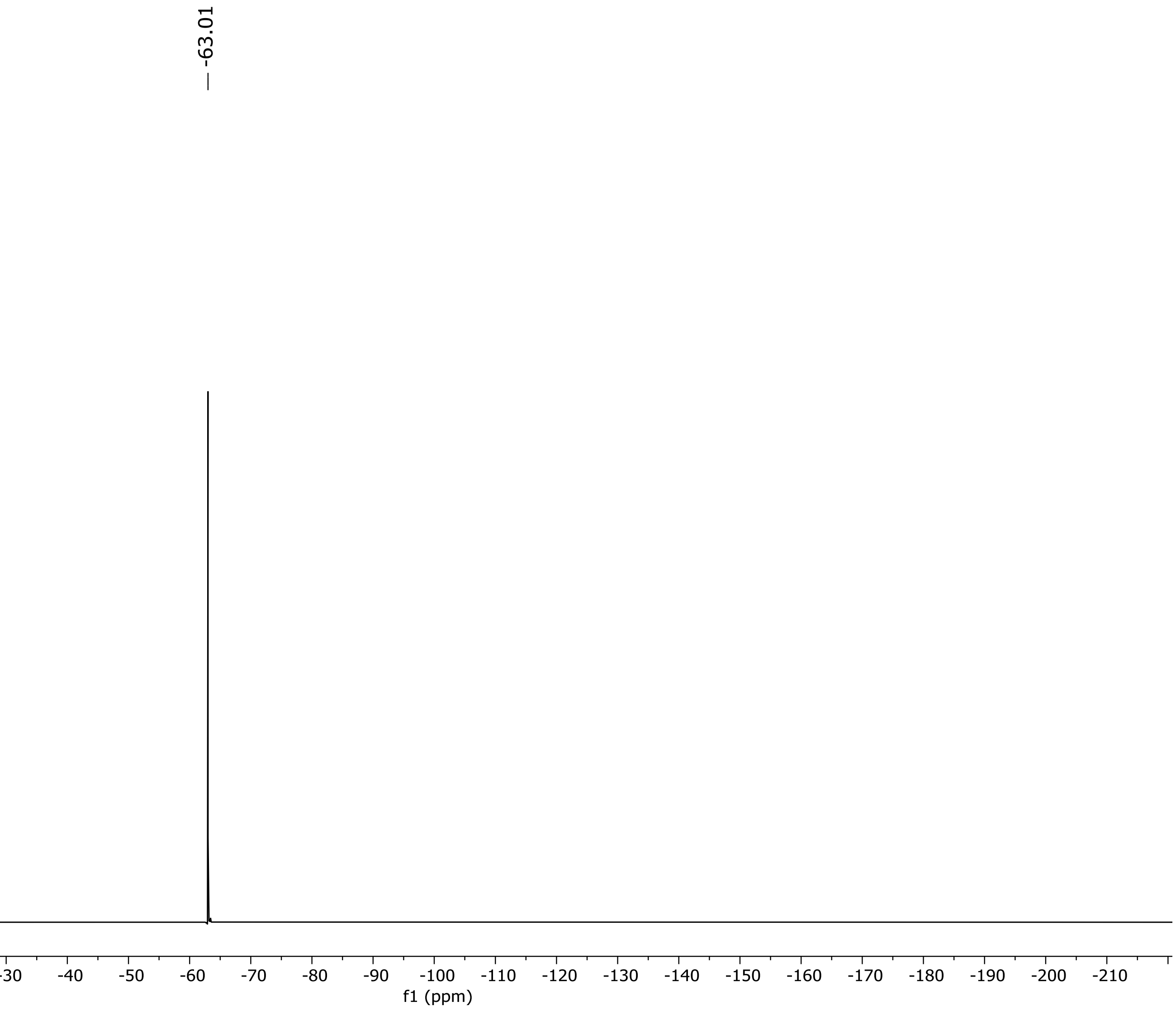


**$^{19}F$ NMR Spectrum of Compound S16 (471 MHz, $CD_3CN$)**

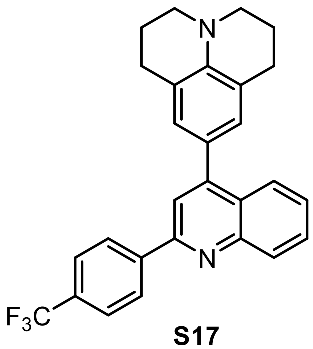

**S17**

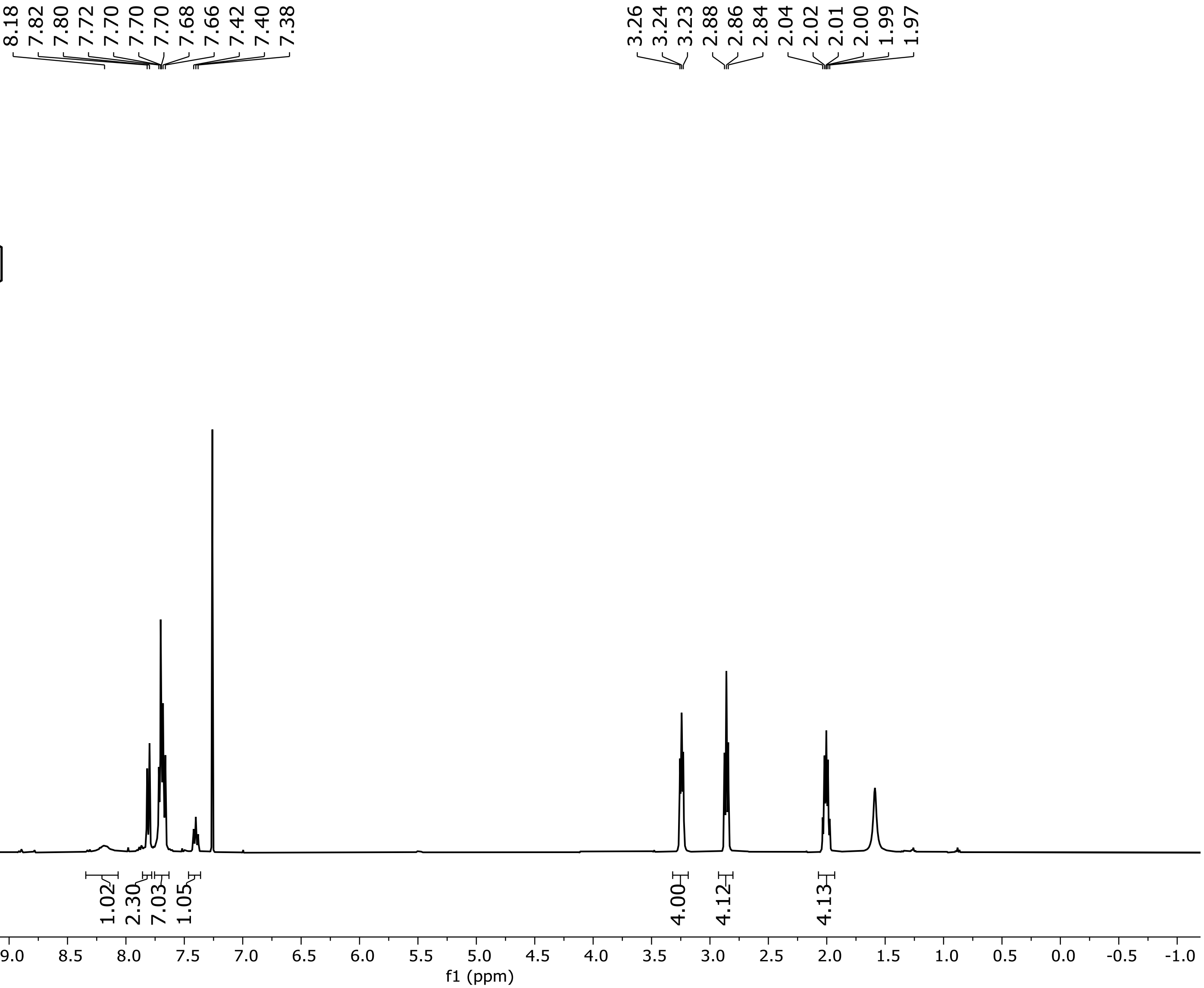


**$^1$H NMR Spectrum of Compound S17 (400 MHz, $CDCl_3$)**

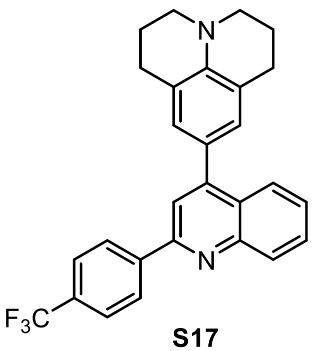

**S17**

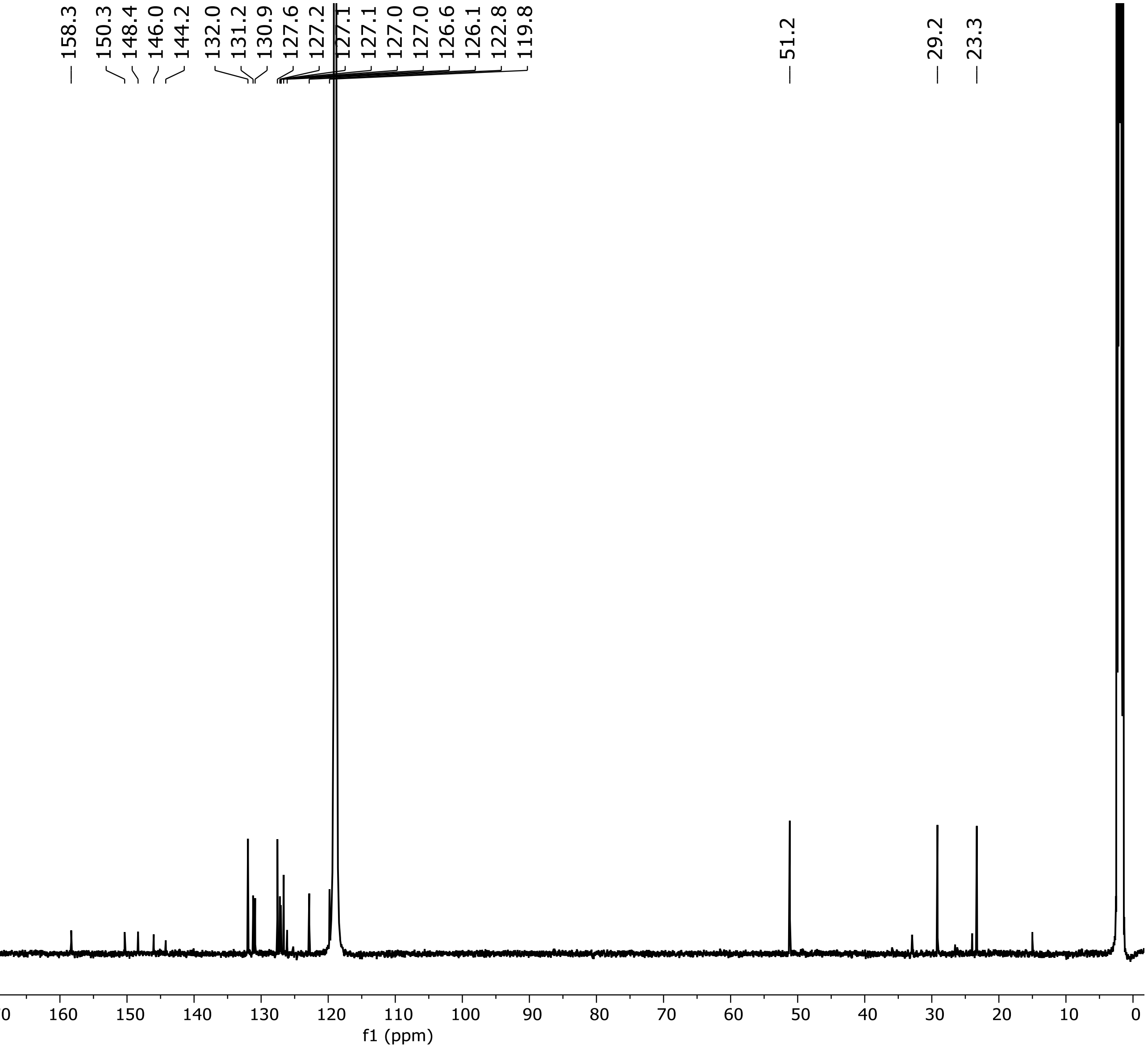


**$^{13}C$ NMR Spectrum of Compound S17 (151 MHz, $CD_3CN$)**

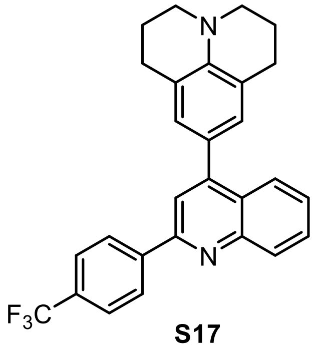


**S17**

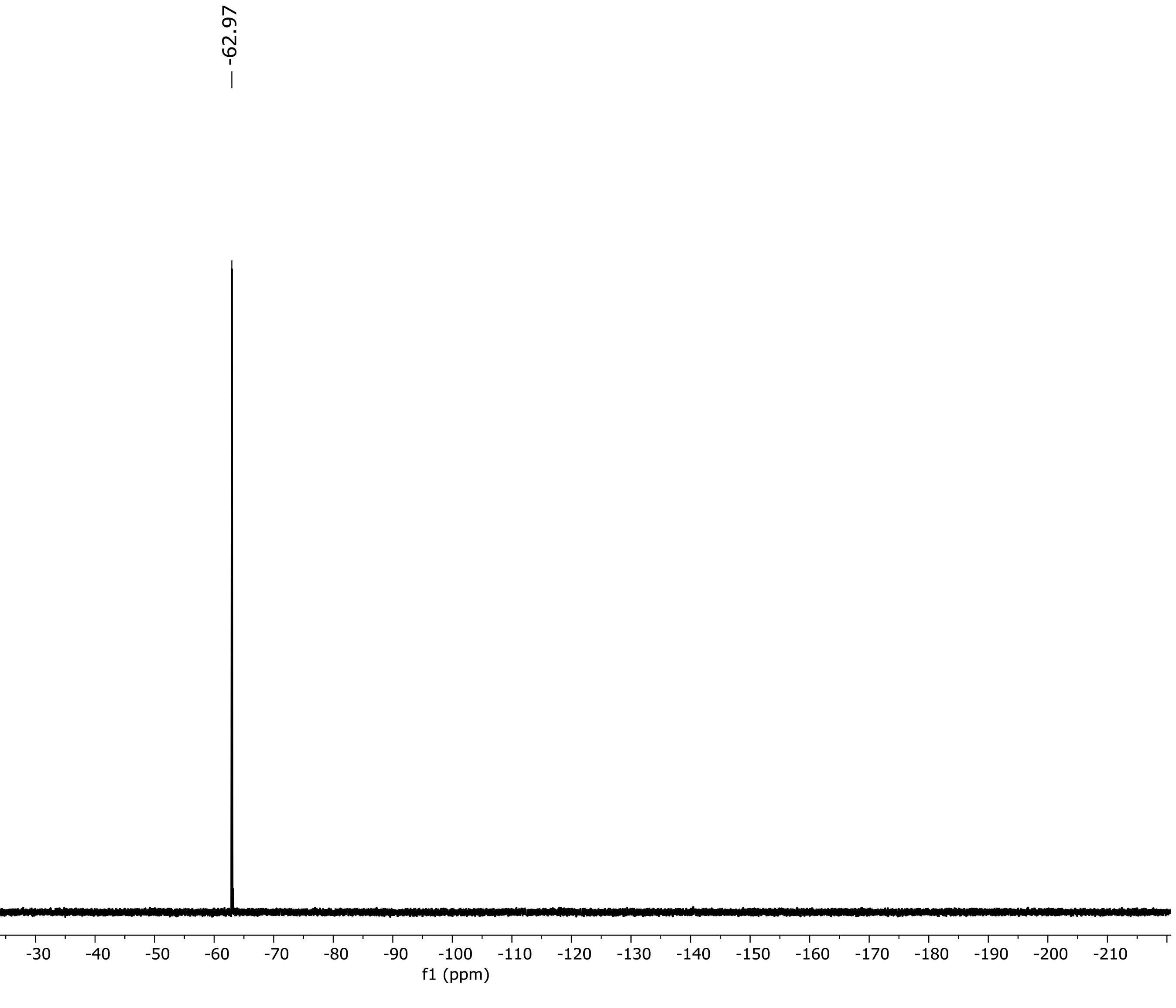


**$^{19}F$ NMR Spectrum of Compound S17 (471 MHz, $CD_3CN$)**

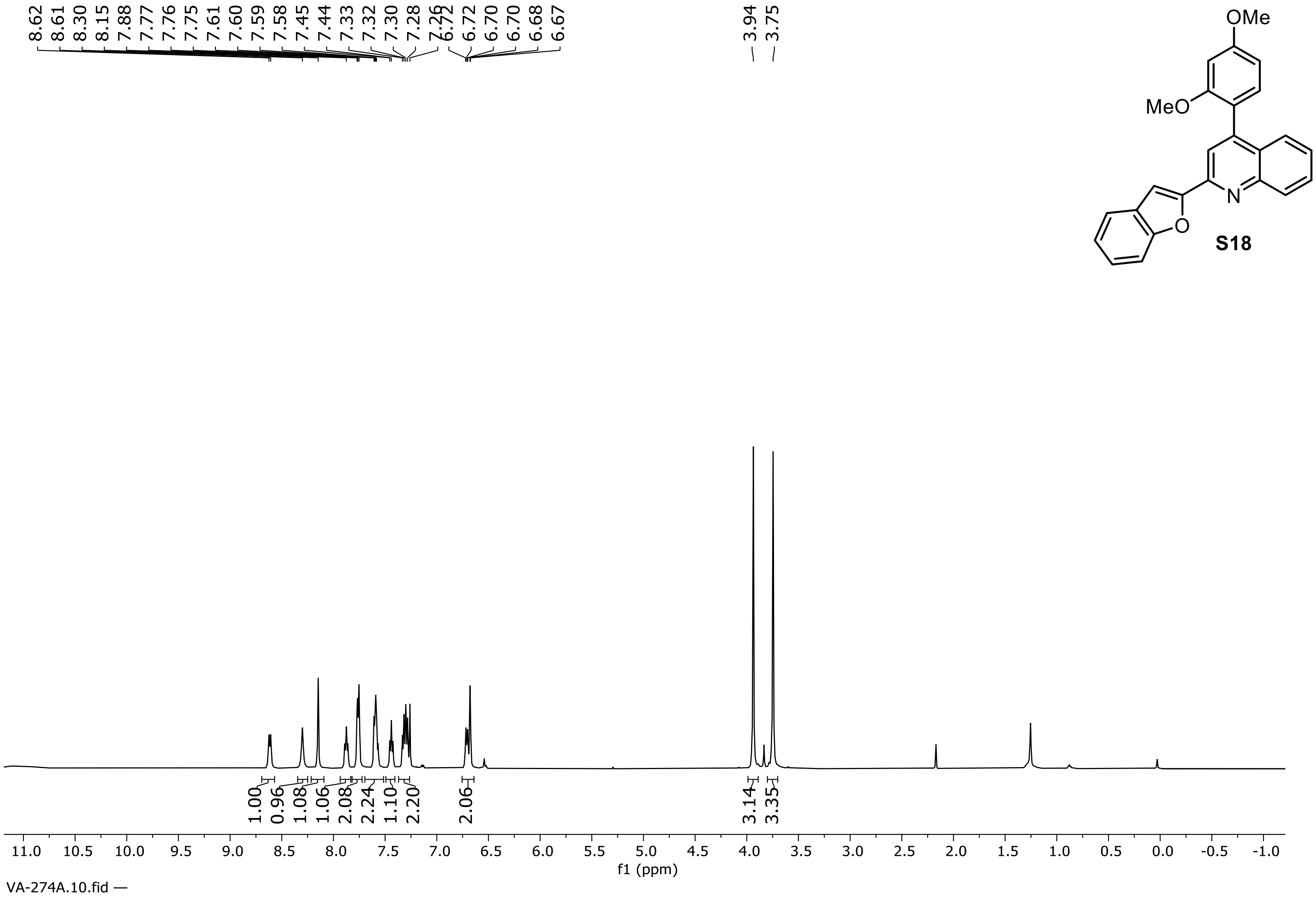


**$^1$H NMR Spectrum of Compound S18 (500 MHz, $CDCl_3$)**

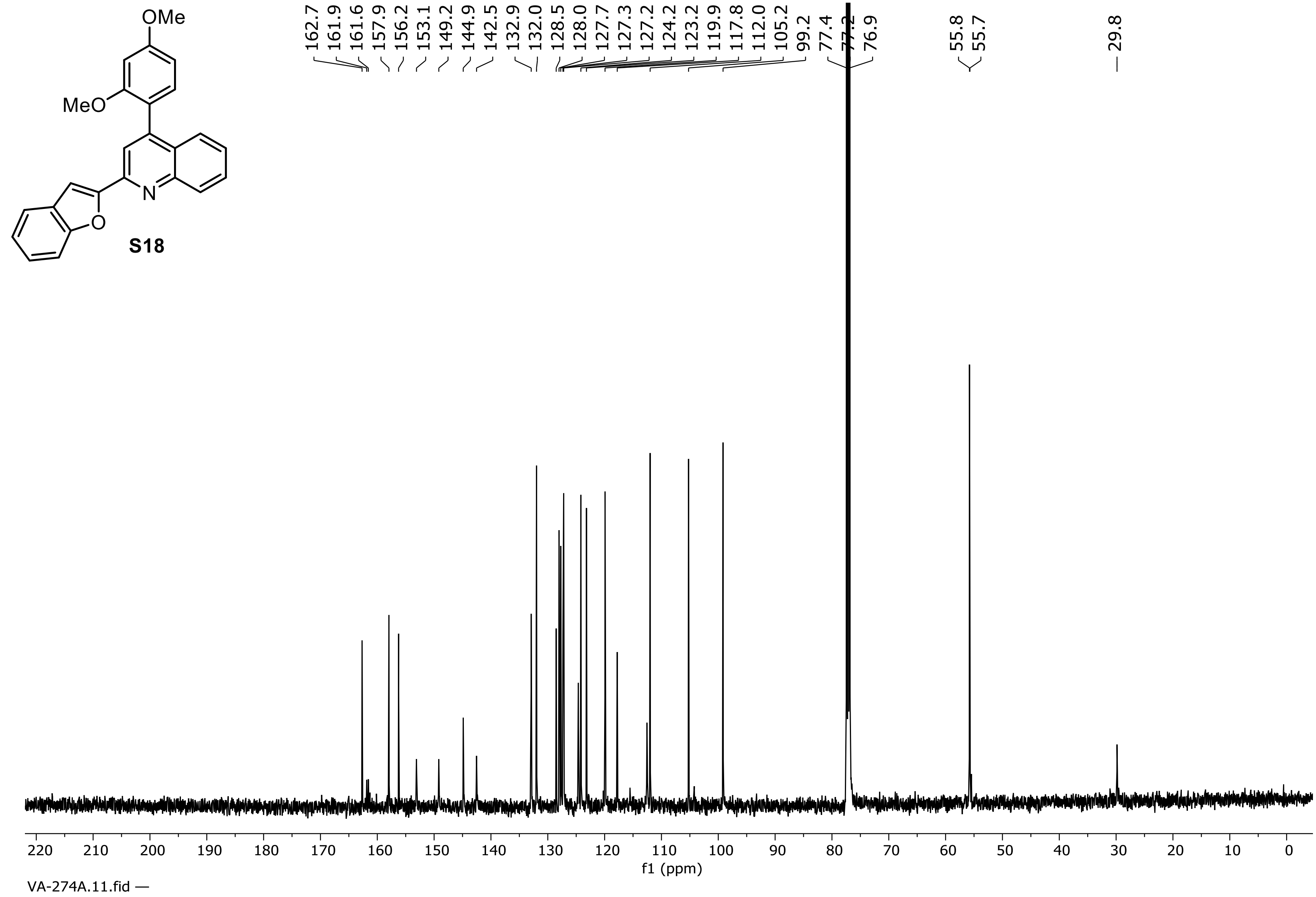


**$^{13}C$ NMR Spectrum of Compound S18 (126 MHz, $CD_3CN$)**

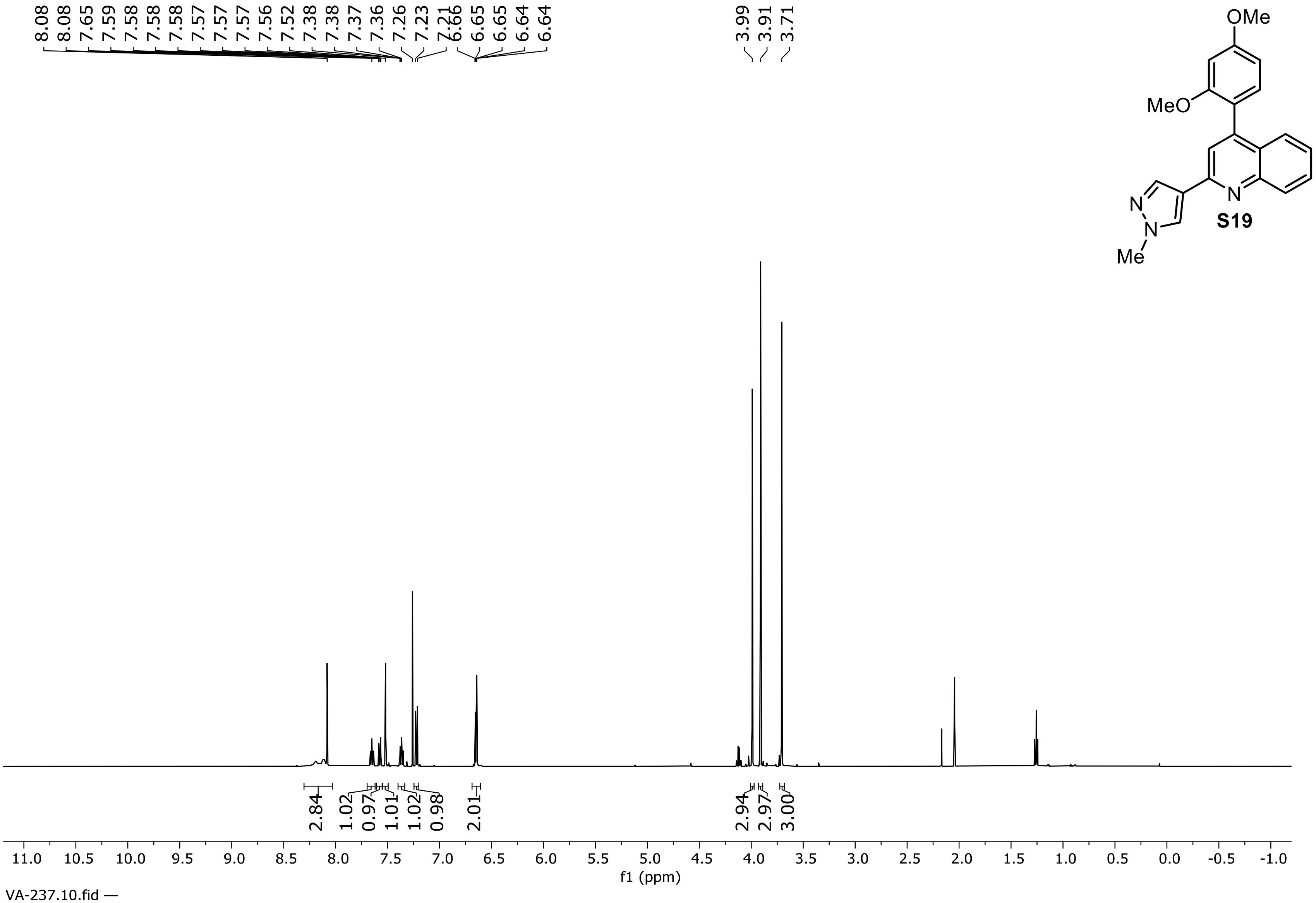


**[1]H NMR Spectrum of Compound S19 (500 MHz, $CDCl_3$)**

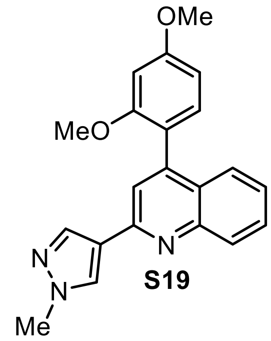


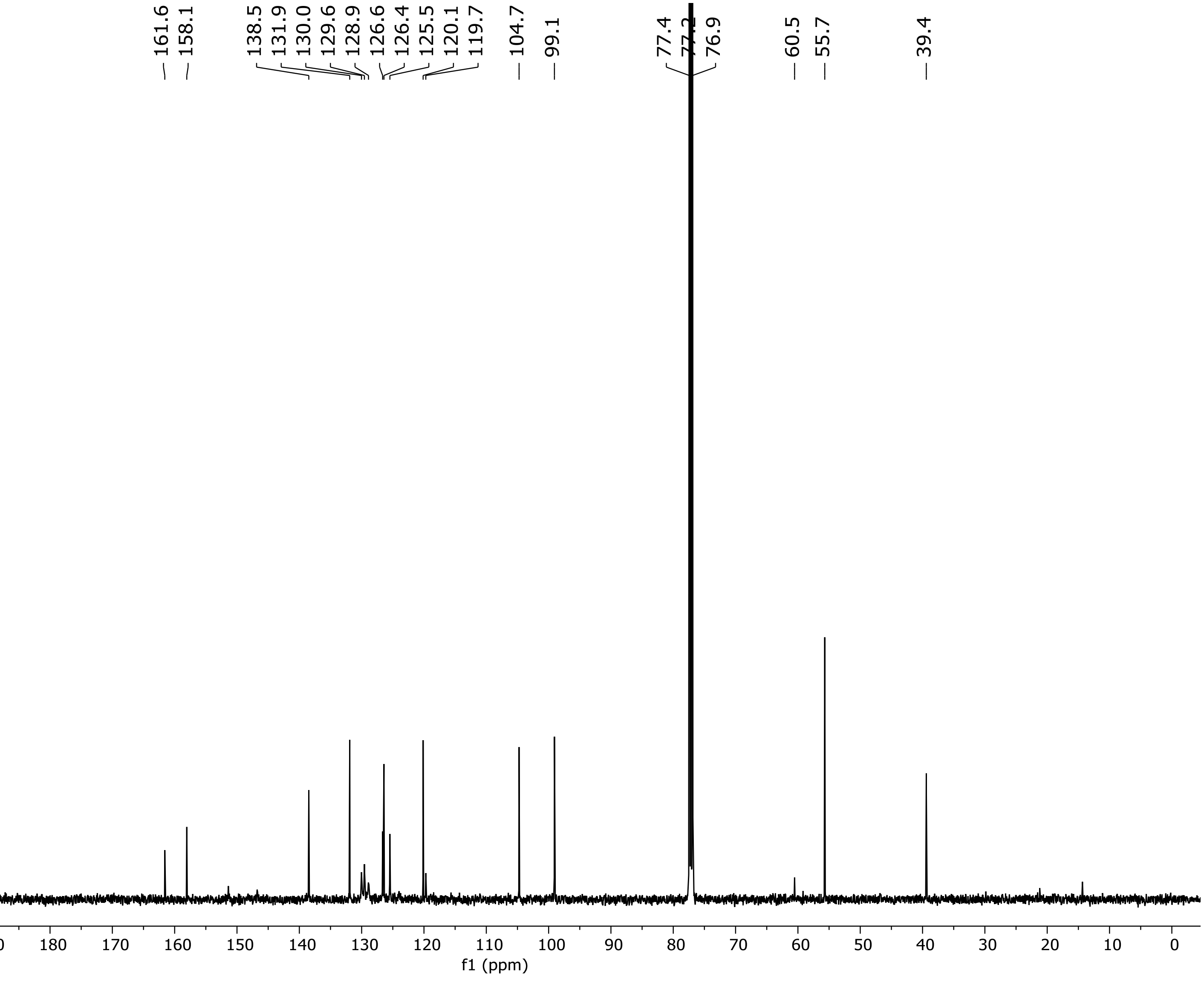


**$^{13}C$ NMR Spectrum of Compound S19 (126 MHz, $CDCl_3$)**

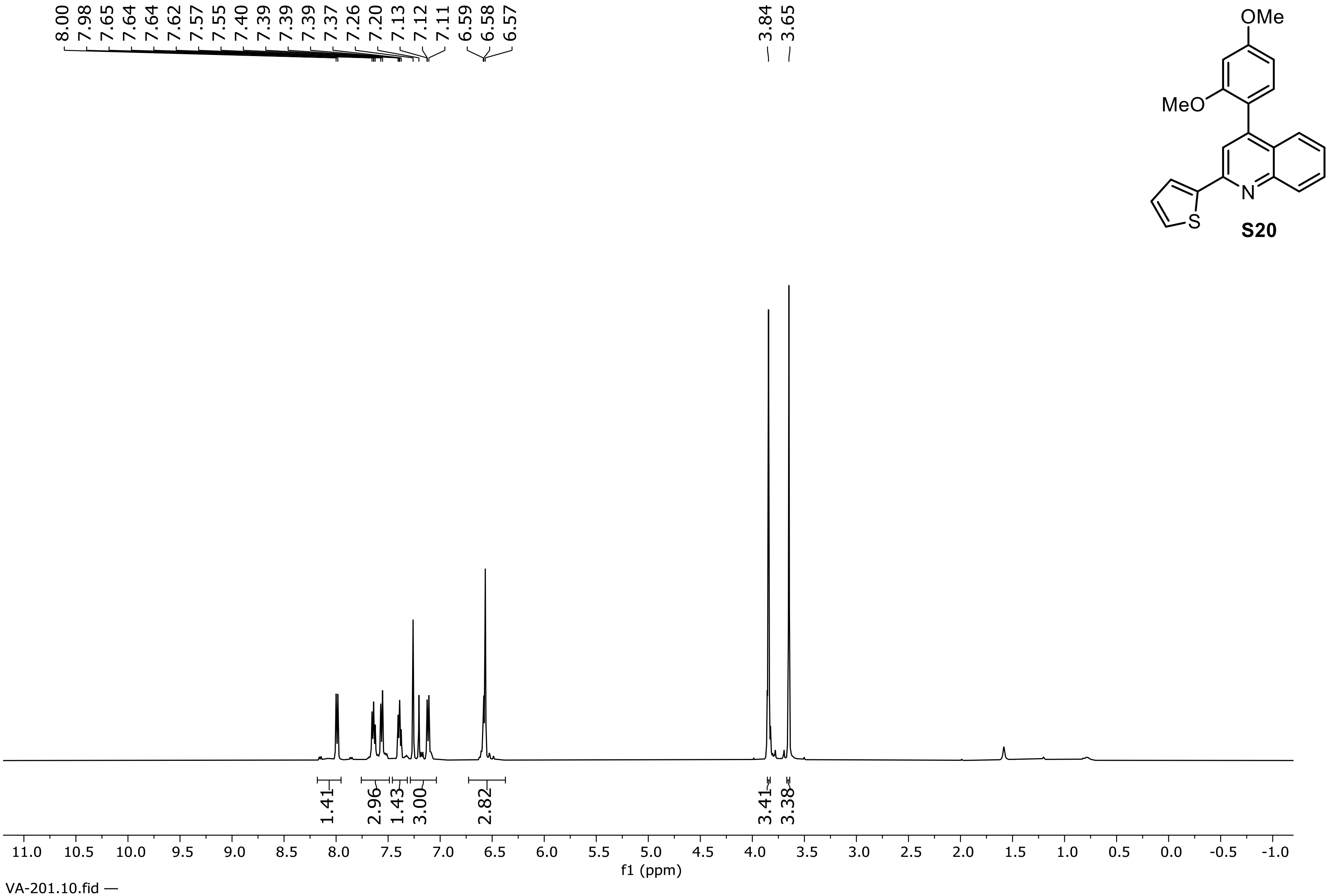


**S20**

**¹H NMR Spectrum of Compound S20 (500 MHz, $CDCl_3$)**

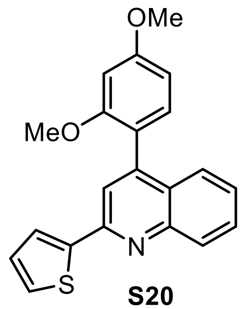


**S20**

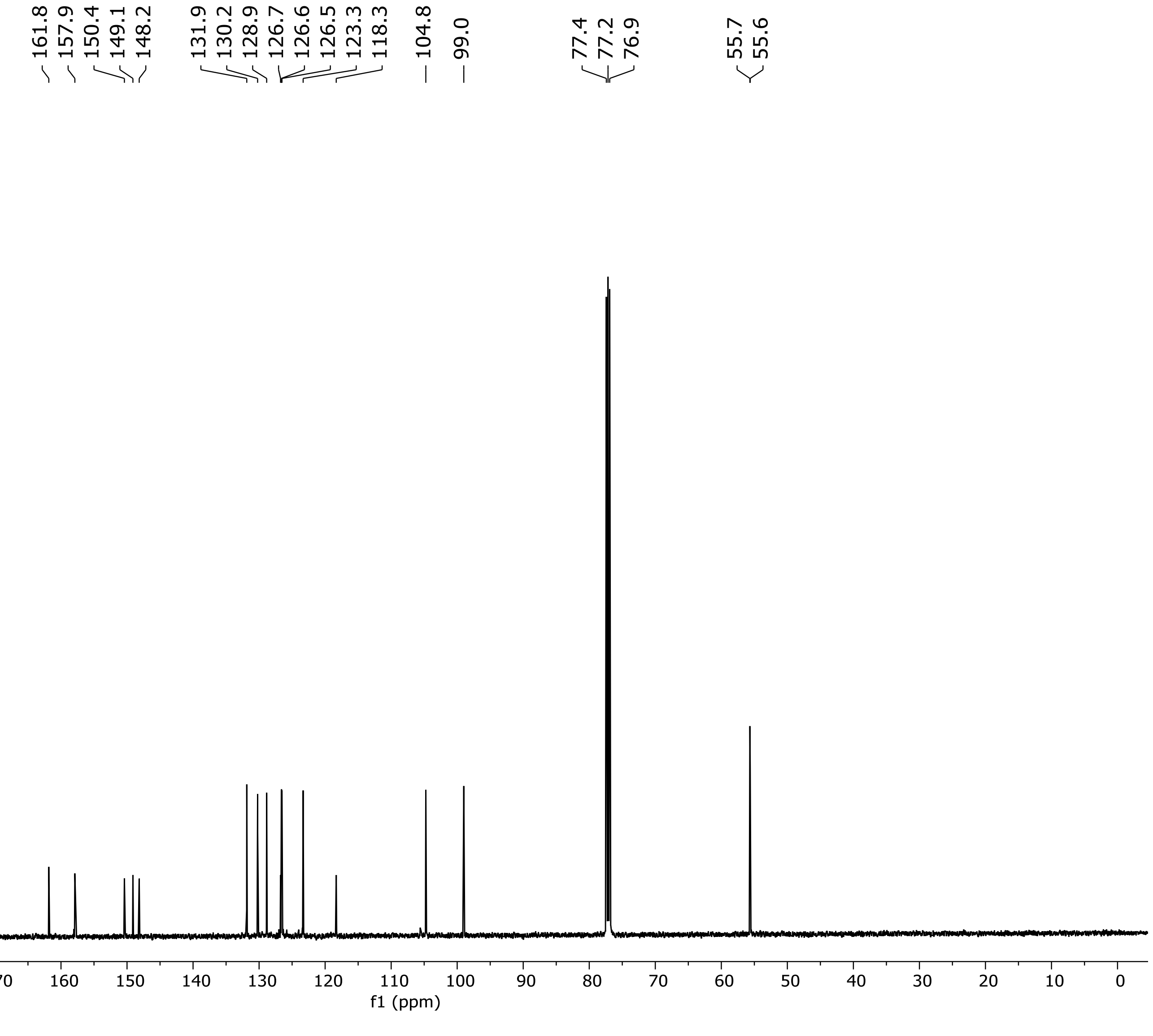


**$^{13}C$ NMR Spectrum of Compound S20 (126 MHz, $CDCl_3$)**

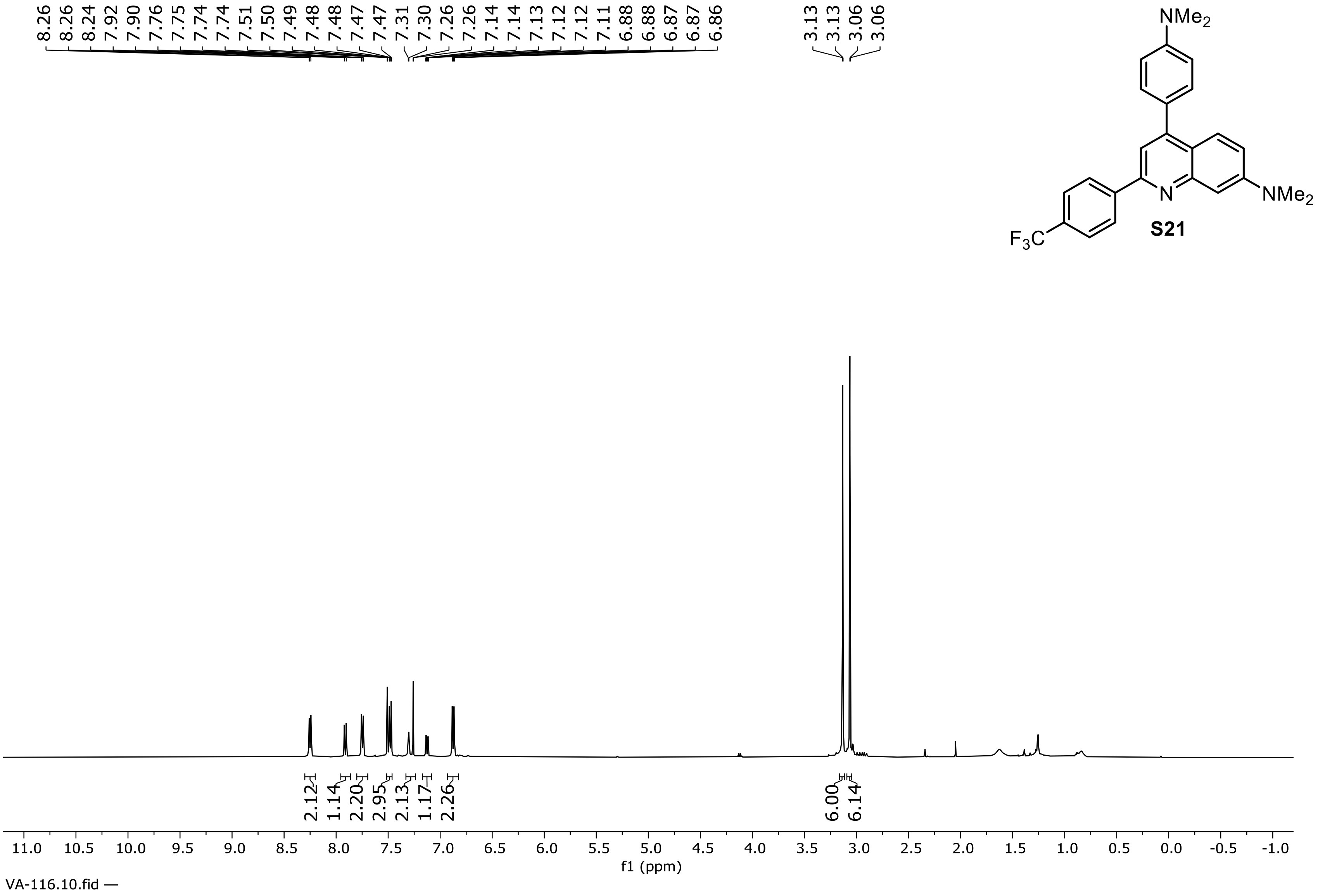


**S21**

**$^1$H NMR Spectrum of Compound S21 (500 MHz, $CDCl_3$)**

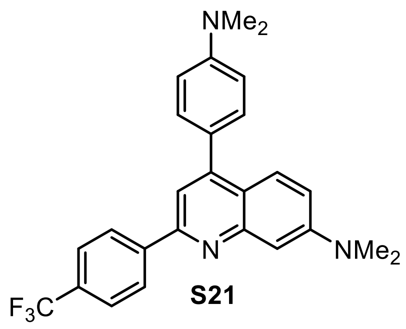


**S21**

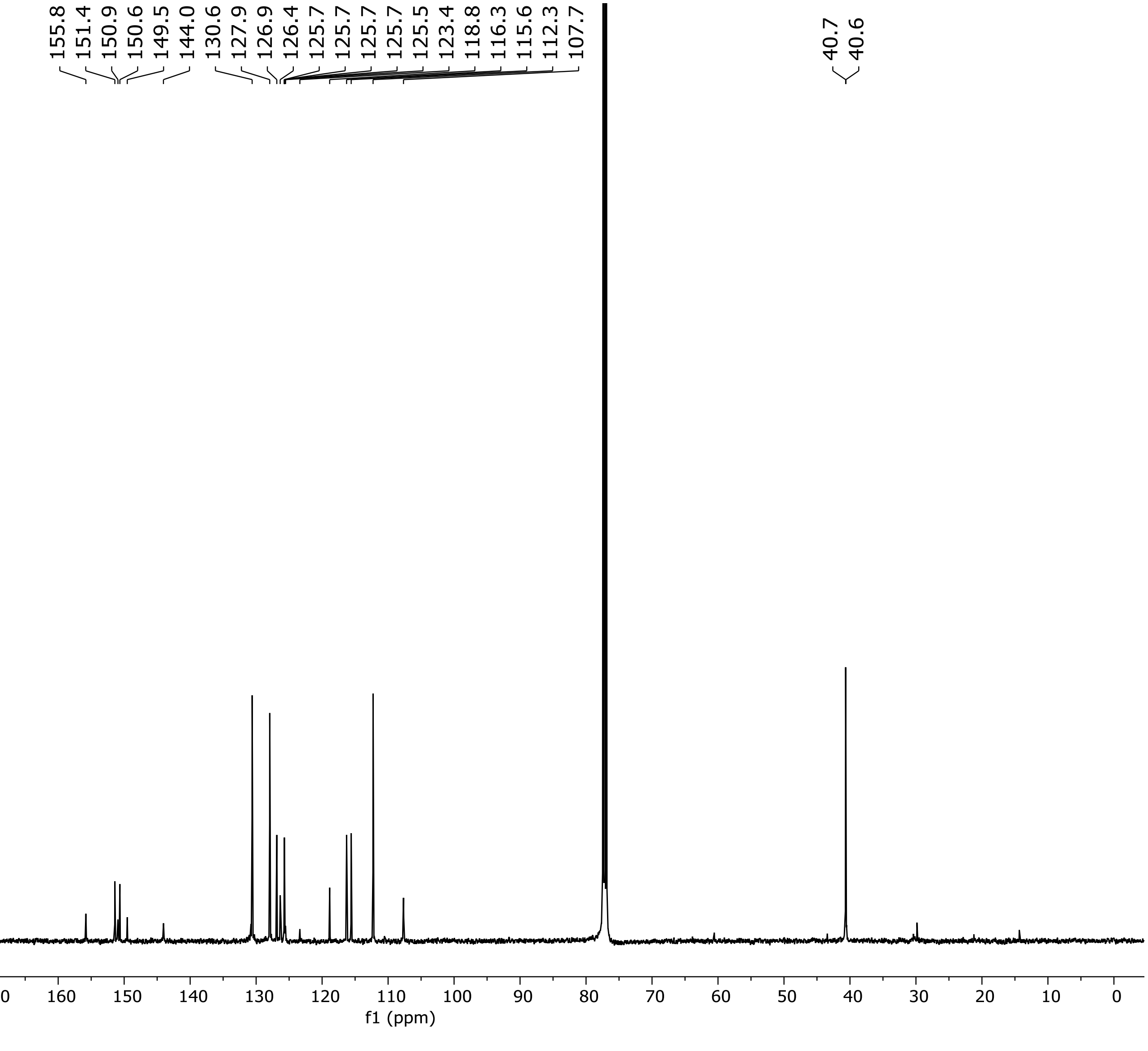


**$^{13}C$ NMR Spectrum of Compound S21 (126 MHz, $CDCl_3$)**

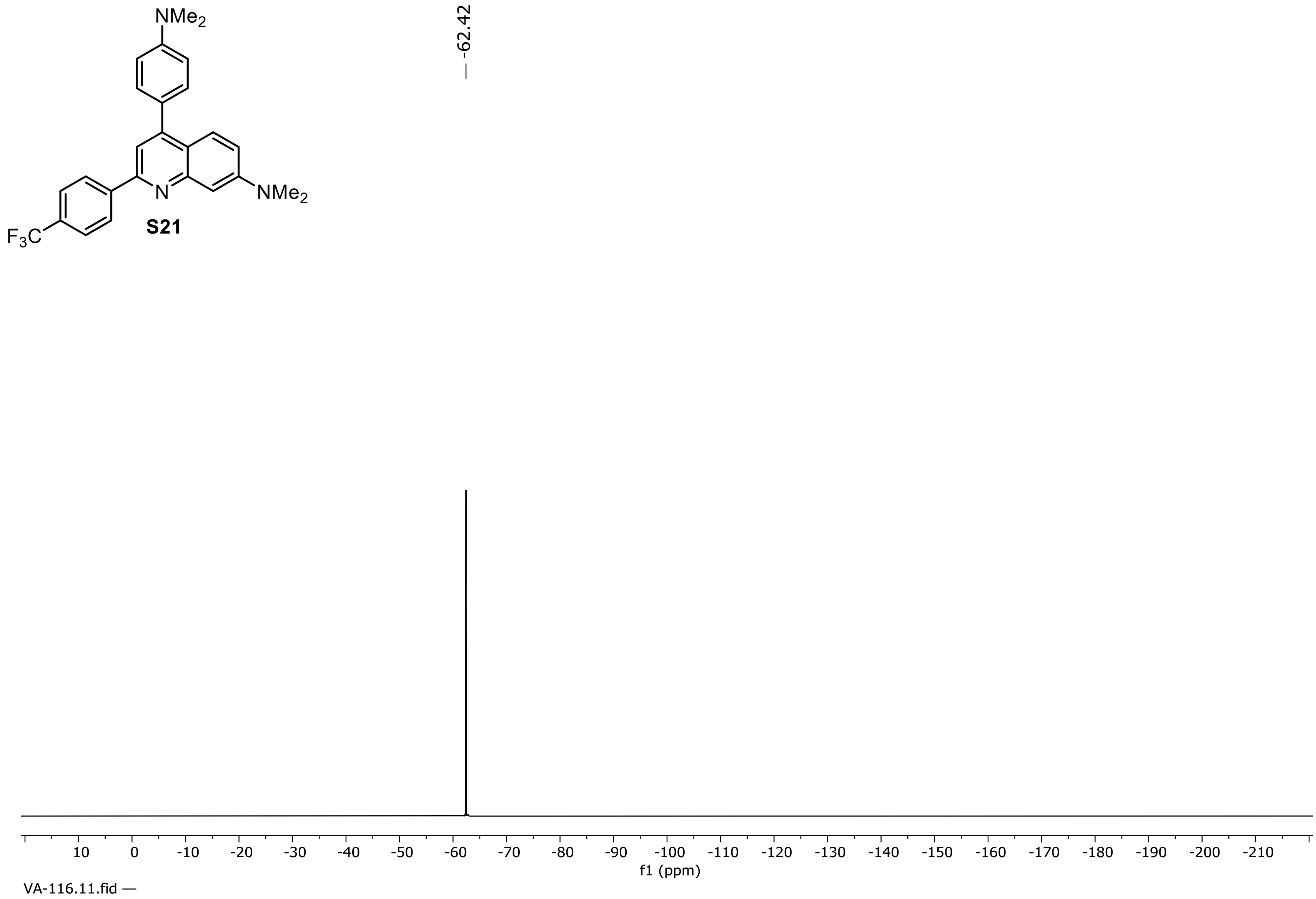


**$^{19}$F NMR Spectrum of Compound S21 (471 MHz, $CDCl_3$)**

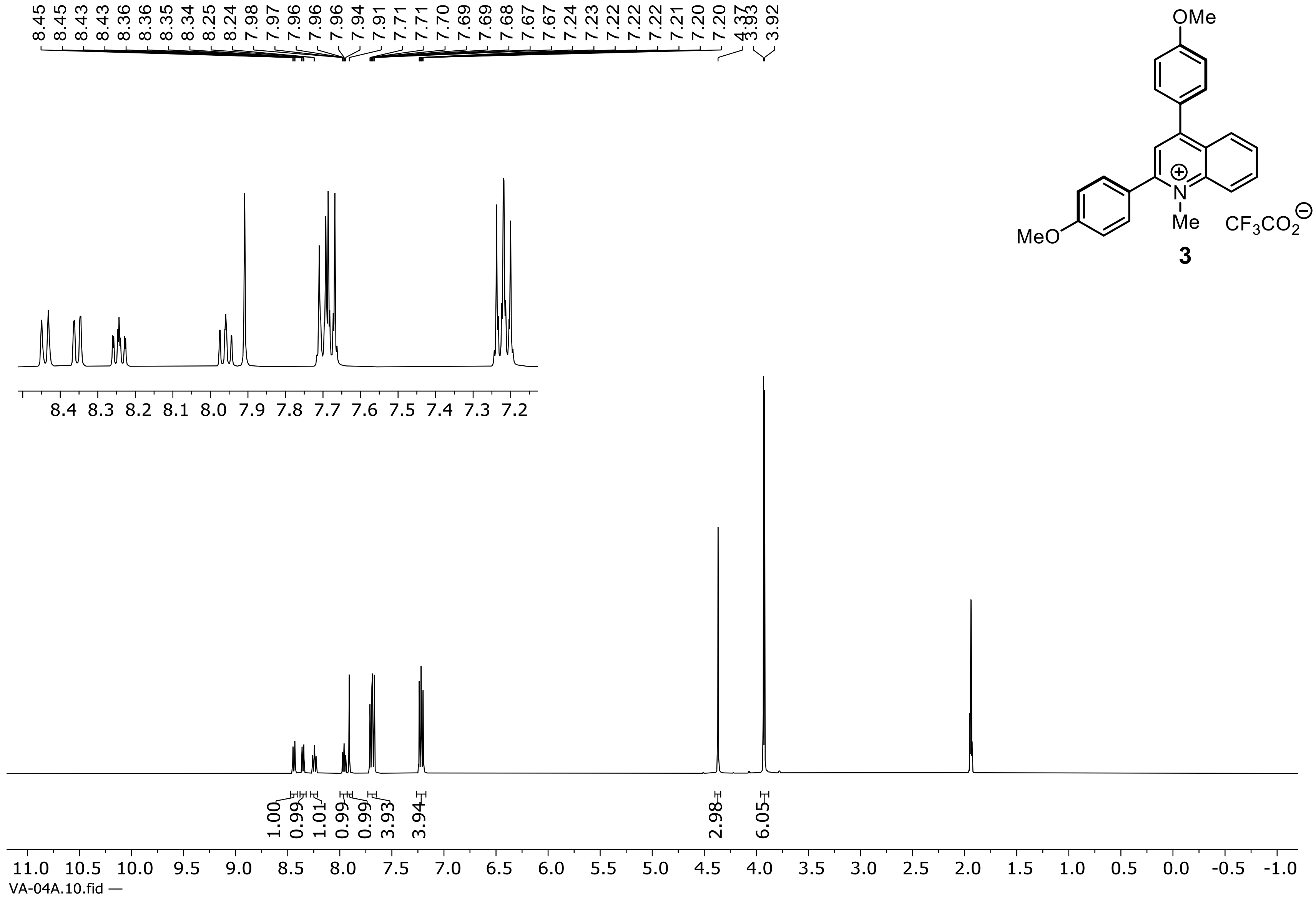


**3**

**$^1$H NMR Spectrum of compound 3 (500 MHz, $CD_3CN$)**

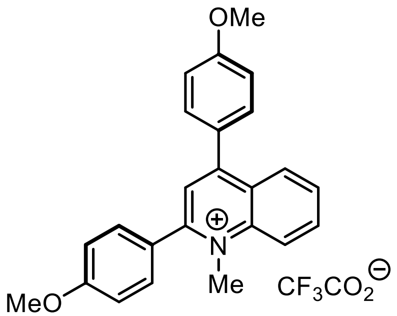


**3**

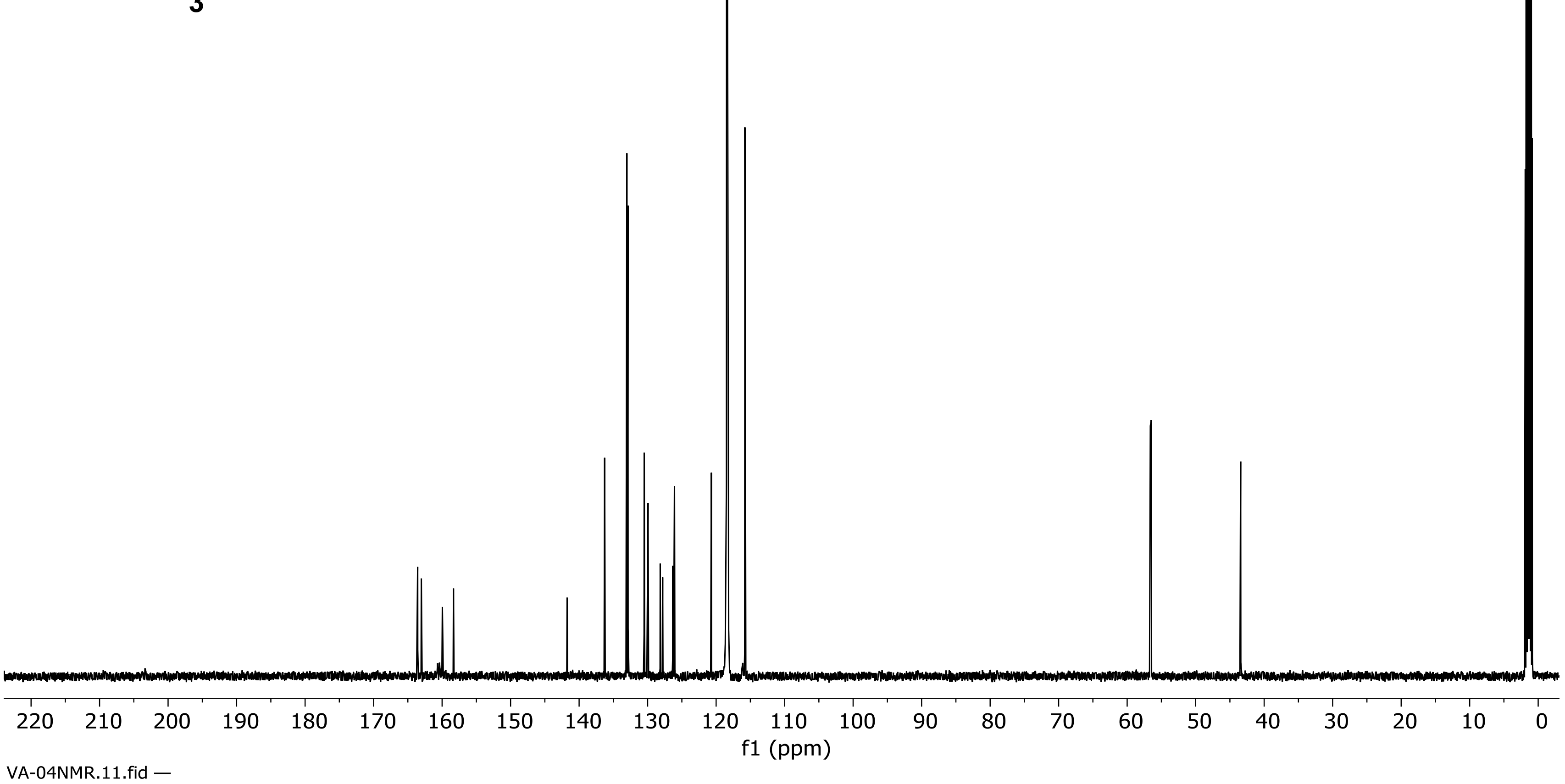


**$^{13}C$ NMR Spectrum of Compound 3 (126 MHz, $CD_3CN$)**

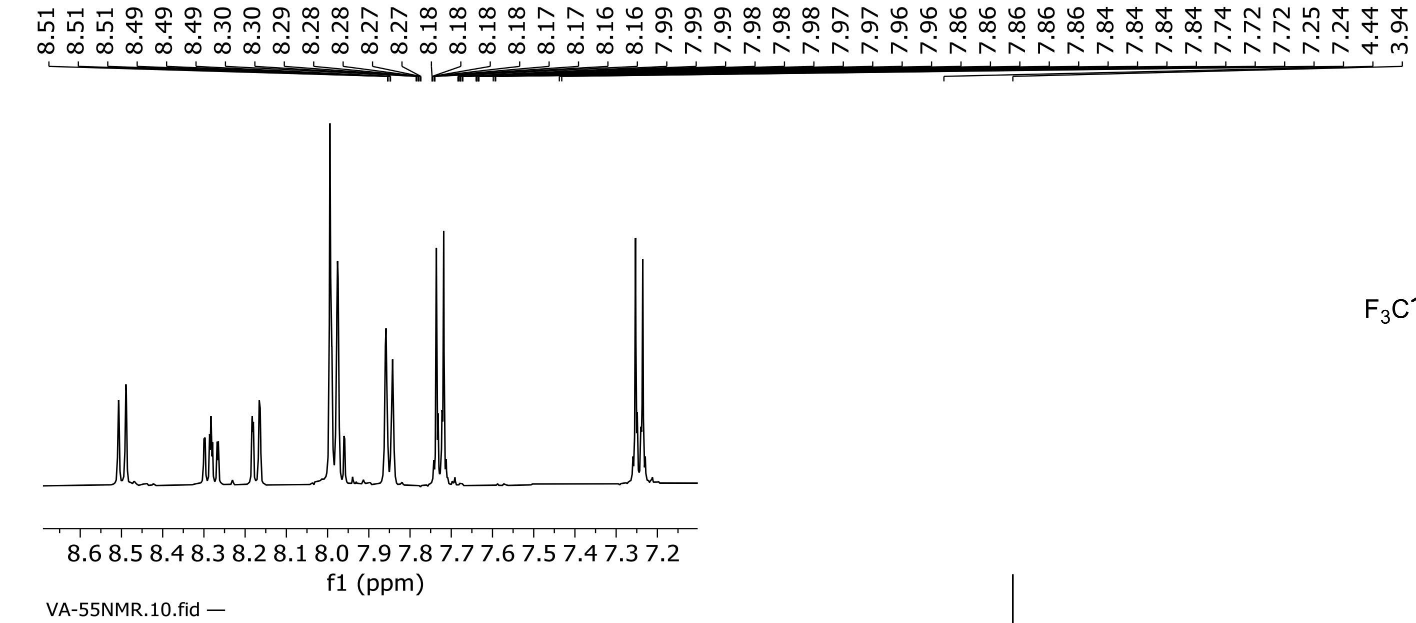


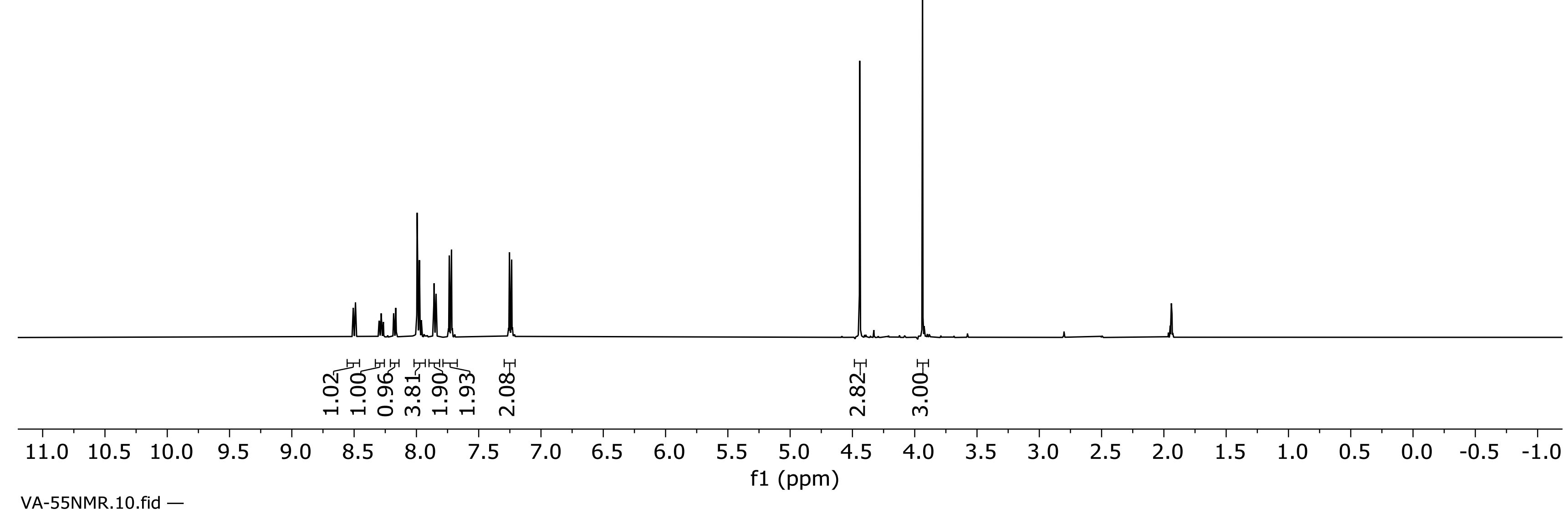


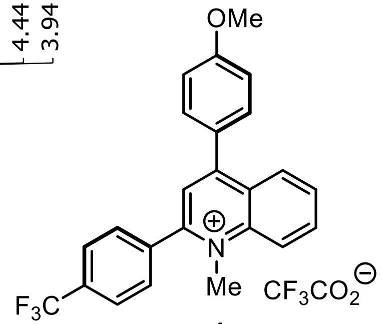


**4**

**$^1$H NMR Spectrum of compound 4 (500 MHz, $CD_3CN$)**

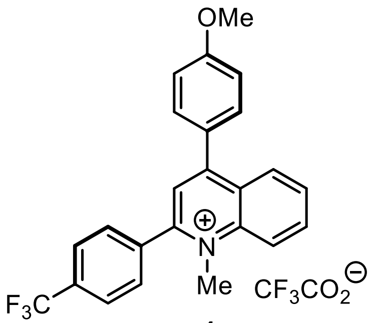

**4**

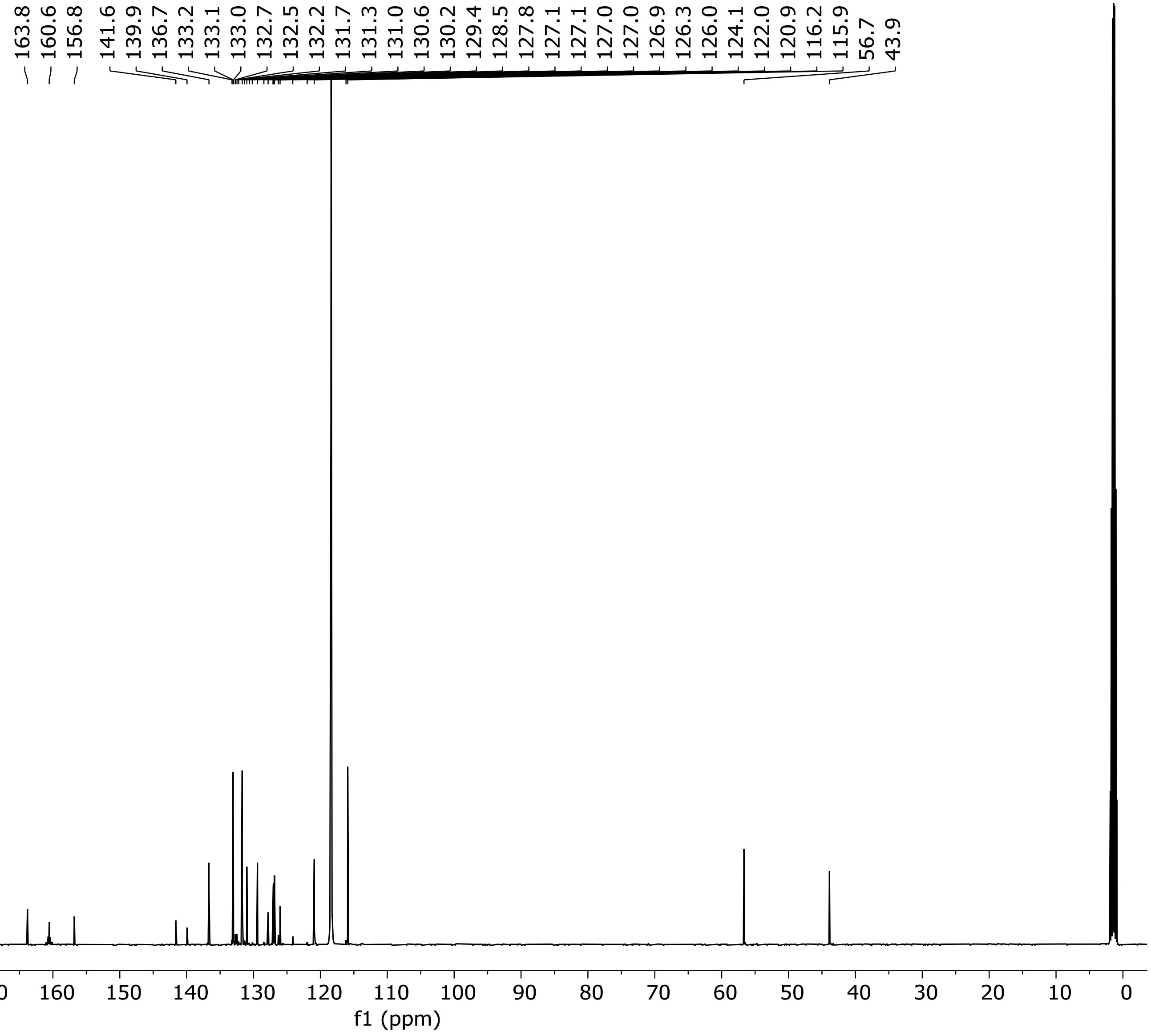


**$^{13}C$ NMR Spectrum of Compound 4 (126 MHz, $CD_3CN$)**

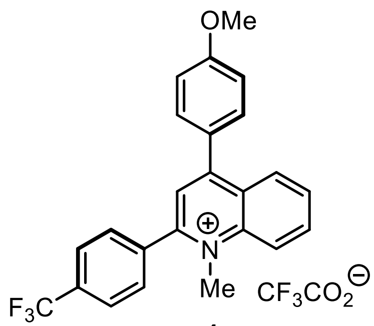


**4**

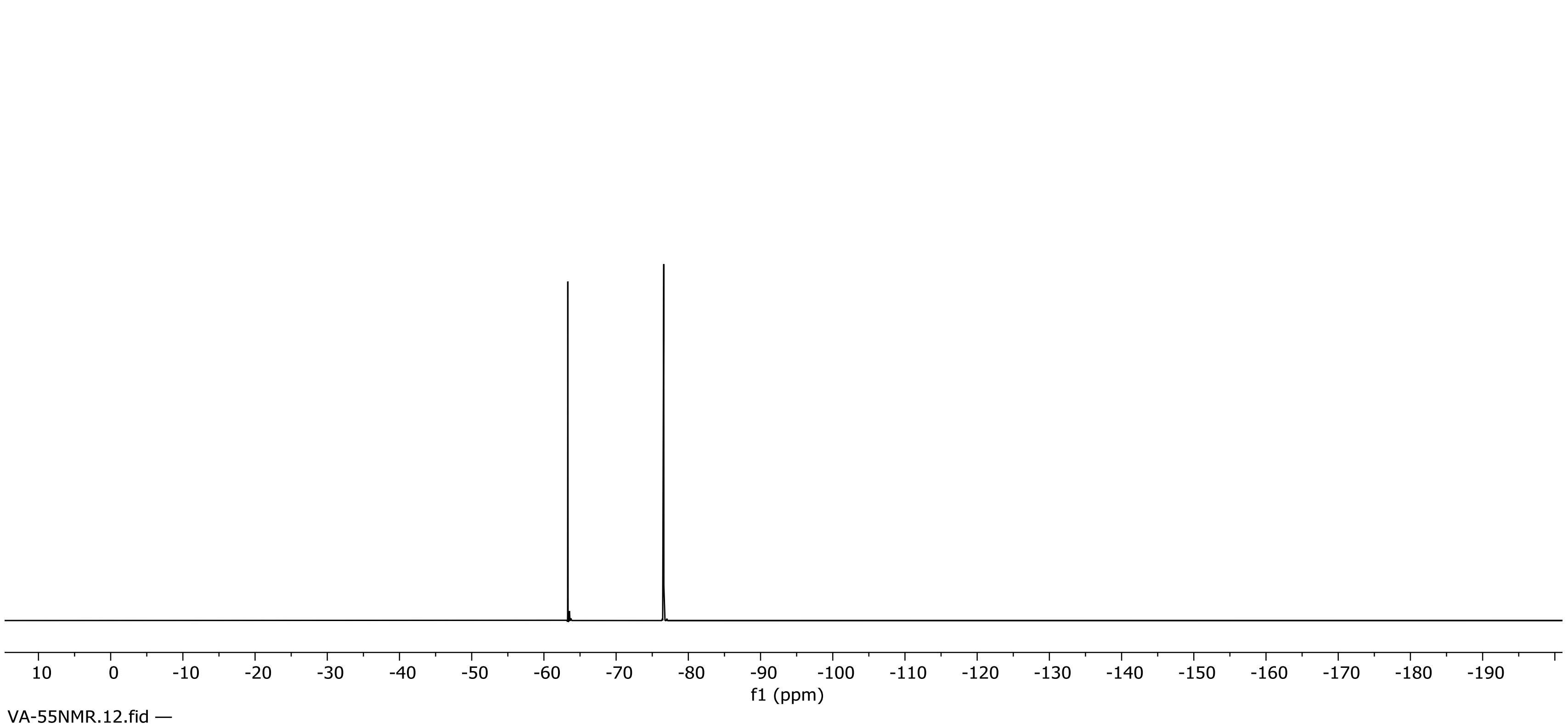


**$^{19}F$ NMR Spectrum of Compound 4 (471 MHz, $CD_3CN$)**

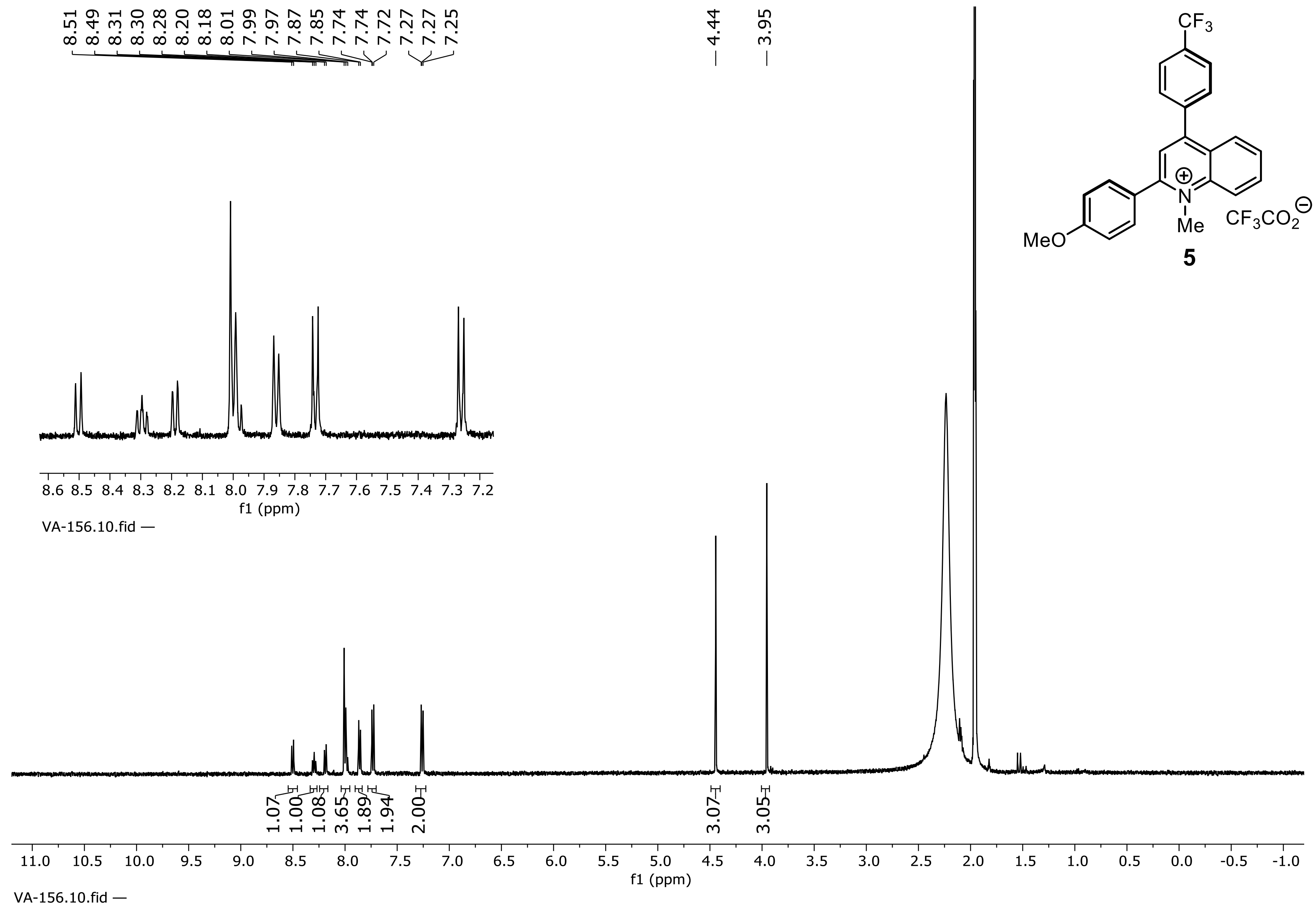


**$^1H$ NMR Spectrum of compound 5 (500 MHz, $CD_3CN$)**

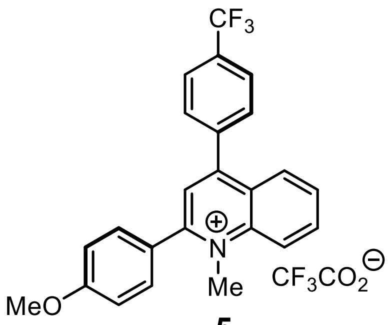


**5**

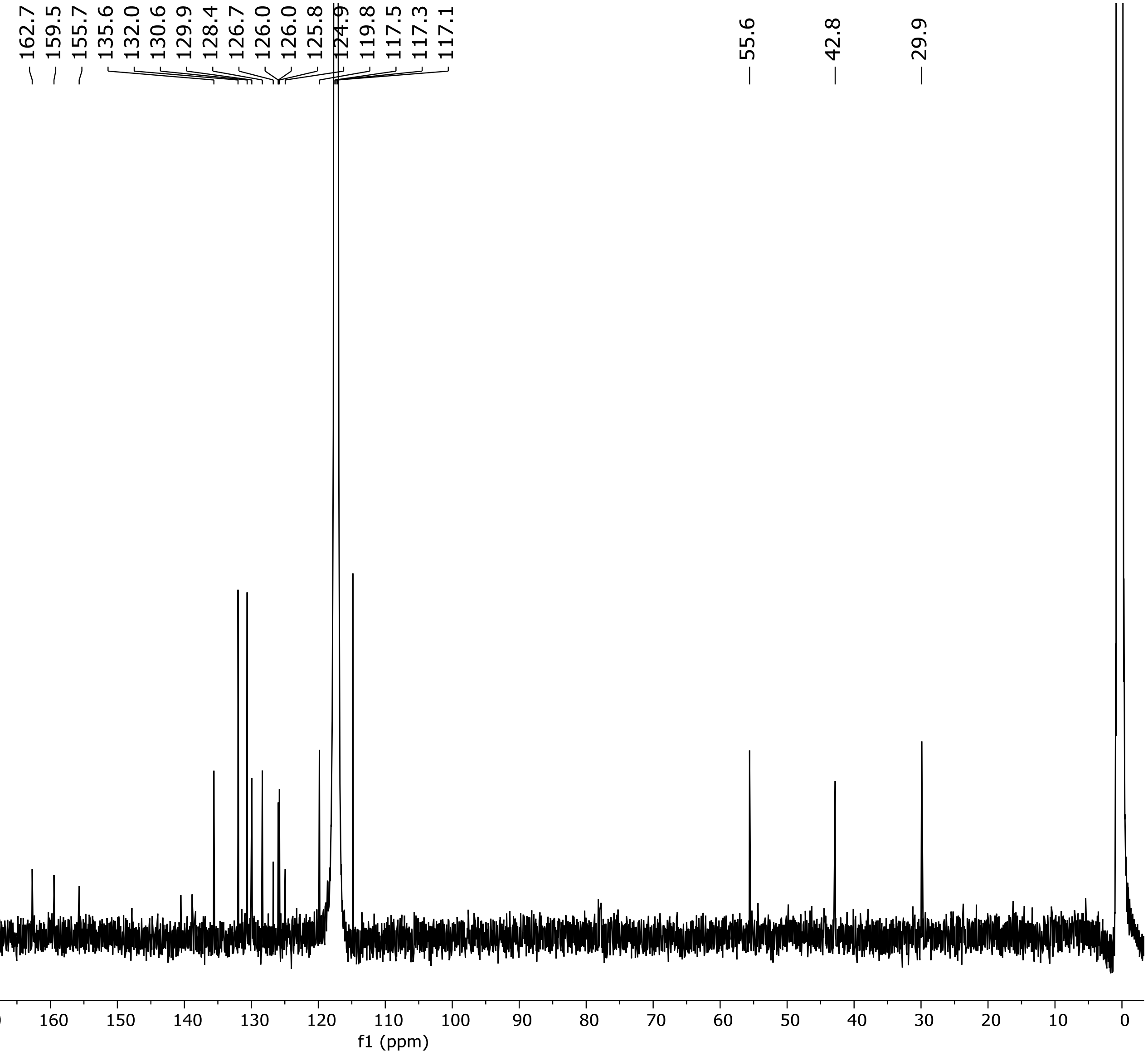


$^{13}$C

**NMR Spectrum of Compound 5 (151 MHz, $CD_3CN$)**

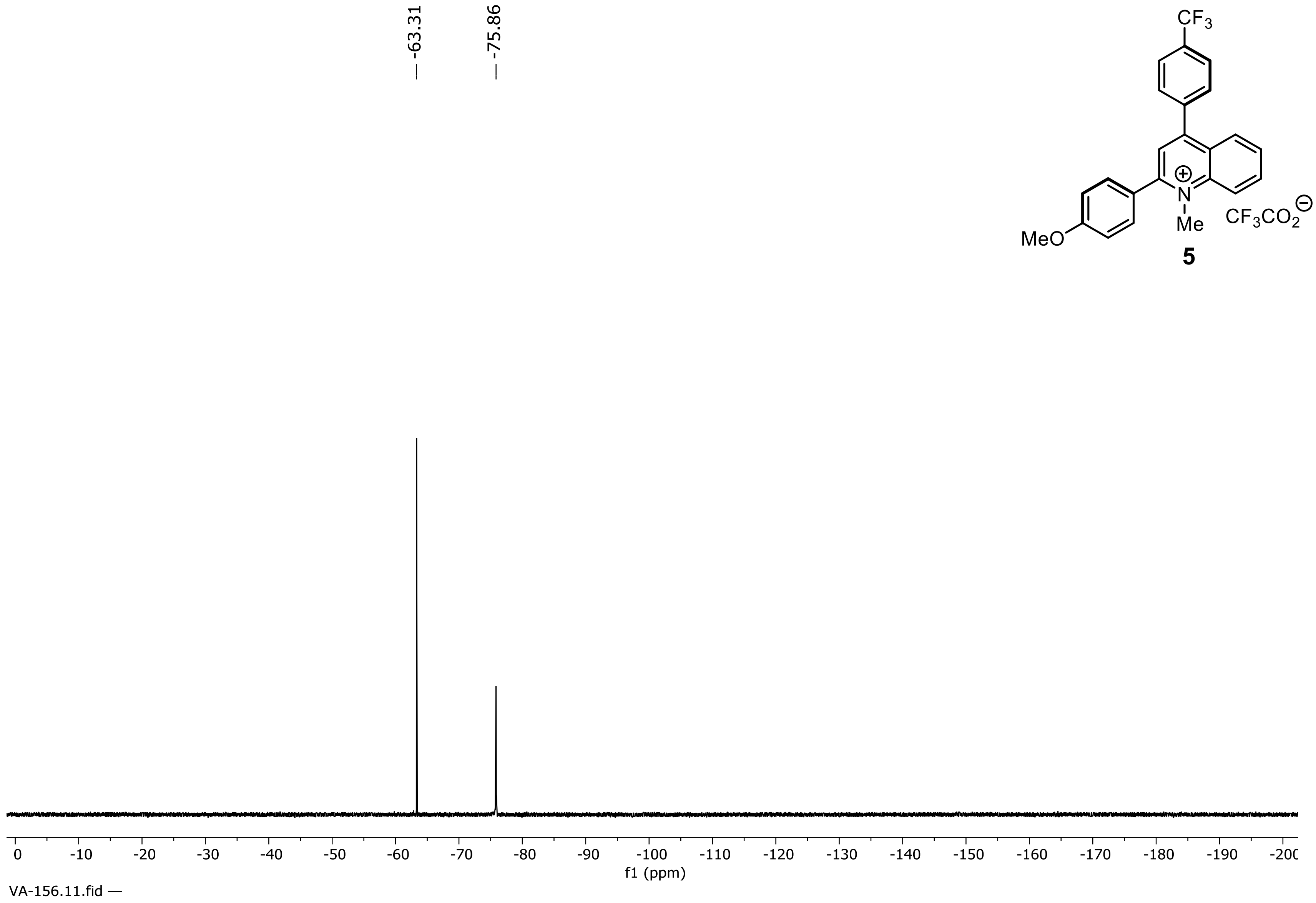


**5**

**$^{19}$F NMR Spectrum of Compound 5 (471 MHz, $CD_3CN$)**

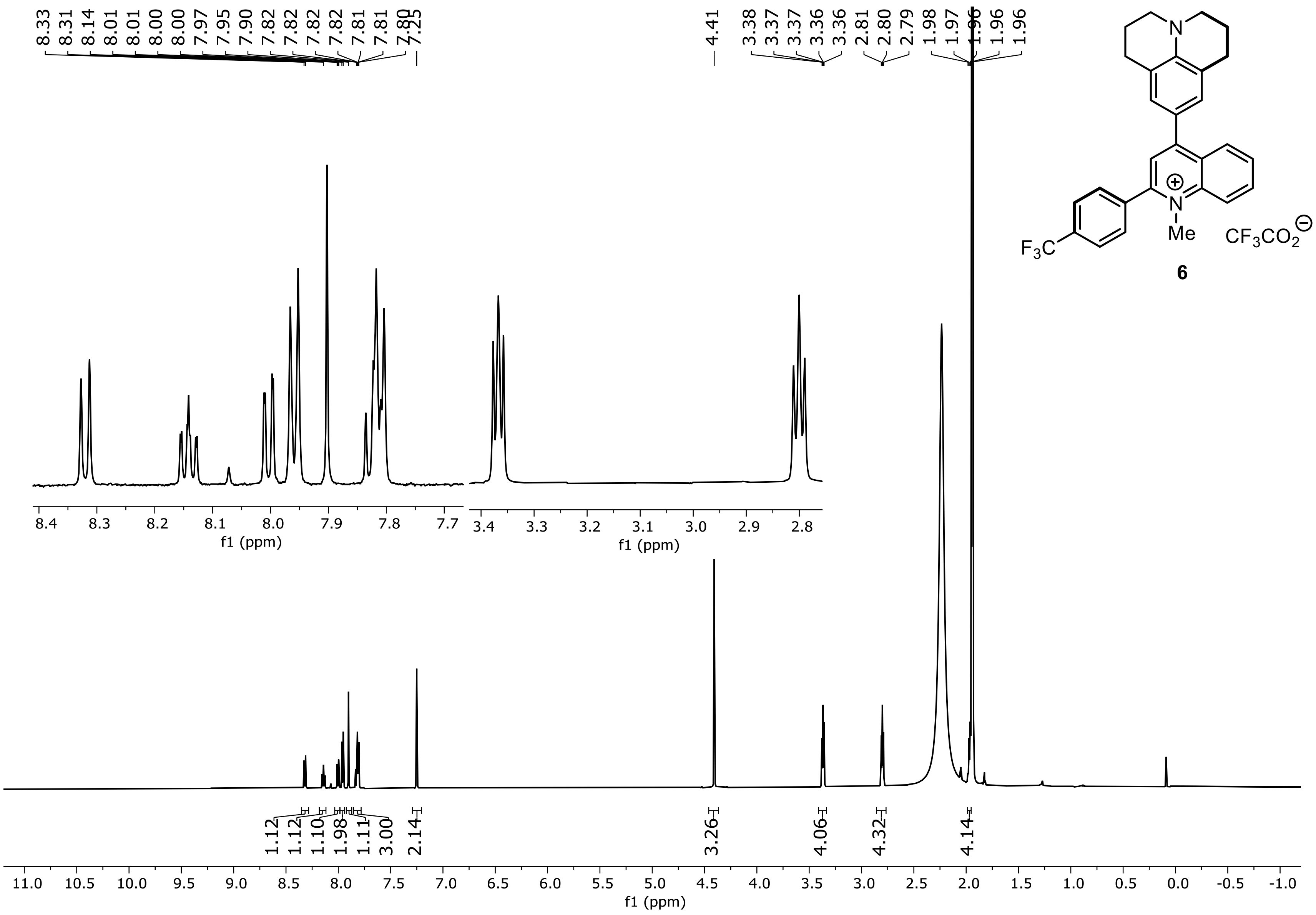


**$^1$H NMR Spectrum of compound 6 (600 MHz, $CD_3CN$)**

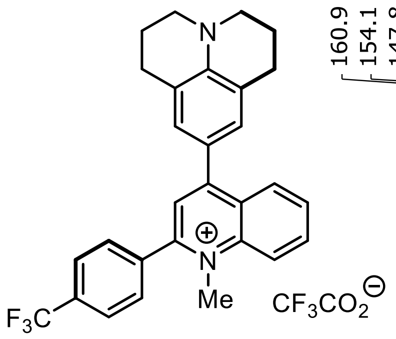


**6**

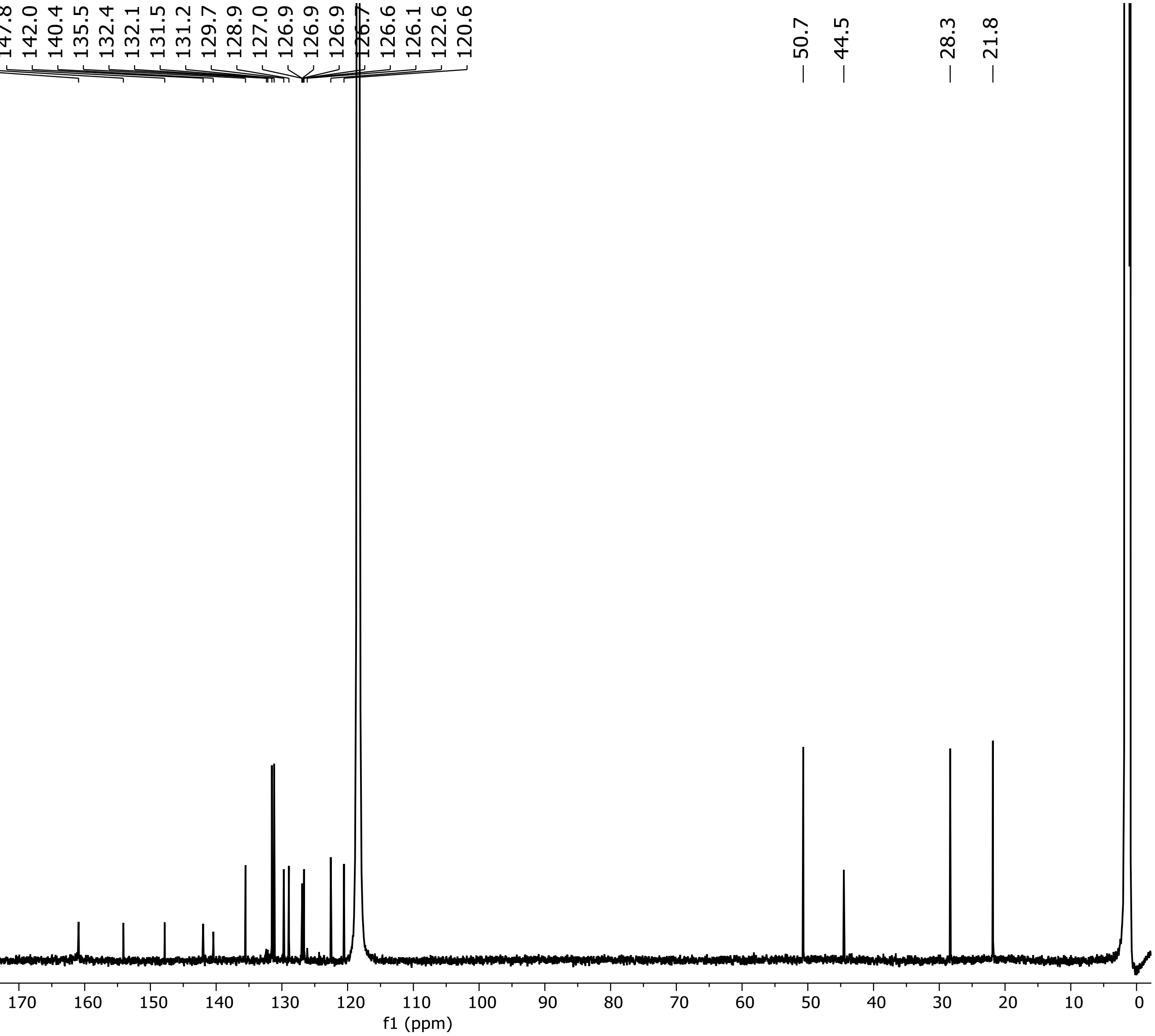


**$^{13}C$ NMR Spectrum of Compound 6 (151 MHz, $CD_3CN$)**

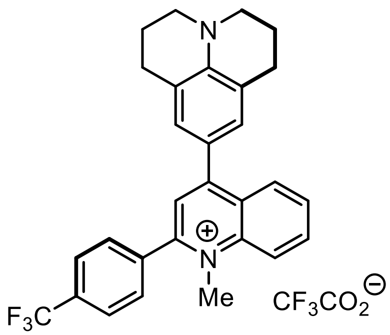

**6**

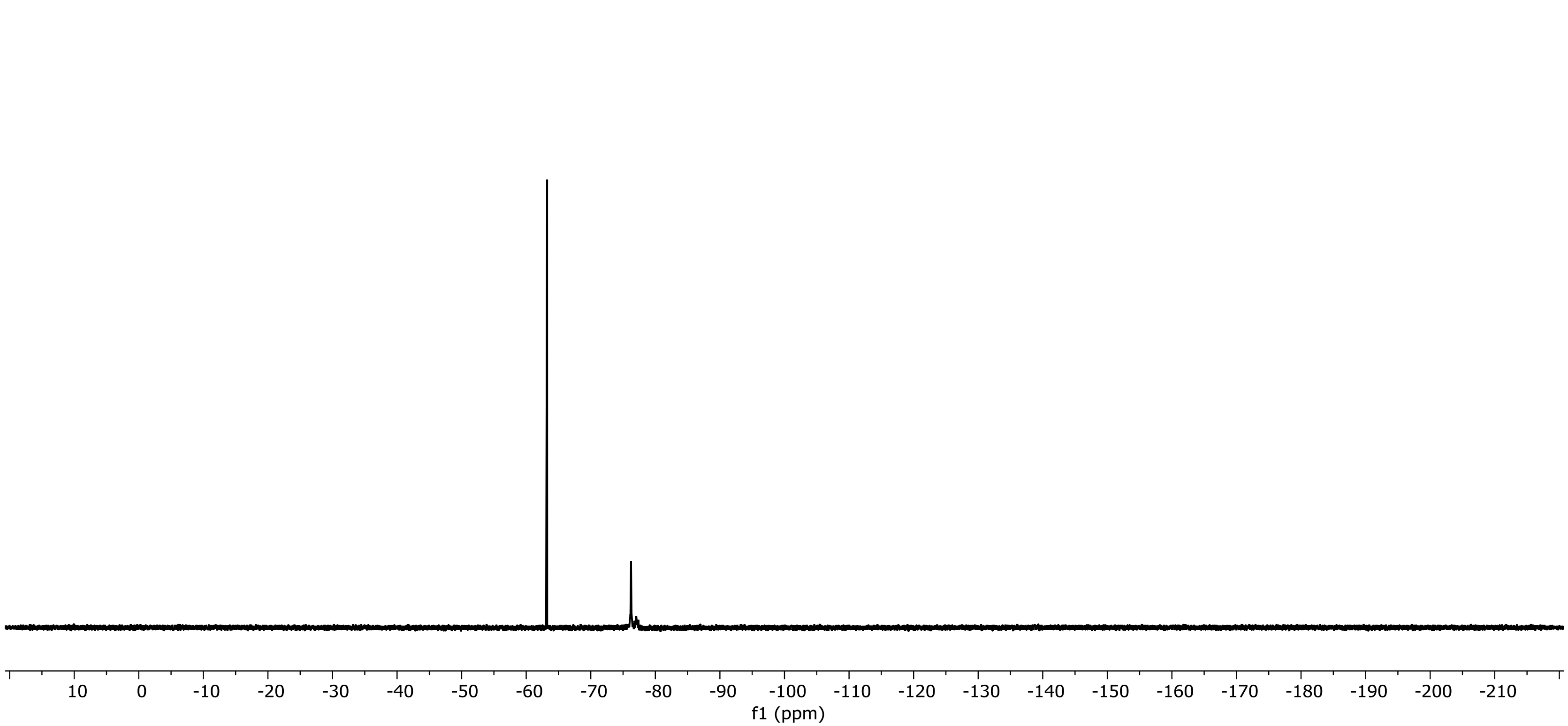


**$^{19}F$ NMR Spectrum of Compound 6 (471 MHz, $CD_3CN$)**

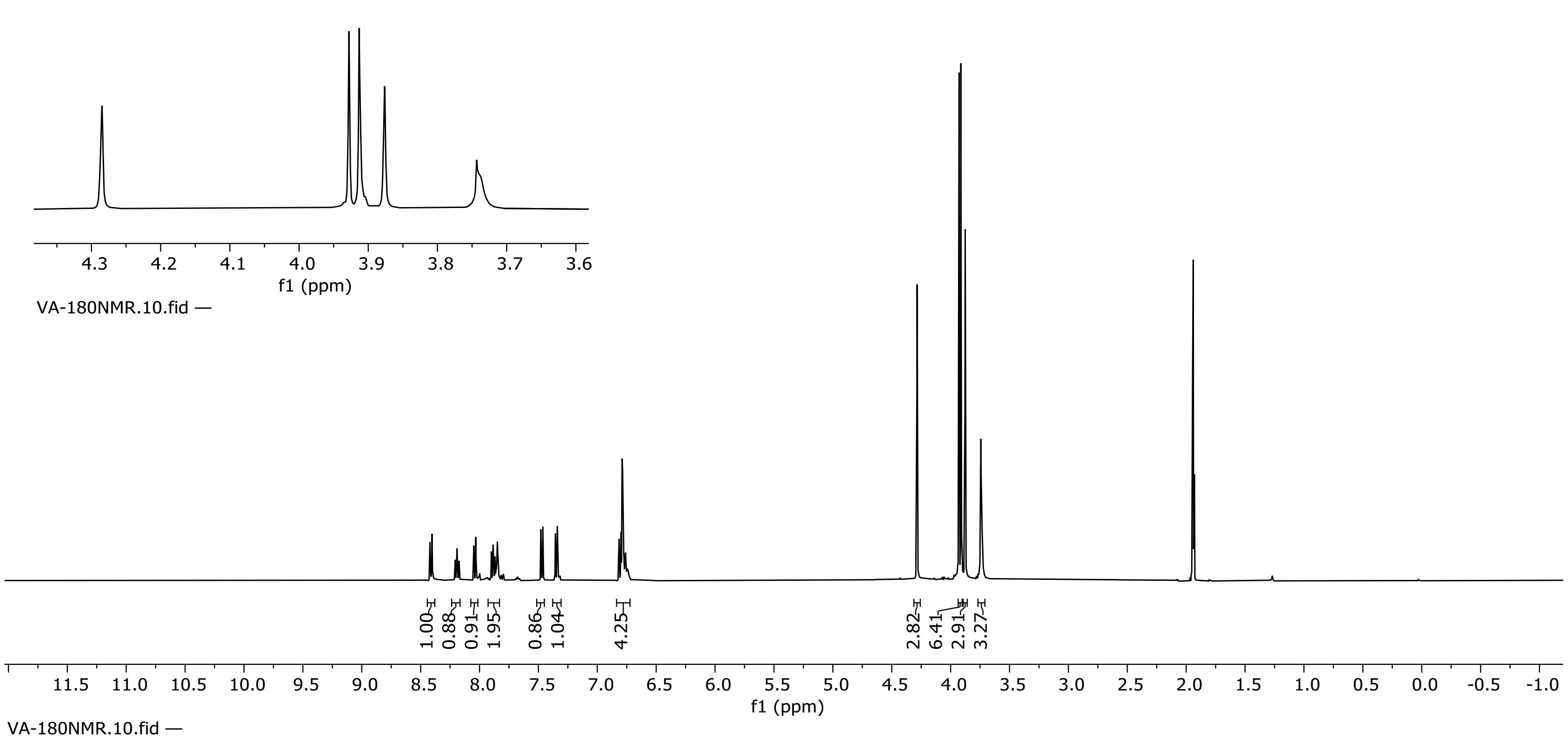

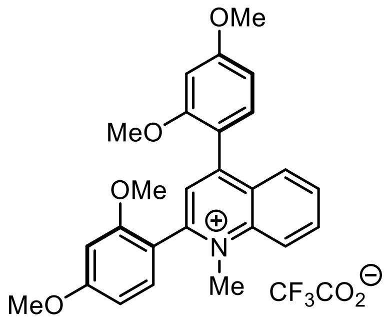

**7**

**[1]H NMR Spectrum of Compound 7 (500 MHz, $CD_3CN$)**

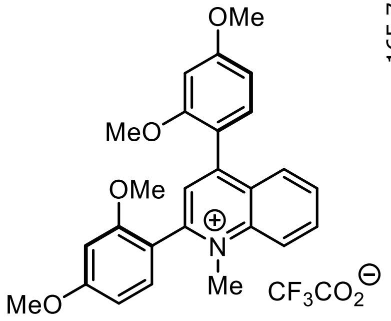

**7**

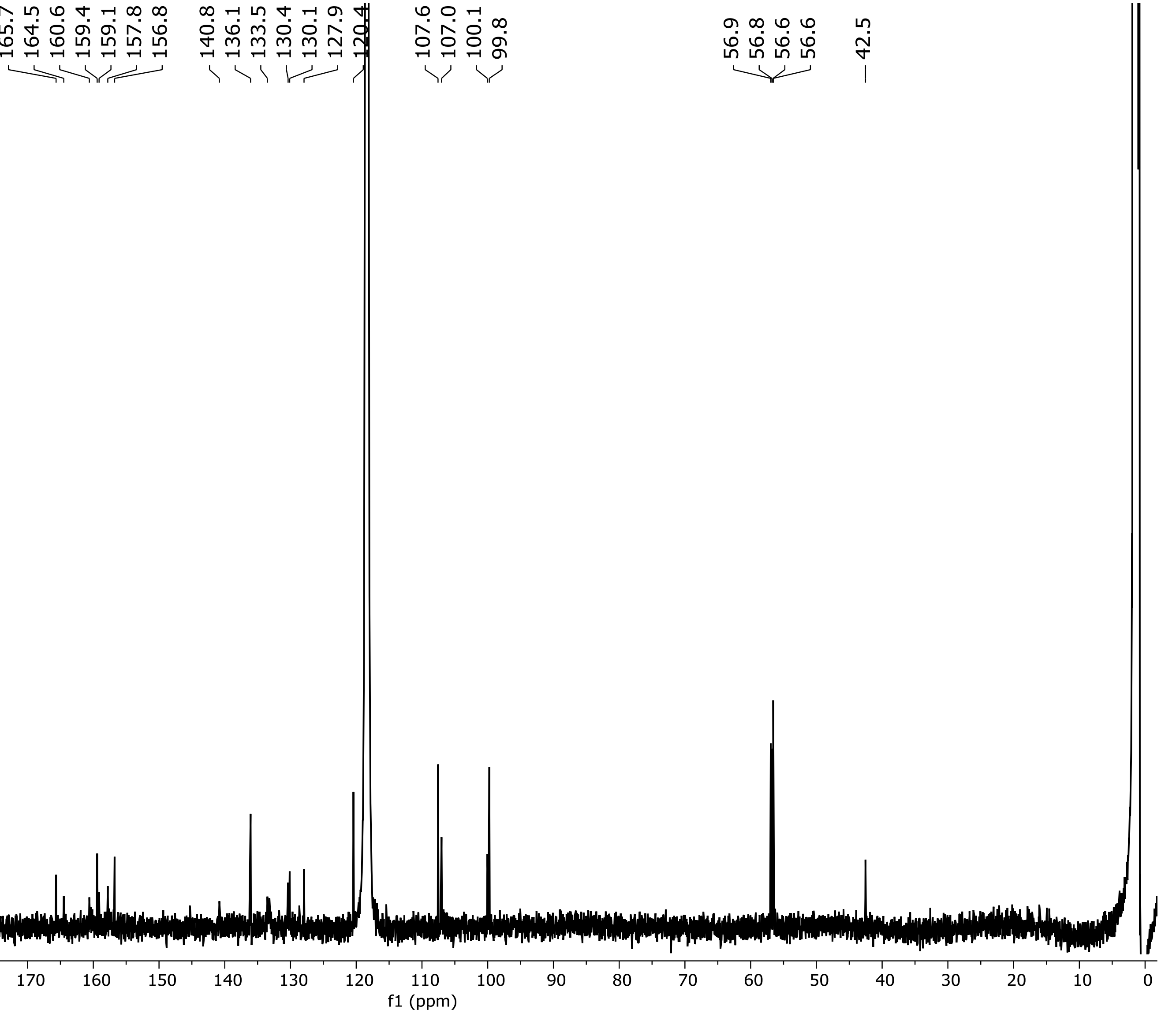


**$^{13}$C NMR Spectrum of Compound 7 (151 MHz, $CD_3CN$)**

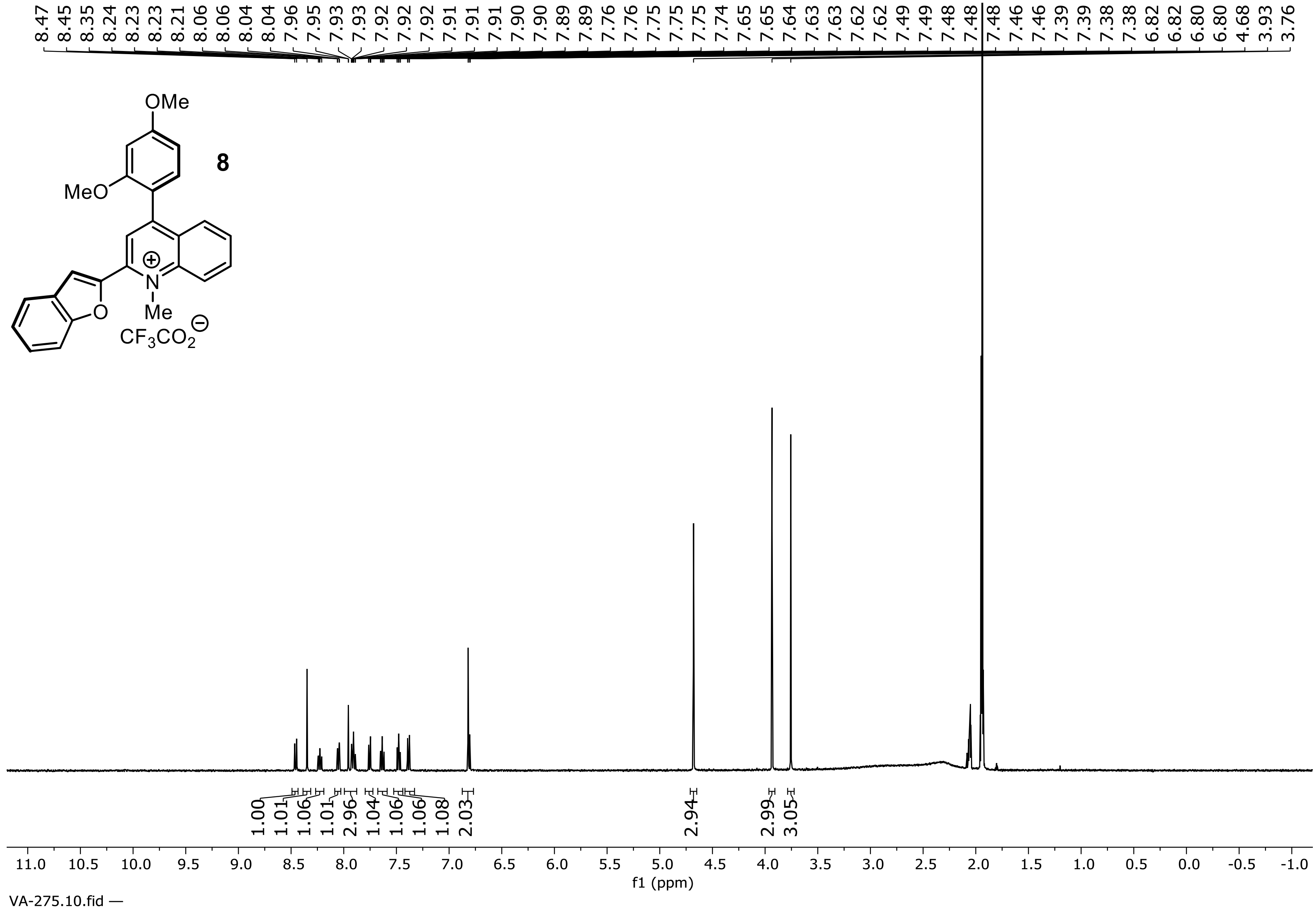


**$^1$H NMR Spectrum of Compound 8 (500 MHz, $CD_3CN$)**

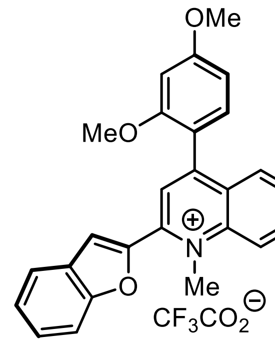


**8**

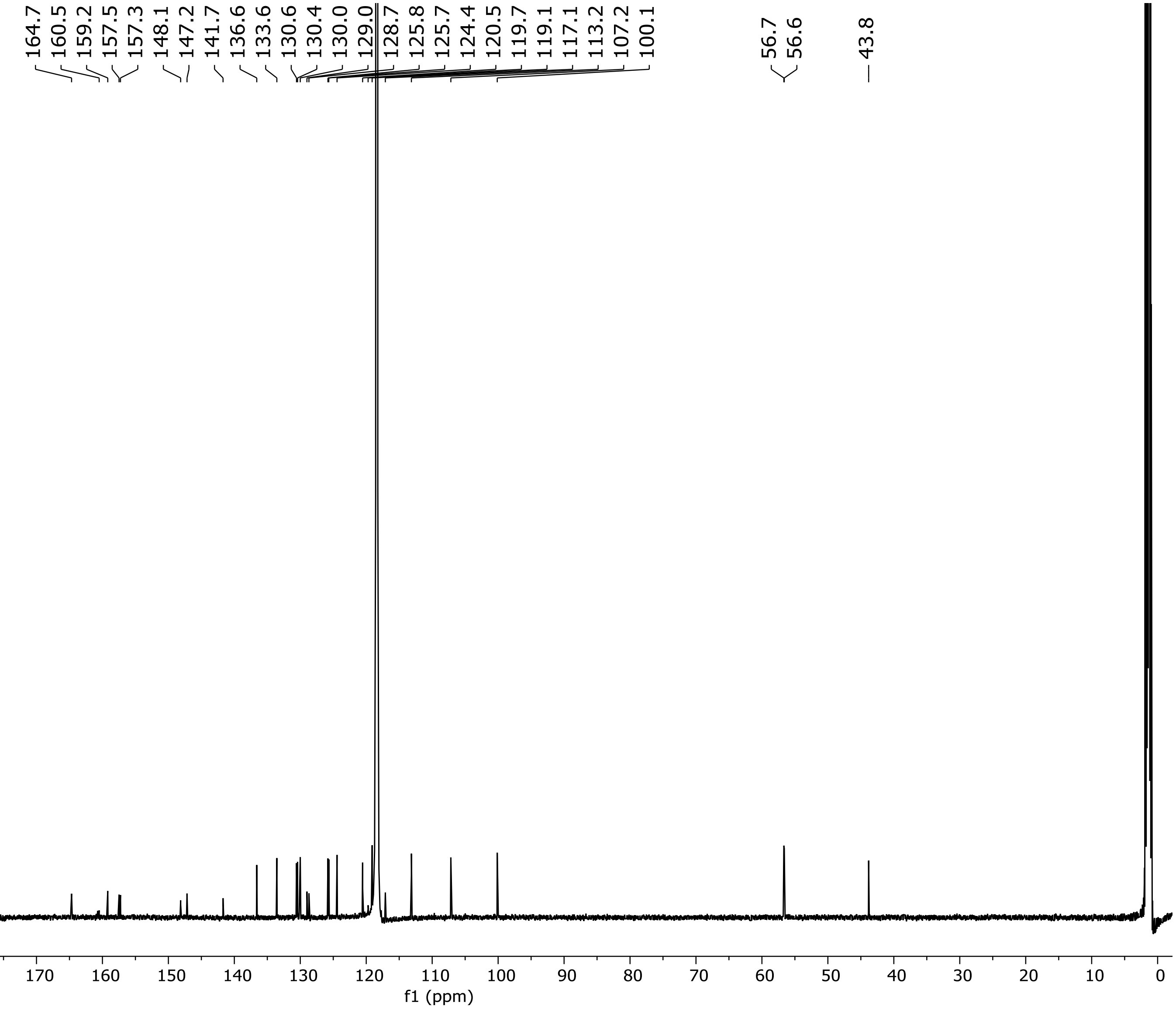


**$^{13}C$ NMR Spectrum of Compound 8 (151 MHz, $CD_3CN$)**

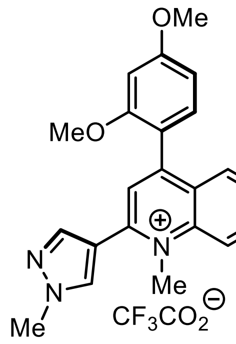

**9**

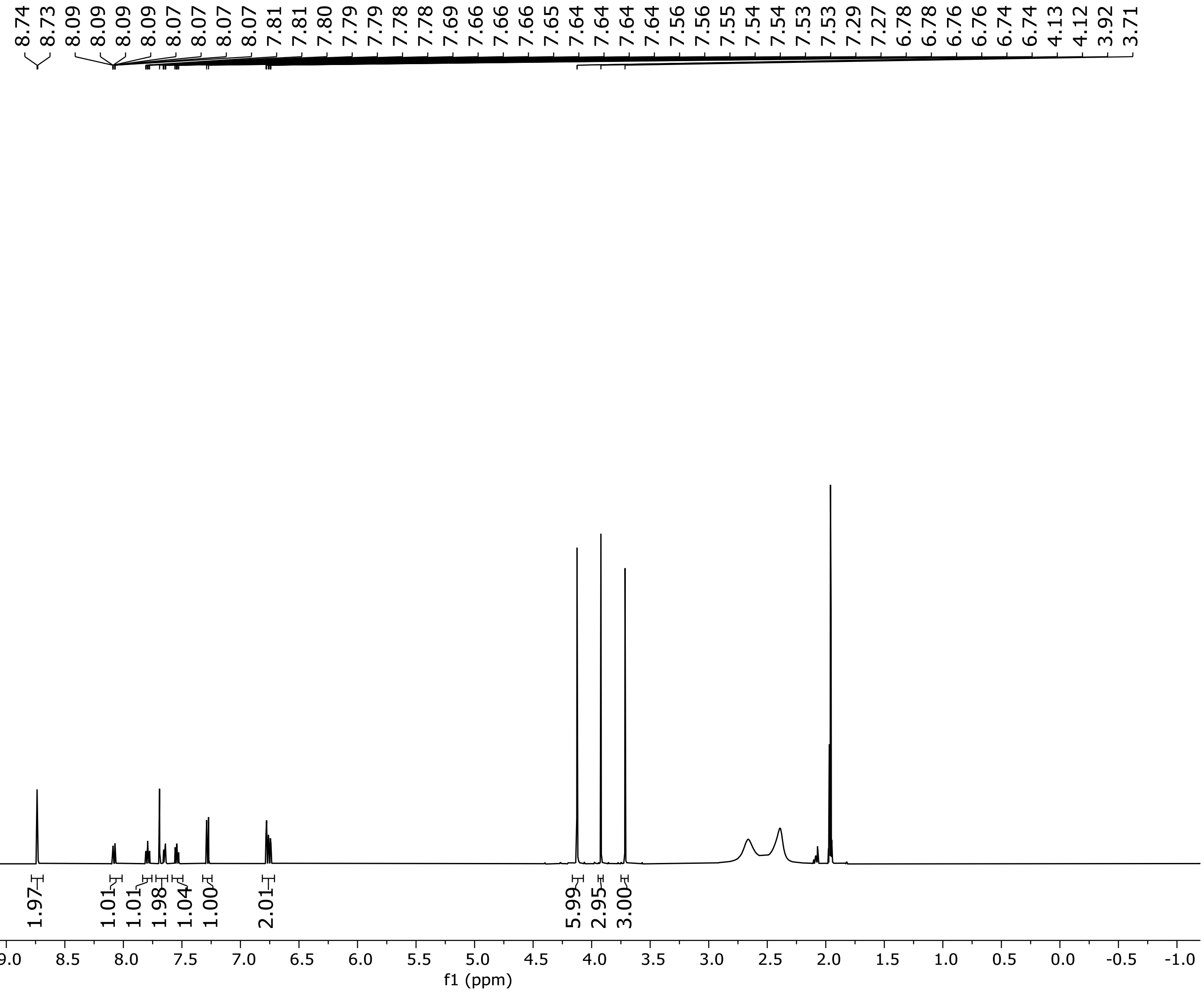


**$^1$H NMR Spectrum of Compound 9 (500 MHz, $CD_3CN$)**

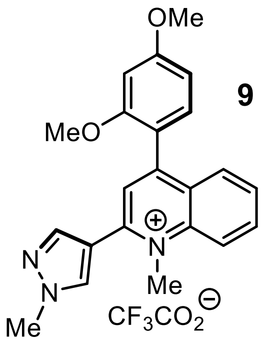


**9**

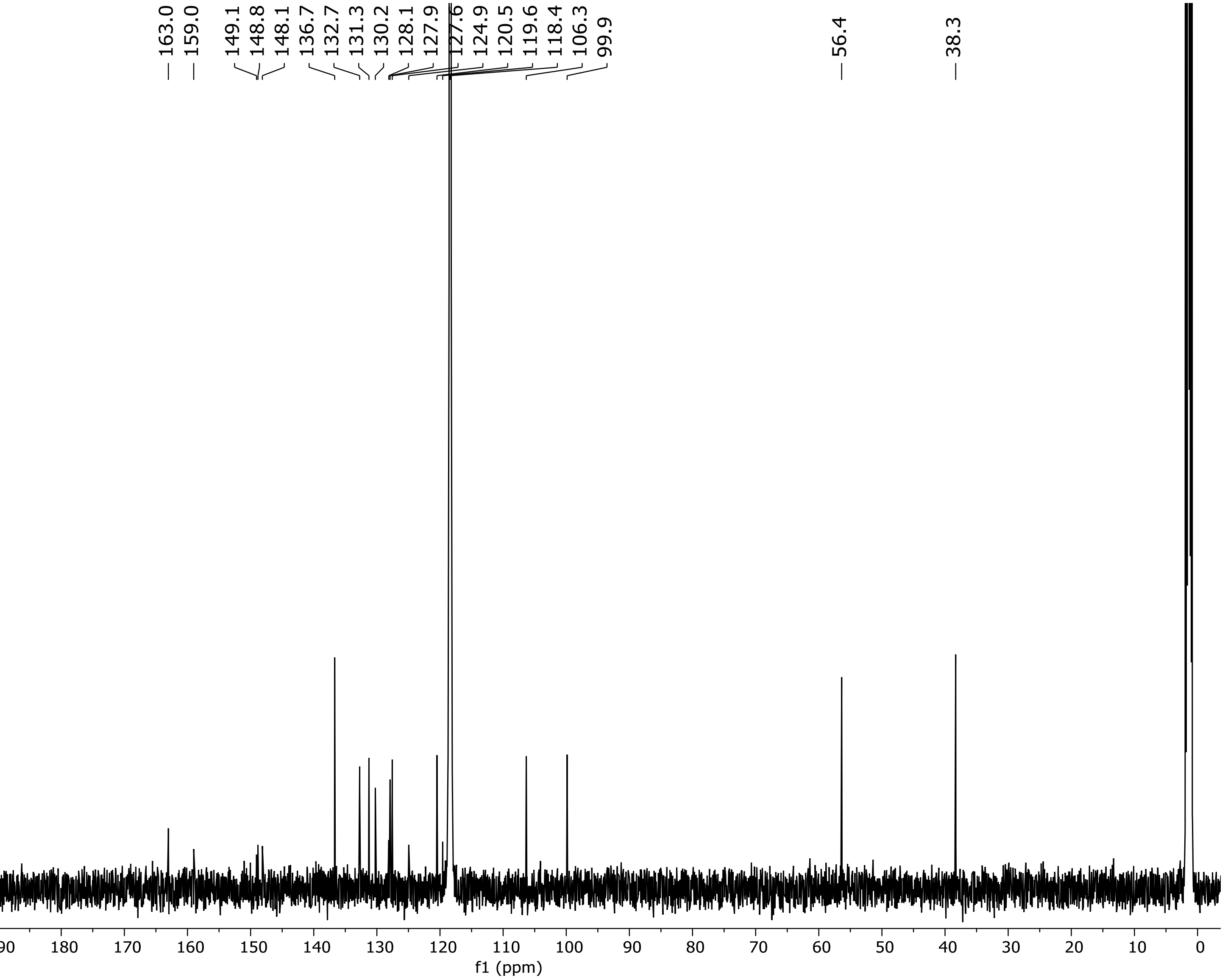


**$^{13}C$ NMR Spectrum of Compound 9 (126 MHz, $CD_3CN$)**

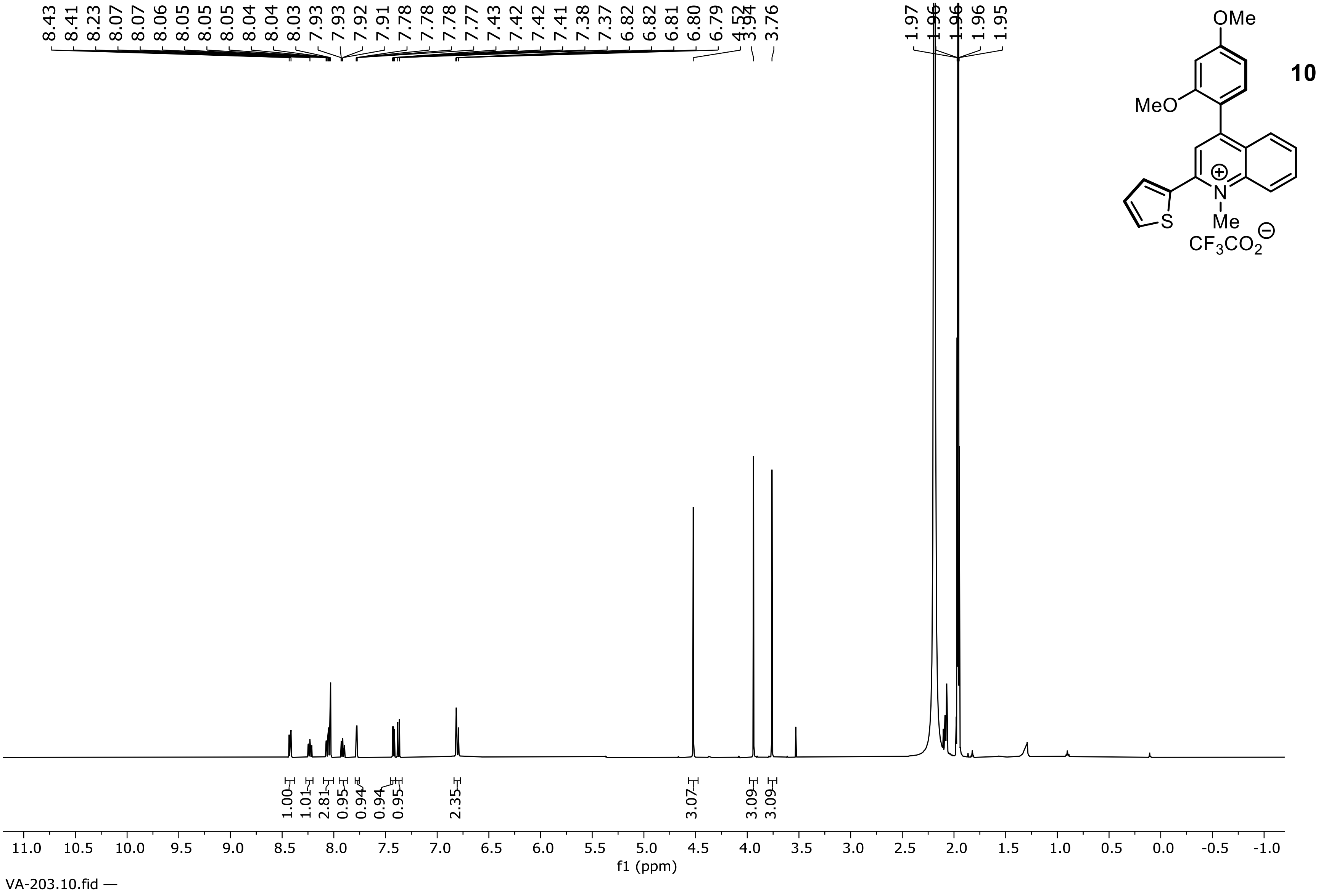

**$^1$H NMR Spectrum of Compound 10 (500 MHz, $CD_3CN$)**

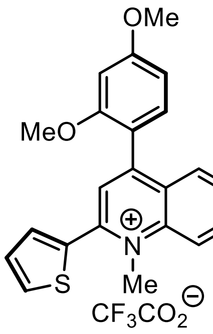


**10**

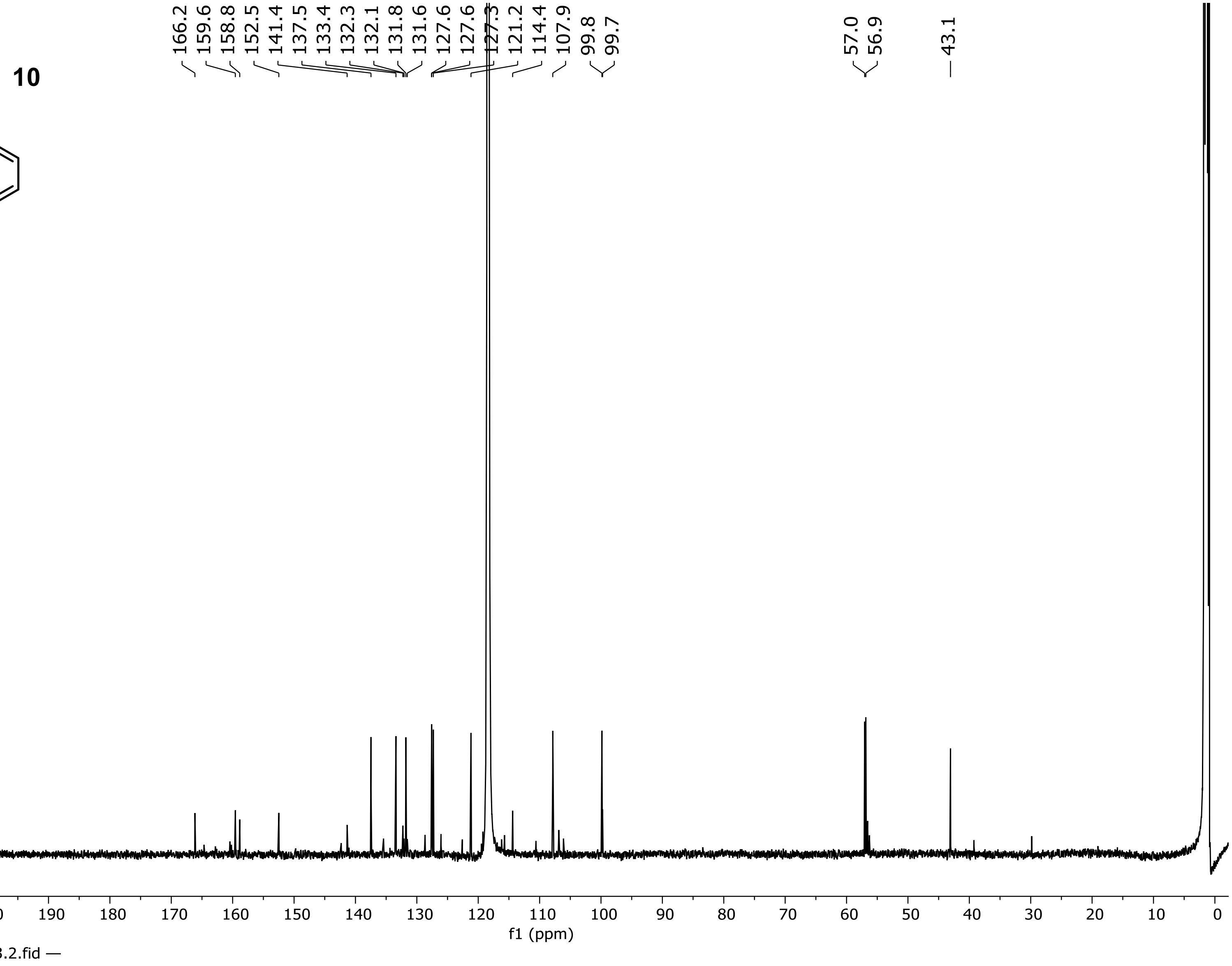


**$^{13}C$ NMR Spectrum of Compound 10 (151 MHz, $CD_3CN$)**

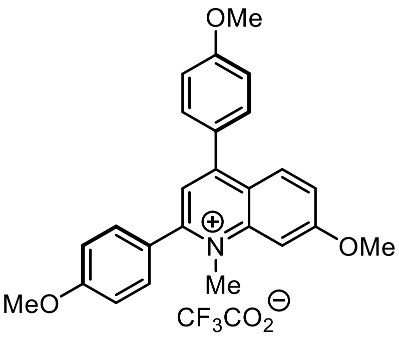


11

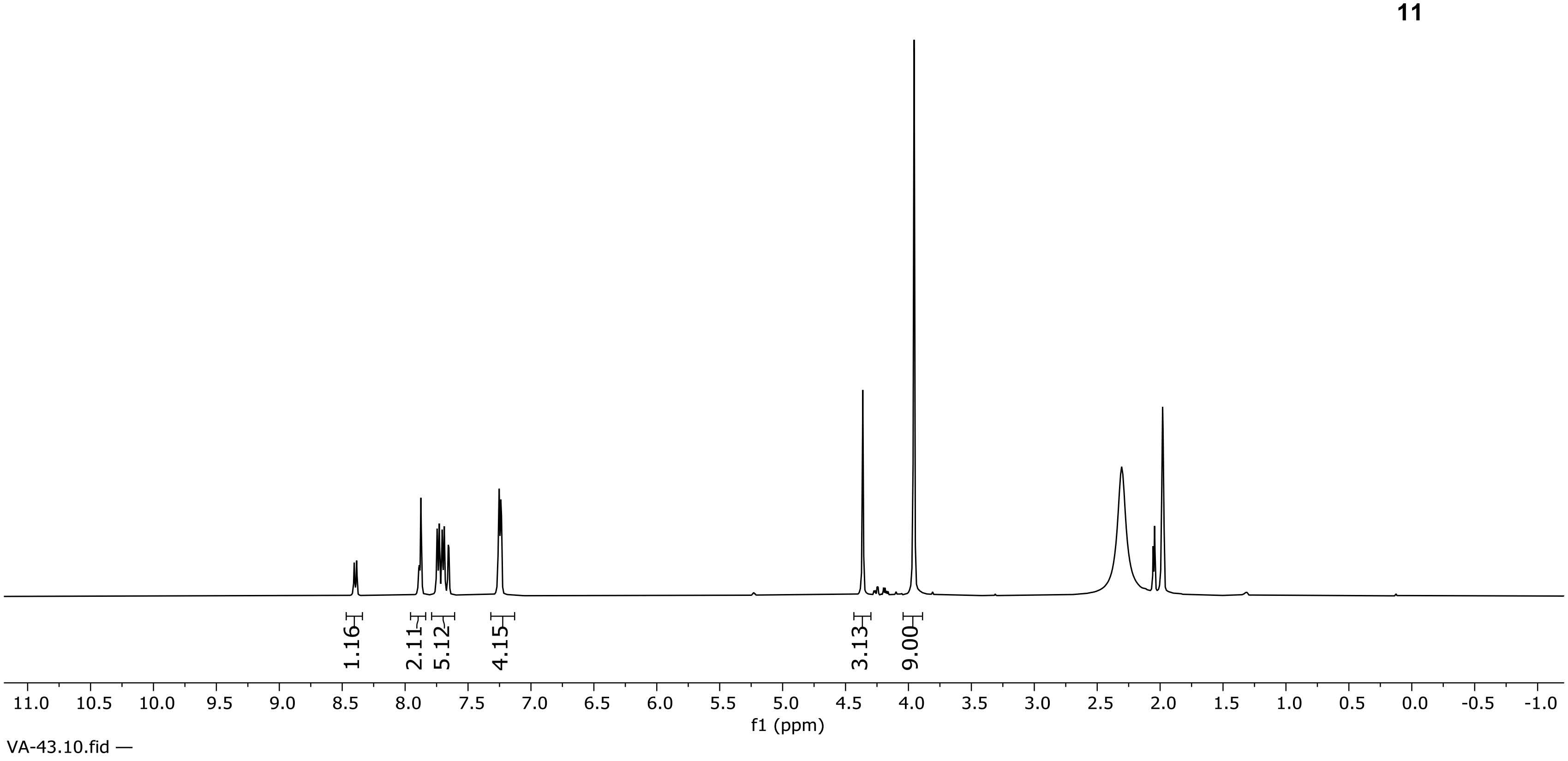


**$^1$H NMR Spectrum of Compound 11 (500 MHz, $CD_3CN$)**

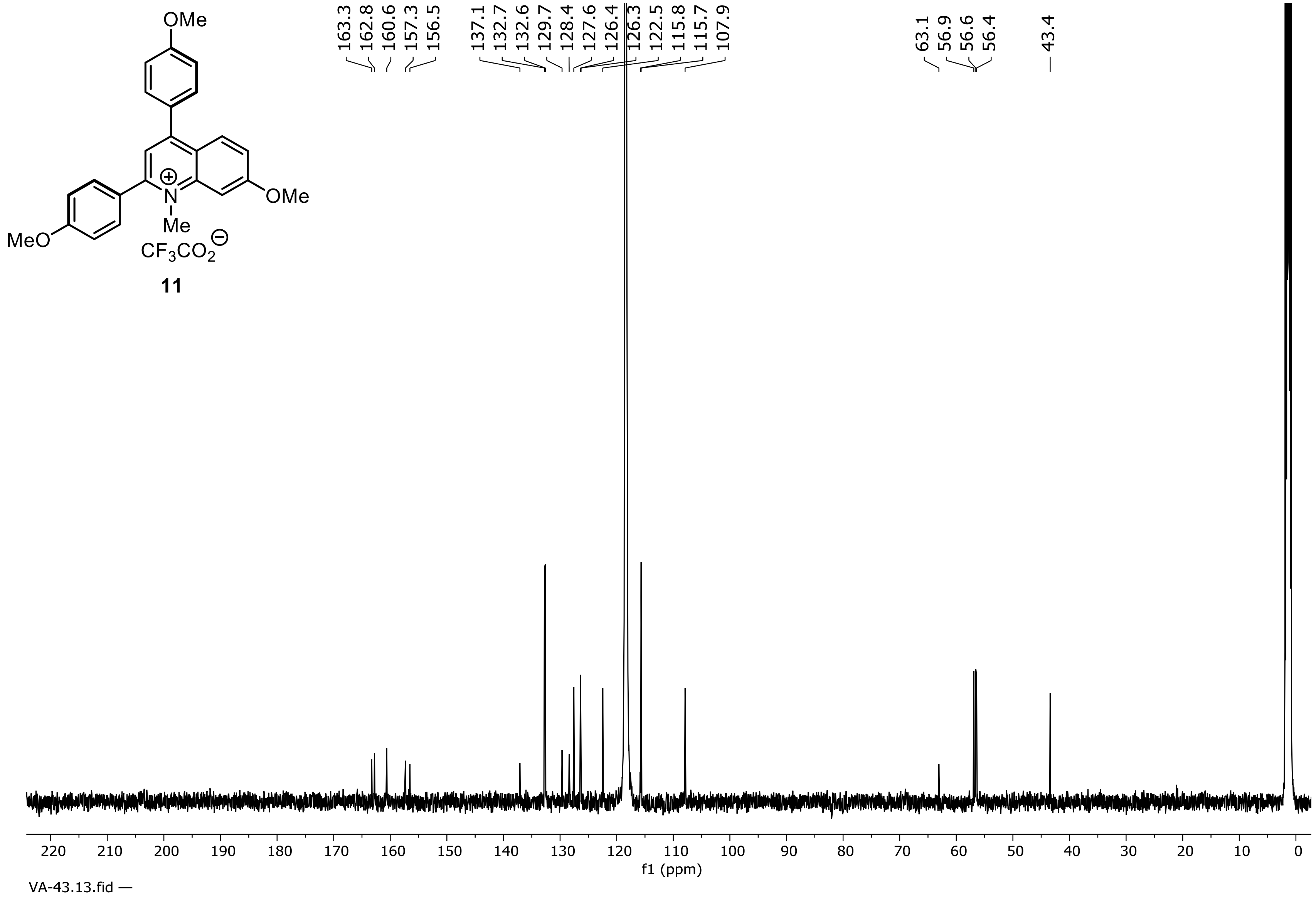


**$^{13}C$ NMR Spectrum of Compound 11 (126 MHz, $CD_3CN$)**

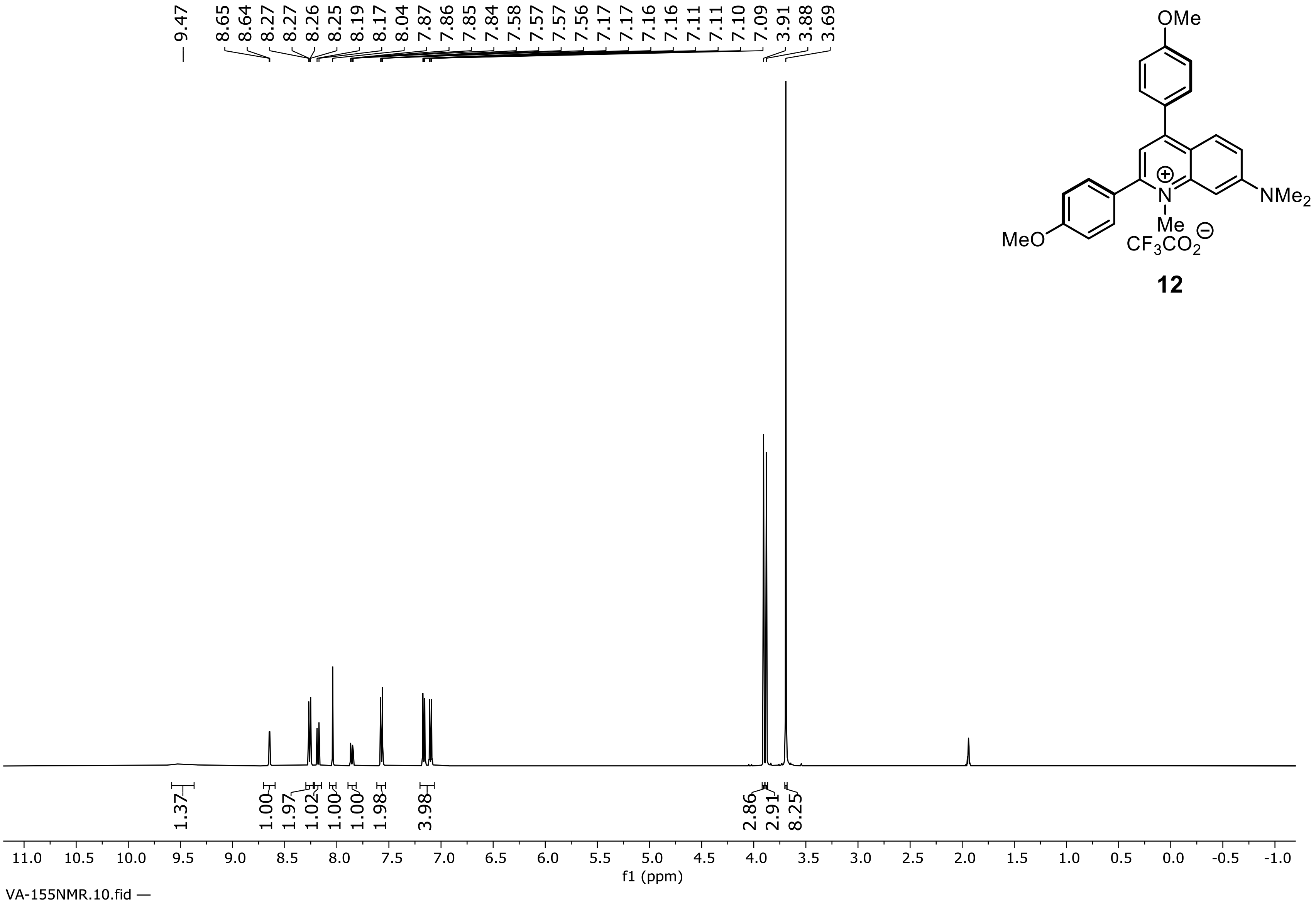


**12**

**¹H NMR Spectrum of Compound 12 (500 MHz, $CD_3CN$)**

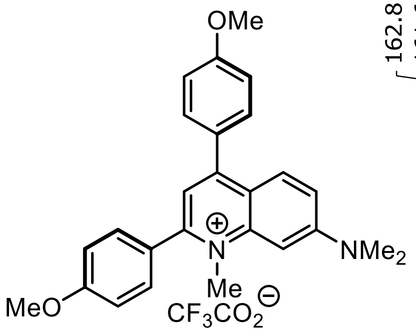


12

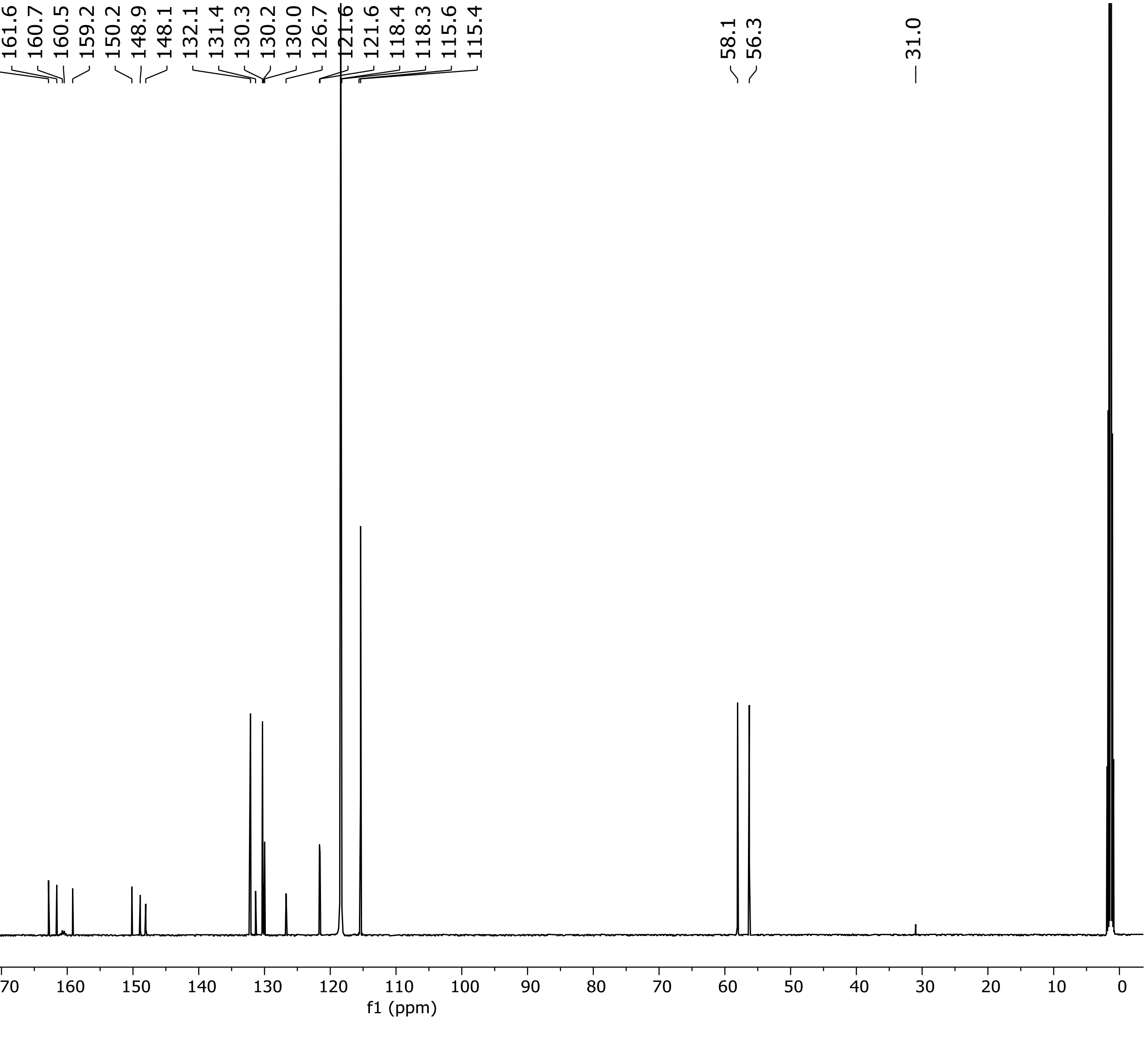


**$^{13}C$ NMR Spectrum of Compound 12 (126 MHz, $CD_3CN$)**

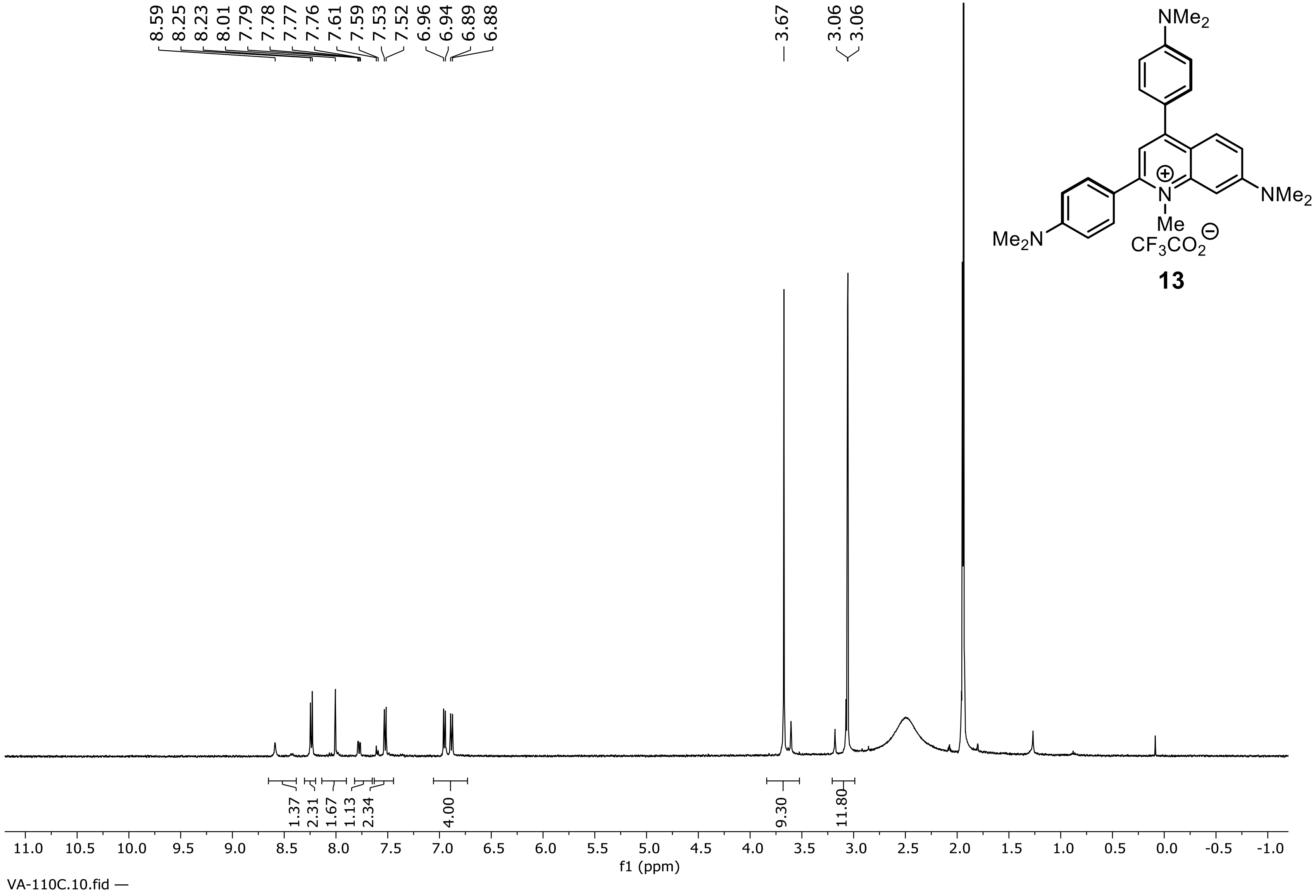


**$^1H$ NMR Spectrum of Compound 13 (500 MHz, $CD_3CN$)**

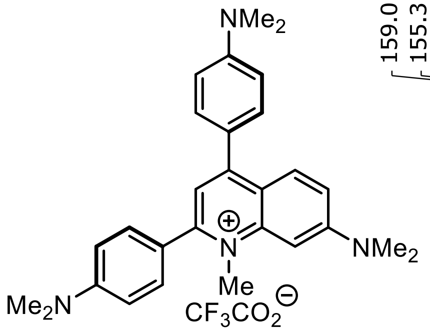


**13**

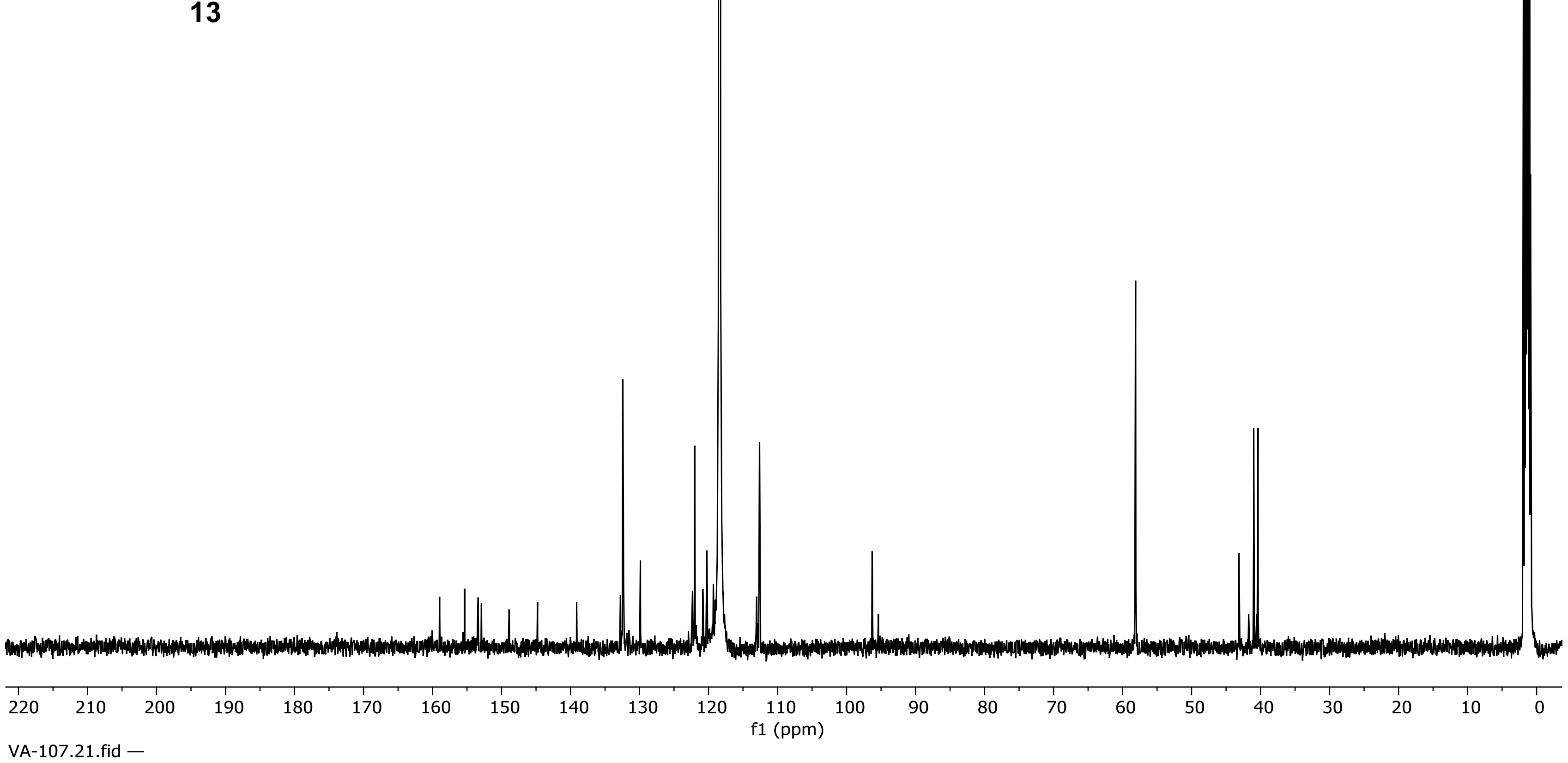


**$^{13}C$ NMR Spectrum of Compound 13 (151 MHz, $CD_3CN$)**

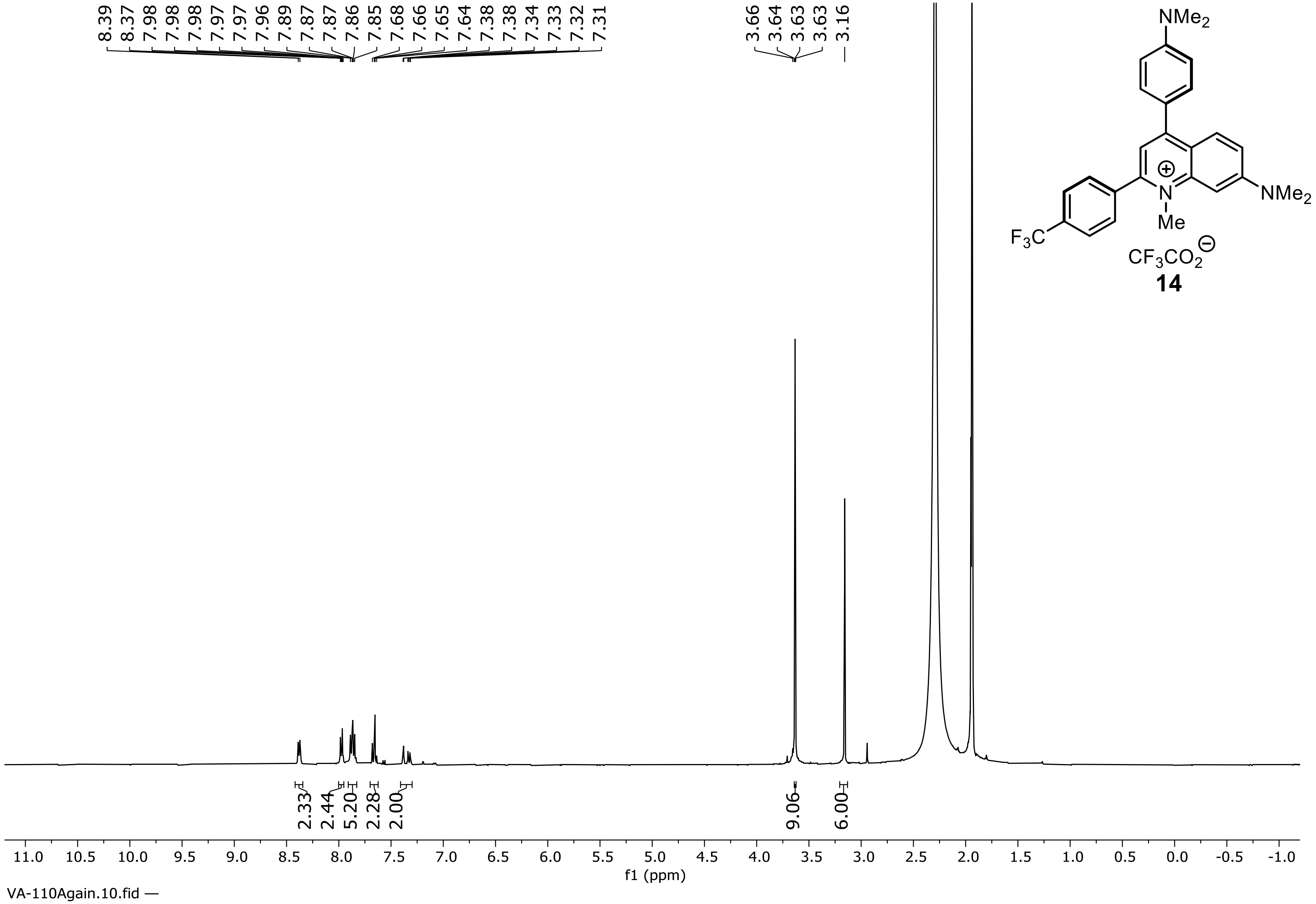


**$^1$H NMR Spectrum of Compound 14 (500 MHz, $CD_3CN$)**

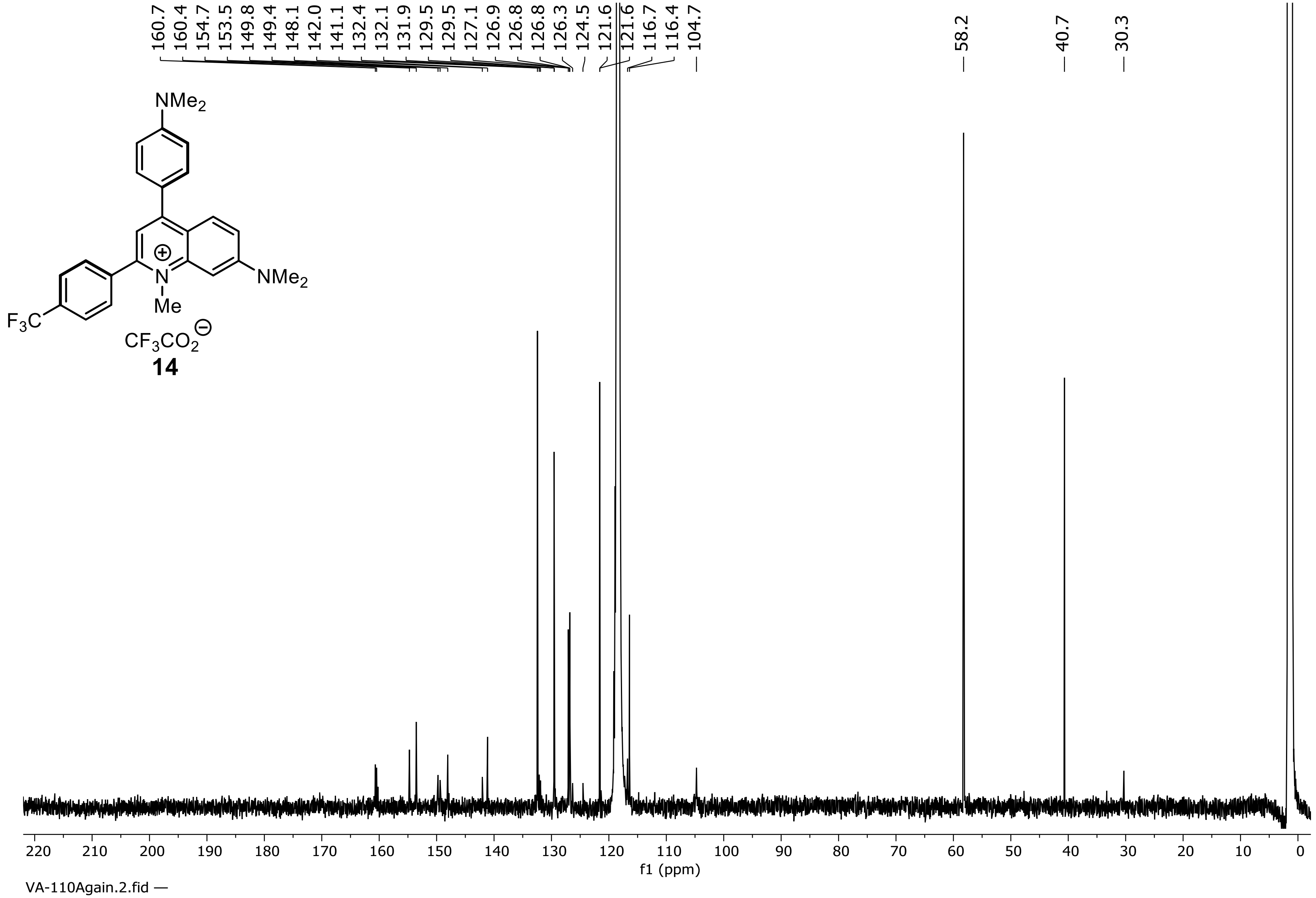


**$^{13}$C NMR Spectrum of Compound 14 (151 MHz, $CD_3CN$)**

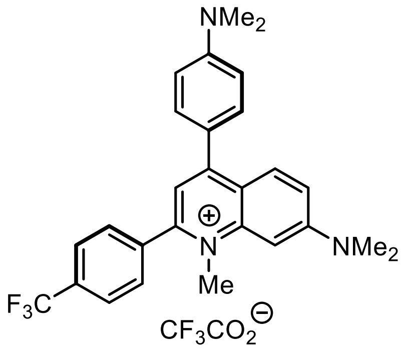


**14**

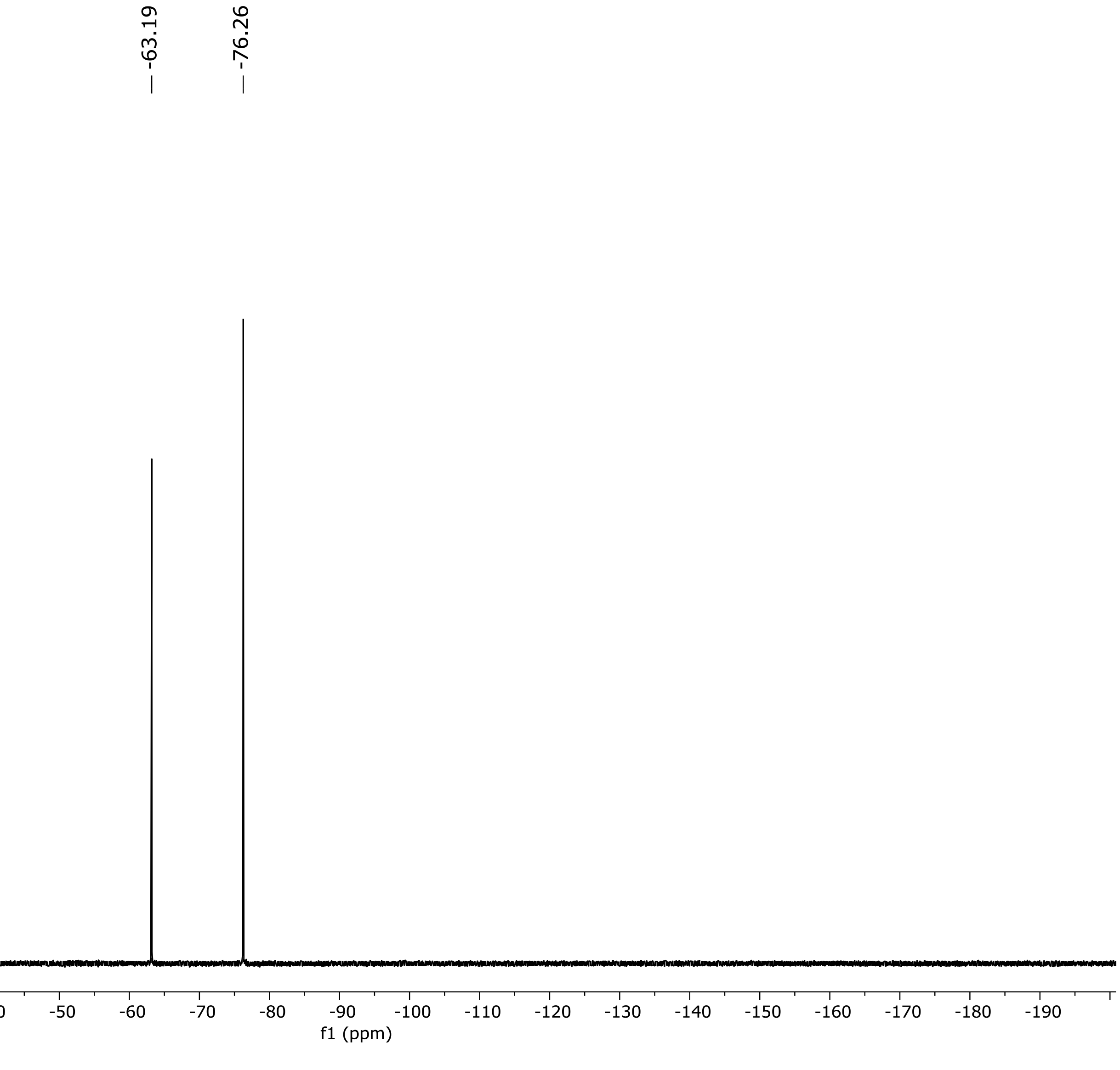


**$^{19}$F NMR Spectrum of Compound 14 (471 MHz, $CD_3CN$)**

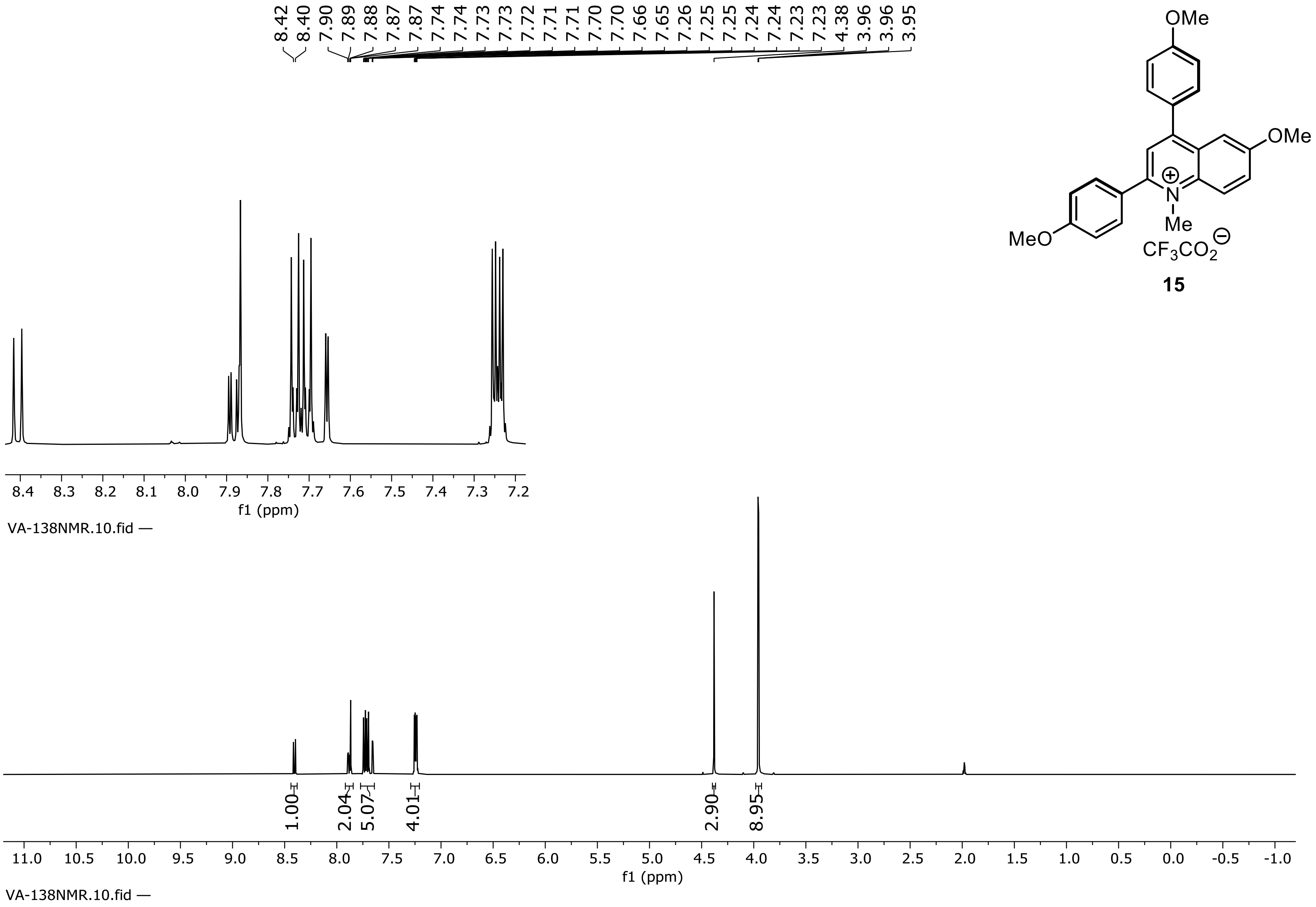


**15**

**[1]H NMR Spectrum of Compound 15 (500 MHz, $CD_3CN$)**

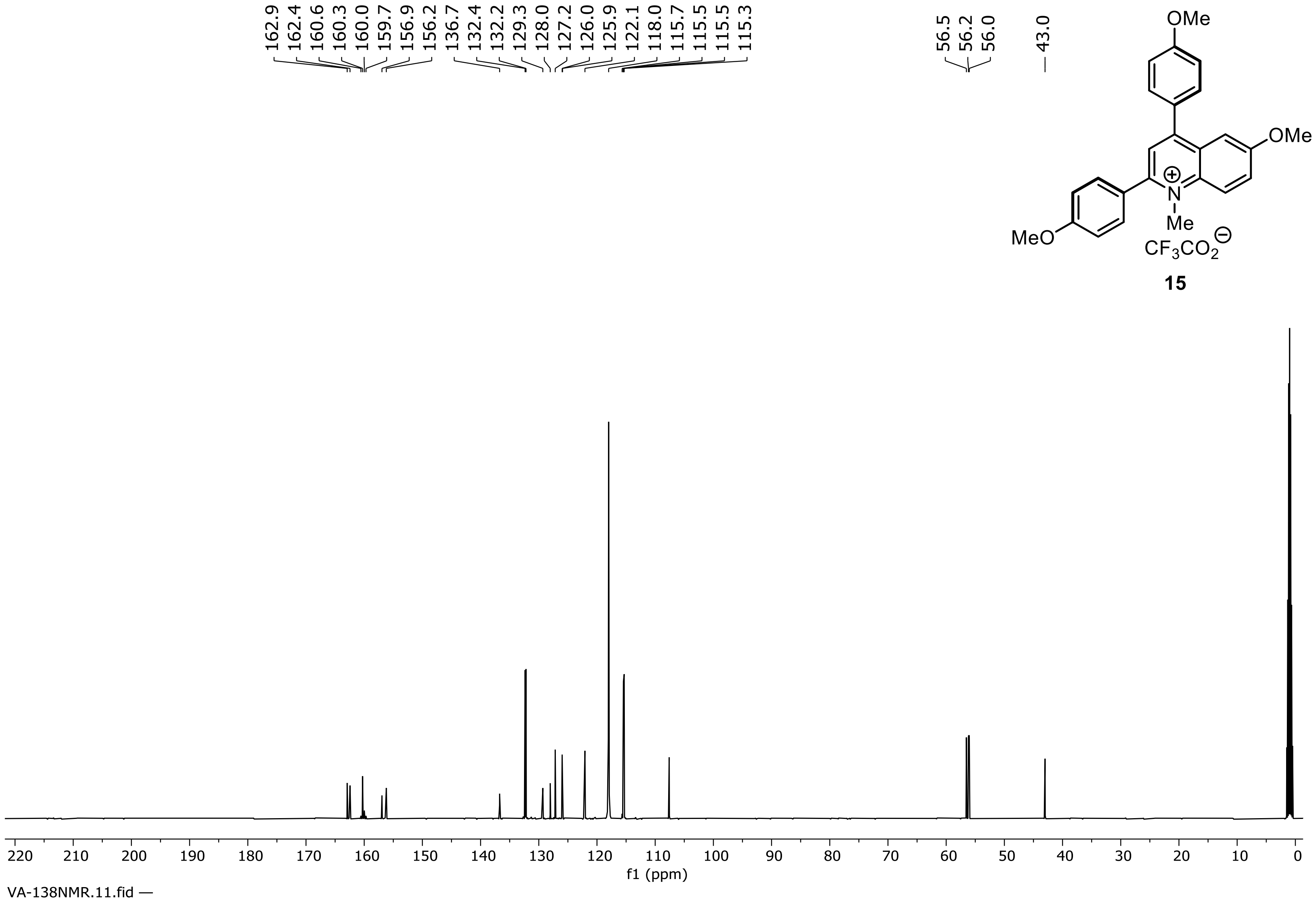


**$^{13}$C NMR Spectrum of Compound 15 (126 MHz, $CD_3CN$)**

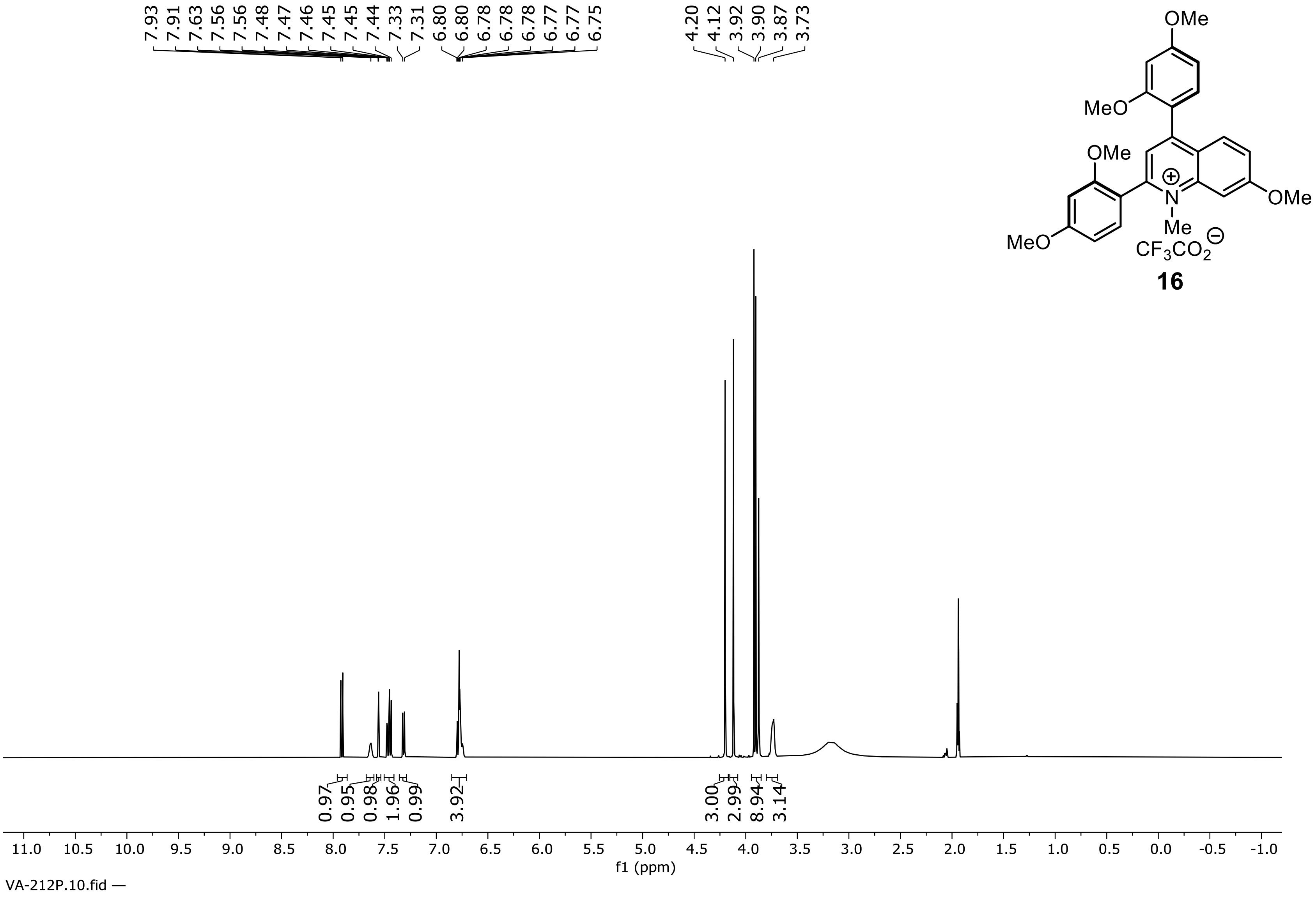


**$^1$H NMR Spectrum of Compound 16 (500 MHz, $CD_3CN$)**

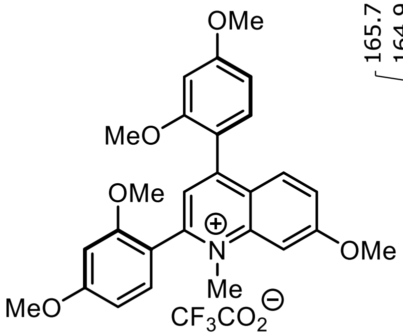

**16**

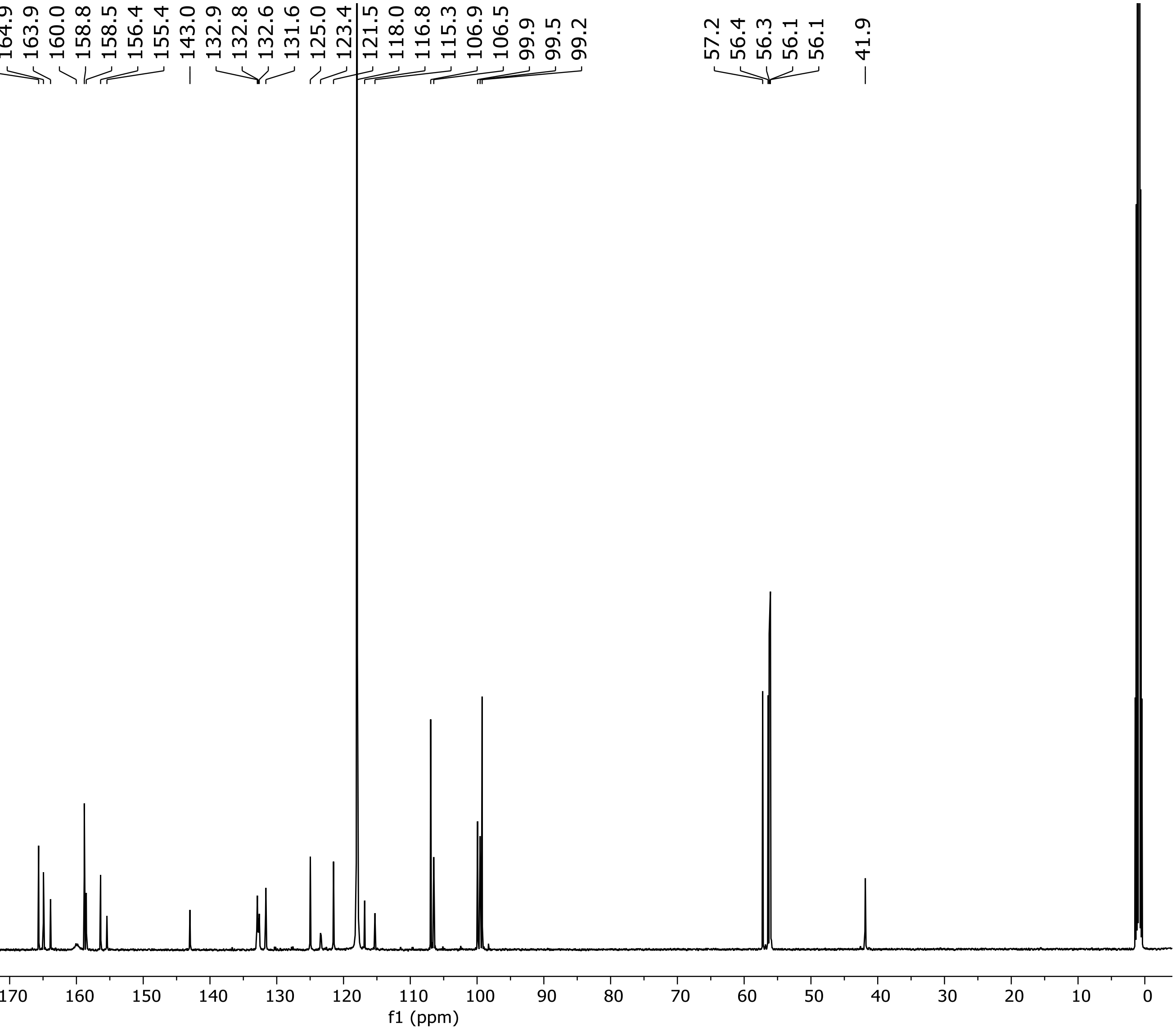


**$^{13}$C NMR Spectrum of Compound 16 (126 MHz, $CD_3CN$)**

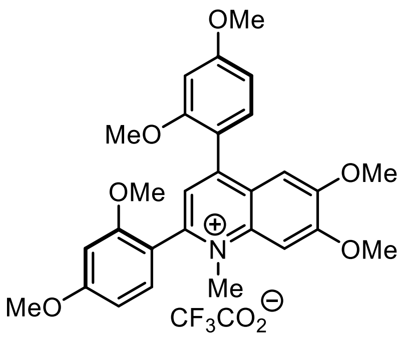


**17**

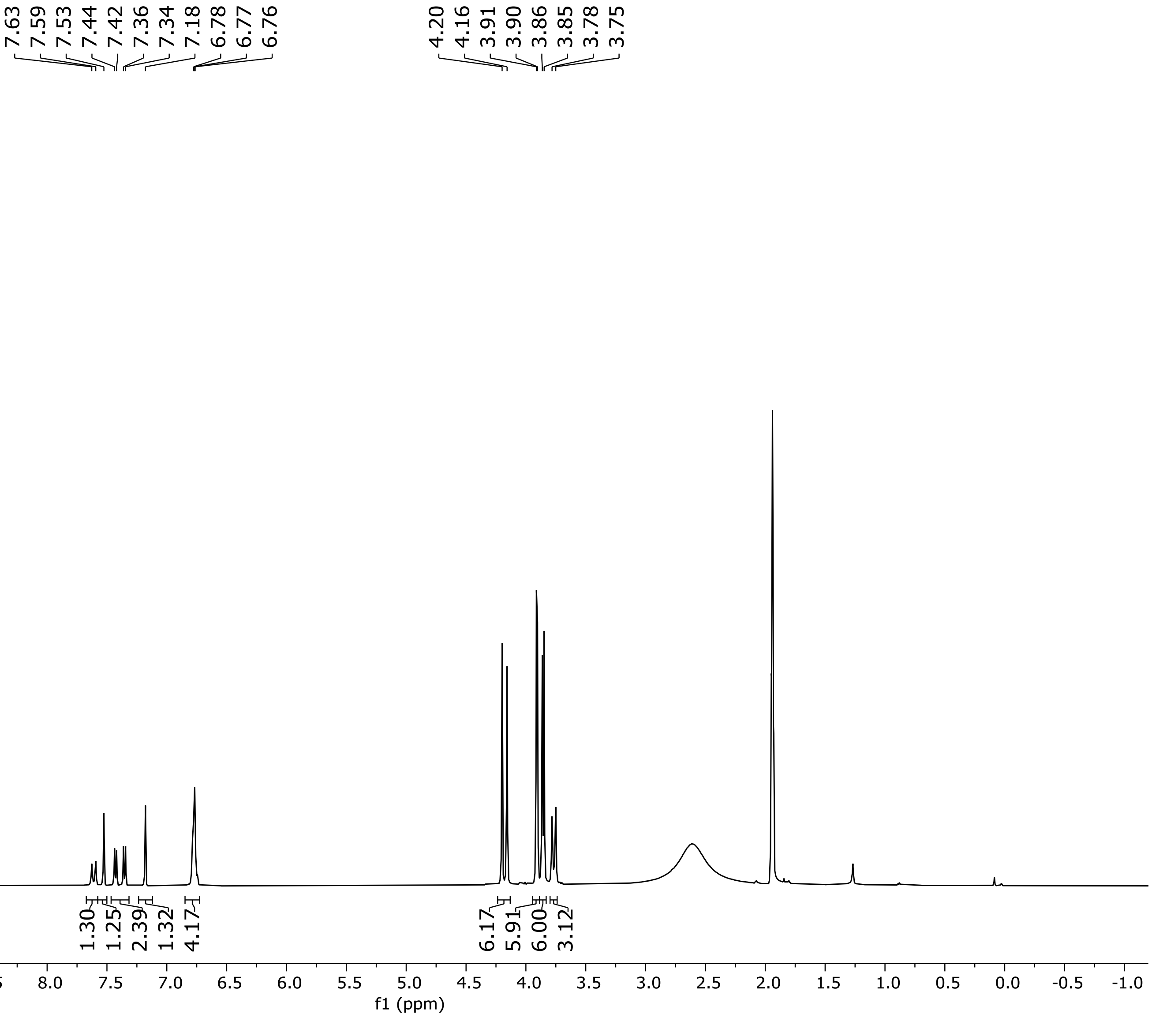


**$^1H$ NMR Spectrum of Compound 17 (500 MHz, $CD_3CN$)**

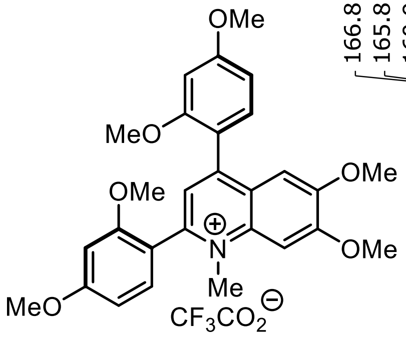

17

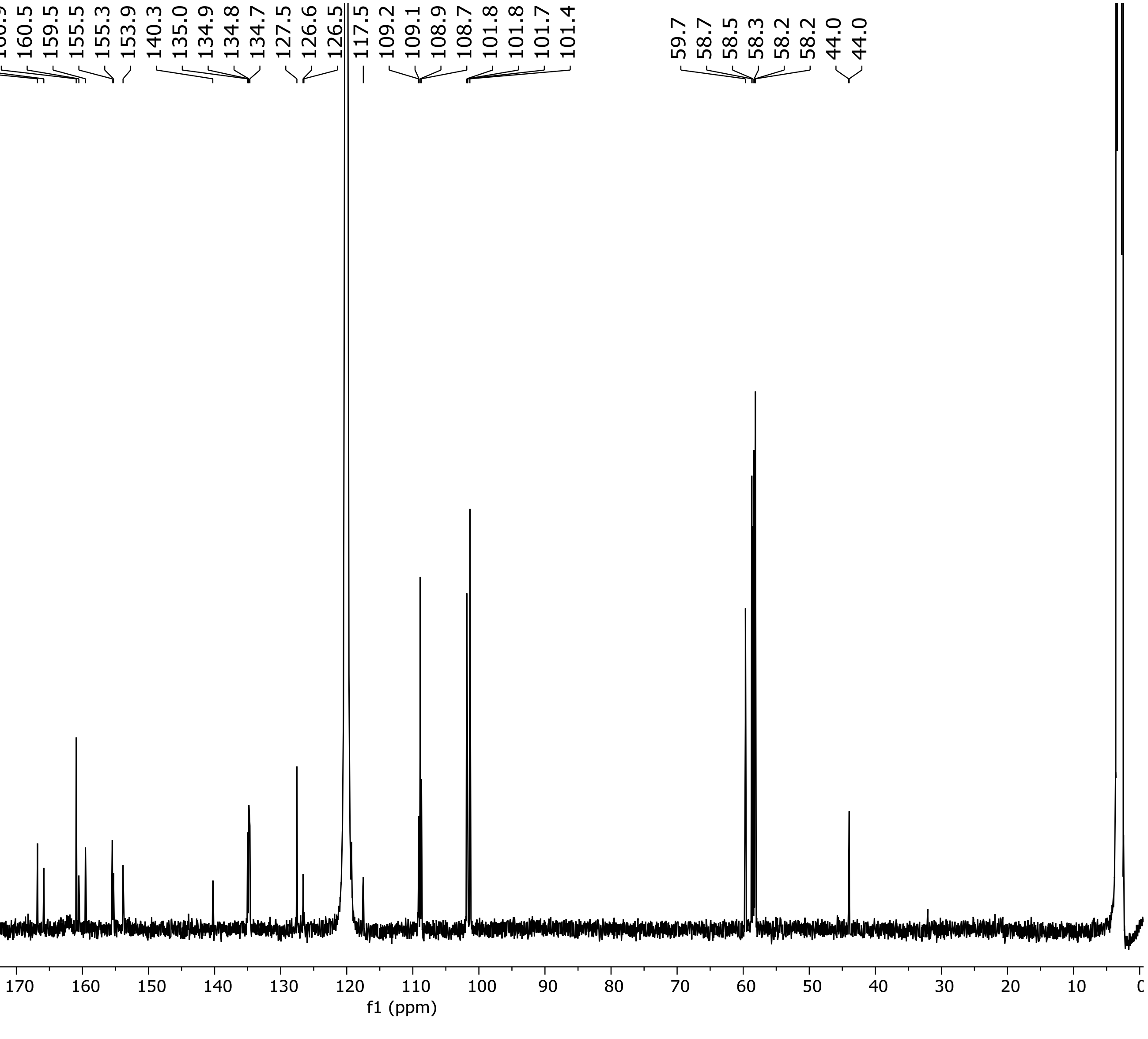


**$^{13}C$ NMR Spectrum of Compound 17 (151 MHz, $CD_3CN$)**

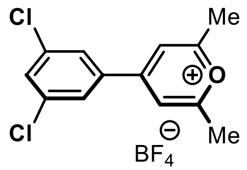


**S22**

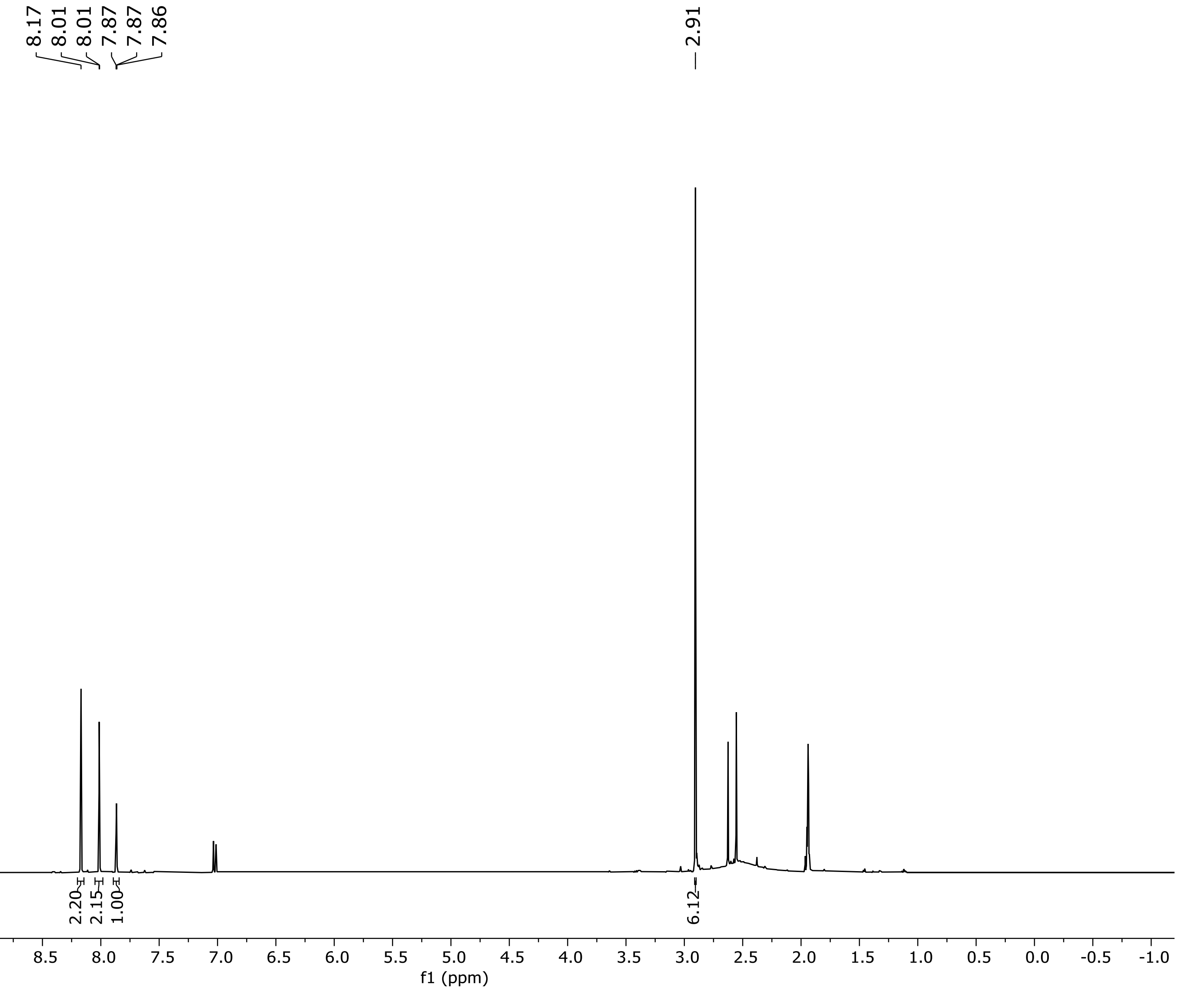


**$^1$H NMR Spectrum of Compound S22 (500 MHz, $CD_3CN$)**

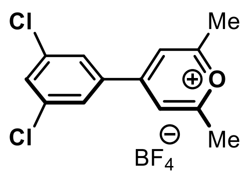


**S22**

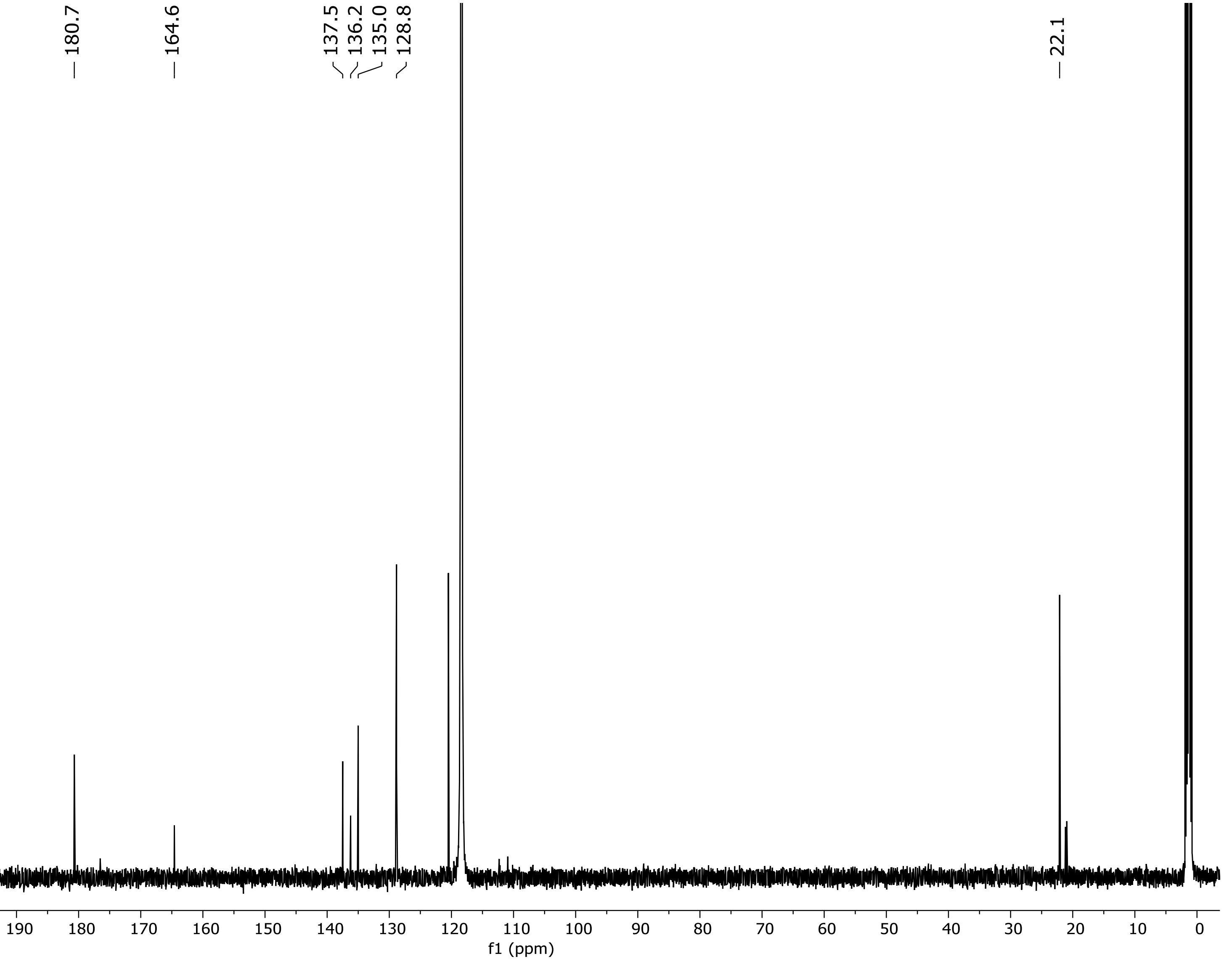


**$^{13}C$ NMR Spectrum of compound S22 (126 MHz, $CD_3CN$)**

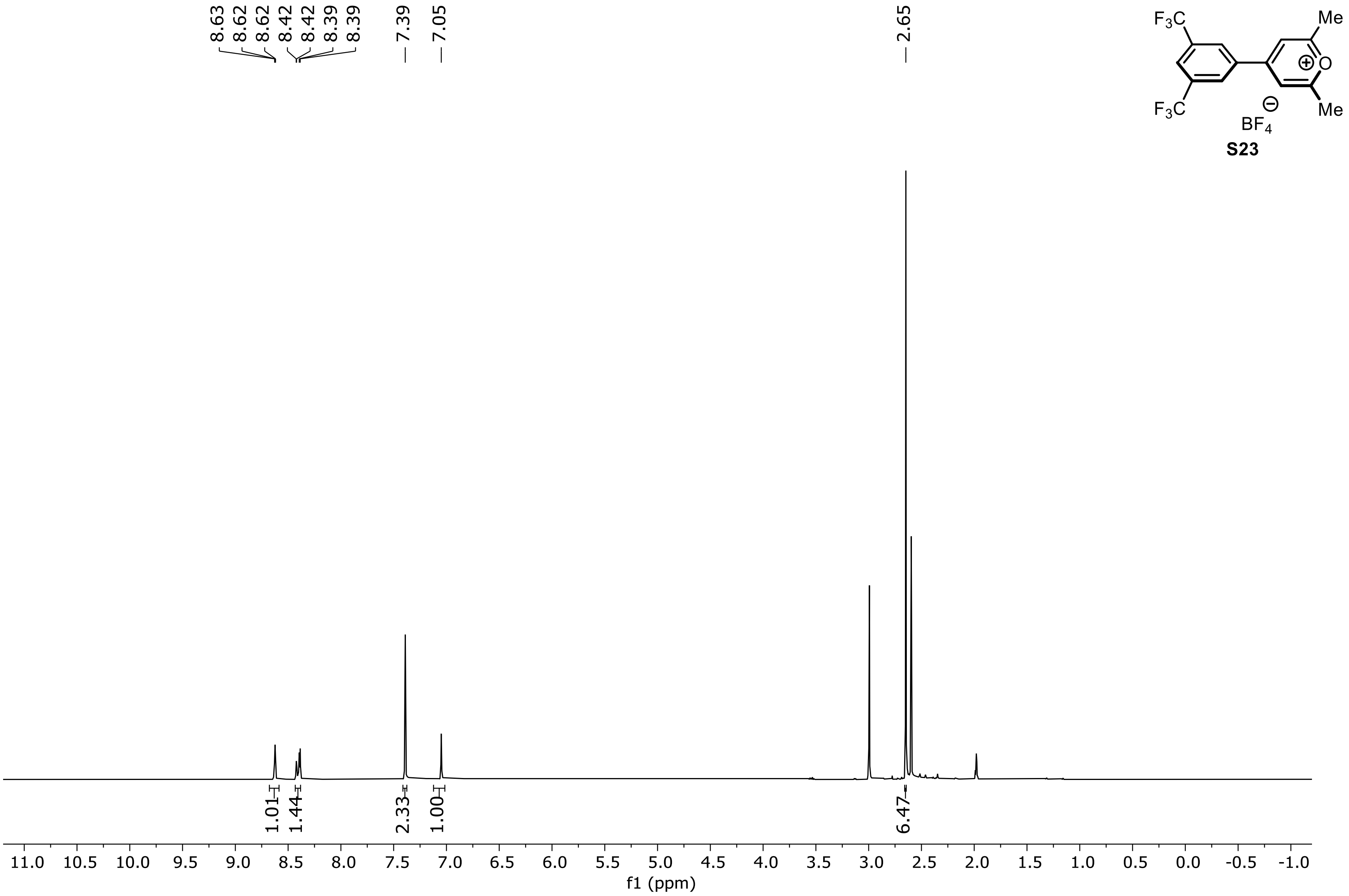


VA-117.20.fid —

**[1]H NMR Spectrum of Compound S23 (500 MHz, $CD_3CN$)**

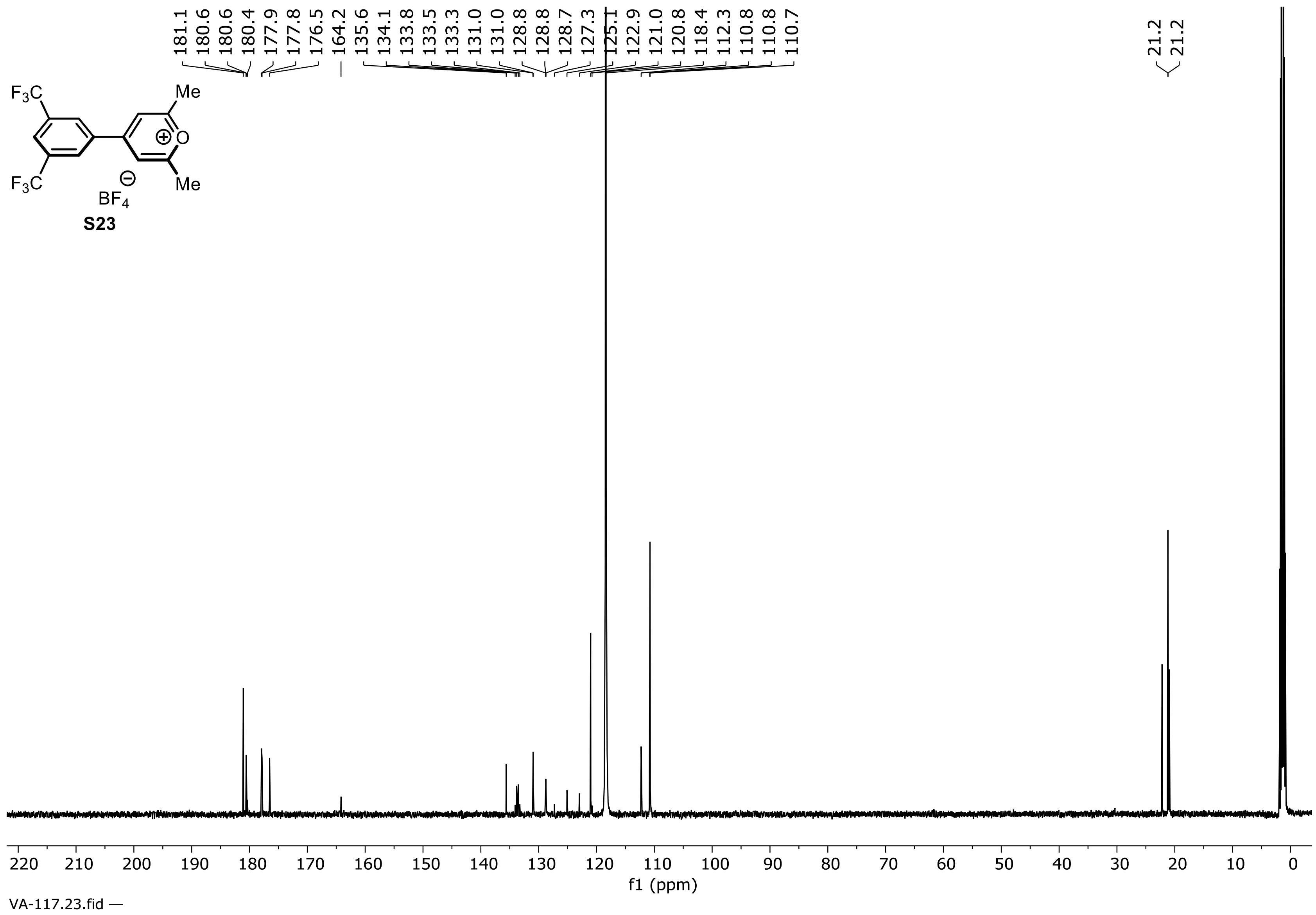


**$^{13}C$ NMR Spectrum of compound S23 (126 MHz, $CD_3CN$)**

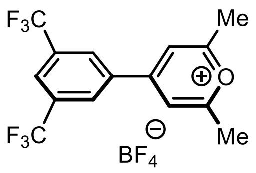


**S23**

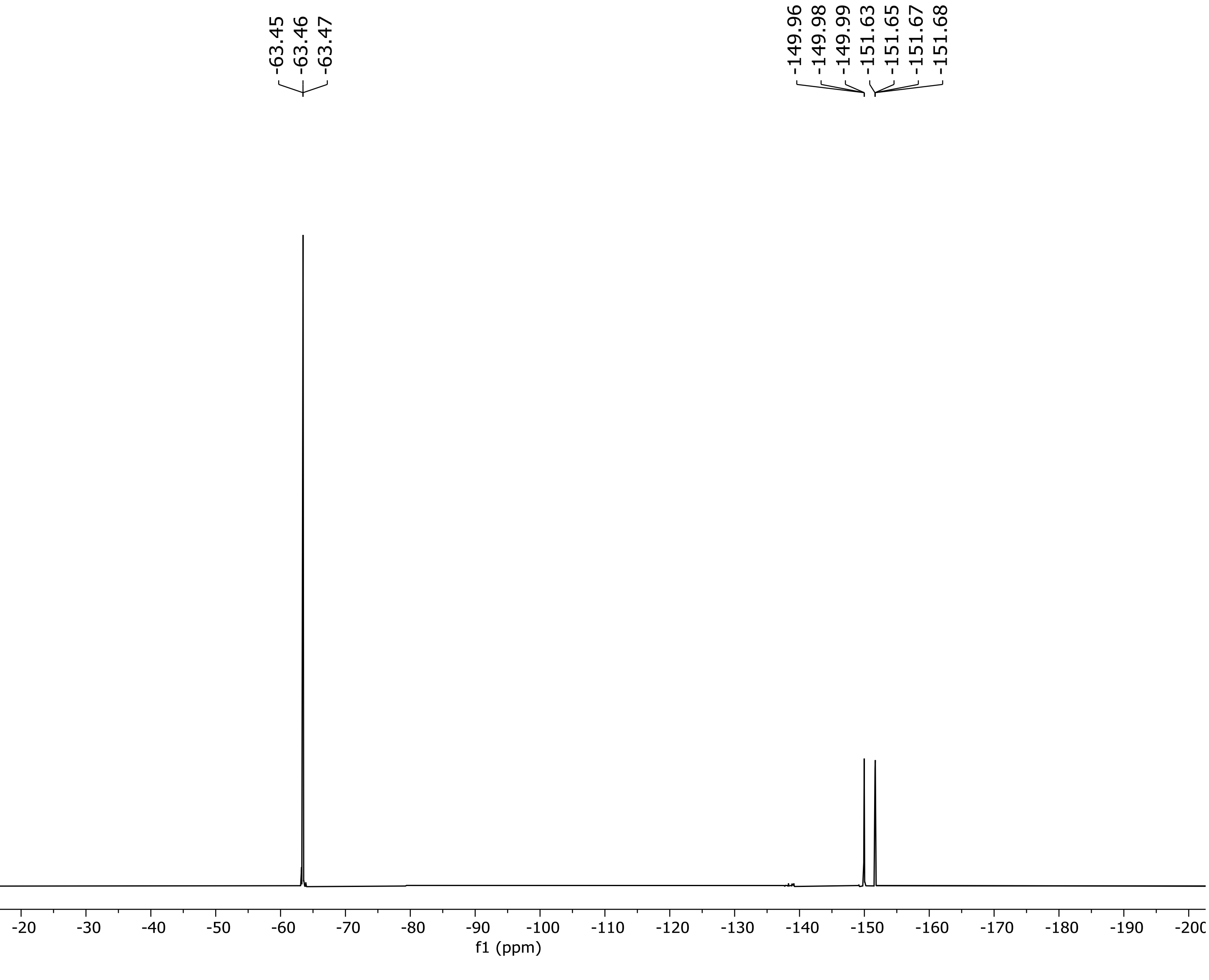


**$^{19}$F NMR Spectrum of compound S23 (471 MHz, $CD_3CN$)**

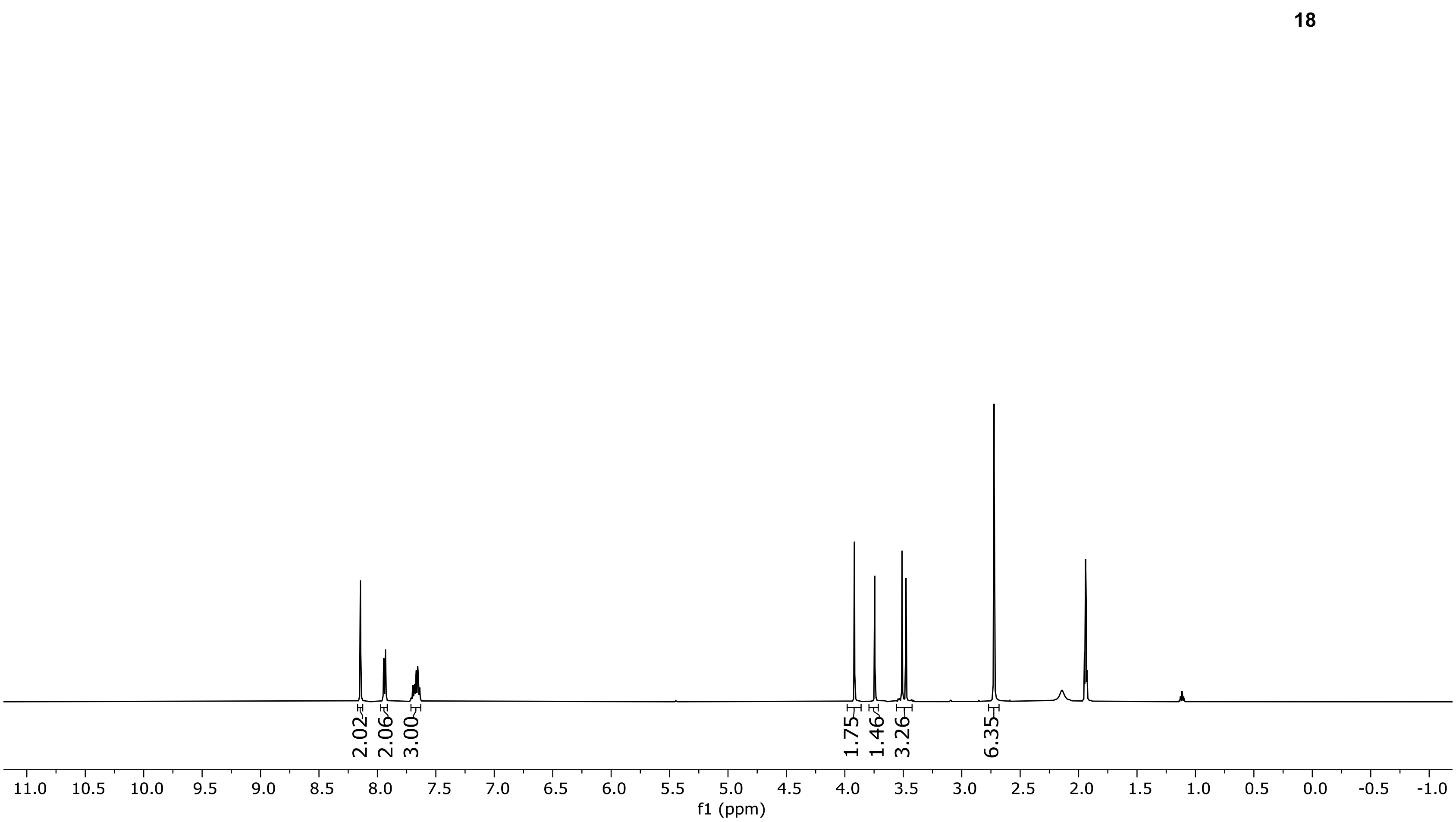


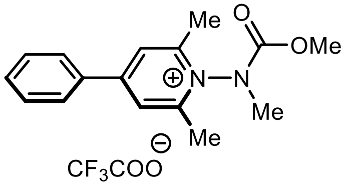


**18**

VA-10P.10.fid —

**$^1$H NMR Spectrum of Compound 18 (500 MHz, $CD_3CN$)**

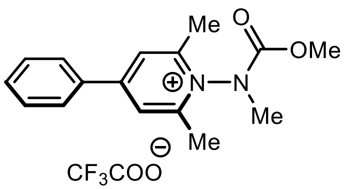


**18**

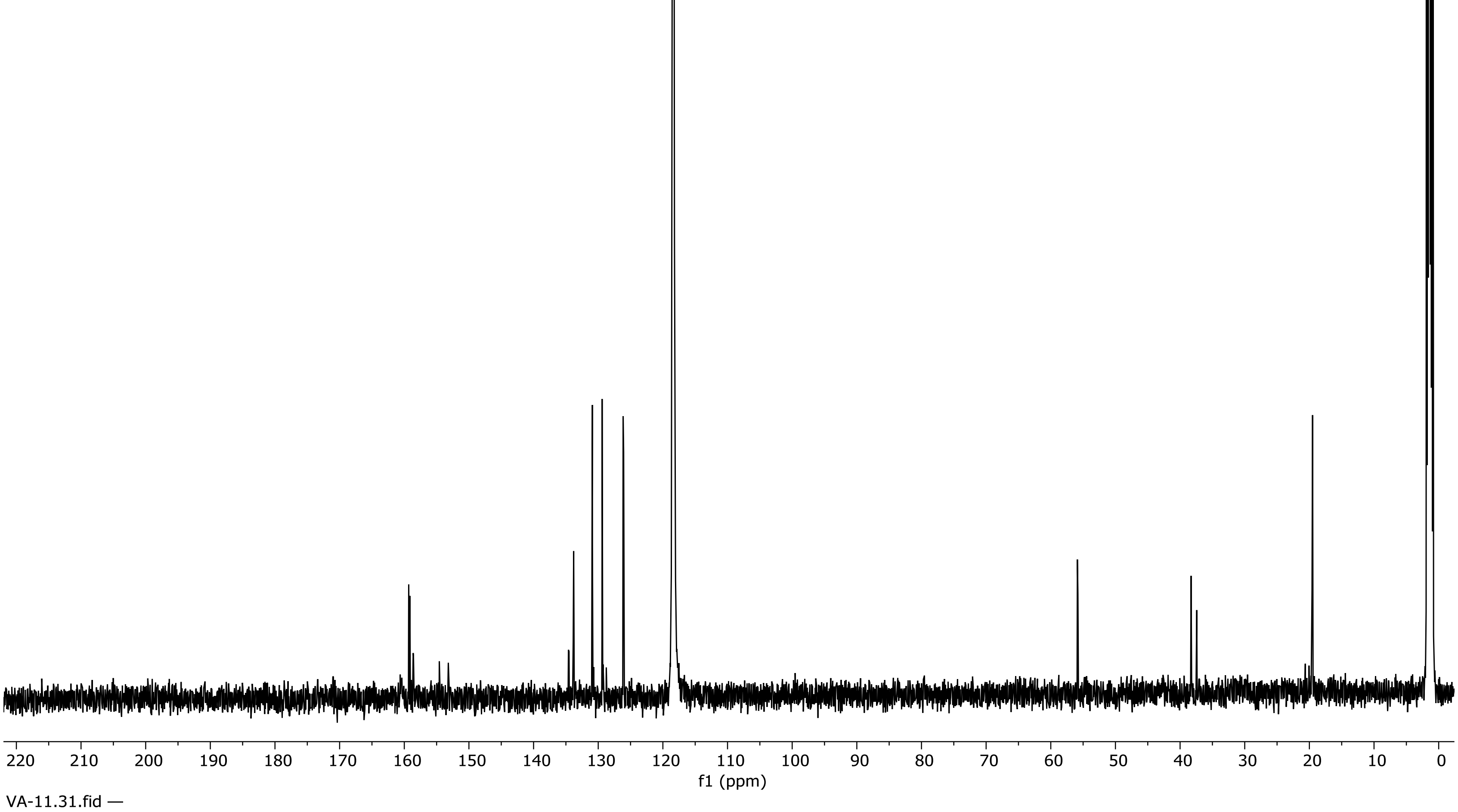


**$^{13}C$ NMR Spectrum of compound 18 (126 MHz, $CD_3CN$)**

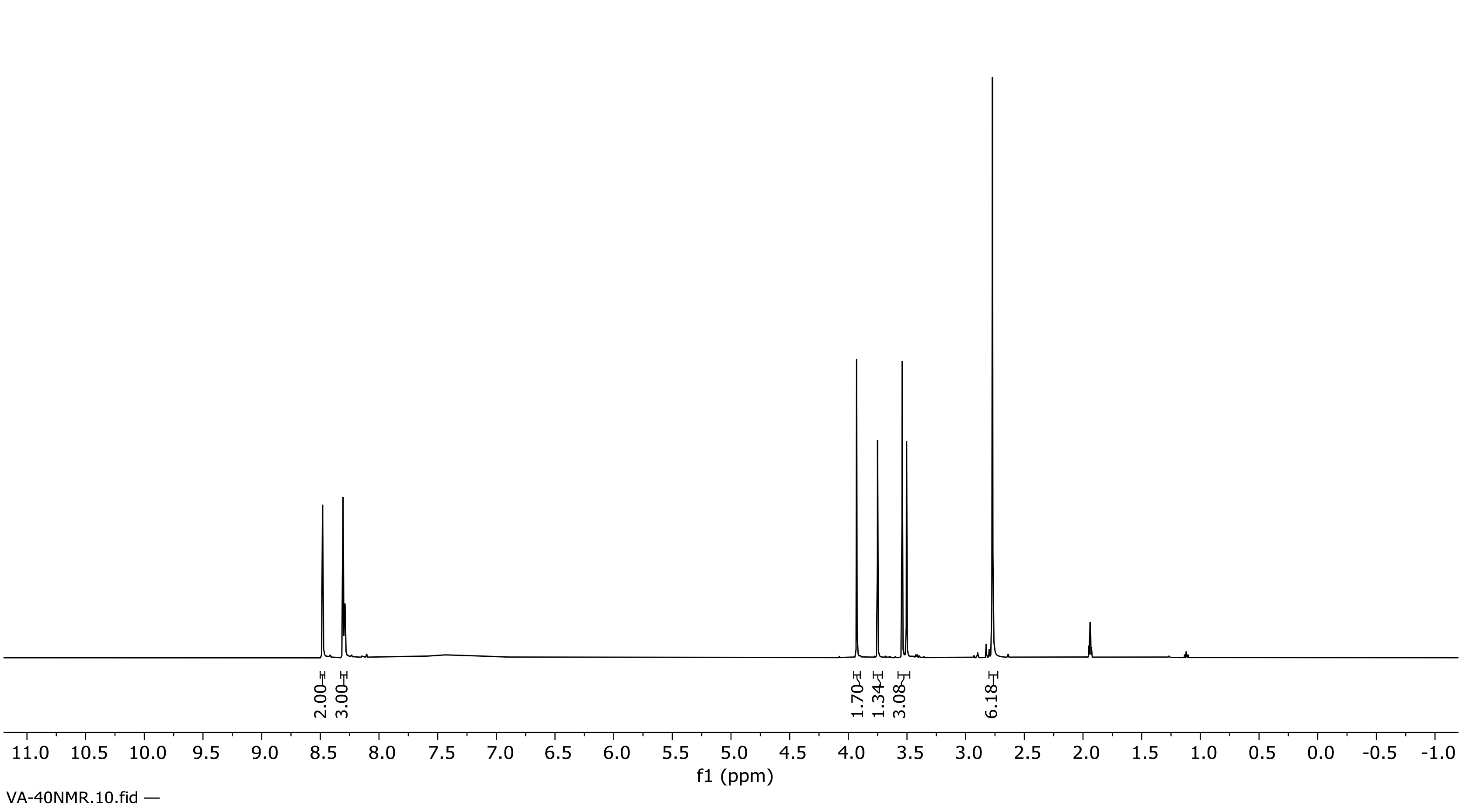


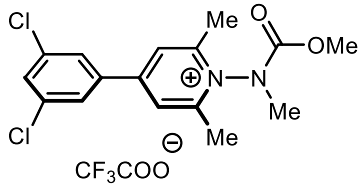


**19**

**¹H NMR Spectrum of Compound 19 (500 MHz, $CD_3CN$)**

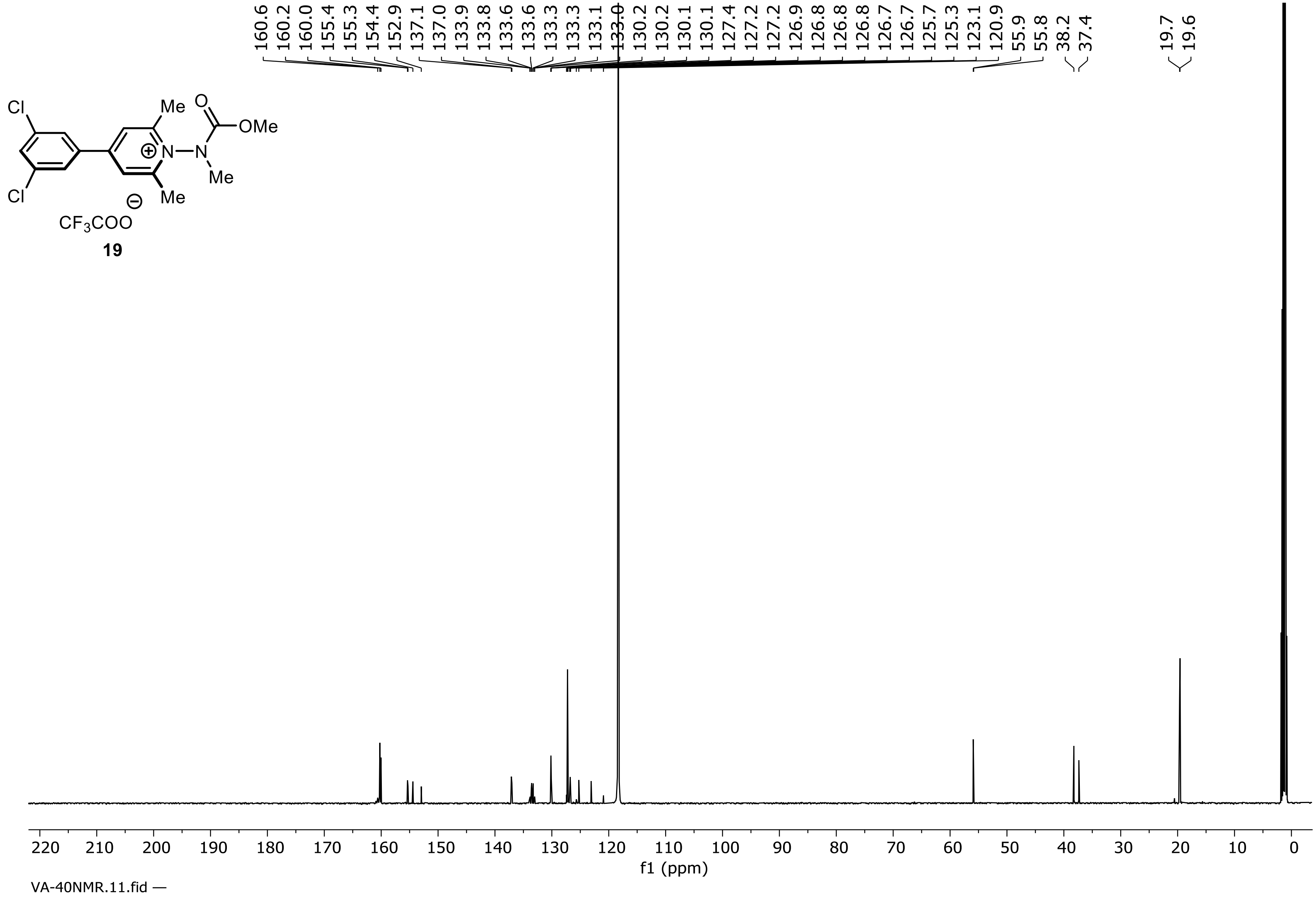


**$^{13}C$ NMR Spectrum of compound 19 (126 MHz, $CD_3CN$)**

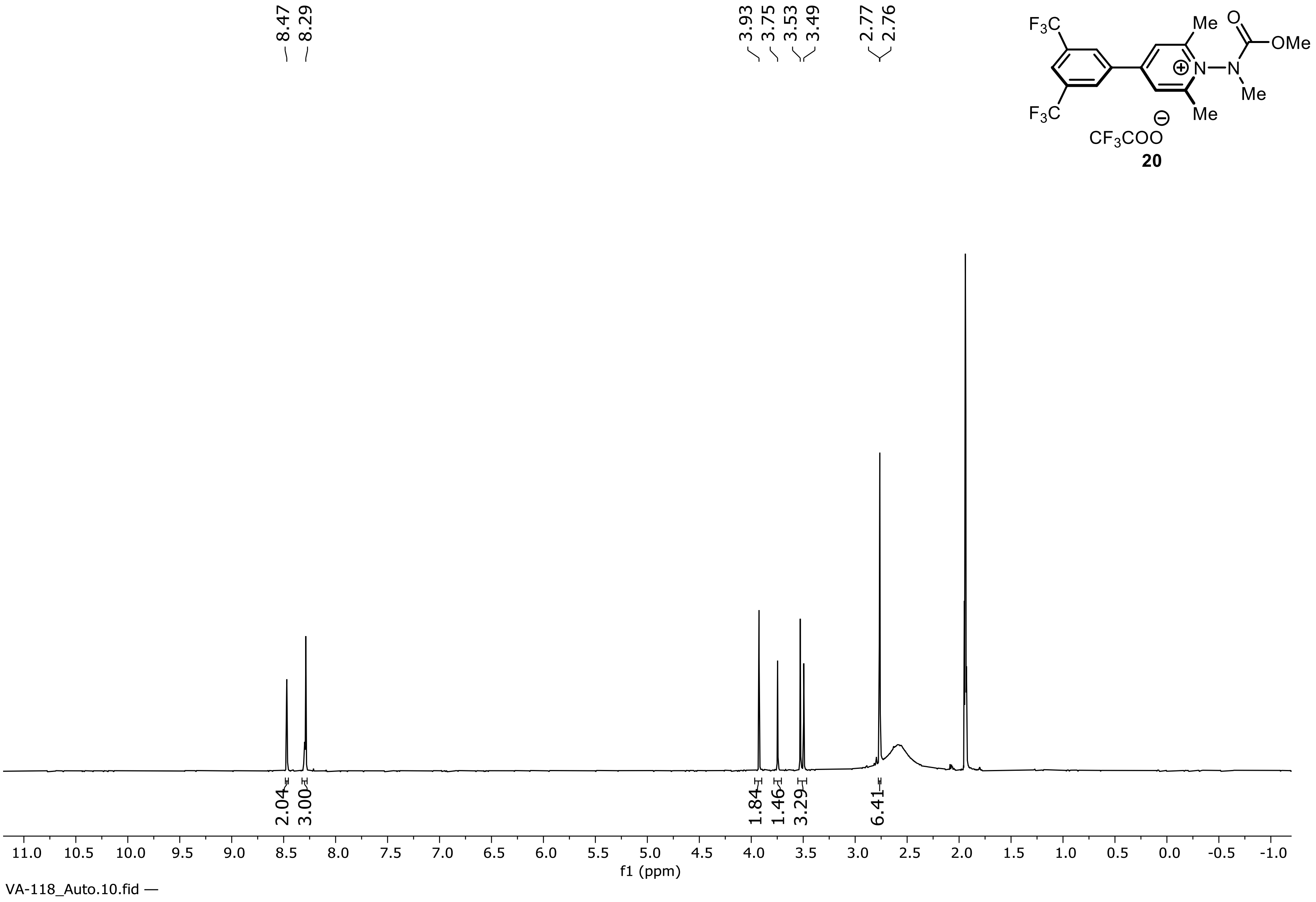


**[1]H NMR Spectrum of Compound 20 (500 MHz, $CD_3CN$)**

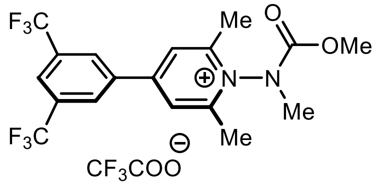


**20**

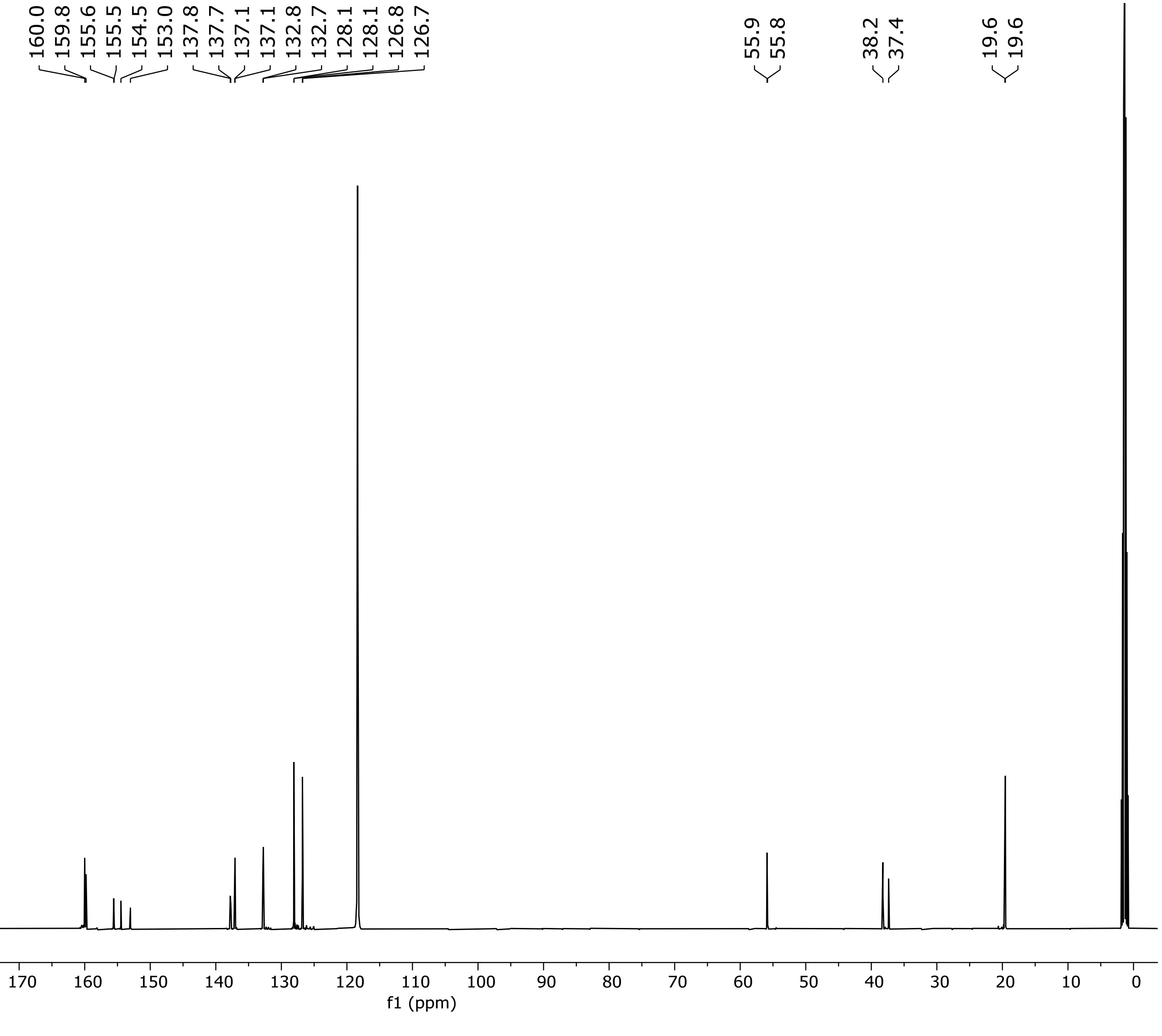


**$^{13}$C NMR Spectrum of compound 20 (126 MHz, $CD_3CN$)**

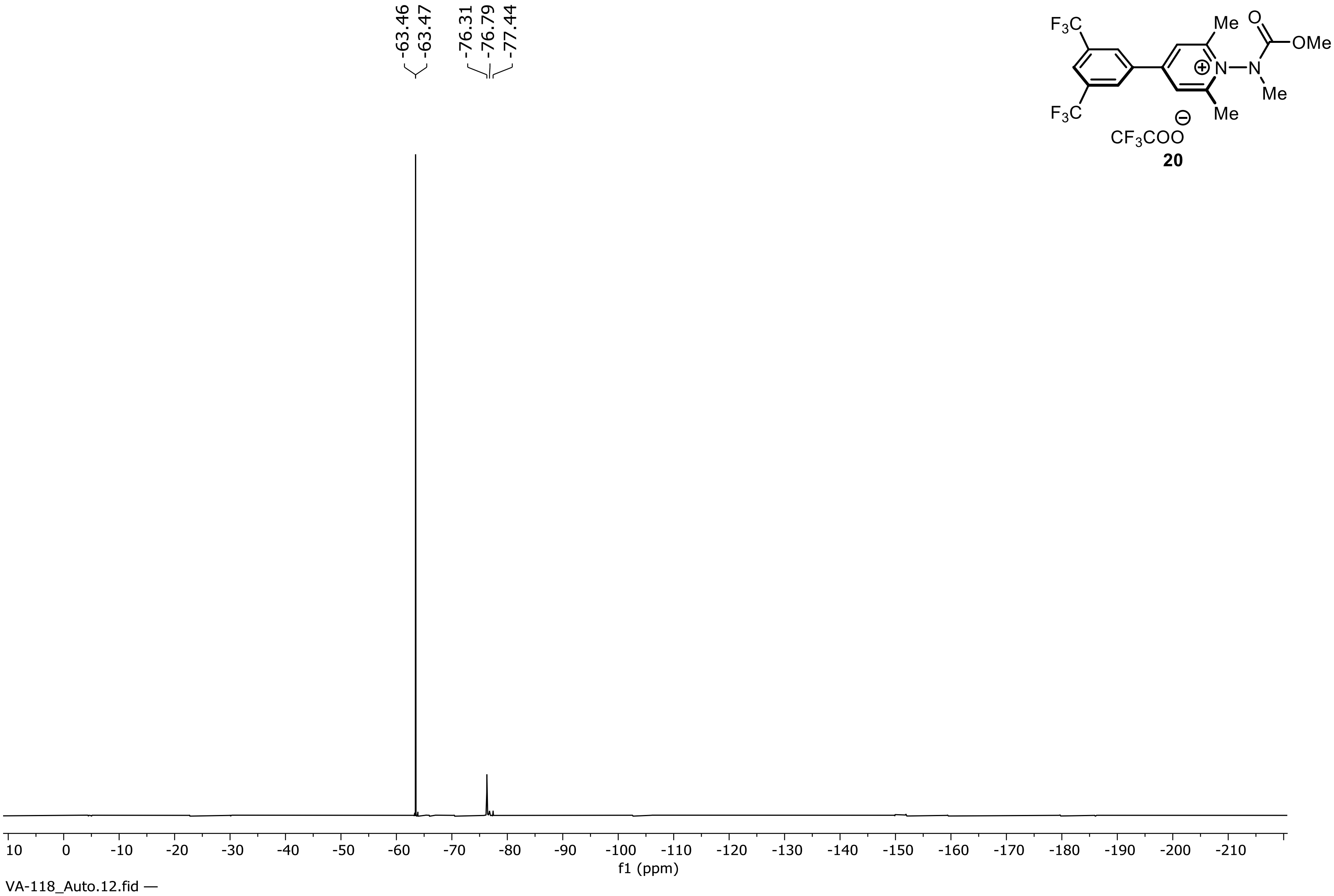


**20**

**$^{19}$F NMR Spectrum of compound 20 (471 MHz, $CD_3CN$)**

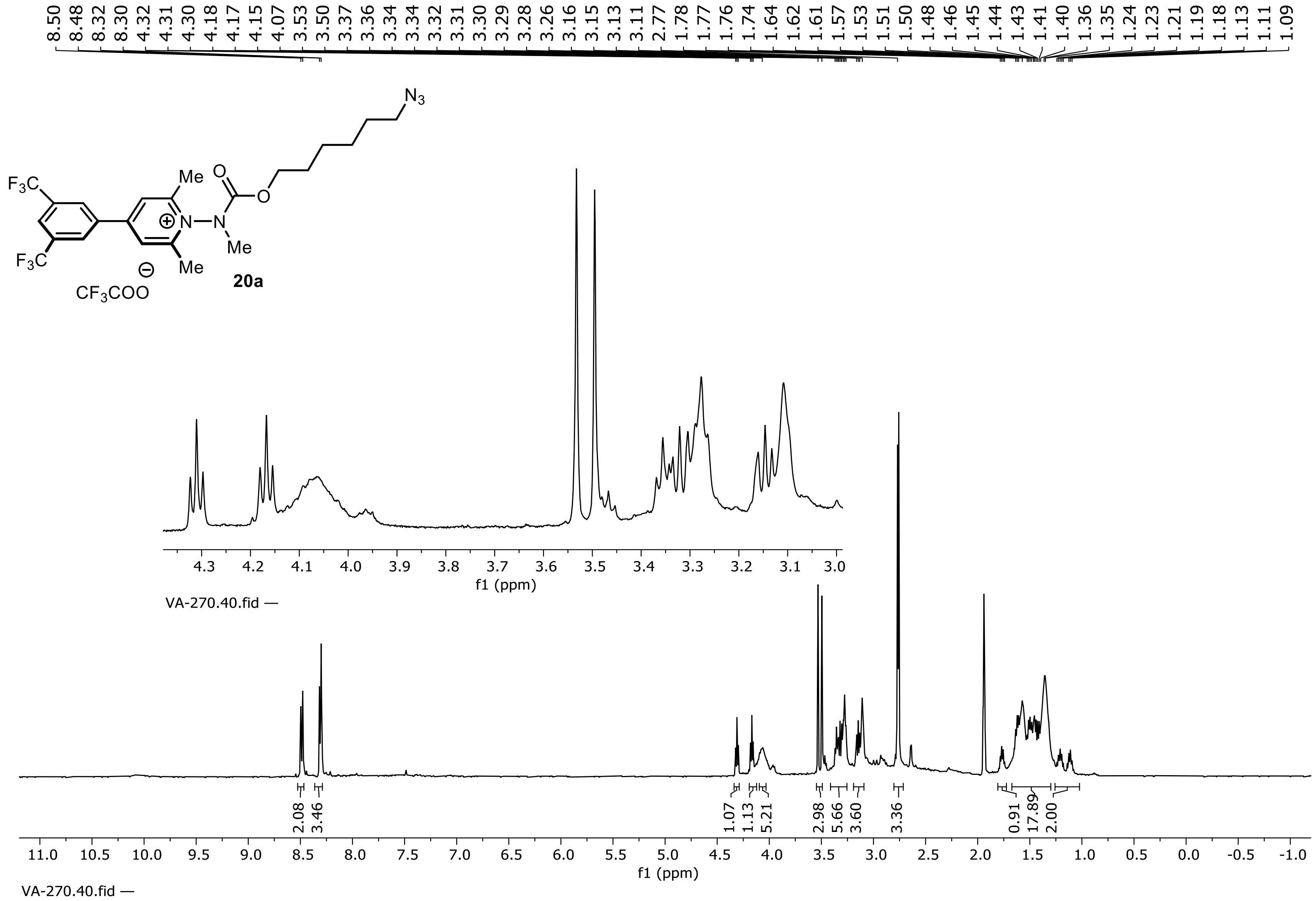


**$^1$H NMR Spectrum of Compound 20a (500 MHz, $CD_3CN$)**

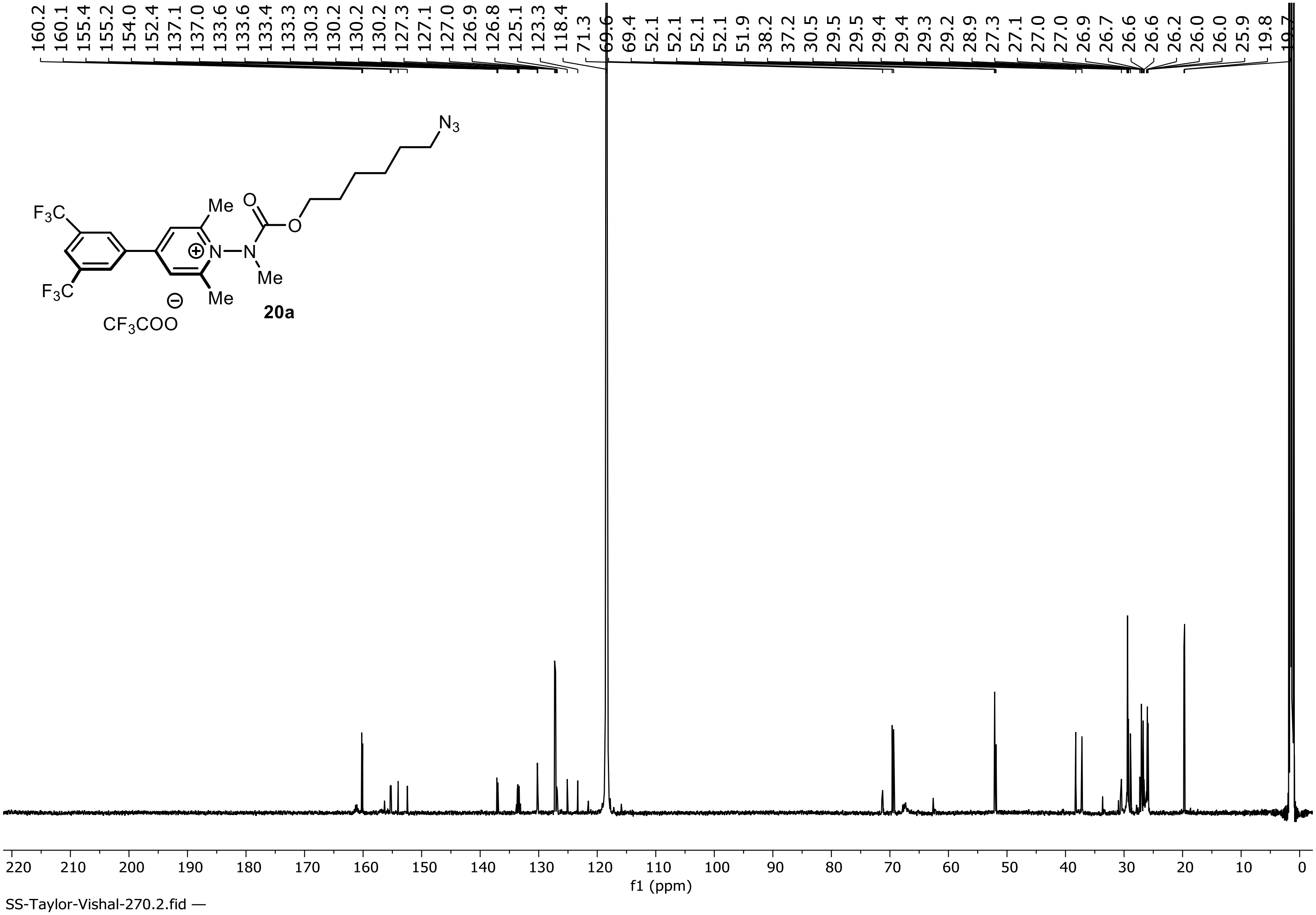


**$^{13}C$ NMR Spectrum of compound 20a (151 MHz, $CD_3CN$)**

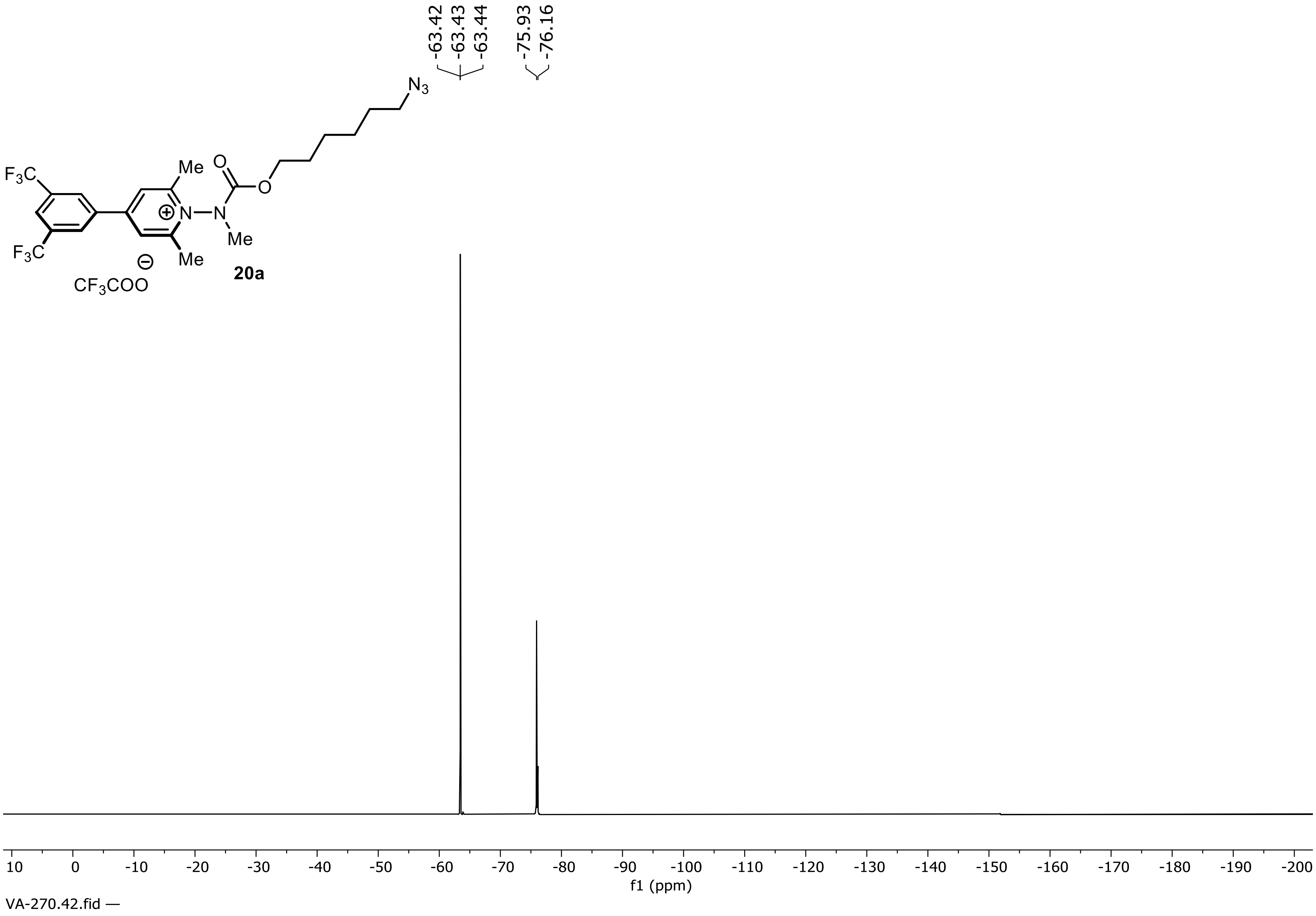


**$^{19}F$ NMR Spectrum of compound 20a (471 MHz, $CD_3CN$)**

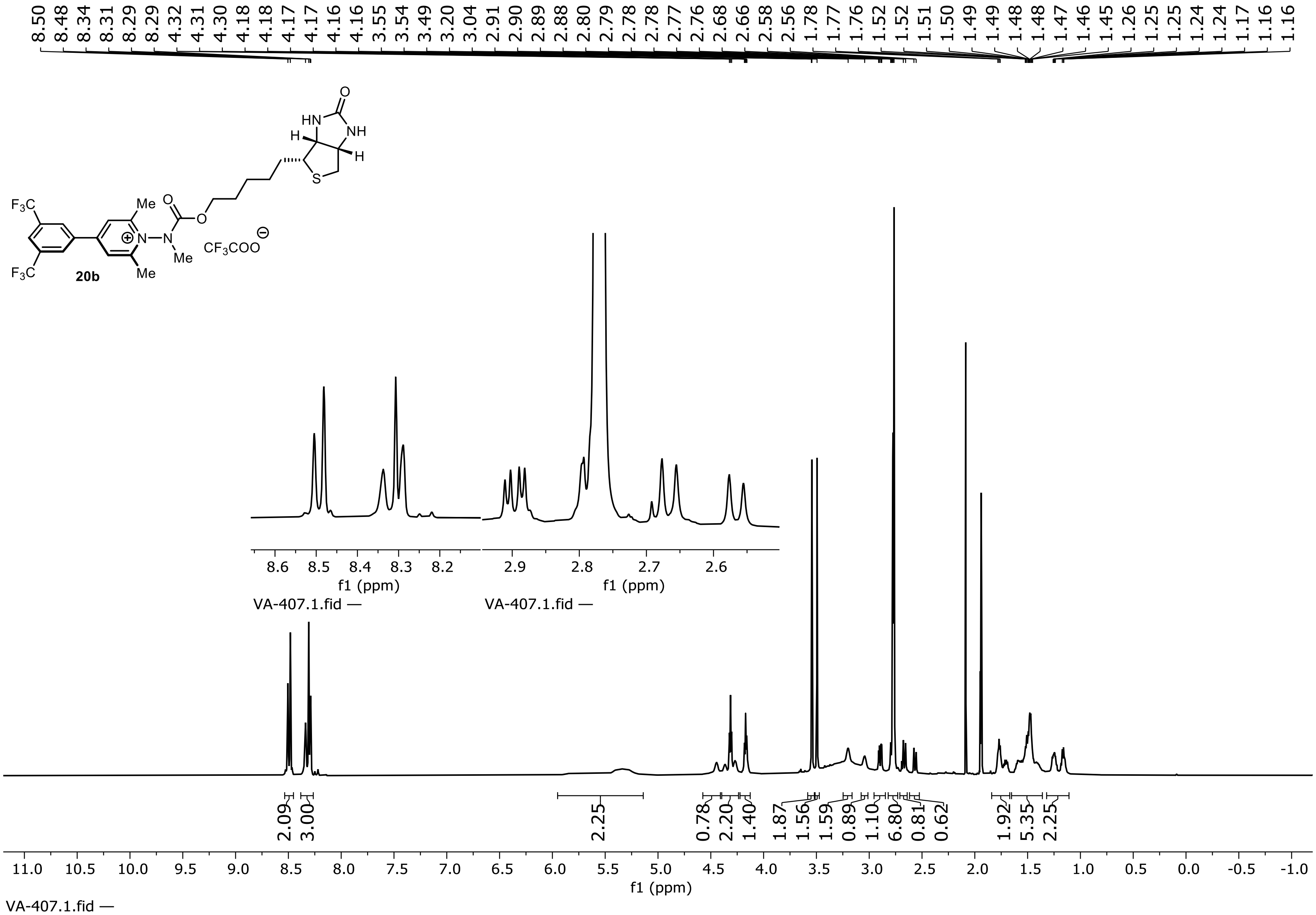


**$^1$H NMR Spectrum of Compound 20b (600 MHz, $CD_3CN$)**

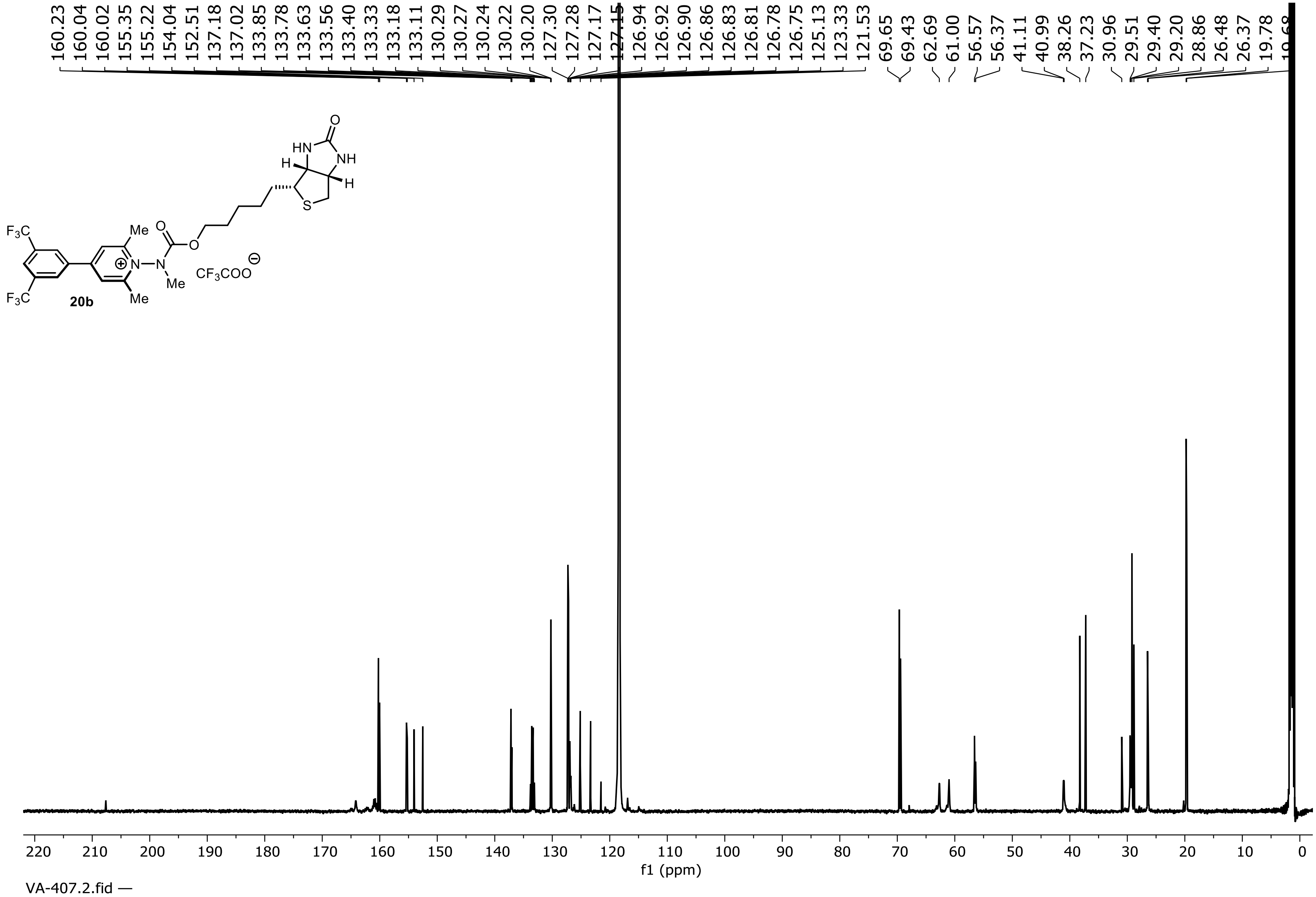


**$^{13}$C NMR Spectrum of compound 20b (151 MHz, $CD_3CN$)**

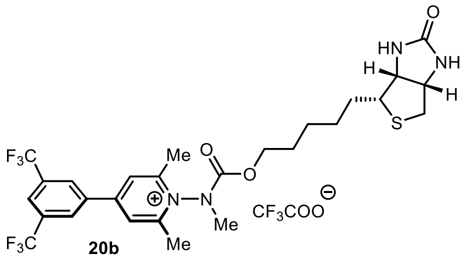

20b

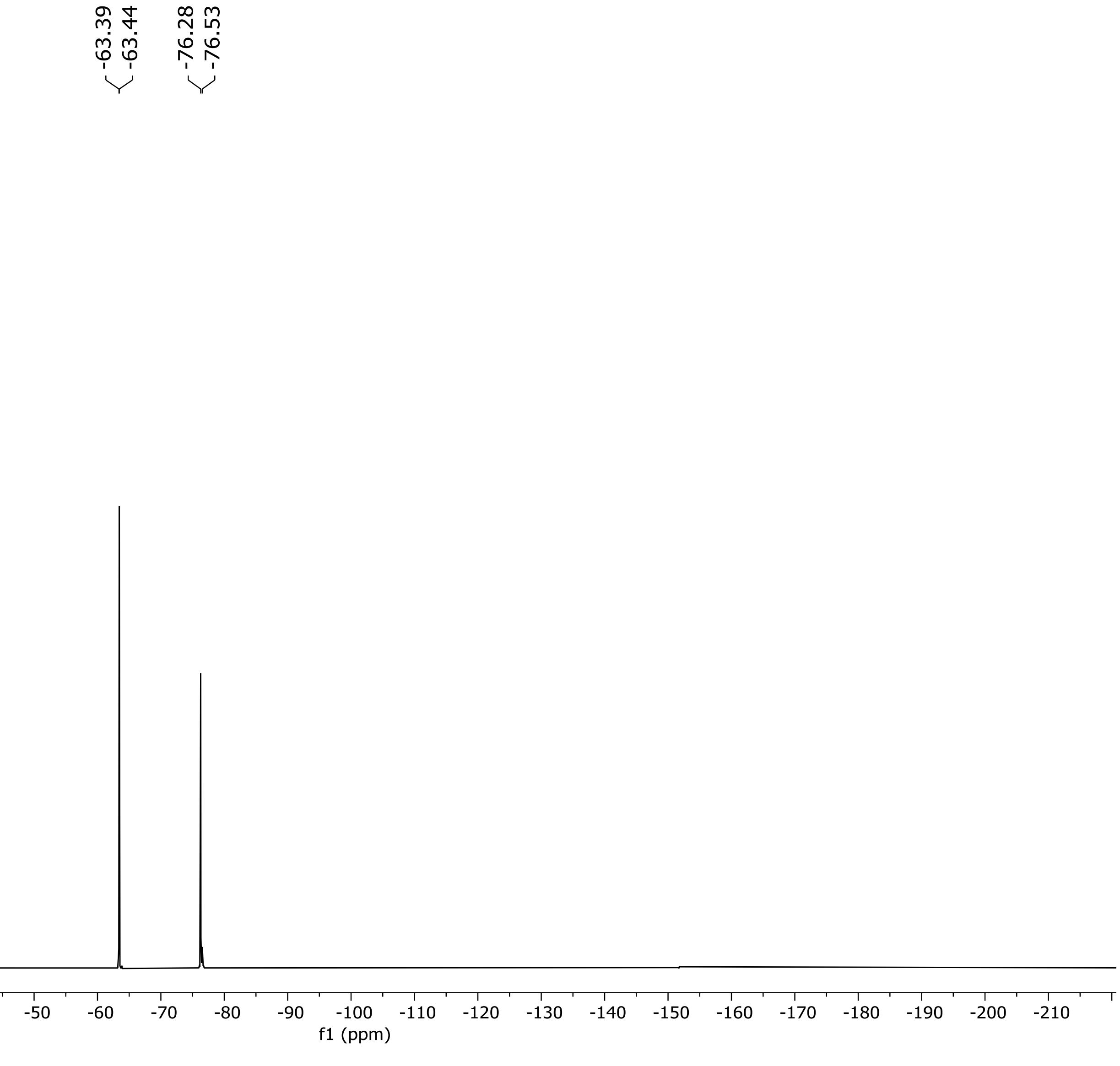


**$^{19}$F NMR Spectrum of compound 20b (471 MHz, $CD_3CN$)**
