## Supplementary material for "Cation-Cation Photosensitization for Protein Ligation and Intracellular Catalysis": Computation Details

**Table of Contents**

**1. Ground State Calculations of select pyridiniums and sensitizers**........... CSI2
**2. References**.......................................................................................CSI11

**Computational Details**

All calculations were performed using the Gaussian 16 computational chemistry software package[1]. Optimized structures were viewed and assessed in gaussview[2].

### 1. Ground State Calculations of select pyridiniums and sensitizers.

The ground state geometries of **16**, **18**, **19**, **20**, **21** were calculated at the UMN12-SX/6-31G(d)/SMD=$H_2O$ theory level. Frequency calculations were performed for all compounds at the same level of theory and revealed no imaginary frequencies in any structures.

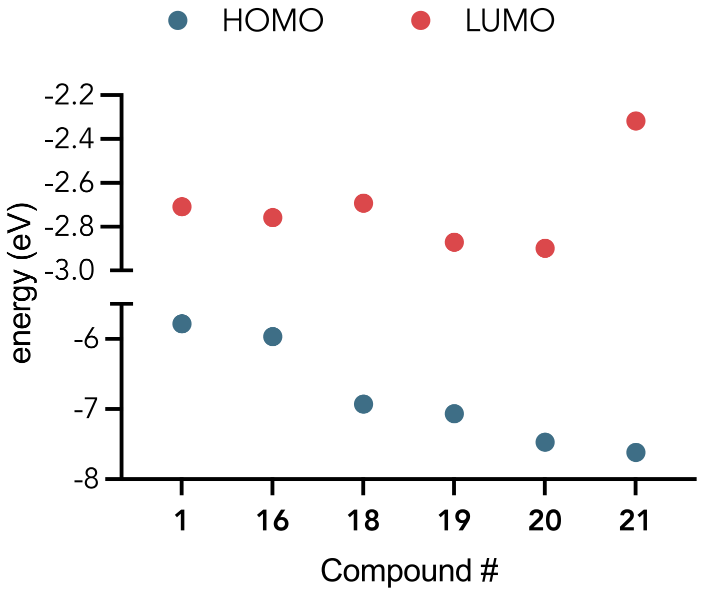


**Figure CSI1**. Calculated HOMO and LUMO energies derived from ground state calculations of catalyst and probe structures **1**, **16**, **18**, **19**, **20**, **21**. See reference 3 for calculations involving probe **1.**

**Ground state geometry for structures 16, 18, 19, 20, 21.**

Geometry of **16**:

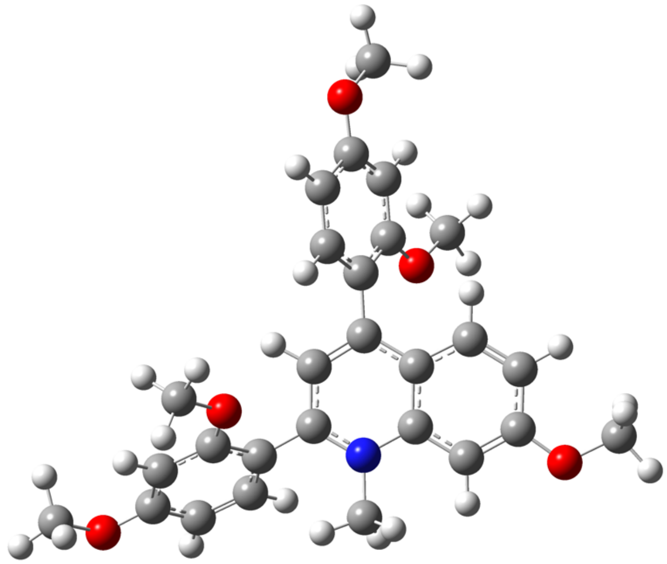

C,0,-2.8966753655,-0.4711073954,0.7128673344
C,0,-4.7241898209,-0.2447549108,2.2307874644
C,0,-5.0051811623,-1.5613266865,0.1722849005
C,0,-5.5405192912,-1.064381838,1.3964502676
C,0,-5.2475636847,0.3055655634,3.4050720862
C,0,-6.8438516374,-1.3711851764,1.8520341921
C,0,-6.54916009,0.0159489313,3.7879134824
H,0,-4.6735819937,0.9736488618,4.0405701421
H,0,-7.459307499,-2.0528440682,1.2674558757
N,0,-3.4003106083,-0.0112510186,1.880699149
C,0,-3.6946735507,-1.2450934403,-0.1321899309
H,0,-3.2611258441,-1.5619758985,-1.0768814885
C,0,-5.8100898141,-2.3288234426,-0.7947284986
C,0,-7.0469290388,-1.854706724,-1.2363161584
C,0,-5.3174824567,-3.5285160329,-1.3640445323
C,0,-7.7962479444,-2.5112157418,-2.2050049495
H,0,-7.4240654252,-0.9147208586,-0.8305212094
C,0,-6.0582956214,-4.2049288978,-2.3231486623
C,0,-7.2917542097,-3.6951528009,-2.7443359328
H,0,-8.7453003634,-2.0919211696,-2.5292163042
H,0,-5.7108917897,-5.1344089961,-2.7679977519
C,0,-7.3496662968,-0.8551683911,3.0182462497

H,0,-8.3544009354,-1.1244581858,3.3361198959
C,0,-1.5413298422,-0.1052469399,0.2716668344
C,0,-1.1505277191,1.2368618112,0.1661959546
C,0,-0.6443319693,-1.1023800892,-0.1604680539
C,0,0.0766408795,1.5928940497,-0.3555602856
H,0,-1.8454963847,2.0174443712,0.4804745597
C,0,0.6023079409,-0.7589884081,-0.684515086
C,0,0.9539193611,0.5883481539,-0.7829482503
H,0,0.3724706956,2.635992277,-0.4517465423
H,0,1.2901504385,-1.5351350163,-1.0055453046
O,0,-7.9233292907,-4.4309877538,-3.6895729978
C,0,-9.175978985,-3.9632173443,-4.1586829157
H,0,-9.0790648927,-2.9763895704,-4.6334425073
H,0,-9.5098402546,-4.6933591187,-4.9020276213
H,0,-9.9109456888,-3.9096586081,-3.3429189737
C,0,-2.548517469,0.6707215553,2.8608993661
H,0,-2.6996778731,0.2058161047,3.8401948277
H,0,-1.5024833289,0.5523358808,2.5800659893
H,0,-2.7991024729,1.7364143288,2.9159571222
O,0,-4.1269357766,-3.9734887412,-0.8980726994
C,0,-3.605150938,-5.1710287246,-1.4456618557
H,0,-4.2806452889,-6.0187340688,-1.2621574346
H,0,-3.4249933157,-5.0688178554,-2.5253718809
H,0,-2.6556854165,-5.348155308,-0.9313027396
O,0,-1.0504789643,-2.3819341202,-0.0052246854
O,0,2.1356654012,1.013559643,-1.2827027862
C,0,3.0591994428,0.0401193222,-1.7394390849
H,0,3.3687057159,-0.6274871031,-0.9227058603
H,0,2.6361438375,-0.5528650759,-2.5627838397
H,0,3.927544831,0.5985195952,-2.1015432985
C,0,-0.1864756674,-3.4207049018,-0.4335357792
H,0,-0.0057946273,-3.3655656709,-1.5163585252
H,0,0.770710779,-3.3880623018,0.1058460175
H,0,-0.7021417657,-4.3551535218,-0.1930227002
O,0,-6.9656662334,0.6017145191,4.9258663989
C,0,-8.2901677997,0.346284619,5.3698302005
H,0,-8.4335348783,-0.7198561861,5.593241465
H,0,-8.4122941643,0.9304554469,6.286436966
H,0,-9.0270857623,0.6748721275,4.624219365

Zero-point correction= 0.508809 (Hartree/Particle)

Thermal correction to Energy= 0.538949
Thermal correction to Enthalpy= 0.539893
Thermal correction to Gibbs Free Energy= 0.448271
Sum of electronic and zero-point Energies= -1474.778755
Sum of electronic and thermal Energies= -1474.748615
Sum of electronic and thermal Enthalpies= -1474.747671
Sum of electronic and thermal Free Energies= -1474.839293

Geometry of **18**:

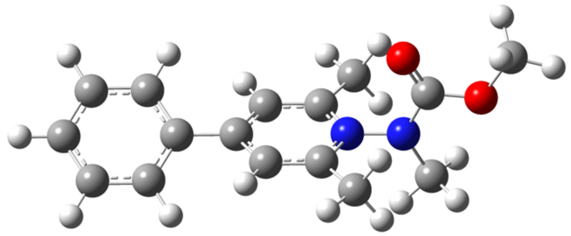

C,0,-2.5065584853,-0.0198007789,-0.5802909919
C,0,-3.1374167582,0.6079945821,0.4924279272
H,0,-0.6385217033,-0.6180364913,-1.4556451409
C,0,-1.1229265411,-0.1447950765,-0.6007267202
C,0,-0.3500803747,0.3728660679,0.4492565385
C,0,1.1176794271,0.2531915055,0.4272882197
C,0,1.9302373914,1.2267634436,1.0251651739
C,0,1.748153172,-0.8339608656,-0.1939320298
C,0,3.3048578175,1.1176554535,1.011382858
C,0,3.1224128729,-0.9504237627,-0.2086777609
H,0,1.1693820783,-1.6310551298,-0.6570829867
N,0,3.8533914892,0.0247759489,0.4055423026
C,0,3.8444847096,-2.0825597893,-0.8483292955
H,0,4.503364251,-1.7209887708,-1.6500936523
H,0,4.4634005424,-2.6233477177,-0.1197079634
H,0,3.1170088399,-2.7766053264,-1.2788606849
N,0,5.2381962781,-0.0979316157,0.4067225769
C,0,5.8830752202,0.4719808818,-0.6559935802
O,0,5.3152716711,1.0518175178,-1.563757761
C,0,5.8837100706,-0.8020195621,1.5091119038
H,0,6.4344529276,-1.6736199636,1.1348975037
H,0,6.5703895924,-0.1317955085,2.0404054723

H,0,5.1043725675,-1.1388059146,2.1993791419
O,0,7.1926345456,0.2981281056,-0.5421084561
C,0,7.9751060831,0.862170013,-1.6003217101
H,0,9.0130707072,0.6372300115,-1.3438124683
H,0,7.7088105068,0.3981265551,-2.5566936557
H,0,7.8196276167,1.9457501651,-1.6500175569
C,0,-0.9932248669,1.0084992715,1.5217013981
C,0,-2.3780391448,1.1176867493,1.5442456681
H,0,-0.4093892078,1.3911454688,2.3597532209
H,0,-4.223778221,0.700620481,0.5087121237
H,0,1.4994345214,2.1111240574,1.4911340692
C,0,4.2147396257,2.1320444972,1.607920407
H,0,4.8537596142,1.6866719405,2.3821437502
H,0,4.8675249104,2.5703664079,0.8401990736
H,0,3.6182641903,2.9301911828,2.0589503714
H,0,-2.8676353577,1.6008550294,2.3899036461
H,0,-3.0959000197,-0.4112720437,-1.409618742

Zero-point correction= 0.329257 (Hartree/Particle)
Thermal correction to Energy= 0.348535
Thermal correction to Enthalpy= 0.349479
Thermal correction to Gibbs Free Energy= 0.281126
Sum of electronic and zero-point Energies= -879.919388
Sum of electronic and thermal Energies= -879.900110
Sum of electronic and thermal Enthalpies= -879.899166
Sum of electronic and thermal Free Energies= -879.967519

Geometry of **19:**

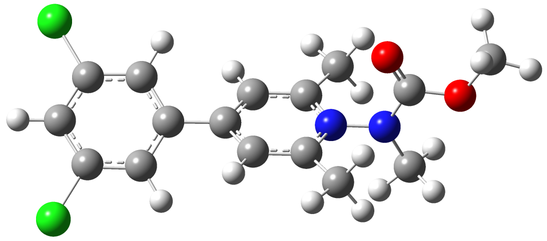

C,0,-2.4844875732,0.0240449059,-0.5809530457
C,0,-3.1406668513,0.6017382143,0.4999153129
H,0,-0.6267506418,-0.5453793134,-1.5005496646
C,0,-1.1040652594,-0.1059304223,-0.6253675194
C,0,-0.3437943845,0.3722536473,0.4475431182
C,0,1.1280432093,0.2532289789,0.4229013905
C,0,1.9337370508,1.2269386071,1.0231805462
C,0,1.7508492197,-0.8326773726,-0.2017859259
C,0,3.3097399117,1.1165608711,1.0092869834
C,0,3.1265538012,-0.9492232523,-0.2164211087
H,0,1.169712909,-1.6269320151,-0.6671962751
N,0,3.8552719544,0.0244592954,0.4010506319
C,0,3.8478376312,-2.0790192117,-0.8598458977
H,0,4.5089570134,-1.7139209052,-1.6580996344
H,0,4.4641293976,-2.6238526311,-0.1320204055
H,0,3.1199842655,-2.7696028419,-1.2951744979
N,0,5.2402190559,-0.1013625337,0.404760052
C,0,5.8879264032,0.4752051102,-0.6533584825
O,0,5.3213872247,1.0610930021,-1.5577345375
C,0,5.8809168,-0.8117472268,1.5062884259
H,0,6.4341674485,-1.6801337894,1.1285355622
H,0,6.5640964339,-0.1439049528,2.0449575737
H,0,5.0985640838,-1.1540656782,2.1904571994
O,0,7.1964840797,0.2991782706,-0.5376938951
C,0,7.9821177694,0.8688740288,-1.5908257502
H,0,9.0191045907,0.6402184167,-1.3337397803
H,0,7.7165712344,0.4115577557,-2.5506096336
H,0,7.8285385967,1.9530095515,-1.6333212492
C,0,-0.9760592646,0.964339216,1.546489214
C,0,-2.3597665221,1.0616414271,1.5540321538
H,0,-0.3994025435,1.3173513589,2.4008253461
H,0,-4.2257238839,0.6901031705,0.5201028277

H,0,1.5005456708,2.1092238492,1.4912402682
C,0,4.2193398337,2.1278660206,1.6100864428
H,0,4.8564450821,1.6788035077,2.383742026
H,0,4.8734980867,2.567447785,0.8443028527
H,0,3.6229003957,2.9251106514,2.0626429407
Cl,0,-3.1513288859,1.7793840328,2.9292631182
Cl,0,-3.4322084137,-0.5499847182,-1.9245564829

Zero-point correction= 0.309629 (Hartree/Particle)
Thermal correction to Energy= 0.331466
Thermal correction to Enthalpy= 0.332410
Thermal correction to Gibbs Free Energy= 0.257066
Sum of electronic and zero-point Energies= -1799.023043
Sum of electronic and thermal Energies= -1799.001206
Sum of electronic and thermal Enthalpies= -1799.000262
Sum of electronic and thermal Free Energies= -1799.075606

Geometry of **20**:

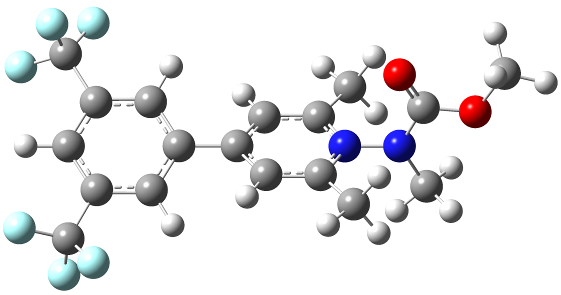

C,0,-2.4955107986,0.0238962101,-0.6057835354
C,0,-3.1484722307,0.5605518248,0.4998731886
H,0,-0.6268427023,-0.5015960887,-1.5229722932
C,0,-1.1131507994,-0.0953851808,-0.6357460567
C,0,-0.3505942741,0.3350419333,0.4572010196
C,0,1.1206797325,0.2133060517,0.437083302
C,0,1.9280960534,1.128633168,1.1219345654
C,0,1.7439439286,-0.8223463635,-0.2678390094
C,0,3.3037955844,1.0139621943,1.1054641148
C,0,3.1192930826,-0.9394337818,-0.2913434767
H,0,1.1638172621,-1.5776272202,-0.7950401806
N,0,3.8492528775,-0.0211153457,0.4041366201

C,0,3.8389401942,-2.0128545873,-1.0267661284
H,0,4.4751350205,-1.5839209647,-1.8134841589
H,0,4.4802028807,-2.5966651327,-0.3530733231
H,0,3.1098152999,-2.6823217219,-1.4918457776
N,0,5.2347226425,-0.14003501,0.3842245902
C,0,5.8694164055,0.5351360184,-0.62305811
O,0,5.2904820827,1.192639834,-1.4682300231
C,0,5.8910739301,-0.9304701356,1.4195716025
H,0,6.4636678687,-1.7485671638,0.9663582498
H,0,6.5593538536,-0.2993218713,2.018142313
H,0,5.1163373168,-1.3509400746,2.0680933086
O,0,7.1801878536,0.3619842674,-0.532479308
C,0,7.9516815857,1.0278006254,-1.5385784979
H,0,8.9926253483,0.7847176839,-1.3126038682
H,0,7.6791202917,0.6553980618,-2.5324663724
H,0,7.7909602192,2.1103466724,-1.4835766902
C,0,-1.0007994096,0.8802363383,1.5682382527
C,0,-2.3869225611,0.9845574022,1.581308711
H,0,-0.4289832052,1.2025165801,2.4394201388
H,0,-4.2352580785,0.6461089123,0.5146539807
C,0,-3.0650526839,1.5190885169,2.8129505089
F,0,-4.2844317777,1.9977512707,2.5457977536
F,0,-3.2146939513,0.5726217885,3.7515997232
F,0,-2.3589386345,2.507441814,3.3758797094
C,0,-3.3117943223,-0.3844120856,-1.8015610141
F,0,-2.6725478177,-1.2750006325,-2.5669396958
F,0,-3.6068717539,0.6630615243,-2.5856518451
F,0,-4.4785913704,-0.9307419674,-1.4374840904
H,0,1.4990342163,1.9703256458,1.6626991317
C,0,4.2137678592,1.9595630126,1.8048394846
H,0,4.8329324228,1.4362897212,2.5461670594
H,0,4.8858372245,2.4579560542,1.0926636672
H,0,3.6183489337,2.7204941712,2.3172823489

Zero-point correction= 0.338670 (Hartree/Particle)
Thermal correction to Energy= 0.365403
Thermal correction to Enthalpy= 0.366347
Thermal correction to Gibbs Free Energy= 0.278509
Sum of electronic and zero-point Energies= -1553.630187
Sum of electronic and thermal Energies= -1553.603453

Sum of electronic and thermal Enthalpies= -1553.602509
Sum of electronic and thermal Free Energies= -1553.690347

Geometry of **21**:

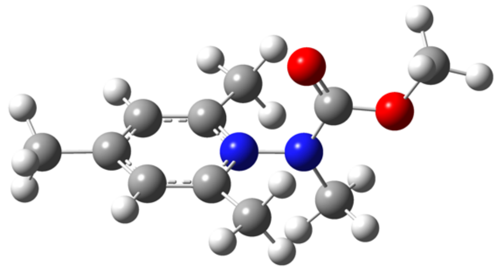

C,0,1.1195041246,0.2643113445,0.4123080513
C,0,1.9258220715,1.2429160316,1.0000270161
C,0,1.745742429,-0.8317823802,-0.1893481694
C,0,3.3019031174,1.1310256758,0.9981546081
C,0,3.1202550715,-0.9529852902,-0.1975717558
H,0,1.1576175413,-1.6131239268,-0.6689939642
N,0,3.8490295549,0.0296016299,0.4080777267
C,0,3.8446994471,-2.0872294986,-0.8309357188
H,0,4.5060226527,-1.7300371387,-1.6326315902
H,0,4.4609682551,-2.6256398153,-0.0983274898
H,0,3.1178538767,-2.7829305309,-1.2599778032
N,0,5.2344267525,-0.0924700871,0.4111310403
C,0,5.8796042229,0.4712857345,-0.6544193007
O,0,5.312626716,1.0496386122,-1.5637395459
C,0,5.8786552784,-0.7979744477,1.5129980173
H,0,6.424183593,-1.6733710587,1.1398826259
H,0,6.5700246423,-0.1301134387,2.0411228425
H,0,5.0992694028,-1.1287282213,2.2061123698
O,0,7.18897816,0.2942349734,-0.5416559799
C,0,7.9713763666,0.8518639733,-1.6032556679
H,0,9.0090843796,0.6245073856,-1.3478185629
H,0,7.7019429855,0.385222567,-2.5574976032
H,0,7.8193697361,1.9357602353,-1.6567518929
H,0,1.4818430212,2.1189291522,1.4709325992
C,0,4.2117165093,2.1443821628,1.5965171734
H,0,4.8481151106,1.6992394929,2.3729899936

H,0,4.867220393,2.5812972094,0.8302815259
H,0,3.6149573871,2.9440883321,2.0445014685
C,0,-0.3682802504,0.3725123964,0.4441735258
H,0,-0.7550341608,-0.2004334176,1.3000911065
H,0,-0.6915681301,1.4122599184,0.5638514808
H,0,-0.8156739275,-0.0511308856,-0.4623783969

Zero-point correction= 0.275014 (Hartree/Particle)
Thermal correction to Energy= 0.291547
Thermal correction to Enthalpy= 0.292491
Thermal correction to Gibbs Free Energy= 0.230396
Sum of electronic and zero-point Energies= -688.375197
Sum of electronic and thermal Energies= -688.358665
Sum of electronic and thermal Enthalpies= -688.357720
Sum of electronic and thermal Free Energies= -688.419816

### 2. References

(1) Gaussian 16, Revision C.01, Frisch, M. J.; Trucks, G. W.; Schlegel, H. B.; Scuseria, G. E.; Robb, M. A.; Cheeseman, J. R.; Scalmani, G.; Barone, V.; Petersson, G. A.; Nakatsuji, H.; Li, X.; Caricato, M.; Marenich, A. V.; Bloino, J.; Janesko, B. G.; Gomperts, R.; Mennucci, B.; Hratchian, H. P.; Ortiz, J. V.; Izmaylov, A. F.; Sonnenberg, J. L.; Williams-Young, D.; Ding, F.; Lipparini, F.; Egidi, F.; Goings, J.; Peng, B.; Petrone, A.; Henderson, T.; Ranasinghe, D.; Zakrzewski, V. G.; Gao, J.; Rega, N.; Zheng, G.; Liang, W.; Hada, M.; Ehara, M.; Toyota, K.; Fukuda, R.; Hasegawa, J.; Ishida, M.; Nakajima, T.; Honda, Y.; Kitao, O.; Nakai, H.; Vreven, T.; Throssell, K.; Montgomery, J. A., Jr.; Peralta, J. E.; Ogliaro, F.; Bearpark, M. J.; Heyd, J. J.; Brothers, E. N.; Kudin, K. N.; Staroverov, V. N.; Keith, T. A.; Kobayashi, R.; Normand, J.; Raghavachari, K.; Rendell, A. P.; Burant, J. C.; Iyengar, S. S.; Tomasi, J.; Cossi, M.; Millam, J. M.; Klene, M.; Adamo, C.; Cammi, R.; Ochterski, J. W.; Martin, R. L.; Morokuma, K.; Farkas, O.; Foresman, J. B.; Fox, D. J. Gaussian, Inc., Wallingford CT, 2016.

(2) GaussView, Version 6, Dennington, Roy; Keith, Todd A.; Millam, John M. Semichem Inc., Shawnee Mission, KS, 2016.

(3) Saha, P.C.; Solanke, P.R.; Biswas, S.; Agarwal, V. Taylor, M.T., Designer Aromatic Cations for Photo-Induced Protein Ligation, Imaging, and Intracellular Catalysis. *BioRxiv*, **2025**, DOI: 10.1101/2025.10.13.681063.
